# Directional Proton Conductance in Bacteriorhodopsin Is Driven by Concentration Gradient, Not Affinity Gradient

**DOI:** 10.1101/2021.10.04.463074

**Authors:** Zhong Ren

## Abstract

Many microorganisms can harvest energy from sun light to establish electrochemical potential across cell membrane by pumping protons outward. Light driven proton pumping against a transmembrane gradient entails exquisite electronic and conformational reconfigurations at fs to ms time scales. However, transient molecular events along the photocycle of bacteriorhodopsin are difficult to comprehend from noisy and inconsistent electron density maps obtained from multiple experiments. A major challenge arises from the coexisting intermediate populations as a heterogenous conformational mixture continuously evolves over 13 decades in time. This study reports a meta-analysis of the recent time-resolved datasets collected by several consortia. By resolving structural heterogeneity, this in-depth analysis substantially improves the quality of the electron density maps, and provides a clear visualization of the isolated intermediates from I to M. The earliest photoproducts revealed by the deconvoluted maps suggest that a proton transfer uphill against 15 pH units is accomplished by the same physics governing the tablecloth trick. While the Schiff base is displaced at the beginning of the photoisomerization within ~30 fs, the proton stays due to its inertia. This affinity-independent early deprotonation builds up a steep proton concentration gradient that subsequently drives the directional proton conductance toward the extracellular medium. This mechanism fundamentally deviates from the widely adopted notion on multiple steps of chemical equilibrium driven by light-induced changes of proton affinity. The method of a numerical resolution of concurrent events from mixed observations is also generally applicable.

**Significance Statement:** Microorganisms can exploit solar energy to offset their cellular acidity from the environment by pumping protons outward under light illumination. The ability to transport ions across the cell membrane in response to light makes this family of small transmembrane proteins a highly desirable toolkit in development of new biotechnologies. It is important to understand how these ion pumps operate at the molecular level. This study finds that the outward proton conductance through bacteriorhodopsin, the most studied model system in the class, is driven by a steep concentration gradient of protons established in the light induced process rather than by an affinity gradient previously sought for decades.

**Summary of Revision:** This is the companion manuscript of another paper already published in PNAS Nexus (Ren, Photoinduced isomerization sampling of retinal in bacteriorhodopsin, *PNAS Nexus*, 1(3), 2022, 10.1093/pnasnexus/pgac103). The original version of this manuscript was submitted to PNAS Nexus on February 18, 2022. The manuscript was reviewed by three reviewers and the Decision Notification was received on April 5, 2022. I appealed the decision to reject the manuscript on May 28, 2022, and the appeal was accepted. A revised version of the manuscript was submitted on July 25, 2022, with an extensive response to the peer review. The editor sent the revised version and the response to peer review back to the three reviewers. Reviewer 3 declined to review the revised manuscript. The editor extended the invitation to several other scientists to review the revised manuscript. All of them declined to review. The second Decision Notification based on the opinions of Reviewers 1 and 2 was received on September 14, 2022. The revised manuscript, the supplementary materials, and all review documents are listed below in the Table of contents. Second revision is underway.

## Introduction

Bacteriorhodopsin (bR) captures photon energy to pump protons outward from the cytoplasm (CP) against the concentration gradient, thus converts light into electrochemical energy. This integral membrane protein of 28 kDa singlehandedly achieves the exact same goal of photosynthesis and cellular respiration combined. This simplicity is widely exploited in optogenetics and bioelectronics (Ashwini et al., 2017; Fenno et al., 2011; Li et al., 2018). A trimeric form of bR on the native purple membrane shares the retinal chromophore and the same protein fold of seven transmembrane helices A-G (Fig. S1), but not necessarily its quaternary structure, with large families of microbial and animal rhodopsins (Ernst et al., 2014; Kandori, 2015). An all-*trans* retinal is covalently connected to Lys216 of helix G through a protonated Schiff base (SB) pointing toward the extracellular side (EC) in the resting state, of which the double bond C_15_=N_ζ_ is also in *trans* (traditionally also noted as *anti* (McCarty, 1970)). Upon absorption of a visible photon, the retinal isomerizes efficiently and selectively to adopt the 13-*cis* configuration (Govindjee et al., 1990). However, the retinal chromophore could also isomerize thermally to 13,15-*cis* (also noted as 13-*cis,15-syn*) configuration in dark, known as dark adaptation (Harbison et al., 1984; Oesterhelt et al., 1973; Smith et al., 1984). Extensive structural events during the ultrafast photoisomerization sampling and regioselectivity were recently revealed by the companion paper of this work (Ren, 2022).

### Photocycle

It has been shown that the blue-shifted species I prior to the photoisomerization arises before 30 fs and remains in 13-*trans* instead of a near 90° configuration about C_13_=C_14_ double bond (Zhong et al., 1996). A variety of molecular events prior to the isomerization have also been detected by vibrational spectroscopy, such as torsions about C_13_=C_14_ and C_15_=N_ζ_, H-out-of-plane wagging at C_14_, and even protein responses (Diller et al., 1995; Kobayashi et al., 2001). A red-shifted intermediate species J in 13-*cis* forms around 430-500 fs (Herbst, 2002; Mathies et al., 1988; Nuss et al., 1985) and decays at 3 ps to a less red-shifted species K (Applebury et al., 1978; van den Berg et al., 1990), which lasts longer than five decades of time therefore becomes the most “stable” intermediate throughout the entire photocycle on a log-time scale. The later period of K could be separated out as an even less red-shifted species KL at 10 ns (Hage et al., 1996; Sasaki et al., 1995; Shichida et al., 1983). The blue-shifted L state emerges from K or KL at 1-2 μs and converts to strongly blue-shifted M states at ~50 μs (Lozier et al., 1975). The L → M transition has been considered as the step for the proton acceptor, the carboxylate group of Asp85, to become neutralized by receiving a proton (Fahmy et al., 1992). This established view has been largely based on peak assignments in resonance Raman (RR) and Fourier transform infrared spectroscopy (FTIR) on wildtype bR and various mutants (Braiman et al., 1988, 1991; Gerwert et al., 1990; Lewis et al., 1974). It has been widely quoted in the literature that the proton transfer from the SB to the proton accepter occurs during the M formation because M is the only deep-violet-absorbing state comparable to the absorption properties of retinal model compounds with unprotonated SB (Heyde et al., 1971). This event of SB deprotonation has been mistakenly inferred from the strong blue shift of M state, the lack of deuteration effect in C_15_=N_ζ_ stretching frequency in M, and the observed protonation of the carboxylate of Asp85 during L → M transition. Deprotonation from the SB is the key event in the mechanism of proton pumping and directional conductance, therefore the centerpiece of this work (Fig. 1 inset). This study finds that this proton transfer spans ten decades in time instead of a single event. The electron density maps of I to M are unscrambled from one another in this analysis. They clearly show extensive conformational changes leading to and developing from the isomerization of the chromophore (Fig. S2).

**Figure 1.**
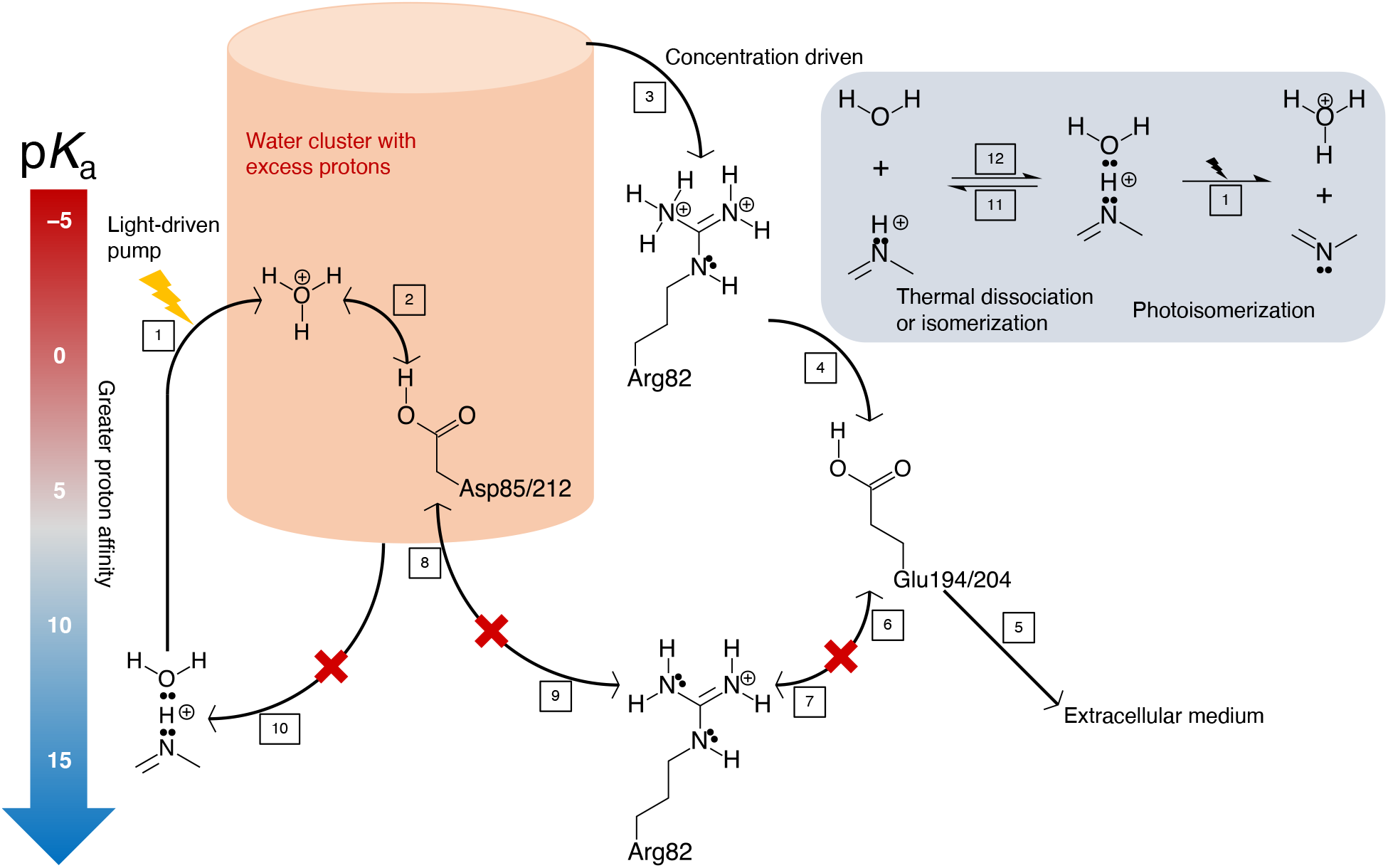
Summary of directional proton conductance in bacteriorhodopsin. The p*K*_a_ values of various proton carrying groups are marked on the left. Those chemical groups positioned lower have greater proton affinities compared to those at higher positions. Black arrows with numbered steps in boxes mark proton conductance pathways. Red crosses indicate blocked pathways. The first step is the light-driven proton pump (inset right) that achieves a high peak power *dH/dt* to gain potential energy of the proton > 15 pH units. Within the water cluster and two carboxylic acids of Asp residues in the inner EC channel, excess protons are being accumulated in the cluster, here portrayed as a bucket, by consecutive photocycles. The rest of the proton conductance is spontaneous downhill flow (Steps 2, 3, 4, and 5). As the super-acidic condition is established, a transient divalent guanidium ion drains the bucket (Step 3) and shuttles protons to carboxylic acids of Glu residues in the proton release complex (Step 4). Finally, protons are released into the EC medium (Step 5). The super-acidic water cluster in the inner EC channel next to the SB is a constant threat of proton backflow (Step 10). The EC and CP half channels are strictly separated during the inevitable pumping motions. The irregular bent helix C ensures its Thr89 follows the motion of the SB and maintains the seal between the EC and CP half channels. Four pathways connect to the monovalent guanidinium group of Arg82 inward and outward. Two uphill pathways on both sides of the guanidinium group are not possible to spontaneously conduct protons (Steps 6 and 8), which also block the other two downhill pathways since the guanidinium group is already protonated (Steps 7 and 9). The only pathway remains open if the super-acidic water cluster causes proton overflow through the divalent guanidinium ion (Step 3). However, a backflow is not possible since the EC medium cannot be super-acidic as long as sufficient water is available. (Inset) Deprotonation from the Schiff base. In the resting state, a proton is situated between the SB N_ζ_ atom and the O atom of Wat402. Two lone pairs of electrons from N and O atoms, respectively, are interacting with the proton. During thermal dissociation or isomerization to the left, the proton stays with the SB with greater probability due to its greater proton affinity (Step 11). This thermal process does not cause deprotonation from the SB. The thermal isomerization to 13,15-*cis* discharges the super-acidic inner EC channel. A thermal dissociation not involving isomerization could revert spontaneously (Step 12). On the other hand, when the SB is abruptly displaced by the photoisomerization as observed here at ~30 fs in I’ species, the proton inertia tips over the balance between these two interactions. The proton cannot overcome its association with the water and accelerate fast enough with the SB. As a result, it stays with the stationary Wat402 to form a transient hydronium ion (Step 1). This process does not revert as the photoisomerization makes the SB no longer accessible from the EC half channel ever since 500 fs. This deprotonation from the SB to the right is not governed by the proton affinities of two competing moieties based on chemical equilibrium. However, the subsequent steps are (Steps 2, 4, and 5). The excess proton of the hydronium with a p*K*_a_ of −1.7 spontaneously transfers to Asp85 to form a carboxylic acid with a p*K*_a_ of 2.2 (Step 2).

### Structure

Although the first low resolution map of bR was revealed in 1975 (Henderson and Unwin, 1975), the majority of structural information awaited technology advances until the 1990s in cryo electron microscopy (Henderson et al., 1990), electron diffraction (Grigorieff et al., 1996), and X-ray crystallography (Pebay-Peyroula et al., 1997). Finally, the atomic resolution of a bR model was achieved (Luecke et al., 1999a). The main technique to study intermediates along the photocycle used to be cryo trapping, including both light illumination of crystals at room temperature followed by rapid freezing and illumination at elevated cryo temperatures. The 13-*cis* retinal and the altered H-bond network have been observed in M state with extensive conformational changes in the anchor helix G (Luecke et al., 1999b), FG loop, EC half of helix C (Takeda et al., 2004), and an enlarged CP pocket (Sass et al., 2000). Numerous attempts have been made to capture intermediates other than M with inconsistent results (Wickstrand et al., 2015).

A new phase began with time-resolved serial crystallography conducted at X-ray free electron lasers (XFELs). The ultrashort X-ray laser pulses open the opportunity to capture transient structural species in the photocycle as short-lived as fs (Chapman, 2019; Wickstrand et al., 2019). It is equally important that serial crystallography enables photo triggering and data collection at room temperature. Authentic signals of conformational changes of greater amplitudes are expected at room temperature compared to cryo trapping. Several international consortia carried out large operations and accumulated abundant time points along the photocycle that spans more than 13 decades of time. Judged only from the extensiveness of the reported signals, time-resolved XFEL data at room temperature unfortunately do not surpass those captured by cryo trapping, which hints at much needed improvements in experimental protocols and data analysis methods. Nogly et al. captured retinal isomerization to 13-*cis* at 10 ps (Nogly et al., 2018). More importantly, they observed that the SB water moves before the isomerization and the SB itself has already displaced at the earliest time point of 49 to 406 fs, which is the key contribution to this analysis. Kovacs et al. contributed datasets at many short time delays (Kovacs et al., 2019). But these datasets contain no function-related signals other than extensive oscillations of low frequencies due to the excessive peak power of the excitation laser pulses (Miller et al., 2020; Ren, 2022). Nango et al. found an ordered water in a newly established H-bond network involving the SB in the 13-*cis* retinal pointing to the CP on ns-μs time scale, which led to their conclusion of the proton path (Nango et al., 2016). These are three major sources of data analyzed in this study (Table S1). Several synchrotron datasets at ms delays also contribute to this analysis (Weinert et al., 2019).

Structural heterogeneity is the common difficulty in both cryo trapping and time-resolved serial crystallography at room temperature. To what extend a specific species can be enriched in crystals depends on the reaction kinetics under a condition of illumination, such as wavelength and temperature. A more stable species K or M can reach a higher fractional concentration at a specific time point due to a greater ratio between the rates going into and exiting from that species. Others will be populated poorly because of their transient nature. This structural heterogeneity causes the practical difficulties in the interpretation of electron density maps and the refinement of these intermediate structures (Ren et al., 2013). An assumption in nearly all previous studies has been that each dataset, at a cryo temperature or at a time delay, is derived from a mixture of a single photoinduced species and the ground state. Therefore, the difference map reveals a pure intermediate structure. This assumption is far from the reality and leads to the more time points the more overinterpretation rather than the more overdetermination. Previous studies using cryo trapping and time-resolved crystallography could not evaluate clear electron density maps of pure intermediate species without a proper deconvolution of heterogeneous species despite the awareness of the extensive possibility of conformational heterogeneity (Ren et al., 2013; Yang et al., 2011). This work is yet another case study to test our general strategy of dynamic crystallography, that is, from the outset every observation at a time point or a temperature setting is treated as a mixture of unknown number of structural species with unknown populations (Methods). The structures of all intermediate species are also unknown except the ground state. The analytical protocol is responsible for a reliable structural interpretation by overdetermination of these unknowns from consistent observations (Ren, 2019; Ren et al., 2013; Yang et al., 2011). Inconsistent observations are identified and isolated.

The previous interpretations of the mixed signals from multiple intermediate species captured in the XFEL datasets presented a rotation of the C_13_=C_14_ double bond in the counterclockwise direction if viewed from the SB end of the retinal (Kovacs et al., 2019; Nogly et al., 2018). An experimentally determined double bond with a torsion angle of 82°, for example the entry 6g7j in Protein Data Bank (PDB), indicates that this twisted double bond conformation is more populated than any other torsion angles at hundreds of fs. Such interpretation is in direct contradiction of molecular dynamics. A double bond conformation spends most time in either energy well of *trans* or *cis*. The transition time between these energy wells is relatively short. A twisted double bond conformation cannot be significantly populated compared to the discrete populations in *trans* and *cis* unless an additional energy well emerges near the torsion angle of 90°, which no evidence supports. The gradual rotation of C_13_=C_14_ double bond presented in the previously refined structures is a misinterpretation of the mixed signals of multiple intermediate species as a single conformer. The gradual rotation does not occur in isomerization, rather it merely reflects the gradual population shift from discrete states of *trans* to *cis*. In stark contrast, perfect *trans* to perfect *cis* isomerization occurs in a single move during the transition of I → J’ at ~500 fs (Ren, 2022). This is not to say that the transitioning double bond never rotates through a twisted conformation passing 90° dihedral angle. This interpretation obeys statistical mechanics and acknowledges that a twisted conformation near 90° is never populated significantly during the isomerization, therefore cannot be observed. The refined structures in *trans* and *cis* states result in two sharp U-turns of the SB N_ζ_ atom not observed in the previous interpretations. The implication of these deconvoluted pure intermediate structures is crucial to the mechanistic understanding of the deprotonation from the SB.

### Mechanism

Understanding the mechanism of proton pumping uphill against gradient has been centered on two entangled aspects: Where are the proton uptake and release pathways? How does the retinal photoisomerization adjust the chemical groups’ ability to hold protons along these pathways? The ability of a chemical group to hold a proton is measured by its value of p*K*_a_, where p stands for −log_10_ function and *K*_a_ is the acid dissociation or ionization constant, one form of the equilibrium constant of a chemical reaction. It is clear that a proton previously retained by the SB is accepted by the carboxylate group of Asp85 in helix C and eventually released to the EC medium. Each of Arg82, Asp85, 212, Glu194, 204, and some waters could play a role (Balashov et al., 1997; Govindjee et al., 1996; Richter et al., 1996a). However, the sequence of events during this proton conductance is unclear thus often mistakenly inferred.

At the resting state of bR, the SB holds a proton tightly with a p*K*_a_ of 13.3 pointing toward the EC channel (Druckmann et al., 1982; Sheves et al., 1986). The nearby proton acceptor Asp85 bridged to the SB by a water stays quite acidic with its p*K*_a_ spanning 2 to 3 range (Balashov et al., 1996a; Brown et al., 1993; Jonas and Ebrey, 1991; Mowery et al., 1979). That water, often numbered as 402, has been identified as the centerpiece of the mechanism (Wickstrand et al., 2015). Therefore, the SB proton is associated with both N and O atoms simultaneously in ≥N_ζ_:H^+^:O<H_2_ with both interactions weaker than an ordinary single bond. One of the competing hypotheses is willing to consider that the proton is first brought to the CP side by the SB isomerization to 13-*cis*, and then conducted through a newly found water other than Wat402 on the CP side, and finally reaches Asp85 via Thr89 (Nango et al., 2016). The difficulty encountered by this hypothesis is the negative p*K*_a_ values of a hydronium ion and a protonated threonine that the proton has to go through, which means that the newfound water and Thr89 are very unlikely to accept a proton from a donor, the protonated SB, with a much higher p*K*_a_. The possibility, close to none, of this hypothesis is not understood. The other competing hypothesis relies on a H-bond network largely similar to that in the resting state to translocate the proton on the SB. The H-bond network from the SB to Wat402 to Asp85 must remain intact long after photoisomerization at ~500 fs (Herbst, 2002; Mathies et al., 1988), which requires two twisted double bonds C_13_=C_14_ and C_15_=N_ζ_ away from *trans* and *cis* configurations (2ntw) for many decades until a successful proton translocation to Asp85 (Lanyi and Schobert, 2007). These highly twisted double bonds cannot exist for that long and they are the direct consequence of the inability to unscramble a mixed observation of multiple conformations (Ren, 2022).

Despite intense studies, fundamental questions on the operating mechanism of this proton pump remain puzzling and unanswered. Q1) How does the protonated SB with a p*K*_a_ of 13.3 transfer its proton to the acidic acceptors with a p*K*_a_ of 2.2 and when? Q2) Which molecular event, the isomerization, or another event, switches the accessibility of the SB from EC to CP and how does the SB avoid reprotonation on the wrong side? Q3) How are protons conducted > 15 Å through the rollercoasting p*K*_a_ values and released into the EC medium from the protonated SB of a high p*K*_a_ through aspartic acids of low p*K*_a_, arginine (strictly speaking, its guanidinium ion) and tyrosines of high p*K*_a_, and glutamic acids of low p*K*_a_ again? Q4) How is a gradient-driven proton backflow prevented in the resting state and during the pumping motions? Q5) How does the retinal chromophore covalently linked to the protein exhibit a swing of absorption maxima nearly as wide as the entire visible spectrum from deep violet to red during its photocycle, while a free retinal compound generally absorbs from near ultraviolet (UV) to green depending on its protonation state? Q6) How does the thermal isomerization occur during dark adaptation that does not require light energy, and why specifically to 13,15-*cis*? What are the physiological functions of the dark and light adaptations, anyway? Among these questions, the greatest puzzle is the affinity-driven proton conductance. No convincing evidence can directly demonstrate a photoinduced reversal of the drastically different proton holding abilities along the EC half channel to facilitate a spontaneous proton conductance that would require an ascending order of p*K*_a_’s. The fundamental flaw in the previous line of thinking is multiple steps of equilibrium driven by proton affinities as the values of p*K*_a_ describe. A proton is spontaneously transferred from one chemical group with a less proton affinity, that is, smaller p*K*_a_, to another with a greater affinity, that is, larger p*K*_a_ (Jardetzky, 1966; Stoeckenius et al., 1979). Such proton conductance was presumed to be accomplished by the means of abundant thermal events of protonation and deprotonation, that is, equilibrium. If a slope of affinity does not satisfy the directional flow of protons, it has been commonly assumed that a photoinduced reversal of proton affinity must have occurred during the photocycle (Neutze et al., 2002; Stoeckenius, 1999). However, the p*K*_a_ of the carboxylic acid of Asp85 was even found reduced by a half unit after photoisomerization to 13-*cis*, that is, the opposite to a spontaneous proton acceptance (Balashov et al., 1996b). This study shows that equilibrium is not how the light-driven proton pump bR works, and the directional proton conductance is not driven by increasing affinities but by a concentration gradient of protons. Photoisomerization of the retinal plays a key role in the establishment of the concentration gradient.

The main findings of this work are summarized in Figs. 1 and S2. Here the ultrafast molecular events revealed from the serial crystallographic datasets demonstrate that the photochemical reaction of bR transfers a proton held by the SB to form a strong acid that barely has any ability to hold a proton. However, this proton transfer is not achieved by a chemical equilibrium therefore not governed by p*K*_a_’s. Instead, before the isomerization occurs, this proton transfer takes place due to the SB displacement within ~30 fs in a single molecular event (Fig. 1 inset). The proton’s inertia prevents it from accelerating as fast as the SB under the same physics of the tablecloth trick (Q1). Rapid photoisomerization flips the already deprotonated SB before 500 fs, thus protects the SB from being reprotonated on the wrong side (Q2). Proton conductance follows a descending order of the proton concentration gradient established in the EC half channel (Q3). The maximum proton concentration established at the end of the EC half channel causes a significant red shift of the retinal absorption (Q5) and drives the dark adaptation (Q6). A built-in seal constructed by an irregular helix reacts to the isomerization and ensures no proton leakage back to the CP channel during the motions of proton pumping (Q4). The inner EC channel is strictly surrounded by chemical groups of high p*K*_a_’s so that protons can only flow out but cannot flow back in even during dark (Q4), known as back pressure established by the function of bR itself (Westerhoff and Dancshazy, 1984). See Supplementary Information (SI) for coherent answers to these questions.

## Results and Discussion

A total of 42 datasets and 36 time points analyzed in this study are divided into two groups: 18 short delays up to 10 ps and 18 long delays ranging from ns to ms, which unfortunately leaves a gap longer than three decades not surveyed (Fig. S2 and Table S1). Difference Fourier maps among these time points and with respect to their corresponding dark datasets are calculated according to protocols previously described (Methods). Singular value decomposition (SVD) of difference maps and the subsequent Ren rotation in a multi-dimensional Euclidean space established by SVD (Ren, 2016, 2019, 2022) are performed separately to 126 difference maps of the short delays and 101 maps of the long delays. This method of “decomposition and deconvolution” results in a clean separation of electron density changes due to coexisting photoexcited species mixed in the observed datasets. The main findings are summarized in a cartoon (Fig. 1) and a Gantt chart (Fig. S2). Eight intermediate structures along the photocycle, now isolated from one another, are refined against reconstituted structure factor amplitudes (Methods; Table S2). The assignments of the previously identified intermediates I to M to these refined structures are largely based on the time stamps of the observed signals. No absorption signatures are directly associated with the X-ray diffraction data, which leaves a possibility of misassignment. However, the temporal order of these intermediates is certain.

### Earliest intermediate I’ and Schiff base deprotonation

Three decomposed map components ***U***_10_, ***U***_14_, and ***U***_17_ of the short delays contain extraordinary structural signals in terms of their extensiveness and quality (Ren, 2022). These signals originate exclusively from a few time points of Nogly et al., too few to fit the time dependency with exponentials. Instead, a spline fitting through these time points estimates the coefficients *c*_10_, *c*_14_, and *c*_17_ in the linear combination of *c*_10_***U***_10_ + *c*_14_***U***_14_ + *c*_17_***U***_17_ for the early states I, J, and their respective precursors I’, J’ (Table S2). The precursor prior to I state is located on the spline trajectory from the origin, that is, the ground state bR at the time point of 0-before the photon absorption, to the first time point of 49 to 406 fs (PDB entry 6g7i). It would be appropriate to name this location on the trajectory I’ as a precursor leading to I state judged only by the time point at ~30 fs.

Five geometric parameters are calculated along the long-chain chromophore from the refined intermediate structures to provide a good metric to evaluate and compare the transient conformations (see Fig. 2 legend for detail). Two single bonds of Lys216 are highly twisted to near 90° in the ground state (Fig. 2b 4^th^ panel), which forms a corner at C_ε_ that deviates from the flat plane of the all-*trans* retinal, protrudes inboard, and makes a van der Waals contact at 3.4 Å with Thr89 in helix C (Fig. 2b 2^nd^ panel). This contact is identified in this study as a seal between the extracellular (EC) and cytoplasmic (CP) half channels. The earliest conformational changes at the chromophore are several single bond rotations due to low energy cost while the double bonds remain rigid at tens of fs (Ren, 2022). The retinal in I’ remains in near perfect all-*trans* configuration, including the Schiff base (SB) double bond C_15_=N_ζ_ although the largest, therefore the fastest, motion observed is around the SB-C_15_=N_ζ_-C_ε_- (Fig. S3). The SB is creased into an S-shape in an expanded retinal binding pocket together with other features revealing the isomerization sampling (Ren, 2022). The slit between Thr89 and the SB increases to 4.3 Å that may break the seal between two half channels, which is inconsequential as argued below. The C_20_ methyl group swings 1.1 Å to the outboard direction, a motion previously known but at much later time (Matsui et al., 2002). These motions are rotations around the long axis of the polyene chain instead of translations (Fig. 2b bottom panel). It is important that the displaced Wat402 (Nogly et al., 2018), now a hydronium ion H_3_O^+^ (see below), remains H-bonded to both Asp residues in I’. However, its association with the SB increases to 3.3 Å that barely qualifies as a H-bond.

**Figure 2.**
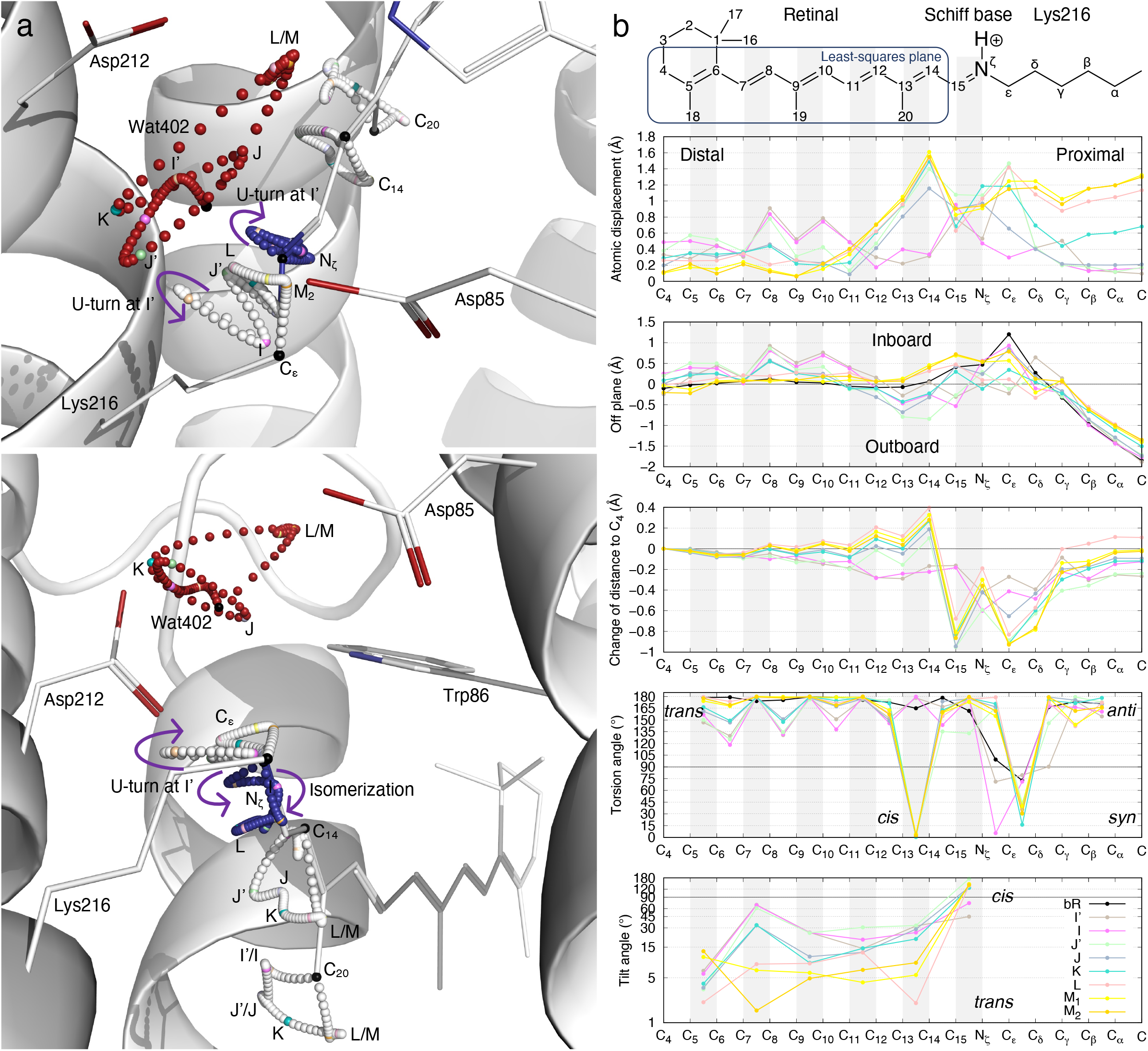
Trajectories of bacteriorhodopsin photocycle. (a) Two orthographical views of refined atomic trajectories. bR structure in the resting state is rendered in white ribbon and sticks. The photocycle trajectories of several atoms of the chromophore C_14_, C_20_, C_ε_, N_ζ_, and Wat402 are rendered as small spheres. Their positions in the ground state bR are in black. The intermediates I’, I, J’, J, K, L, M_1_, and M_2_ are colored differently. This color scheme (key in b) is consistently applied to all figures in this paper. Other small spheres along the atomic trajectories are in white for carbon, blue for nitrogen, and red for oxygen. (b) Conformational parameters calculated from the refined chromophore. The chemical structure of the chromophore on top is aligned to the horizontal axis. Double bonds are shaded in gray. 1) Atomic displacements along the polyene chain and the lysyl side chain of Lys216 with respect to the ground state structure demonstrate the overall amplitude of conformational change of the chromophore (top panel). 2) The retinal in the ground state is largely flat except C_15_ and the methyl groups of C_16_ and C_17_. Five consecutive double bonds from C_5_ to C_14_ of the retinal are largely coplanar. A plane is least-squares fitted to C_4_ through C_14_ of the resting state (boxed on the chemical structure on top). This retinal plane is largely parallel to the three-fold axis of bR trimer. Atoms along the polyene chain and the lysyl side chain of Lys216 are located on both sides of this retinal plane. The side toward the three-fold axis of the bR trimer is called inboard; the opposite side is outboard. Distances from the atoms along the long-chain chromophore to the retinal plane in the ground state are calculated for all intermediates. This parameter shows the curvature of the refined conformation. The creased retinal in the early intermediates and the inboard protruding corner at C_ε_ in the resting state are clearly shown (2^nd^ panel). 3) Distances to atom C_4_ are calculated for all refined chromophores. Changes in these distances with respect to the resting state show the shortened chromophore in I’ and I. Once isomerization to 13-*cis* occurs, the segment from C_15_ to C_δ_ around the SB becomes significantly closer to the β-ionone ring, while the distal segment of the retinal from C_14_ and beyond stretches (3^rd^ panel). 4) The torsion angles of single and double bonds quantify *anti/syn* or *trans/cis* for the ground state and all intermediates. Only a single bond can be twisted with its torsion angle near 90°. A double bond would spend very little time at a torsion angle near 90° during isomerization compared to *trans* or *cis* configurations thus not easily observed (4^th^ panel). 5) Six double bonds of the chromophore are refined to either *cis* or *trans* configurations in the ground state and all intermediates with small deviations from the ideal geometry. Therefore, each double bond is least-squares fitted with a plane. The fitted plane of each double bond tilts from that of the ground state as the conformation evolves through the intermediates and measures the local tilting of the retinal (bottom panel).

Due to the availability of the time stamp, a conformational change captured in a time-resolved experiment can be converted to the velocity of an atomic displacement that would further hint the minimum acceleration and force required to achieve the observed conformation changes at the time of the measurement. It can be shown that a force > 500 pN is required to keep the proton accelerating as fast as N_ζ_ that has been displaced by 0.7 Å within 30 fs. A force of this magnitude could unfold a-helices or even an entire protein (Su and Purohit, 2009; Takahashi et al., 2018). Accelerating the proton with the SB is not achievable given the charge on the proton and its H-bond association with Wat402. Compared to the regular single bond N-H in an amine group at a p*K*_a_ of 40, the coordination bond N_ζ_:H^+^ in the protonated SB at its p*K*_a_ of 13.3 is much weaker. However, the proton association with the SB weaker than a covalent bond is not the reason why it departs from the SB. The proton associates with both the SB N_ζ_ and O of Wat402 simultaneously in ≥N_ζ_:H^+^:O<H_2_, where the proton H^+^ does not contribute any electron in either association. The proton most likely departs from the water during a thermal dissociation because of the negative p*K*_a_ of a hydronium (Fig. 1 inset). The positive charge of the proton is delocalized in an aqueous system, thus the proton associates with waters in significantly covalent nature (Swanson and Simons, 2009). Therefore, the proton likely associates with both partners in comparable strength due to the delocalized positive charge in ≥N_ζ_:H-O^+^<H_2_. One of these two associations must break within the first 30 fs before photoisomerization. Which of the two comparable interactions survives the abrupt tare is determined by the inertia of the proton. At the beginning of the photoisomerization, the proton has to stay with the stationary partner instead of accelerating with the departing partner due to its inertia (Q1; Fig. 1 inset). Therefore, I’ captures the starting moment of a reliable deprotonation from the SB that does not depend on thermal equilibrium. Here the 0.6 Å increase from the SB to Wat402 is attributed to the separation of the proton from the SB rather than from Wat402. The hydronium 402 has been shown possible by magic angle spinning NMR (Friedrich et al., 2020). Since several water molecules have been observed in the inner EC channel, possibilities of other proton complexes also exist, such as H_5_O_2_^+^ and even more extensive clusters with excess protons and delocalized positive charges (Marx et al., 1999; Mathias and Marx, 2007; Swanson and Simons, 2009).

This conjecture here, however, is not based on an observation of the scattered X-rays by the proton. Direct evidence supporting this water cluster with excess protons reside in FTIR observations. Complex kinetic behaviors of several broad band continua at the frequencies > 1700 cm^−1^, 2200-2500 cm^−1^, and > 2700 cm^−1^ were recorded by various FTIR experiments and interpreted as protonated water clusters (Garczarek et al., 2004, 2005; Lorenz-Fonfria et al., 2017; Wang and El-Sayed, 2001), which agrees well with what will be argued below. A super-acidic water cluster harboring excess protons in the inner EC channel must be established before proton release. Reliable deprotonations from the SB in consecutive photocycles continuously overcharge the inner EC channel. The momentarily negative going signals in these FTIR continua indicate a discharge event from a super-acidic water cluster maintained by consecutive proton pumping. These FTIR continua were originally thought within the active site by the SB (Wang and El-Sayed, 2001), and later unfortunately were misassigned to a water cluster in the proton release complex on the EC surface of the protein on the basis that some of signals disappeared from the mutant E204Q (Garczarek et al., 2004). Removal of one terminal carboxylate near the exit of the EC channel (Fig. S1) in the mutant E204Q certainly could affect proton release. Shutting down one proton release pathway certainly would affect the kinetics of the upstream water cluster in the inner EC channel. Furthermore, no structural basis exists to establish a water cluster with more than one excess proton associated with the proton release complex. Therefore, the FTIR continua most likely originate from a super-acidic water cluster in the inner EC channel because a well-known proton barrier surrounds the inner EC channel (Fig. S1). Maintaining the inner EC channel super charged is required for the proper function of the proton pump (see below).

The proton acceptor Asp85 “quickly” accepts a proton from the hydronium 402, not necessarily the very proton originated from the SB in the same photocycle. The neutralization kinetics of the carboxylate of Asp85 was shown in tens of μs to ms during M formation (Braiman et al., 1988, 1991; Gerwert et al., 1990). This seems reasonable given that the p*K*_a_ difference between the aspartic acid and hydronium is < 4 pH units. See mutant functions that alter this kinetics in SI. However, the deprotonation event from the SB cannot be directly inferred from the vibrational signatures due to the neutralization of the carboxylic proton acceptors in the Asp residues. In other words, the “proton transfer” from the SB to the proton acceptor, as commonly stated in the literature, is not a single event. Instead, it spans ten decades in time with multiple asynchronous molecular events.

This extremely early deprotonation from the SB at ~30 fs even before the completion of the photoisomerization directly contradicts the consensus view long established by spectroscopy. Interestingly, it agrees with an early finding that proton movement to or from the SB does not occur during J formation although it was not an intended hint to suggest that the proton has already departed from the SB before J state (Nuss et al., 1985). Nevertheless, I have to engage this debate under the premise of photoisomerized states J’ and J later.

### Intermediate I and Schiff base inaccessibility

The retinal in I state remains in near perfect all-*trans*, including the SB. However, the major difference from the precursor is that the single bond N_ζ_-C_ε_ is now in a perfect *syn* conformation, and the anchor Lys216 has largely returned to its resting conformation (Figs. 2b 4^th^ panel and S4). The SB makes a sharp U-turn toward inboard before 500 fs (Figs. 2a and S3) and closes the slit with Thr89 to 3.8 Å and ends the breakage in the seal that is too brief to result in a proton leakage (Q3). The SB is near the apex of the first U-turn in I’, while near the apex of the second U-turn in I state immediately before isomerization (Fig. 2a). During the second U-turn, the already deprotonated SB is pointing toward inboard while Wat402, currently a hydronium, is displaced outboard but remains H-bonded to both Asp85 and 212. The SB and Wat402 are not only beyond a H-bond distance, but also in a bad geometry for proton exchange (Fig. S4d). The accessibility of the SB in I state is already very limited while the SB remains in the EC half channel but has been reoriented toward inboard despite that the isomerization has yet to occur (Q2). This ultrafast U-turn is how the SB gets rid of the proton by a mechanism of the same physics that governs the tablecloth trick (Q1). Here the tablecloth trick is not an analogy to the SB deprotonation. Both are governed by the exact same Newton mechanics: A certain mass, such as a planetary body, a dinning set, or a proton, does not accelerate fast enough unless a sufficient force is exerted on it. Holding Wat402 in place by Asp85 and 212 symmetrically from inboard and outboard, respectively, is as important as the ultrafast shaking of the SB to its successful deprotonation. See SI for mutant functions when the symmetric holding of Wat402 is disrupted. The U-turn of the SB marks the peak power, or the maximum rate of enthalpy change *dH/dt*, of the entire photochemical reaction (see Concluding Remarks). The spectrograms derived from the transient absorption or transmittance clearly support that the photocycle reaches its peak power within the first tens of fs (Kahan et al., 2007; Kobayashi et al., 2001). If a deprotonation is unsuccessful at the peak power of the photocycle, there will be no further opportunity to translocate a proton from a fair base to a decent acid uphill against 11 pH units under equilibrium. The mainstream view on this deprotonation mechanism that requires drastic changes in proton affinities of multiple partners is thermodynamically impossible. The same is true that a proton transfer is energetically impossible from the protonated SB at a p*K*_a_ of 13.3 to form a hydronium, a strong acid of a p*K*_a_ of −1.7, under equilibrium. The deprotonated SB is extremely vulnerable at this moment in I state despite its limited accessibility, as it will be quickly reprotonated if it lingers on in the EC channel to allow an equilibrium. See SI for this case in D85N mutant (Song et al., 1993). As shown below, the isomerization to a 13-*cis* retinal, described in excited state dynamics as a barrierless reaction (Birge, 1990; Gozem et al., 2017), prevents such an equilibrium in a futile photocycle. However, some of the photocycles are indeed unsuccessful as its quantum yield indicates if a reprotonation event does occur before the isomerization.

Incidentally, all previous hypotheses on the mechanism of SB deprotonation imply a formation of hydronium at one point or another (Lanyi and Schobert, 2007; Nango et al., 2016). It would require a donor group with a p*K*_a_ more negative than −1.7 to reliably form a hydronium by the means of equilibrium. None of the previous hypotheses have successfully explained how this is possible.

### Intermediate J’ and photoisomerization

J’ is the first report of a complete photoisomerization to perfect 13-*cis* at ~700 fs. This specific isomer of retinal in bR is selected from numerous possibilities during isomerization sampling by the regioselectivity of the retinal binding pocket (Ren, 2022). Near perfect 13-*cis* is successfully refined in both structures of J’ and J (Fig. 2b 4^th^ panel) that depict the process of photoisomerization started in an expanded retinal binding pocket at ~30 fs and finished in a contracted one at ~20 ps (Ren, 2022). However, the SB double bond C_15_=N_ζ_ is momentarily distorted from the *trans* configuration in the refined structure of J’ with a torsion angle of ~130°. This is the largest distortion captured for this double bond, which could, nevertheless, indicate some mixed signals not thoroughly unscrambled. The *trans* double bond C_15_=N_ζ_ is promptly restored in J to achieve 13-*cis*,15-*trans* configuration that differs from the 13,15-*cis* configuration in dark adapted retinal (Fig. 2b 4^th^ panel). The SB N_ζ_ is rotating clockwise looking from the proximal to distal direction in the entire process of the isomerization I’ → I → J’ → J (Fig. 2a). A photocycle could be wasted if reprotonation occurs prior to the isomerization at ~500 fs as the less-than-perfect quantum yield indicates that it could happen sometimes even in wildtype. The isomerization completes the accessibility switch for the SB in J’ from the EC half channel to the CP half channel (Q2), contrary to the switching mechanism based on affinity (Lórenz-Fonfría and Kandori, 2009).

The absorption property of bR during its entire photocycle has been thoroughly and repeatedly measured. If the photocycle of bR is divided into two halves I → J → K → L → M and N → O → bR, that is, a division according to the protonation state of the SB. Each half starts with a swing from a strong transitional blue shift to a strong transitional red shift and continues with less red shift to more blue shift until the strongest blue shift at the end (Fig. 3 bottom). However, the first half during unprotonated SB ends with a deep-violet-absorbing M state, while the second half during protonated SB finishes in a yellow-absorbing ground state. The apparent periodicity of similar color change repeats twice in the entire photocycle. Here, the color shift during a transition should not be confused with the color shift of each intermediate state with respect to the ground state (Fig. 3 top). Unfortunately, the periodicity of the absorption property went unnoticed, and the implication of the absorption property was mistakenly inferred despite the right conclusions were reported originally. In 1973, Oesterhelt & Hess first demonstrated that bR photoconverts between two absorption maxima 568 and 412 nm (Oesterhelt and Hess, 1973), the ground state and the later named M state, respectively. The next year, Lewis et al. concluded from their RR experiments that the SB is protonated in the ground state and unprotonated in the 412-nm-absorbing M state (Lewis et al., 1974). Ever since then, these conclusions stood the test of time. However, the SB in M state is neutral, unprotonated and the SB deprotonates during M formation are two entirely different concepts. The former does not imply the latter. The former is not being disputed here, nor any disagreement on the neutralization of the proton acceptor Aps85 in M state. The strong blue shift from 568 to 412 nm was only observed in M state, and therefore it was mistaken as the only state with a deprotonated SB since some model compounds usually absorb in near UV if they carry unprotonated SB (Heyde et al., 1971). However, a strict correlation between an absorption maximum at 412 nm and the molecular event of SB deprotonation was never established. An environmental change occurring to the same retinal isomer could cause greater blue shift than a change in protonation state. For example, the absorption of an unprotonated, neutral SB in an all-*trans* retinal *in vacuo*, that is, without interaction with solvent, peaks at 487 nm in cyan, while the absorption of a protonated SB in an all-*trans* retinal peaks at 444 nm in methanol or even peaks in near UV in ethanol, further blue shifted from the neutral SB *in vacuo* (Andersen et al., 2005; Heyde et al., 1971). Surprisingly, protonation state is a rather minor factor compared to the geometry and environment of the chromophore. Therefore, absorption maxima other than 412 nm in no way indicate a protonated SB prior to M formation. Quite the contrary, the stepwise blue shifts during J^625^ → K^610^ → KL^596^ → L^550^ → M^412^ (Fig. 3) indicate that the strong influences from the stereochemical distortion and charge redistribution over the chromophore subside gradually (see intermediate structures J, K, L, M_1_, and M_2_ below) and give way to the influence of the neutral, unprotonated SB throughout this process over 12 decades. Immediately before and after photoisomerization, I’, I, J’, and J species are highly unusual and strained in their conformations (Fig. 2b). The rapid swing from a strong transitional blue shift to a strong transitional red shift indicates the violent conformational changes at the peak power of the photocycle. The second half of the photocycle also starts with proton transfer and reisomerization that could cause some unsettling conformational strain and a local power maximum, therefore, once again a swing from blue shift to red shift. The absorption spectrograms provide a more complete picture when the photocycle reaches its peak power (Kahan et al., 2007; Kobayashi et al., 2001). The normal blue shift induced by the deprotonated SB is only a minor factor hence masked by other major factors until the chromophore is relaxed enough. Nevertheless, the second strongest blue shift of I^460^ could be partially caused by the SB deprotonation but also affected by its strained conformation. Therefore, the blue-absorbing I state marks the initial deprotonation from the SB that is going through a violent conformational struggle of isomerization. The deep-violet-absorbing M state exhibits an unprotonated SB in a relaxed 13-*cis* retinal.

**Figure 3.**
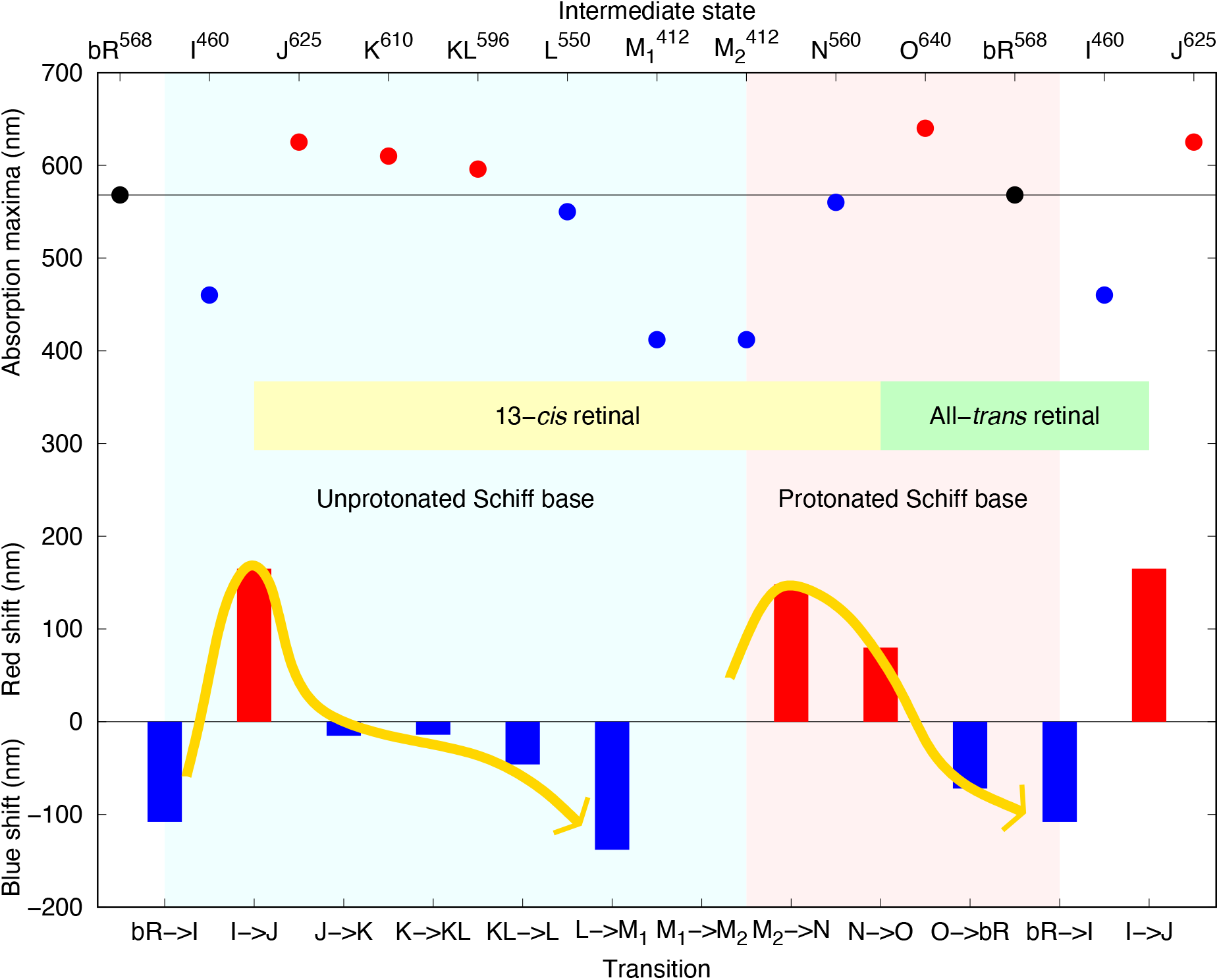
Periodic transitional color shift in two halves of bacteriorhodopsin photocycle. Absorption maxima of the intermediate states are marked as superscripts and plotted on top. With respect to the ground state, red or blue shift of each intermediate state is represented by a colored dot. Color shifts during transitions from one state to next are plotted as red and blue bars at the bottom. The periodic trends coincide with two halves of the photocycle featuring the unprotonated and protonated SB shaded in light blue and pink, respectively. Each protonation state starts with a rapid swing from a strong transitional blue shift to a strong transitional red shift and continues with a gradual change from red shift to more and more blue shift. A transitional color shift shall not be confused with the color shift of each state with respect to the absorption maximum of the ground state.

Interpretation of spectroscopic data under an erroneous structural basis could be misleading. Wat402 is no longer connected to the SB since tens fs in I’. The SB has departed from the EC half channel, the chamber that houses the resting SB in the ground state, ever since J’ in hundreds fs. It is critically important to the function of this proton pump that the SB is isolated from its home space after the completion of the accessibility switching until nearly the end of the photocycle. Arg82, Asp85, 212, and everything used to be around the SB are no longer relevant after isomerization at J’. Unfortunately, spectroscopic data were largely interpreted in its home space, or an environment similar to the ground state. No consideration could be given to the creased S-shape of the retinal gradually flattening in the expanded and contracted retinal binding pocket (Ren, 2022). Inconsistency existed throughout the history of spectroscopic interpretation. The literature on the subject of deprotonation from the SB largely evades direct evidence that supports a lasting protonated SB prior to M formation. Some of the earliest RR studies already noticed that the proton on the SB is removed before reaching M state (Lewis et al., 1974) and that the signature of deprotonation from the SB does not follow the kinetic rise of M state, but appears before 8 μs when there is no sign of M (Marcus and Lewis, 1977). To verify the peak assignment to the vibrational mode of the SB double bond C_15_=N_ζ_, parallel RR experiments were often conducted on normal and deuterated specimens. The frequency down shift of the SB double bond stretching in heavy water has been considered as an indication of H-bond strength to the SB. Much smaller difference Δ*v*_H-D_ was measured in K state compared to the ground state (Kandori et al., 2002), which indicates the already broken H-bond between Wat402 and the SB and directly supports an early deprotonation prior to K. The strongest evidence in vibrational spectroscopy that led to SB deprotonation at M formation is the lack of deuteration effect in C_15_=N_ζ_ stretching frequency only in M state. But this frequency down shift is also very small in K from deuterated samples (Kandori et al., 2002; Terner and El-Sayed, 1985). This small down shift in K when deuterated could hint not only the water has been lost but also the proton or deuteron except some remaining deuteration effect due to the still strained chromophore in K.

It is essentially unclear what exactly is the vibrational frequency of C_15_=N_ζ_ in a photoexcited retinal during its photocycle, especially, the early photocycle that is undergoing a rapid, complicated evolution through many transient states with strained chromophore far from any equilibrium states (Fig. 2). The dependency of the C_15_=N_ζ_ stretching frequency on the absorption maximum of the intermediate states was noticed long ago (Argade and Rothschild, 1983). Therefore, the frequency assignment could have been complicated by the coincidence that both a deuteronated (as opposed to protonated) C_15_=N_ζ_:D^+^ in bR and the unprotonated, neutral double bond C=N in a model compound vibrate at two indistinguishable frequencies around 1622±2 cm^−1^ (Heyde et al., 1971; Lewis et al., 1974; Stockburger et al., 1979). In addition, the vibration frequencies of N_ζ_:H^+^ and N_ζ_:D^+^ in the ground state and K were assigned by polarized FTIR experiments, and their dipolar orientation was said tilted in the ground state and parallel to the membrane in K (Kandori et al., 1998, 2002). This could be a misassignment due to the complication from another N-H bond parallel to the membrane in K since this bond N_ζ_:H^+^ or N_ζ_:D^+^ does not exist in K. Even if it does, this bond should be more perpendicular than parallel to the membrane.

### Intermediate J and traveling water or hydronium 402

The refined structure of J shows a perfect 13-*cis* and C_15_=N_ζ_ *trans* in a contracted retinal binding pocket (Ren, 2022). Due to the 13-*cis* configuration, C_20_ methyl group protrudes to a CP direction that is further tilted toward outboard from its original direction. No additional consequence is immediately obvious until the retinal flattens and returns to its original plane at later intermediates. At this moment, Trp182 and the retinal anchor Lys216 stay in their resting positions. Wat402, perhaps currently still a hydronium ion, is well defined in both Js. Contrary to the previous conclusion (Wickstrand et al., 2015), this traveling water is never disordered throughout the surveyed portion of the photocycle up to M_2_ despite its large trajectory spanning greater than 3 Å (Fig. 2a). Wat402 is observed with both strong difference electron density and its refined electron density. Its association with Asp85 and 212 remains unwavering throughout the photocycle except one increase to 4.1 Å from Asp212 during J’, the first moment after isomerization. This observation demonstrates that Wat402, a hydronium through a large part of the photocycle, is an integral part of the moving cluster that accepts the proton from the SB. Most likely, this water will remain ordered throughout the rest of the photocycle not yet surveyed.

The main cause to this contradiction is that a light induced change at any given time point consists of a mixture of multiple conformations. As the most straightforward example, the dramatic motion of Wat402 appears to be a disorder in an experimental map prior to deconvolution. An experimental map is a spatial average over the diffracting crystals and a temporal average over the X-ray exposures. In case of serial crystallography, the range of this average is an extensive pool of crystals potentially with diverse individualities (Ren et al., 2018). The only common feature among multiple species of bR intermediates is the negative density on the original location of Wat402 due to its departure. Multiple positive features at various locations are of low occupancies without a proper deconvolution. In this analysis, a linear combination of several SVD components unscrambles the mixture so that each state along the photocycle presents a pure conformational species, a time-independent structure. Is there an experimental solution to this problem of structural heterogeneity? Would a better time resolution improve an experimental difference map so that a traveling water molecule and other moving parts of the structure would always be temporally resolved? The answer is both surprisingly and conclusively no, even if, for the sake of argument, an ultimate time resolution of true zero is achievable by two zero-width impulses of pump and probe (Strictly speaking, each crystal must also be a spatial impulse, that is, zero size.). Because at no time throughout the entire photocycle, the traveling water is synchronized throughout the entire crystal, even if every retinal in the crystal absorbs a visible photon at the exact same time zero. There is no difference between crystal and solution regarding this premise in thermodynamics, which has been long recognized as Moffat laid the foundation of time-resolved crystallography (Moffat, 1989, 2001). In conclusion, a numerical resolution of structural heterogeneity, as exemplified here, is the last and the only resort (Ren et al., 2013).

### Five-decade long-lived intermediate K and global change in intermediate L

This work continues with another joint analysis of the time-resolved datasets at long delays > 10 ns collected mainly by Nango et al. with contributions from Nogly et al., Kovacs et al., and Weinert et al. (Table S1). These datasets at the long delays are relatively noisier and contain systematic errors from one experiment to another. The analytical procedure of SVD and the subsequent Ren rotation (Ren, 2016, 2019, 2022) is capable of isolating time-resolved signals from these error and noise components. SVD of 101 difference Fourier maps identifies 19 major components (Fig. S5a). Four of them contain outstanding signals of structural changes (Figs. 4 and S6). Two others also carry structural signals but without a clear time dependency (Figs. S7 and S8). The number of datasets at long delays warrants an exponential fitting to the SVD coefficients (Fig. S9), which models the reaction scheme of the photocycle better (Schmidt et al., 2010) compared to the spline fitting applied to the limited number of short delays (Ren, 2022). The time constants obtained from the exponential fitting agree well with the transition times between intermediates K, L, M_1_, and M_2_ known from spectroscopy (Ernst et al., 2014). Coefficients for relatively pure states are estimated and the refinement protocol based on reconstituted difference maps again produce atomic coordinates of these states (Methods).

**Figure 4.**
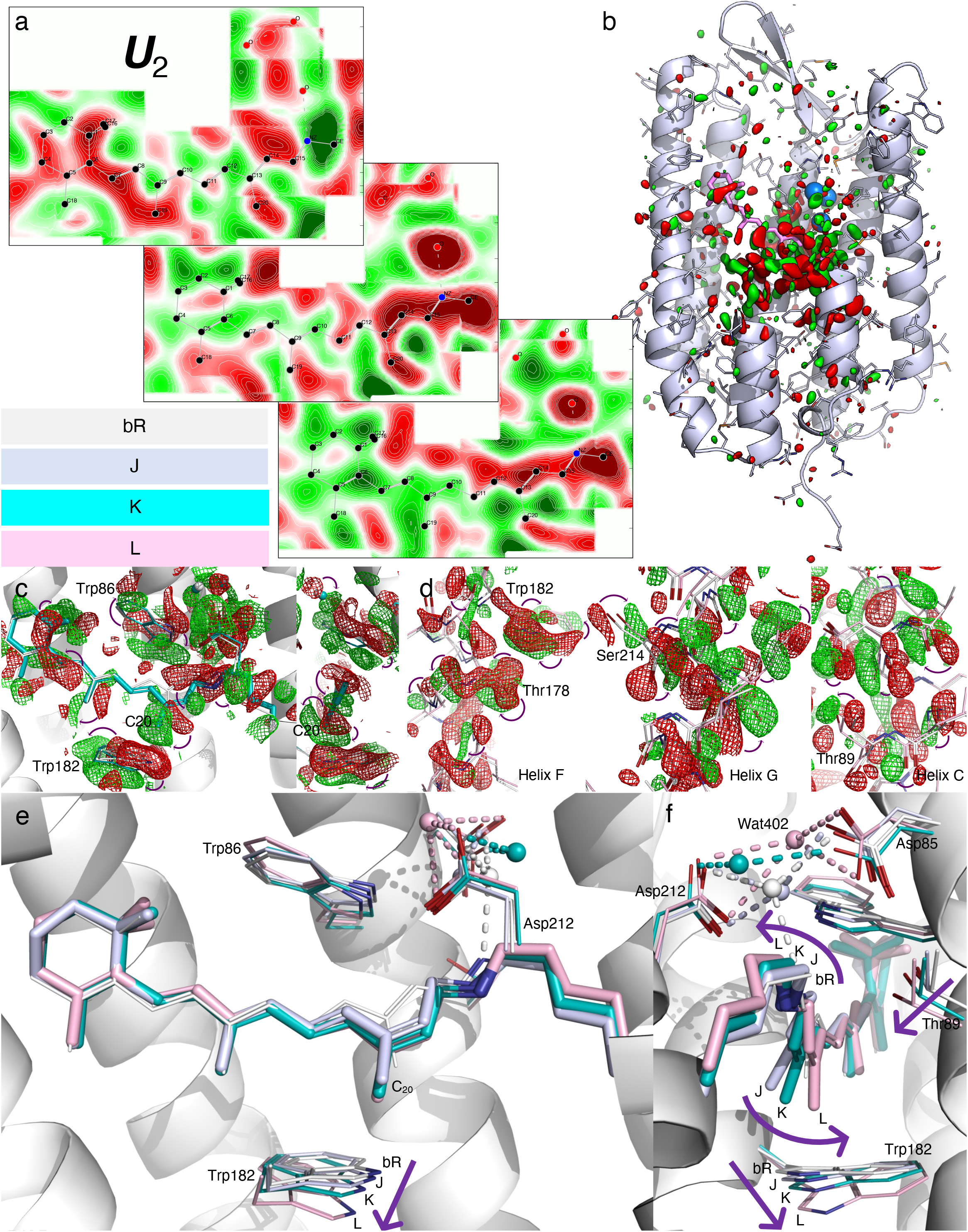
Intermediates J, K, and L. (a) Cross sections of component map ***U***_2_ of the long delays > 10 ns. The middle cross section is an integration ±0.2 Å around the surface through the retinal. The top cross section is an integration 1.4-1.8 Å outboard from the retinal surface and the bottom is an integration 0.6-1 Å inboard. Green and red indicate electron density gain and loss, respectively. Stronger signals are around the proximal segment of the retinal, including Wat402. Positive and negative sheets of densities flank the distal segment of the retinal from inboard and outboard sides, respectively. This component is nearly constant in all long-delay difference maps with a dark dataset as reference (Fig. S5). (b) Overview of the component map ***U***_2_ of the long delays contoured at ±3*σ*. The extraordinary signal-to-noise level is shown as the association with specific elements of the structure. Some parts of the structure are omitted to show the interior. (c) Two orthographical views of the reconstituted difference map K - bR from ***U***_1_, ***U***_2_, and ***U***_3_ of the long delays (Fig. S6abc). The map is contoured at ±2.5*σ* in green and red mesh, respectively. Atomic displacements are indicated by the arrows marking the negative and positive pairs of densities. (d) Reconstituted difference map L - bR from ***U***_2_, ***U***_3_, and ***U***_6_ of the long delays (Fig. S6bcd). The map is contoured at ±3.5*σ* in green and red mesh, respectively. The strongest signals are concentrated on the irregularities of helices F, G, and C (Fig. S12). (e and f) Two orthographical views of the refined retinal in J, K, and L compared with the resting state in white. The retinal is less creased in K and flattened in L compared to the previous intermediates (Fig. 2b 2^nd^ panel). C_20_ methyl group is less and less tilted from its orientation in the resting state (Fig. 2b bottom panel). Wat402 with good electron density (Figs. S10 and S11) remains its H-bonds with both Asp85 and 212 in both K and L. Thr89 follows the motion of the SB toward outboard.

Emerged from J around 10 ps, K lasts more than five decades into μs, which suggests that it is a struggle to achieve the next state L over a significant energy barrier. The creased retinal ever since I’ at the very first time point (Ren, 2022) is relaxing in K (blue; Figs. 4ef and S10), and results in three consecutive transitional blue shifts giving way to the influence of the deprotonated SB (Fig. 3). This slow and difficult relaxation could be driven in part by the energy stored in the creased 13-*cis* retinal and leads to global changes over the entire molecule. First, C_20_ methyl group is tilting less at 20° from its resting orientation (Fig. 2b bottom panel). Although protruded toward the CP direction due to the 13-*cis* configuration, the tilted C_20_ in J is not pointing to Trp182 directly (Fig. 4ef). Therefore, Trp182 is little affected in J. As C_20_ swings back in K, it starts to push Trp182 toward the CP and causes displacements in the main chain. The positive density that indicates the position of C_20_ in the reconstituted difference map K - bR forms an arc shape centered on the polyene chain (Fig. 4c). This observation strongly suggests that K is a long process, rather than a discrete state, with C_20_ distributed along the arc leading toward the next state L, in which the methyl group is fully erected (Fig. 4ef). Second, the entire chromophore in K, except the most distal segment beyond C_8_, is in a flat conformation with the double bond C_13_=C_14_ in *cis* and the single bond C_ε_-C_δ_ in *syn*. 12 consecutive torsion angles from C_8_ to C_α_ deviate no more than 20° from *trans/cis* or *anti/syn*. The root-mean-squared deviation (rmsd) is 8.6° (Fig. 2b 4^th^ panel). To achieve such energetically favorable conformation in the chromophore, the retinal anchor Lys216 is driven toward the EC direction, which causes a significant shift of the main chain in the vicinity. Third, again due to the 13-*cis* configuration, Trp86 on the EC side moves with the proximal segment of the retinal toward the CP direction, which also causes movement in the main chain. As a result, the H-bond Asp85O_δ_-Thr89O_γ_ in K is strengthened and refined to the shortest distance of 2.5 Å among all intermediates and the ground state. This strengthened H-bond during the long lifetime of K was previously detected by FTIR (Kandori et al., 1999). All three developments above are the coherent progresses in a long and hard work toward the goal to establish the state L with global consequences.

After five decades of time since tens of ps, the large energy barrier is finally overcome around several μs as described by the component map ***U***_3_ (Fig. S6c) in the global transition of K → L. This transition is modeled here at a time constant of 2 μs (Fig. S9). The potential energy in the creased retinal established within tens of fs after photon absorption is spent to trigger and drive this transition, at least in part. Other sources of potential energy, such as that stored in the bent helix C (Fig. S12a), could also contribute to the global transition. Strong signals in the reconstituted difference map L - bR (Fig. 4d) support the refined structure of L (pink; Fig. S11). The refined structure of L shows that the 13-*cis* retinal is finally flattened as its potential energy is consumed. 12 consecutive torsion angles from C_5_ to C_δ_, including the SB, deviate from *trans/cis* or *syn/anti* no more than 23° with an rmsd of 10° (Fig. 2b 4^th^ panel). As a small cost, a couple of single bonds of Lys216 have to deviate from the perfect *syn* or *anti* conformation like those in the flat all-*trans* retinal in the resting state. The plane of the flattened 13-*cis* retinal has largely returned to the original plane of the all-*trans* retinal (Fig. 2b 2^nd^ and bottom panels). The distorted single bonds along the S-shaped polyene chain ever since tens of fs in I’ (Fig. 2b 4^th^ panel) support the isomerization sampling caused by a charge separation (Nogly et al., 2018; Ren, 2022). This conformational distortion and charge distribution could be major factors that influence absorption property. The fully flattened retinal suggests that the separate charge has recombined in L. As a result, L state continues the blue-shifting trend and achieves a significant step to green-absorbing at 550 nm from the last state KL (Fig. 3). Nevertheless, the absorption maximum of L state is still long way from where an unprotonated SB in a free retinal compound should absorb in near UV, which in no way implies that the SB is holding on to its proton. C_20_ methyl group is pointing directly to Trp182, which causes helix F to displace (Figs. 4d and S12b). Leu93 in helix C is also affected by the protruding C_20_ methyl group back into the original retinal plane. On the other hand, the fully erected C_20_ displaces the anchor helix G with equally strong signals (Figs. 4d and S12c).

The L state represents the end goal of the global conformational changes due to the retinal isomerization. Additional changes after L are no longer global (see below). C_δ_-C_ε_ of the anchor moves outboard and could break the seal with Thr89 once again. Last time, a potential breakage occurs in I’ at tens of fs that is too brief for any proton leak (Fig. 2b 2^nd^ panel). This time, the ns-μs time scale allows the protein to react. It is clearly observed that Thr89 and helix C always follow C_δ_-C_ε_ outboard in J, K, and L and maintain a distance no more than 3.6 Å because of its tendency to be straightened (Figs. 4f and S12a). This motion was also captured previously in cryo-trapped intermediates (Royant et al., 2000). Therefore, the seal between two half channels is kept intact (Q4). Here the mechanism of these motions of helices is attributed to several irregularities in these helices (Fig. S12), the kinked helix C (Grigorieff et al., 1996), the stretched helix F, and the π helical segment in G (Pebay-Peyroula et al., 1997). Contrary to the previous belief (Luecke et al., 1999a), it is our general conclusion also from studies of other protein mechanisms that irregularities in secondary structures not merely fulfill some structural requirements but offer a wide variety of mechanical properties to achieve the functions of protein nanodevices (Ren et al., 2016; Shin et al., 2019). These motions of the helical segments collectively result in a tighter EC channel and an open CP channel in L (Fig. S13).

### Intermediates M_1_, M_2_ and proton release

An increase of the component map ***U***_1_ separating the long delays > 10 μs from the earlier ones is modeled as the L → M transition at a time constant of 27 μs. The M state could be split into two by an increase of the component map ***U***_6_ at a time constant of 290 μs (Fig. S9). Signals in both ***U***_1_ and ***U***_6_ are distributed over the EC half of the molecule instead of globally (Fig. S6ad). Two similar structures M_1_ and M_2_ (yellow and orange) are refined according to the reconstituted difference maps M_1_ - bR and M_2_ - bR, respectively (Figs. S14 and S15). Since the L state, the SB is moving inboard mainly by twisting the torsion angle around N_ζ_-C_ε_. C_ε_ in M_1_ and M_2_ has moved 0.6 and 0.8 Å, respectively, from its position in L. It seems that the conformation continues to make local adjustments while it is approaching a point poised to isomerize back to all-*trans* (Fig. S16c). Two M substates achieve another significant step of blue shift to reach deep-violet-absorbing at 412 nm (Fig. 3). This is the most blue-shifted state that an unprotonated SB can attain when it is tethered to the protein unlike an unprotonated SB in a free retinal compound absorbing in near UV (Q5) (Heyde et al., 1971). In the meanwhile, Thr89 and helix C are being pushed back toward inboard, which restores the kink of helix C in the resting state (Fig. S16abc). Thr89 is kept 3.1 Å away from the SB in L, M_1_, and M_2_ constantly so that the seal is well maintained between two half channels (Q4). Compared to the slightly longer distance between Thr89 and the SB at 3.6 Å in J and K, it is obvious that Thr89 and helix C react to the retreating SB during J and K, but the SB is actively pushing Thr89 and helix C from L to M (Fig. S16c). As discussed below, during active pumping, a drastic proton concentration gradient is present across this seal that divides two half channels. A well-maintained seal is crucial to the efficiency of the pump. Various mutants with substitutions for Thr89 that could result in a proton leak are discussed in SI. They retain ~2/3 of the pumping activity compared to the wildtype (Marti et al., 1991).

The new development in the retinal displaces helix G further toward the EC direction as previously observed (Takeda et al., 2004). On the other hand, the stretched helix F in L springs back in M and becomes shorter than that in the resting state (Fig. S12b). The EC segment of helix F turns around to move outboard since L (Fig. 4d). However, the EC segments of helices C and G squeeze even tighter, while Arg82 swings out of the way as observed previously (Luecke et al., 2000; Sass et al., 2000). These motions over the EC half of the molecule from tens of μs to ms are directly related to the pathway of the proton release (Q3). Residues in the EC half channel are not arranged in a monotonic order of their p*K*_a_ values. Quite the contrary, acidic residues Asp85 and 212 with the smallest p*K*_a_ values are at the end of the half channel next to the SB with the greatest p*K*_a_. Another layer of acidic residues Glu9, 194, and 204 are at the EC surface. It is puzzling that Arg82, Tyr57, and Tyr83 are located between these two acidic layers (Figs. 5 and S1). It had been depicted multiple times by similar illustrations in the past that a high p*K*_a_ layer is sandwiched between two acidic layers with low p*K*_a_ values in the EC half channel. Yet, the question remains: Why are the residues with high and low p*K*_a_ values arranged in the way they are? How are protons conducted through the proton barrier of the guanidinium group of Arg82 that is already protonated and the phenol hydroxyl groups of the Tyr residues that cannot be protonated anymore? No evidence shows that such rollercoasting p*K*_a_ values would change in any meaningful way by the light-induced conformations (Brown et al., 1993). Therefore, the previous proposal on an affinity-driven proton conductance is fundamentally flawed.

**Figure 5.**
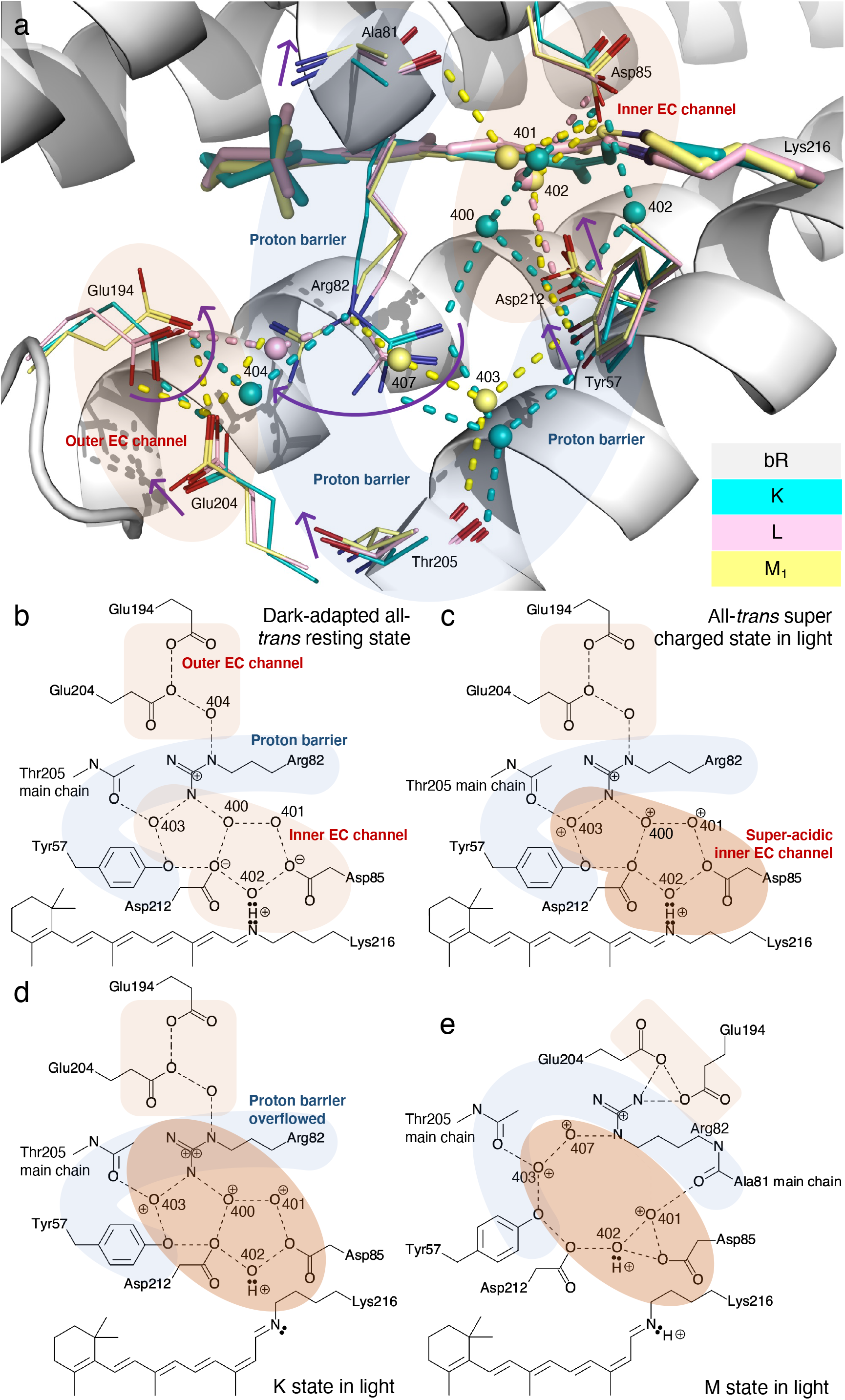
H-bond network in EC half channel. The EC half channel is decorated with acidic residues, such as Asp85 and 212 at the end, called inner EC channel shaded in pink, and Glu9, 194, and 204 at the EC surface, called outer EC channel also shaded in pink. Arg82, Tyr57, and 83 are located between these two layers that form a proton barrier shaded in blue (Fig. S1). (a) The refined structures of K in blue, L in pink, and M_1_in yellow are rendered in the framework of the resting state in white. Several arrows mark the motions of K → L → M_1_. (b) Flattened network of dark-adapted resting state. Dashed lines mark H-bonds and salt bridges. Hydrogens, except the proton on the SB, are omitted due to the ambiguity in their associations with other atoms. 400 numbers are waters. A dual-pentagon of H-bond network is characteristic in the resting state. The guanidinium group of Arg82 is already protonated with a positive charge at a neutral pH. The phenol hydroxyl groups of Tyr cannot be protonated anymore. These residues form a proton barrier shaded in blue between the inner aspartic acids and the outer glutamic acids shaded in pink. The H-bond network in the EC half channel connects the inner acidic residues with the outer ones through the proton barrier of Arg82, Tyr57, and the main chain carbonyls of Ala81 and Thr205. The inner EC channel is discharged. No proton conductance can occur in this resting state. (c) Flattened network in an all-*trans* state in light. This beginning state of each photocycle exists after light adaptation during continuous pumping activity in light. The retinal has returned to its all-*trans* configuration ready to be excited for the next photocycle. Crystallographic structures in the ground state most likely depict this state. The absorption maximum in the yellow indicates the inner EC channel is charged with excess protons. At the maximum capacity, the carboxylic acids of Aps residues are neutralized, and several waters could have also been charged to become hydronium ions as shaded in brown. Although a six-proton capacity is observed in this state (two carboxylic acids and four hydronium ions 400-403), it is unknown how many excess protons are required to drive the proton conductance. (d) Flattened network of K state in light. The dual-pentagon H-bond network is disrupted therefore not observed in its intact form in all excited states except in K. Proton conductance through the proton barrier is driven by excess protons in the super-acidic inner EC channel shaded in brown. The guanidinium group of Arg82 is protonated for a second time under the extraordinary proton concentration. (e) Flattened network of M state in light. The guanidinium group swings toward EC and makes direct contact with the glutamic acids. The second extra charge on the guanidinium group is conducted to the EC medium through the proton release complex driven by affinity gradient (Fig. 1). However, the guanidinium group retains one extra proton.

Here a concentration-driven, instead of affinity-driven, proton conductance is hypothesized. Proton release becomes synchronized to the photocycle only when a super-acidic inner EC channel is established by the first a few photocycles after a long dark period that pump protons into the inner EC channel, but no proton release is possible yet. This startup process is a part of the light adaptation. It was shown that the time constant for dark adaptation is 14 min (Fahr and Bamberg, 1982). Other experiments demonstrated that dark adaptation is a process depending on temperature, pH, and lipid environment, nevertheless, a process far slower than the photocycle of bR (Dencher et al., 1983; Ohno et al., 1977). This can be explained as the steep electrochemical gradient across the SB linkage drives the thermal dual-isomerization specifically to 13,15-*cis* without the need of light (Q6). It is yet unknown whether the isomerizations at C_13_=C_14_ and C_15_=N_ζ_ occur simultaneously or consecutively. The thermal isomerization could discharge the inner EC channel by carrying a proton to the CP side without the ultrafast photophysical process (Fig. 1 inset). The dark-adapted state consists of 1/2 to 2/3 of the chromophores in 13,15-*cis* (Fahr and Bamberg, 1982; Harbison et al., 1984; Maeda et al., 1977; Scherrer et al., 1989), which seems to hint that discharging pathways other than the proton-gradient-driven thermal isomerization also play a role. Those alternative pathways discharge more excess protons at higher temperature due to greater thermal motions so that the fraction of 13,15-*cis* chromophore in dark-adapted state decreases (Scherrer et al., 1989). Despite the exquisite arrangement of the proton barrier and the irregular, bent helix C constantly pressing its Thr89 on the moving part of the chromophore, the super-acidic proton pool cannot be maintained indefinitely in dark. The slow discharge at least four decades slower than the photocycle by the acid driven isomerization may not be an unintended defect of the proton pump. Quite the opposite, the super acidic condition deep inside a protein could be harmful and needs to be discharged while the pump is not in use. I suggest that this could be a form of photoprotection (Q6).

After a long dark period, a light adaptation is necessary for the normal function of bR. This startup includes not only a reset of the chromophore to all-*trans* configuration but also the buildup of a proton concentration at the end of the EC half channel during the first a few photocycles without any proton release. Multiple groups in the inner EC channel, including various waters, could be all protonated after consecutive photocycles, that is, a puddle of protons establishes a super-acidic inner EC channel as long as the seal of Thr89 is well maintained and the discharging processes are slow enough. This is possible because the proton pumping step is a photophysical process instead of a chemical equilibrium (see above). Although the capacity for the excess protons is observed as six or more (Fig. 5cd), it is unknown how many excess protons are necessary to drive proton release (see below). That is to say, two states sharing the same or very similar all-*trans* retinal are differentiated - the dark-adapted state with all-*trans* SB and a discharged inner EC channel (Fig. 5b) and the beginning state of the photocycle in light with a super charged inner EC channel (Fig. 5c). The structure and function of the all-*trans* light state is perhaps the one usually observed by crystallography and spectroscopy. The existence of the super-acidic inner EC channel is implied from various spectroscopic data. On the other hand, the dark-adapted and discharged state is harder to observe.

Although the first a few photocycles are productive during light adaptation, they are not capable of releasing any proton before an established proton concentration gradient. An established super-acidic inner EC channel triggers a concentration-driven proton conductance (Fig. 5d). Proton release from the wildtype bR was observed before proton uptake as if it is coupled with, if not faster than, the protonation of the proton acceptor Asp85 (Braiman et al., 1988, 1991; Gerwert et al., 1990; Zimanyi et al., 1992), which strongly suggests that the newly pumped proton is not the one released to the EC medium in the same photocycle. Here the concentration-driven proton conductance argues that the newly pumped proton causes a proton overflow from the already super-acidic inner EC channel (Figs. 1 and 5de). Therefore, the observed “early release” is actually a very late release during the next photocycle or several cycles after the very proton has been pumped into the inner EC channel (Fig. 5e). The existence of a super-acidic inner EC channel during active proton pumping and releasing is also evidenced by the most red-absorbing O state at 640 nm in the entire photocycle, the first state of all-*trans* retinal that has just brought an additional proton into the proton pool on the EC side (Fig. 3). A free retinal compound with a SB protonated by anilines absorbs as far as green light < 515 nm (Heyde et al., 1971). The extra red shift of O state indicates a very acidic condition compared to its predecessor N absorbing at 560 nm and the successor back to the beginning of the photocycle absorbing at 568 nm. The last transitional blue shift of O^640^ → bR^568^ caused by a discharge of the inner EC channel is consistent with the consensus view that the proton transfer from Asp85 or 212 to the proton release complex occurs at the end of the photocycle (Richter et al., 1996b). Because of this super-acidic inner EC channel necessary for proton release, the range of absorption maxima during the photocycle is extended from deep violet to red, nearly as wide as the entire visible spectrum (Q5).

The inner EC channel has been called a proton cage at the active site of bR (Friedrich et al., 2020), which reflects the same fact of the proton barrier. Here I argue that the super-acidic inner EC channel causes a second protonation of the guanidinium group because of three electron-rich nitrogens (Fig. 5d). The observation to corroborate this speculation is the motions of Arg82 in helix C, Glu194 at the end of helix F, and Glu204 in helix G during K → L → M_1_→ M_2_. The monovalent guanidinium cation during the early photocycle has no salt bridge to Glu194 and 204 until M state. It appears that Asp85 and 212 are continuously protonated by proton pumping thus neutralized in the inner EC channel, or consecutively so as previously suggested (Dioumaev et al., 1999; Zscherp et al., 2001). Several hydronium ions make the inner EC channel positively charged. The observed kinetics of the negative going FTIR continua explain that the excess protons harbored by the water cluster in the inner EC channel are being discharged by the observed motions of Arg82 during M (Garczarek et al., 2004, 2005; Lorenz-Fonfria et al., 2017; Wang and El-Sayed, 2001). An extra positive charge on the short-lived divalent guanidinium drives itself away from the inner EC channel and toward the negative charged Glu residues in the outer EC channel (Fig. 5a). Driven by the electrostatic force, the EC segments of helices C and G move closer to each other in the refined M structures, where Arg82 is sandwiched in between (Fig. 5). Molecular dynamics simulations found that the orientation of Arg82 is correlated with the charge difference between the inner and outer acids (Ge and Gunner, 2016). The contact between Glu194, 204, and the guanidinium discharges the second extra proton to the outer EC channel and the EC medium (Fig. 5e). A monovalent guanidinium cation is unable to discharge its proton to the carboxylates of the Glu residues due to their low proton affinity (Fig. 1). Therefore, Arg82 is a unidirectional proton valve that conducts protons outward, but only under sufficient proton concentration in the inner EC channel. After dark adaptation, Arg82 prevents a concentration-driven proton leak back into the cell, because the EC medium cannot become super acidic as long as sufficient water is available (Fig. 1). See SI for mutant functions when the proton barrier is breached by substitutes for Arg82.

## Concluding Remarks

Under the guidance of Jardetzky’s allostery model of transmembrane pump (Jardetzky, 1966) and the common line of thinking led by Stoeckenius (Stoeckenius et al., 1979), evidences were sought in the past half century to explain how the absorbed photon energy is used to alter the proton affinity landscape along its pathway so that protons can flow in one direction along an ascending order of p*K*_a_ values (Braiman et al., 1996; Brown and Lanyi, 1996; Govindjee et al., 1994; Neutze et al., 2002). Neither a hydraulic powered noria or an animal powered sakia in the old time, nor an electric powered fluid pump in the modern days, works under such an operating principle that requires energy for a reconstruction of the landscape. Quite the contrary, the landscape is a given constant and energy is absorbed by the to-be-pumped substance to gain its potential energy, such as water is lifted by a noria or pressurized by an electric pump. The energized substance will then flow freely along the largely constant landscape governed by thermodynamics. In the meanwhile, it is critical to prevent a backflow along a far steeper gradient than the forward-driving gradient. The molecular pump bacteriorhodopsin (bR) for protons works under the exact same principle that fundamentally differs from what Jardetzky and Stoeckenius once imagined. Other transmembrane molecular pumps and transporters likely work in the same way at this philosophical level disregarding any molecular detail. Alteration of affinities to the pumped substance cannot be an effective strategy because it is against the chemical nature along the translocation pathway and therefore energetically difficult to achieve. For example, the p*K*_a_ values differ more than 11 pH units at the resting state between the protonated Schiff base (SB) and the carboxylic acids, the proton acceptors. That is to say, a photoinduced reduction of one million times to the proton affinity of the protonated SB must occur simultaneously with an increase of one million times to the proton affinity of the carboxylic acids. Even if these changes in proton affinity are achievable, a significant fraction of the SB will remain protonated at a prolonged equilibrium, that is, failed attempts of proton transfers. The proton levee, consisting of a few tyrosine residues, main chain carbonyls, and most importantly, Arg82, is strategically constructed around the inner EC channel to prevent proton backflow during a power blackout. Their high p*K*_a_ values are not meant to be changed even though small changes could be detected. In this study, the assumption of an affinity-driven proton conductance is replaced by a mechanism of concentration-driven proton conductance that is not necessarily synchronous to the early photocycles. The photophysical nature of the deprotonation from the SB guarantees a proton concentration high enough to drive the directional proton conductance.

All chemical reactions involve energy transfer or conversion, known as enthalpy change *DH*. This flow of energy in a unit time is the temporal rate of enthalpy change *dH/dt* or power as a function of time. This function is usually far from a constant during the course of a given chemical reaction. The peak power of a chemical reaction could be orders of magnitude greater than its average power. This is particularly true for each individual molecular event compared to a molecular ensemble. Consider the photochemical reaction of the retinal isomerization in bR. Its peak power is reached at the U-turn of the SB prior to photoisomerization, which accomplishes the initial deprotonation of each photocycle. The protein structure of bR prevents a photochemical reaction from reaching its equilibrium as the photoisomerization of the retinal switches the accessibility of the SB to a different compartment. This is usually unachievable in chemical reactions of small molecules.

## Acknowledgements

This work is supported in part by the grant R01EY024363 from National Institutes of Health. The following database and software are used in this work: CCP4 (ccp4.ac.uk), Coot (www2.mrc-lmb.cam.ac.uk/Personal/pemsley/coot), dynamiX™ (Renz Research, Inc.), gnuplot (gnuplot.info), PDB (rcsb.org), PHENIX (phenix-online.org), PyMOL (pymol.org), Python (python.org), and SciPy (scipy.org).

## Data Availability

The refined structures of I’, I, J’, J, K, L, M_1_, and M_2_ are deposited to PDB-Dev (pdb-dev.wwpdb.org) with accession codes PDBDEV_00000129, 138, 139, 140, 144, 145, 146, and 147, respectively.

## References

Adams, P.D., Afonine, P.V., Bunkóczi, G., Chen, V.B., Davis, I.W., Echols, N., Headd, J.J., Hung, L.-W., Kapral, G.J., Grosse-Kunstleve, R.W., et al. (2010). PHENIX: a comprehensive Python-based system for macromolecular structure solution. Acta Crystallogr. D Biol. Crystallogr. D66, 213–221. https://doi.org/10.1107/S0907444909052925.

Andersen, L.H., Nielsen, I.B., Kristensen, M.B., El Ghazaly, M.O.A., Haacke, S., Nielsen, M.B., and Petersen, M.Å. (2005). Absorption of Schiff-base retinal chromophores in vacuo. J. Am. Chem. Soc. 127, 12347–12350. https://doi.org/10.1021/ja051638j.

Applebury, M.L., Peters, K.S., and Rentzepis, P.M. (1978). Primary intermediates in the photochemical cycle of bacteriorhodopsin. Biophys. J. 23, 375–382. https://doi.org/10.1016/S0006-3495(78)85456-3.

Argade, P.V., and Rothschild, K.J. (1983). Quantitative analysis of resonance Raman spectra of purple membrane from Halobacterium halobium: L550 intermediate. Biochemistry 22, 3460–3466. https://doi.org/10.1021/bi00283a024.

Ashwini, R., Vijayanand, S., and Hemapriya, J. (2017). Photonic potential of haloarchaeal pigment bacteriorhodopsin for future electronics: A review. Curr. Microbiol. 74, 996–1002. https://doi.org/10.1007/s00284-017-1271-5.

Balashov, S.P., Govindjee, R., Kono, M., Imasheva, E., Lukashev, E., Ebrey, T.G., Crouch, R.K., Menick, D.R., and Feng, Y. (1993). Effect of the arginine-82 to alanine mutation in bacteriorhodopsin on dark adaptation, proton release, and the photochemical cycle. Biochemistry 32, 10331–10343. https://doi.org/10.1021/bi00090a008.

Balashov, S.P., Imasheva, E.S., Govindjee, R., and Ebrey, T.G. (1996a). Titration of aspartate-85 in bacteriorhodopsin: what it says about chromophore isomerization and proton release. Biophys. J. 70, 473–481. https://doi.org/10.1016/S0006-3495(96)79591-7.

Balashov, S.P., Imasheva, E.S., Govindjee, R., Sheves, M., and Ebrey, T.G. (1996b). Evidence that aspartate-85 has a higher pKa in all-trans than in 13-cis bacteriorhodopsin. Biophys. J. 71, 1973–1984. https://doi.org/10.1016/S0006-3495(96)79395-5.

Balashov, S.P., Imasheva, E.S., Ebrey, T.G., Chen, N., Menick, D.R., and Crouch, R.K. (1997). Glutamate-194 to cysteine mutation inhibits fast light-induced proton release in bacteriorhodopsin. Biochemistry 36, 8671–8676. https://doi.org/10.1021/bi970744y.

van den Berg, R., Du-Jeon-Jang, Bitting, H.C., and El-Sayed, M.A. (1990). Subpicosecond resonance Raman spectra of the early intermediates in the photocycle of bacteriorhodopsin. Biophys. J. 58, 135–141. https://doi.org/10.1016/S0006-3495(90)82359-6.

Berman, H.M., Kleywegt, G.J., Nakamura, H., and Markley, J.L. (2012). The Protein Data Bank at 40: Reflecting on the past to prepare for the future. Structure 20, 391–396. https://doi.org/10.1016/j.str.2012.01.010.

Birge, R.R. (1990). Photophysics and molecular electronic applications of the rhodopsins. Annu. Rev. Phys. Chem. 41, 683–733. https://doi.org/10.1146/annurev.pc.41.100190.003343.

Braiman, M.S., Mogi, T., Marti, T., Stern, L.J., Khorana, H.G., and Rothschild, K.J. (1988). Vibrational spectroscopy of bacteriorhodopsin mutants: light-driven proton transport involves protonation changes of aspartic acid residues 85, 96, and 212. Biochemistry 27, 8516–8520. https://doi.org/10.1021/bi00423a002.

Braiman, M.S., Bousche, O., and Rothschild, K.J. (1991). Protein dynamics in the bacteriorhodopsin photocycle: submillisecond Fourier transform infrared spectra of the L, M, and N photointermediates. Proc. Natl. Acad. Sci. 88, 2388–2392. https://doi.org/10.1073/pnas.88.6.2388.

Braiman, M.S., Dioumaev, A.K., and Lewis, J.R. (1996). A large photolysis-induced pKa increase of the chromophore counterion in bacteriorhodopsin: implications for ion transport mechanisms of retinal proteins. Biophys. J. 70, 939–947. https://doi.org/10.1016/S0006-3495(96)79637-6.

Brown, L.S., and Lanyi, J.K. (1996). Determination of the transiently lowered pKa of the retinal Schiff base during the photocycle of bacteriorhodopsin. Proc. Natl. Acad. Sci. 93, 1731–1734. https://doi.org/10.1073/pnas.93.4.1731.

Brown, L.S., Bonet, L., Needleman, R., and Lanyi, J.K. (1993). Estimated acid dissociation constants of the Schiff base, Asp-85, and Arg-82 during the bacteriorhodopsin photocycle. Biophys. J. 65, 124–130. https://doi.org/10.1016/S0006-3495(93)81064-6.

Chandonia, J.-M., and Brenner, S.E. (2006). The impact of structural genomics: expectations and outcomes. Science 311, 347–351. https://doi.org/10.1126/science.1121018.

Chapman, H.N. (2019). X-ray free-electron lasers for the structure and dynamics of macromolecules. Annu. Rev. Biochem. 88, 35–58. https://doi.org/10.1146/annurev-biochem-013118-110744.

Dencher, N.A., Kohl, K.D., and Heyn, M.P. (1983). Photochemical cycle and light-dark adaptation of monomeric and aggregated bacteriorhodopsin in various lipid environments. Biochemistry 22, 1323–1334. https://doi.org/10.1021/bi00275a002.

Dickopf, S., Alexiev, U., Krebs, M.P., Otto, H., Mollaaghababa, R., Khorana, H.G., and Heyn, M.P. (1995). Proton transport by a bacteriorhodopsin mutant, aspartic acid-85-->asparagine, initiated in the unprotonated Schiff base state. Proc. Natl. Acad. Sci. 92, 11519–11523. https://doi.org/10.1073/pnas.92.25.11519.

Diller, R., Maiti, S., Walker, G.C., Cowen, B.R., Pippenger, R., Bogomolni, R.A., and Hochstrasser, R.M. (1995). Femtosecond time-resolved infrared laser study of the J-K transition of bacteriorhodopsin. Chem. Phys. Lett. 241, 109–115. https://doi.org/10.1016/0009-2614(95)00598-X.

Dioumaev, A.K., Brown, L.S., Needleman, R., and Lanyi, J.K. (1999). Fourier transform infrared spectra of a late intermediate of the bacteriorhodopsin photocycle suggest transient protonation of Asp-212. Biochemistry 38, 10070–10078. https://doi.org/10.1021/bi990873+.

Druckmann, S., Ottolenghi, M., Pande, A., Pande, J., and Callender, R.H. (1982). Acid-base equilibrium of the Schiff base in bacteriorhodopsin. Biochemistry 21, 4953–4959. https://doi.org/10.1021/bi00263a019.

Ernst, O.P., Lodowski, D.T., Elstner, M., Hegemann, P., Brown, L.S., and Kandori, H. (2014). Microbial and animal rhodopsins: Structures, functions, and molecular mechanisms. Chem. Rev. 114, 126–163. https://doi.org/10.1021/cr4003769.

Fahmy, K., Weidlich, O., Engelhard, M., Tittor, J., Oesterhelt, D., and Siebert, F. (1992). Identification of the proton acceptor of Schiff base deprotonation in bacteriorhodopsin: A Fourier-transform-infrared study of the mutant Asp85 →Glu in its natural lipid environment. Photochem. Photobiol. 56, 1073–1083. https://doi.org/10.1111/j.1751-1097.1992.tb09731.x.

Fahr, A., and Bamberg, E. (1982). Photocurrents of dark-adapted bacteriorhodopsin on black lipid membranes. FEBS Lett. 140, 251–253. https://doi.org/10.1016/0014-5793(82)80906-X.

Fenno, L., Yizhar, O., and Deisseroth, K. (2011). The development and application of optogenetics. Annu. Rev. Neurosci. 34, 389–412. https://doi.org/10.1146/annurev-neuro-061010-113817.

Friedrich, D., Brünig, F.N., Nieuwkoop, A.J., Netz, R.R., Hegemann, P., and Oschkinat, H. (2020). Collective exchange processes reveal an active site proton cage in bacteriorhodopsin. Commun. Biol. 3, 4. https://doi.org/10.1038/s42003-019-0733-7.

Garczarek, F., Wang, J., El-Sayed, M.A., and Gerwert, K. (2004). The assignment of the different infrared continuum absorbance changes observed in the 3000–1800-cm-1 region during the bacteriorhodopsin photocycle. Biophys. J. 87, 2676–2682. https://doi.org/10.1529/biophysj.104.046433.

Garczarek, F., Brown, L.S., Lanyi, J.K., and Gerwert, K. (2005). Proton binding within a membrane protein by a protonated water cluster. Proc. Natl. Acad. Sci. 102, 3633–3638. https://doi.org/10.1073/pnas.0500421102.

Ge, X., and Gunner, M.R. (2016). Unraveling the mechanism of proton translocation in the extracellular half-channel of bacteriorhodopsin: Proton translocation in bacteriorhodopsin. Proteins Struct. Funct. Bioinforma. 84, 639–654. https://doi.org/10.1002/prot.25013.

Gerwert, K., Hess, B., Soppa, J., and Oesterhelt, D. (1989). Role of aspartate-96 in proton translocation by bacteriorhodopsin. Proc. Natl. Acad. Sci. 86, 4943–4947. https://doi.org/10.1073/pnas.86.13.4943.

Gerwert, K., Souvignier, G., and Hess, B. (1990). Simultaneous monitoring of light-induced changes in protein side-group protonation, chromophore isomerization, and backbone motion of bacteriorhodopsin by time-resolved Fourier-transform infrared spectroscopy. Proc. Natl. Acad. Sci. 87, 9774–9778. https://doi.org/10.1073/pnas.87.24.9774.

Glynn, C., and Rodriguez, J.A. (2019). Data-driven challenges and opportunities in crystallography. Emerg. Top. Life Sci. ETLS20180177. https://doi.org/10.1042/ETLS20180177.

Govindjee, R., Balashov, S.P., and Ebrey, T.G. (1990). Quantum efficiency of the photochemical cycle of bacteriorhodopsin. Biophys. J. 58, 597–608. https://doi.org/10.1016/S0006-3495(90)82403-6.

Govindjee, R., Balashov, S.P., Ebrey, T., Oesterhelt, D., Steinberg, G., and Sheves, M. (1994). Lowering the intrinsic pKa of the chromophore’s Schiff base can restore its light-induced deprotonation in the inactive Tyr-57→Asn mutant of bacteriorhodopsin. J. Biol. Chem. 269, 14353–14354. https://doi.org/10.1016/S0021-9258(17)36626-7.

Govindjee, R., Misra, S., Balashov, S.P., Ebrey, T.G., Crouch, R.K., and Menick, D.R. (1996). Arginine-82 regulates the pKa of the group responsible for the light-driven proton release in bacteriorhodopsin. Biophys. J. 71, 1011–1023. https://doi.org/10.1016/S0006-3495(96)79302-5.

Gozem, S., Luk, H.L., Schapiro, I., and Olivucci, M. (2017). Theory and simulation of the ultrafast double-bond isomerization of biological chromophores. Chem. Rev. 117, 13502–13565. https://doi.org/10.1021/acs.chemrev.7b00177.

Grigorieff, N., Ceska, T.A., Downing, K.H., Baldwin, J.M., and Henderson, R. (1996). Electron-crystallographic refinement of the structure of bacteriorhodopsin. J. Mol. Biol. 259, 393–421. https://doi.org/10.1006/jmbi.1996.0328.

Hackett, N.R., Stern, L.J., Chao, B.H., Kronis, K.A., and Khorana, H.G. (1987). Structure-function studies on bacteriorhodopsin. V. Effects of amino acid substitutions in the putative helix F. J. Biol. Chem. 262, 9277–9284. https://doi.org/10.1016/S0021-9258(18)48077-5.

Hage, W., Kim, M., Frei, H., and Mathies, R.A. (1996). Protein dynamics in the bacteriorhodopsin photocycle: A nanosecond step-scan FTIR investigation of the KL to L transition. J. Phys. Chem. 100, 16026–16033. https://doi.org/10.1021/jp9614198.

Harbison, G.S., Smith, S.O., Pardoen, J.A., Winkel, C., Lugtenburg, J., Herzfeld, J., Mathies, R., and Griffin, R.G. (1984). Dark-adapted bacteriorhodopsin contains 13-cis, 15-syn and all-trans, 15-anti retinal Schiff bases. Proc. Natl. Acad. Sci. 81, 1706–1709. https://doi.org/10.1073/pnas.81.6.1706.

Henderson, R., and Unwin, P.N.T. (1975). Three-dimensional model of purple membrane obtained by electron microscopy. Nature 257, 28–32. https://doi.org/10.1038/257028a0.

Henderson, R., Baldwin, J.M., Ceska, T.A., Zemlin, F., Beckmann, E., and Downing, K.H. (1990). Model for the structure of bacteriorhodopsin based on high-resolution electron cryo-microscopy. J. Mol. Biol. 213, 899–929. https://doi.org/10.1016/S0022-2836(05)80271-2.

Henry, E.R., and Hofrichter, J. (1992). Singular value decomposition: Application to analysis of experimental data. In Numerical Computer Methods, (Academic Press), pp. 129–192.

Herbst, J. (2002). Femtosecond infrared spectroscopy of bacteriorhodopsin chromophore isomerization. Science 297, 822–825. https://doi.org/10.1126/science.1072144.

Heyde, M.E., Gill, D., Kilponen, R.G., and Rimai, L. (1971). Raman spectra of Schiff bases of retinal (models of visual photoreceptors). J. Am. Chem. Soc. 93, 6776–6780. https://doi.org/10.1021/ja00754a012.

Jardetzky, O. (1966). Simple allosteric model for membrane pumps. Nature 211, 969–970. https://doi.org/10.1038/211969a0.

Jonas, R., and Ebrey, T.G. (1991). Binding of a single divalent cation directly correlates with the blue-to-purple transition in bacteriorhodopsin. Proc. Natl. Acad. Sci. 88, 149–153. https://doi.org/10.1073/pnas.88.1.149.

Jung, Y.O., Lee, J.H., Kim, J., Schmidt, M., Moffat, K., Šrajer, V., and Ihee, H. (2013). Volume-conserving trans–cis isomerization pathways in photoactive yellow protein visualized by picosecond X-ray crystallography. Nat. Chem. 5, 212–220. https://doi.org/10.1038/nchem.1565.

Kahan, A., Nahmias, O., Friedman, N., Sheves, M., and Ruhman, S. (2007). Following photoinduced dynamics in bacteriorhodopsin with 7-fs impulsive vibrational spectroscopy. J. Am. Chem. Soc. 129, 537–546. https://doi.org/10.1021/ja064910d.

Kandori, H. (2015). Ion-pumping microbial rhodopsins. Front. Mol. Biosci. 2. https://doi.org/10.3389/fmolb.2015.00052.

Kandori, H., Kinoshita, N., Shichida, Y., and Maeda, A. (1998). Protein structural changes in bacteriorhodopsin upon photoisomerization as revealed by polarized FTIR spectroscopy. J. Phys. Chem. B 102, 7899–7905. https://doi.org/10.1021/jp981949z.

Kandori, H., Kinoshita, N., Yamazaki, Y., Maeda, A., Shichida, Y., Needleman, R., Lanyi, J.K., Bizounok, M., Herzfeld, J., Raap, J., et al. (1999). Structural change of threonine 89 upon photoisomerization in bacteriorhodopsin as revealed by polarized FTIR spectroscopy. Biochemistry 38, 9676–9683. https://doi.org/10.1021/bi990713y.

Kandori, H., Belenky, M., and Herzfeld, J. (2002). Vibrational frequency and dipolar orientation of the protonated Schiff base in bacteriorhodopsin before and after photoisomerization. Biochemistry 41, 6026–6031. https://doi.org/10.1021/bi025585j.

Kobayashi, T., Saito, T., and Ohtani, H. (2001). Real-time spectroscopy of transition states in bacteriorhodopsin during retinal isomerization. Nature 414, 531–534. https://doi.org/10.1038/35107042.

Kovacs, G.N., Colletier, J.-P., Grünbein, M.L., Yang, Y., Stensitzki, T., Batyuk, A., Carbajo, S., Doak, R.B., Ehrenberg, D., Foucar, L., et al. (2019). Three-dimensional view of ultrafast dynamics in photoexcited bacteriorhodopsin. Nat. Commun. 10, 3177. https://doi.org/10.1038/s41467-019-10758-0.

Lanyi, J.K., and Schobert, B. (2007). Structural changes in the L photointermediate of bacteriorhodopsin. J. Mol. Biol. 365, 1379–1392. https://doi.org/10.1016/j.jmb.2006.11.016.

Lewis, A., Spoonhower, J., Bogomolni, R.A., Lozier, R.H., and Stoeckenius, W. (1974). Tunable laser resonance Raman spectroscopy of bacteriorhodopsin. Proc. Natl. Acad. Sci. 71, 4462–4466. https://doi.org/10.1073/pnas.71.11.4462.

Li, Y.-T., Tian, Y., Tian, H., Tu, T., Gou, G.-Y., Wang, Q., Qiao, Y.-C., Yang, Y., and Ren, T.-L. (2018). A review on bacteriorhodopsin-based bioelectronic devices. Sensors 18, 1368. https://doi.org/10.3390/s18051368.

Liebschner, D., Afonine, P.V., Baker, M.L., Bunkóczi, G., Chen, V.B., Croll, T.I., Hintze, B., Hung, L.-W., Jain, S., McCoy, A.J., et al. (2019). Macromolecular structure determination using X-rays, neutrons and electrons: recent developments in Phenix. Acta Crystallogr. Sect. Struct. Biol. 75, 861–877. https://doi.org/10.1107/S2059798319011471.

Logunov, S.L., El-Sayed, M.A., Song, L., and Lanyi, J.K. (1996). Photoisomerization quantum yield and apparent energy content of the K intermediate in the photocycles of bacteriorhodopsin, its mutants D85N, R82Q, and D212N, and deionized blue bacteriorhodopsin. J. Phys. Chem. 100, 2391–2398. https://doi.org/10.1021/jp9515242.

Lórenz-Fonfría, V.A., and Kandori, H. (2009). Spectroscopic and kinetic evidence on how bacteriorhodopsin accomplishes vectorial proton transport under functional conditions. J. Am. Chem. Soc. 131, 5891–5901. https://doi.org/10.1021/ja900334c.

Lorenz-Fonfria, V.A., Saita, M., Lazarova, T., Schlesinger, R., and Heberle, J. (2017). pH-sensitive vibrational probe reveals a cytoplasmic protonated cluster in bacteriorhodopsin. Proc. Natl. Acad. Sci. 114, E10909–E10918. https://doi.org/10.1073/pnas.1707993114.

Lozier, R.H., Bogomolni, R.A., and Stoeckenius, W. (1975). Bacteriorhodopsin: a light-driven proton pump in Halobacterium Halobium. Biophys. J. 15, 955–962. https://doi.org/10.1016/S0006-3495(75)85875-9.

Luecke, H., Schobert, B., Richter, H.-T., Cartailler, J.-P., and Lanyi, J.K. (1999a). Structure of bacteriorhodopsin at 1.55 Å resolution. J. Mol. Biol. 291, 899–911. https://doi.org/10.1006/jmbi.1999.3027.

Luecke, H., Schobert, B., Richter, H.-T., Cartailler, J.-P., and Lanyi, J.K. (1999b). Structural changes in bacteriorhodopsin during ion transport at 2 angstrom resolution. Science 286, 255–260. https://doi.org/10.1126/science.286.5438.255.

Luecke, H., Schobert, B., Cartailler, J.-P., Richter, H.-T., Rosengarth, A., Needleman, R., and Lanyi, J.K. (2000). Coupling photoisomerization of retinal to directional transport in bacteriorhodopsin. J. Mol. Biol. 300, 1237–1255. https://doi.org/10.1006/jmbi.2000.3884.

Maeda, A., Iwasa, T., and Yoshizawa, T. (1977). Isomeric composition of retinal chromophore in dark-adapted bacteriorhodopsin. J. Biochem. (Tokyo) 82, 1599–1604. https://doi.org/10.1093/oxfordjournals.jbchem.a131855.

Marcus, M.A., and Lewis, A. (1977). Kinetic resonance Raman spectroscopy: Dynamics of deprotonation of the Schiff base of bacteriorhodopsin. Science 195, 1328–1330. https://doi.org/10.1126/science.841330.

Marti, T., Otto, H., Mogi, T., Rösselet, S.J., Heyn, M.P., and Khorana, H.G. (1991). Bacteriorhodopsin mutants containing single substitutions of serine or threonine residues are all active in proton translocation. J. Biol. Chem. 266, 6919–6927. https://doi.org/10.1016/S0021-9258(20)89590-8.

Marx, D., Tuckerman, M.E., Hutter, J., and Parrinello, M. (1999). The nature of the hydrated excess proton in water. Nature 397, 601–604. https://doi.org/10.1038/17579.

Mathias, G., and Marx, D. (2007). Structures and spectral signatures of protonated water networks in bacteriorhodopsin. Proc. Natl. Acad. Sci. 104, 6980–6985. https://doi.org/10.1073/pnas.0609229104.

Mathies, R., Brito Cruz, C., Pollard, W., and Shank, C. (1988). Direct observation of the femtosecond excited-state cis-trans isomerization in bacteriorhodopsin. Science 240, 777–779. https://doi.org/10.1126/science.3363359.

Matsui, Y., Sakai, K., Murakami, M., Shiro, Y., Adachi, S., Okumura, H., and Kouyama, T. (2002). Specific damage induced by X-ray radiation and structural changes in the primary photoreaction of bacteriorhodopsin. J. Mol. Biol. 324, 469–481. https://doi.org/10.1016/S0022-2836(02)01110-5.

McCarty, C.G. (1970). Chapter 9 syn-anti isomerizations and rearrangements. In The Chemistry of the Carbon-Nitrogen Double Bond, (John Wiley & Sons, Ltd), p. 363.

Miller, R.J.D., Paré-Labrosse, O., Sarracini, A., and Besaw, J.E. (2020). Three-dimensional view of ultrafast dynamics in photoexcited bacteriorhodopsin in the multiphoton regime and biological relevance. Nat. Commun. 11, 1240. https://doi.org/10.1038/s41467-020-14971-0.

Moffat, K. (1989). Time-Resolved Macromolecular Crystallography. Annu. Rev. Biophys. Biophys. Chem. 18, 309–332. https://doi.org/10.1146/annurev.bb.18.060189.001521.

Moffat, K. (2001). Time-resolved biochemical crystallography: A mechanistic perspective. Chem Rev 101, 1569–1582. https://doi.org/10.1021/cr990039q.

Mogi, T., Stern, L.J., Marti, T., Chao, B.H., and Khorana, H.G. (1988). Aspartic acid substitutions affect proton translocation by bacteriorhodopsin. Proc. Natl. Acad. Sci. 85, 4148–4152. https://doi.org/10.1073/pnas.85.12.4148.

Mowery, P.C., Lozier, R.H., Chae, Q., Tseng, Y.-W., Taylor, M., and Stoeckenius, W. (1979). Effect of acid pH on the absorption spectra and photoreactions of bacteriorhodopsin. Biochemistry 18, 4100–4107. https://doi.org/10.1021/bi00586a007.

Nango, E., Royant, A., Kubo, M., Nakane, T., Wickstrand, C., Kimura, T., Tanaka, T., Tono, K., Song, C., Tanaka, R., et al. (2016). A three-dimensional movie of structural changes in bacteriorhodopsin. Science 354, 1552–1557. https://doi.org/10.1126/science.aah3497.

Neutze, R., Pebay-Peyroula, E., Edman, K., Royant, A., Navarro, J., and Landau, E.M. (2002). Bacteriorhodopsin: a high-resolution structural view of vectorial proton transport. Biochim. Biophys. Acta BBA - Biomembr. 1565, 144–167. https://doi.org/10.1016/S0005-2736(02)00566-7.

Nogly, P., Weinert, T., James, D., Carbajo, S., Ozerov, D., Furrer, A., Gashi, D., Borin, V., Skopintsev, P., Jaeger, K., et al. (2018). Retinal isomerization in bacteriorhodopsin captured by a femtosecond x-ray laser. Science 361, eaat0094. https://doi.org/10.1126/science.aat0094.

Nuss, M.C., Zinth, W., Kaiser, W., Kolling, E., and Oesterhelt, D. (1985). Femtosecond spectroscopy of the first events of the photochemical cycle in bacteriorhodopsin. Chem. Phys. Lett. 117, 1–7. https://doi.org/10.1016/0009-2614(85)80393-6.

Oesterhelt, D., and Hess, B. (1973). Reversible photolysis of the purple complex in the purple membrane of Halobacterium halobium. Eur. J. Biochem. 37, 316–326. https://doi.org/10.1111/j.1432-1033.1973.tb02990.x.

Oesterhelt, D., Meentzen, M., and Schuhmann, L. (1973). Reversible dissociation of the purple complex in bacteriorhodopsin and identification of 13-cis and all-trans-retinal as its chromophores. Eur. J. Biochem. 40, 453–463. https://doi.org/10.1111/j.1432-1033.1973.tb03214.x.

Ohno, K., Takeuchi, Y., and Yoshida, M. (1977). Effect of light-adaptation on the photoreaction of bacteriorhodopsin from Halobacterium halobium. Biochim. Biophys. Acta BBA - Bioenerg. 462, 575–582. https://doi.org/10.1016/0005-2728(77)90102-5.

Otto, H., Marti, T., Holz, M., Mogi, T., and Stern, L.J. (1990). Substitution of amino acids Asp-85, Asp-212, and Arg-82 in bacteriorhodopsin affects the proton release phase of the pump and the pK of the Schiff base. Proc. Natl. Acad. Sci. 87, 1018–1022. https://doi.org/10.1073/pnas.87.3.1018.

Pebay-Peyroula, E., Rummel, G., Rosenbusch, J.P., and Landau, E.M. (1997). X-ray structure of bacteriorhodopsin at 2.5 Angstroms from microcrystals grown in lipidic cubic phases. Science 277, 1676–1681. https://doi.org/10.1126/science.277.5332.1676.

Ren, Z. (2013a). Reaction trajectory revealed by a joint analysis of Protein Data Bank. PLoS ONE 8, e77141. https://doi.org/10.1371/journal.pone.0077141.

Ren, Z. (2013b). Reverse engineering the cooperative machinery of human hemoglobin. PLoS ONE 8, e77363. https://doi.org/10.1371/journal.pone.0077363.

Ren, Z. (2016). Molecular events during translocation and proofreading extracted from 200 static structures of DNA polymerase. Nucleic Acids Res. 6, 1–13. https://doi.org/10.1093/nar/gkw555.

Ren, Z. (2019). Ultrafast structural changes decomposed from serial crystallographic data. J. Phys. Chem. Lett. 10, 7148–7163. https://doi.org/10.1021/acs.jpclett.9b02375.

Ren, Z. (2022). Photoinduced isomerization sampling of retinal in bacteriorhodopsin. PNAS Nexus https://doi.org/10.1093/pnasnexus/pgac103.

Ren, Z., Perman, B., Srajer, V., Teng, T.-Y., Pradervand, C., Bourgeois, D., Schotte, F., Ursby, T., Kort, R., Wulff, M., et al. (2001). A molecular movie at 1.8 Å resolution displays the photocycle of photoactive yellow protein, a eubacterial blue-light receptor, from nanoseconds to seconds. Biochemistry 40, 13788–13801. https://doi.org/10.1021/bi0107142.

Ren, Z., Chan, P.W.Y., Moffat, K., Pai, E.F., Royer, W.E., Šrajer, V., and Yang, X. (2013). Resolution of structural heterogeneity in dynamic crystallography. Acta Cryst D69, 946–959. https://doi.org/10.1107/S0907444913003454.

Ren, Z., Ren, P.X., Balusu, R., and Yang, X. (2016). Transmembrane helices tilt, bend, slide, torque, and unwind between functional states of rhodopsin. Sci. Rep. 6, 34129. https://doi.org/10.1038/srep34129.

Ren, Z., Ayhan, M., Bandara, S., Bowatte, K., Kumarapperuma, I., Gunawardana, S., Shin, H., Wang, C., Zeng, X., and Yang, X. (2018). Crystal-on-crystal chips for in situ serial diffraction at room temperature. Lab. Chip 18, 2246–2256. https://doi.org/10.1039/C8LC00489G.

Richter, H.T., Needleman, R., and Lanyi, J.K. (1996a). Perturbed interaction between residues 85 and 204 in Tyr-185-->Phe and Asp-85-->Glu bacteriorhodopsins. Biophys. J. 71, 3392–3398. https://doi.org/10.1016/S0006-3495(96)79532-2.

Richter, H.-T., Needleman, R., Kandori, H., Maeda, A., and Lanyi, J.K. (1996b). Relationship of retinal configuration and internal proton transfer at the end of the bacteriorhodopsin photocycle. Biochemistry 35, 15461–15466. https://doi.org/10.1021/bi9612430.

Royant, A., Edman, K., Ursby, T., Pebay-Peyroula, E., Landau, E.M., and Neutze, R. (2000). Helix deformation is coupled to vectorial proton transport in the photocycle of bacteriorhodopsin. Nature 406, 645–648. https://doi.org/10.1038/35020599.

Sasaki, J., Yuzawa, T., Kandori, H., Maeda, A., and Hamaguchi, H. (1995). Nanosecond time-resolved infrared spectroscopy distinguishes two K species in the bacteriorhodopsin photocycle. Biophys. J. 68, 2073–2080. https://doi.org/10.1016/S0006-3495(95)80386-3.

Sass, H.J., Büldt, G., Gessenich, R., Hehn, D., Neff, D., Schlesinger, R., Berendzen, J., and Ormos, P. (2000). Structural alterations for proton translocation in the M state of wild-type bacteriorhodopsin. Nature 406, 649–653. https://doi.org/10.1038/35020607.

Schaffer, J.E., Kukshal, V., Miller, J.J., Kitainda, V., and Jez, J.M. (2021). Beyond X-rays: an overview of emerging structural biology methods. Emerg. Top. Life Sci. ETLS20200272. https://doi.org/10.1042/ETLS20200272.

Scherrer, P., Mathew, M.K., Sperling, W., and Stoeckenius, W. (1989). Retinal isomer ratio in dark-adapted purple membrane and bacteriorhodopsin monomers. Biochemistry 28, 829–834. https://doi.org/10.1021/bi00428a063.

Schmidt, M., Rajagopal, S., Ren, Z., and Moffat, K. (2003). Application of singular value decomposition to the analysis of time-resolved macromolecular X-ray data. Biophys. J. 84, 2112–2129. https://doi.org/10.1016/S0006-3495(03)75018-8.

Schmidt, M., Graber, T., Henning, R., and Srajer, V. (2010). Five-dimensional crystallography. Acta Crystallogr. A 66, 198–206. https://doi.org/10.1107/S0108767309054166.

Sheves, M., Albeck, A., Friedman, N., and Ottolenghi, M. (1986). Controlling the pKa of the bacteriorhodopsin Schiff base by use of artificial retinal analogues. Proc. Natl. Acad. Sci. 83, 3262–3266. https://doi.org/10.1073/pnas.83.10.3262.

Shichida, Y., Matuoka, S., Hidaka, Y., and Yoshizawa, T. (1983). Absorption spectra of intermediates of bacteriorhodopsin measured by laser photolysis at room temperatures. Biochim. Biophys. Acta BBA - Bioenerg. 723, 240–246. https://doi.org/10.1016/0005-2728(83)90123-8.

Shin, H., Ren, Z., Zeng, X., Bandara, S., and Yang, X. (2019). Structural basis of molecular logic OR in a dual-sensor histidine kinase. Proc. Natl. Acad. Sci. 116, 19973–19982. https://doi.org/10.1073/pnas.1910855116.

Smith, S.O., Myers, A.B., Pardoen, J.A., Winkel, C., Mulder, P.P.J., Lugtenburg, J., and Mathies, R. (1984). Determination of retinal Schiff base configuration in bacteriorhodopsin. Proc. Natl. Acad. Sci. 81, 2055–2059. https://doi.org/10.1073/pnas.81.7.2055.

Song, L., El-Sayed, M.A., and Lanyi, J.K. (1993). Protein catalysis of the retinal subpicosecond photoisomerization in the primary process of bacteriorhodopsin photosynthesis. Science 261, 891–894. https://doi.org/10.1126/science.261.5123.891.

Šrajer, V., Ren, Z., Teng, T.-Y., Schmidt, M., Ursby, T., Bourgeois, D., Pradervand, C., Schildkamp, W., Wulff, M., and Moffat, K. (2001). Protein conformational relaxation and ligand migration in myoglobin: A nanosecond to millisecond molecular movie from time-resolved Laue X-ray diffraction. Biochemistry 40, 13802–13815. https://doi.org/10.1021/bi010715u.

Stockburger, M., Klusmann, W., Gattermann, H., Massig, G., and Peters, R. (1979). Photochemical cycle of bacteriorhodopsin studied by resonance Raman spectroscopy. Biochemistry 18, 4886–4900. https://doi.org/10.1021/bi00589a017.

Stoeckenius, W. (1999). Bacterial rhodopsins: Evolution of a mechanistic model for the ion pumps. Protein Sci. 8, 447–459. https://doi.org/10.1110/ps.8.2.447.

Stoeckenius, W., Lozier, R.H., and Bogomolni, R.A. (1979). Bacteriorhodopsin and the purple membrane of halobacteria. Biochim. Biophys. Acta BBA - Rev. Bioenerg. 505, 215–278. https://doi.org/10.1016/0304-4173(79)90006-5.

Su, T., and Purohit, P.K. (2009). Mechanics of forced unfolding of proteins. Acta Biomater. 5, 1855–1863. https://doi.org/10.1016/j.actbio.2009.01.038.

Swanson, J.M.J., and Simons, J. (2009). Role of charge transfer in the structure and dynamics of the hydrated proton. J. Phys. Chem. B 113, 5149–5161. https://doi.org/10.1021/jp810652v.

Takahashi, H., Rico, F., Chipot, C., and Scheuring, S. (2018). α-helix unwinding as force buffer in spectrins. ACS Nano 12, 2719–2727. https://doi.org/10.1021/acsnano.7b08973.

Takeda, K., Matsui, Y., Kamiya, N., Adachi, S., Okumura, H., and Kouyama, T. (2004). Crystal structure of the M intermediate of bacteriorhodopsin: Allosteric structural changes mediated by sliding movement of a transmembrane helix. J. Mol. Biol. 341, 1023–1037. https://doi.org/10.1016/j.jmb.2004.06.080.

Terner, J., and El-Sayed, M.A. (1985). Time-resolved resonance Raman spectroscopy of photobiological and photochemical transients. Acc. Chem. Res. 18, 331–338. https://doi.org/10.1021/ar00119a002.

Tittor, J., Schweiger, U., Oesterhelt, D., and Bamberg, E. (1994). Inversion of proton translocation in bacteriorhodopsin mutants D85N, D85T, and D85,96N. Biophys. J. 67, 1682–1690. https://doi.org/10.1016/S0006-3495(94)80642-3.

Tsin, A., Betts-Obregon, B., and Grigsby, J. (2018). Visual cycle proteins: Structure, function, and roles in human retinal disease. J. Biol. Chem. 293, 13016–13021. https://doi.org/10.1074/jbc.AW118.003228.

Ursby, T., and Bourgeois, D. (1997). Improved estimation of structure-factor difference amplitudes from poorly accurate data. Acta Crystallogr. A 53, 564–575. https://doi.org/10.1107/S0108767397004522.

Wang, J., and El-Sayed, M.A. (2001). Time-resolved Fourier transform infrared spectroscopy of the polarizable proton continua and the proton pump mechanism of bacteriorhodopsin. Biophys. J. 80, 961–971. https://doi.org/10.1016/S0006-3495(01)76075-4.

Weinert, T., Skopintsev, P., James, D., Dworkowski, F., Panepucci, E., Kekilli, D., Furrer, A., Brünle, S., Mous, S., Ozerov, D., et al. (2019). Proton uptake mechanism in bacteriorhodopsin captured by serial synchrotron crystallography. Science 365, 61–65. https://doi.org/10.1126/science.aaw8634.

Westerhoff, H., and Dancshazy, Z. (1984). Keeping a light-driven proton pump under control. Trends Biochem. Sci. 9, 112–117. https://doi.org/10.1016/0968-0004(84)90107-5.

Wickstrand, C., Dods, R., Royant, A., and Neutze, R. (2015). Bacteriorhodopsin: Would the real structural intermediates please stand up? Biochim. Biophys. Acta 1850, 536–553. https://doi.org/10.1016/j.bbagen.2014.05.021.

Wickstrand, C., Nogly, P., Nango, E., Iwata, S., Standfuss, J., and Neutze, R. (2019). Bacteriorhodopsin: Structural insights revealed using X-ray lasers and synchrotron radiation. Annu. Rev. Biochem. 88, 59–83. https://doi.org/10.1146/annurev-biochem-013118-111327.

Yang, X., Ren, Z., Kuk, J., and Moffat, K. (2011). Temperature-scan cryocrystallography reveals reaction intermediates in bacteriophytochrome. Nature 479, 428–432. https://doi.org/10.1038/nature10506.

Zhong, Q., Ruhman, S., Ottolenghi, M., Sheves, M., Friedman, N., Atkinson, G.H., and Delaney, J.K. (1996). Reexamining the primary light-induced events in bacteriorhodopsin using a synthetic C13=C14-locked chromophore. J Am Chem Soc 118, 12828–12829. https://doi.org/10.1021/ja961058+.

Zimanyi, L., Varo, G., Chang, M., Ni, B., Needleman, R., and Lanyi, J.K. (1992). Pathways of proton release in the bacteriorhodopsin photocycle. Biochemistry 31, 8535–8543. https://doi.org/10.1021/bi00151a022.

Zscherp, C., Schlesinger, R., and Heberle, J. (2001). Time-resolved FT-IR spectroscopic investigation of the pH-dependent proton transfer reactions in the E194Q mutant of bacteriorhodopsin. Biochem. Biophys. Res. Commun. 283, 57–63. https://doi.org/10.1006/bbrc.2001.4730.

