## Supplementary materials for "Directional Proton Conductance in Bacteriorhodopsin Is Driven by Concentration Gradient, Not Affinity Gradient"

Zhong Ren

Department of Chemistry, University of Illinois at Chicago, Chicago, IL 60607, USA
Renz Research, Inc., Westmont, IL 60559, USA


ORCID 0000-0001-7098-3127

#### Supplementary Information

*Mutant functions*

Under the mechanism of proton conductance driven by concentration gradient presented here, a number of mutant functions are explained. How mutants behave, in return, further support these hypotheses. Light adaptations in D85N and D85T mutants convert only ~50% of the chromophores into all-*trans* unlike the complete process in the wildtype (Tittor et al., 1994). With 50% all-*trans* retinal that is the active isomer for proton pumping, the pumping activity is still abolished in D85N at neutral pH (Mogi et al., 1988). The remaining proton acceptor Asp212 is H-boned to the phenol hydroxy group of Tyr57 (Fig. S17). The mutant D85N has a smaller proton capacity in its inner EC channel. After two successful photocycles, Asp212 and Wat402 are both protonated. However, proton conductance is stopped at the carbonyl groups =O of both side chains of Asn85 and Asp212, the amide group $-NH_2$ of Asn85, and the hydroxy groups -OH of Tyr57 and 185, since none of them can be protonated further. The smaller proton capacity in its inner EC channel is easily filled and drives a much faster thermal isomerization so that the light adaptation could only achieve 50% functional chromophores. The lack of a proton outlet causes the SB to be reprotonated quickly during I′ or I before isomerization. In addition, the smaller proton capacity completely filled with two protons slows down the photoisomerization by at least four-fold and with anther 20-fold slower second phase (Song et al., 1993), which allows reprotonation of the SB still in the EC channel before isomerization. The proton pump stalls with repeated futile photocycles. Photovoltage measurement across a vesicle membrane incorporated with D85N mutant protein showed a movement of positive charge into the CP side before 1 μs that the exact timing was too fast to be resolved, and later the positive charge moves back to the EC side at ~10 ms (Otto et al., 1990). The net charge movement is near zero. This agrees well with the repeated futile photocycles described here due to either a failed deprotonation from the SB or a rapid reprotonation before isomerization. The isomerization and reisomerization during I → J′ and N → O bring a proton into the CP half channel and back into the EC side. The accelerated reprotonation of the SB on the EC side and the tardy isomerization due to the proton outlet shutdown indicate an even more acidic inner EC channel despite a missing acidic group. This is consistent with the further red shifts to absorption maxima of 595-617 nm for D85N mutant compared to the wildtype in ground state (Logunov et al., 1996; Otto et al., 1990; Tittor et al., 1994).

It has been shown that the proton pumping activity of D85N can be somewhat restored at an alkaline pH (Dickopf et al., 1995), which is consistent with the concentration-driven proton conductance proposed here. The phenol hydroxy group of Tyr57, a weak acid, could start to conduct proton to Wat403 and Arg82 only at an alkaline pH by deprotonation. On the other hand, some activity remains in D212N, which suggests a proton outlet going through Asp85, Wat401, 400, Arg82, and beyond (Fig. S17). In addition, Asp to Glu mutants are harder to assess because of their severe structural impact. Proton pumping activity is also abolished in D212A (Mogi et al., 1988) perhaps due to a more mobile Wat402 or a hydronium that reprotonates the SB before isomerization or fails to withhold the proton during the SB U-turn. This mutant function further underscores the importance that Wat402 or a hydronium ion maintains its H-bonds to both Asp 85 and 212 throughout the photocycle as observed in this study despite its large traveling trajectory (Fig. 2a).

Mutants at the site of Arg82 are crucial tests to the mechanisms proposed here. M formation in the wildtype is pH-dependent at a few μs under alkaline pH and 85 μs at neutral pH (Balashov et al., 1993; Zimanyi et al., 1992), which nicely illustrates the concentration-gradient-driven proton conductance proposed here as faster proton release takes place driven by greater proton gradient. The proton barrier would be breached in R82A mutant (Fig. 5a). Therefore, the EC half channel must be filled with a continuous water network. Very little proton gradient can be established along the water filled channel during continuous pumping activity. That is why both dark and light adapted states of R82A mutant blue shift ~5 nm from the corresponding states of the wildtype as the breached proton barrier cannot hold a proton pool (Balashov et al., 1993). R82A exhibits the same rate of proton uptake as the wildtype, but a rapid M formation without a pH dependency between neutral pH and pH10. Such rapid M formation is even faster than the wildtype under alkaline condition (Balashov et al., 1993). This is obviously due to the inability to establish the super-acidic inner EC channel without an intact proton barrier so that the protonation of Asp85 is accelerated compared to the wildtype. More interestingly, the fast phase of proton release is absent in R82A. Instead, a slow relaxation of 30 ms after proton uptake was observed (Balashov et al., 1993). Under the mechanism of a concentration-gradient-driven proton conductance, this slow relaxation is the spontaneous movement of the newly pumped proton in the same photocycle driven by a small gradient. Therefore, the "early release" in the fast phase of proton release cannot be observed without a well-established super-acidic inner EC channel in R82A. The "early release" observed from the wildtype is actually a much slower proton movement, in which a proton pumped into the super-acidic inner channel several photocycles ago is "squeezed out" by a newly pumped proton. This proton released from the wildtype appears to be coupled with or even faster than the protonation of Asp85.

Similar to R82A, another mutant R82Q also features rapid, biphasic formation of M state at 1 and 160 μs but delayed proton release after uptake at neutral pH (Govindjee et al., 1996). However, unlike R82A, proton release from R82Q shows its strong pH-dependency. At neutral pH, the majority of proton release occurs late at 23 ms. If the pH increases to 8, most proton release precedes proton uptake at a few ms. The amide group on the side chain of the neutral residue Gln82 could be protonated by hydroniums in the inner EC channel. However, the amide group is not a sufficient proton barrier with its p$K_a$ of -1.4 that could only establish a proton puddle less acidic compared to that of the wildtype maintained by a divalent guanidinium ion, that is, a shallower bucket than that in Fig. 1. Another important factor to a smaller proton puddle is that Gln side chain is two bonds shorter than Arg side chain. As a result, rapid proton release at a few ms is possible with a sufficient proton gradient under high pH. However, proton release is delayed to 23 ms as the proton gradient decreases under neutral pH. Again, the proton apparently released early is one of the protons accumulated in the inner EC channel, not the one freshly pumped in the same photocycle.

Substitutions for Thr89 next to the SB on the inboard side that plays a crucial role to seal off two half channels have great impact to the absorption property, the dark and light adaptations, and the pumping activity (Marti et al., 1991). The absorption peaks of T89V and T89A mutants broaden or even show a bimodal shape. Both exhibit large blue shifts with respect to the wildtype. But it is hard to determine the exact value of the blue shift due to the broadened and bimodal peaks. Nevertheless, it is clear that the large blue shifts result from the leaky seal between two half channels that is incapable of maintaining the required super-acidic inner EC channel so that a protonated SB absorbs in the green as it should be. It is not surprising that both mutants retain ~2/3 of the wildtype activity when operating with a leaky seal. If a single methyl group from the Ala residue is too small to seal off two half channels, why could not a Val residue in the position of Thr do the exact same job? The structural mechanism of Thr89 here to play the role of a seal is as same as that of the next residue Thr90 that prevents byproducts during the photoinduced isomerization sampling (Ren, 2022). Both Thr89 and Thr90 form H-bonds to main chain carbonyl groups in the same helix C, which fix their side chains toward the desired orientations (Fig. S18). A Val substitute would be free to choose any other rotameric positions therefore not act as a good seal despite the similar size and shape to a Thr side chain. Unlike the dark adaptation of the wildtype in which the pH-gradient-driven isomerization reduces the all-*trans* composition from 100% to 50% or even 33%, the dark adaptation of T89A mutant reduces the all-*trans* composition slightly from 42% to 36% (Marti et al., 1991) because of the lack of driving force. But it remains puzzling that T89V mutant does not reduce the all-*trans* composition during its dark adaptation, instead, it increases the composition from 75% to 92%. I suspect that the side chain of Val89 would adopt an opposite rotameric position compared to Thr89 and be positioned more toward the CP side of the SB under the pressure of the straightening helix C, rather than situated on the inboard side in the wildtype, so that the isomer selection is affected.

See also Fig. S12b for P186L mutant with a reduced pumping activity less than half of the wildtype (Hackett et al., 1987).

*Supplementary Q&A*

This study addresses six key questions regarding the operating mechanisms of the light-driven proton pump bR in addition to the question of isomerization sampling (Ren, 2022). Answers to these questions are somewhat scattered in the main text while evidence are presented to support these answers. Here the answers are recapped without the interruption of the technical details of the supporting data.

Q1) How does the protonated SB with a p$K_a$ of 13.3 transfer its proton to the acidic acceptors with a p$K_a$ of 2.2 and when?

A1) Which of the two comparable weak interactions with the proton in $\geqslant N_\zeta:H^+:O<H_2$ survives an abrupt tare while the retinal prepares to undergo photoisomerization is determined by the inertia of the proton (Fig. 1 inset). The SB nitrogen makes a rapid U-turn at ~30 fs (Figs. 2a and S3 inset). It is decisive that the proton stays with the stationary partner and dissociates from the accelerating partner, the SB, due to its inertia at the beginning of the retinal photoisomerization. A force > 500 pN is required to accelerate the proton as fast as the SB. Breaking one of the three equivalent bonds in $H\text{-}O^+<H_2$ is nearly as costly as breaking the proton association with the SB, where the positive charge of the proton may have delocalized. Therefore, the proton joins Wat402 to form a hydronium ion due to its inertia instead of keeping up to maintain the SB protonated. Such a proton transfer uphill over 15 pH units within tens of fs is not achieved by a chemical equilibrium, thus not governed by p$K_a$ values. This proton transfer requires a high peak power d$H$/d$t$ localized at the SB. On the other hand, holding Wat402 in place by both Asp85 and 212 symmetrically from inboard and outboard is as important as the ultrafast shaking of the SB. This proton transfer uses the same mechanism of the tablecloth trick. The early deprotonation from the SB is a separate event from the protonation of Asp85 ten decades later at the formation of M state. This light-driven reaction of a proton pump differs markedly from a thermal driven chemical equilibrium that occurs in the dual-isomerization to form 13,15-*cis* during dark adaptation (Fig. 1 inset).

Q2) Which molecular event, the isomerization, or another event, switches the accessibility of the SB from EC to CP and how does the SB avoid reprotonation on the wrong side?

A2) Immediately before isomerization in the I state, the SB is reoriented toward the inboard while Wat402, a hydronium at this moment, is displaced to the outboard. This geometry is very bad for proton exchange (Fig. S4). Therefore, the accessibility of the SB is already limited even before isomerization. However, accessibility switch of the SB is finally accomplished by the photoisomerization during I $\rightarrow$ J′. The photoisomerization must be quick enough to prevent a chemical equilibrium in the EC half channel. If the equilibrium is reached before isomerization, the SB is reprotonated from the EC side, and the quantum yield of the proton pump is sacrificed. Accelerated reprotonation from the EC channel and tardy isomerization occur in D85N mutant and abolish proton pumping activity.

Q3) How are protons conducted > 15 Å through the rollercoasting p$K_a$ values and released into the EC medium from the protonated SB of a high p$K_a$ through aspartic acids of low p$K_a$, arginine (strictly speaking, its guanidinium ion) and tyrosines of high p$K_a$, and glutamic acids of low p$K_a$ again?

A3) The EC half channel is obviously not lined with residues of p$K_a$ values in a monotonic order (Figs. 5 and S1). Despite numerous observations of p$K_a$ changes, no sufficient evidence supports photoinduced changes to form an ascending order of p$K_a$ values either. Proton is therefore not conducted along an ascending order of p$K_a$ values driven by the proton affinity (Stoeckenius, 1999; Stoeckenius et al., 1979). Instead, proton conductance is driven by concentration gradient, not by affinity gradient. The guanidinium ion of the arginine and tyrosines of high p$K_a$, together with several main chain carbonyls, are arranged to function as a proton levee so that protons could only overflow outward from a super-acidic inner EC channel established after several successful cycles of proton pumping (Fig. 1). No proton is released during the first a few cycles of productive pumping until a sufficient proton gradient is established. Under a super-acidic condition in the inner EC channel, the guanidinium group of Arg82 is protonated for a second time to form a divalent cation. While it swings outward, the extra proton is discharged to Glu194, 204, and a water cluster in the outer EC channel, the proton release complex. Protons could not flow backward into the cell as long as sufficient water is available in the EC medium. The rollercoasting p$K_a$ values ensure the concentration-driven outward conductance of protons and prevent the concentration-driven inward conductance during dark, that is, a unidirectional valve.

Q4) How is a gradient-driven proton backflow prevented in the resting state and during the pumping motions?

A4) A potential leak of protons back into the cell during dark, known as back pressure, is prevented by the exact mechanism that conducts proton outward (A3). However, the chance of proton leak increases dramatically during the molecular motions of proton pumping because the inner EC channel becomes super acidic and the retinal and its surrounding are in constant motions. Lys216 forms a corner at $C_\varepsilon$ that protrudes inboard and makes a van der Waals contact with Thr89 in helix C (Figs. 2b and S12a). Pressed by the corner at $C_\varepsilon$, helix C is kinked and bends away. This contact is a seal between two half channels. This seal could be broken briefly in I′ state but inconsequential. $C_\delta$-$C_\varepsilon$ moves outboard in L state once again due to a flattened 13-*cis* retinal. But this motion is slow enough to allow the spring-loaded helix C to react. Helix C straightens and maintains the contact between the chromophore and Thr89 to prevent a concentration-gradient-driven proton leak back into the CP (Figs. 4f and S16c). During the M states, the corner at $C_\varepsilon$ is reestablished. It pushes Thr89 and helix C to restore its kink.

Q5) How does the retinal chromophore covalently linked to the protein exhibit a swing of absorption maxima nearly as wide as the entire visible spectrum from deep violet to red during its photocycle, while a free retinal compound generally absorbs from near ultraviolet (UV) to green depending on its protonation state?

A5) The range of absorption maxima is extended considerably and shifted to the red side while retinal is incorporated into bR. This is due the super-acidic inner EC channel charged with excess protons right next to the SB. This super charged proton pool is established by the photophysical deprotonation from the SB that does not rely on proton affinities (A1). This excess proton pool drives the directional proton conductance in the EC half channel.

Q6) How does the thermal isomerization occur during dark adaptation that does not require light energy, and why specifically to 13,15-*cis*? What are the physiological functions of the dark and light adaptations, anyway?

A6) It would require a large part of the visual cycle involving multiple enzymes to restore the 11-*cis* retinal chromophore back into visual rhodopsins from the photoisomerization product all-*trans* retinal (Tsin et al., 2018). The thermal isomerization from all-*trans* to 13,15-*cis* during dark adaptation of bR does not just happen without the driving force from the super-acidic inner EC channel established during light adaptation. Despite the exquisite arrangement of the proton barrier and the irregular, bent helix C constantly pressing its Thr89 on the moving part of the chromophore, the super-acidic proton pool cannot be maintained indefinitely. The slow discharge at least four decades slower than the photocycle by the acid driven isomerization does not sacrifice the overall efficiency. The super acidic proton pool could be harmful and needs to be discharged while the pump is not in use. Therefore, the physiological function of the dark adaptation of bR is very like a form of photoprotection. The function of the light adaptation is to establish the require proton concentration gradient for directional proton conductance.

**Methods**

From the outset, the key presumption is that every crystallographic dataset, at a given temperature and a given time delay after the triggering of the photochemical reaction, captures a mixture of unknown number of intermediate species at unknown fractions. Needless to say, all structures of the intermediates are also unknown except the structure at the ground state that has been determined and well refined by static crystallography. A simultaneous solution of all these unknowns requires multiple datasets that are collected at various temperatures or time delays so that a common set of intermediate structures are present in these datasets with variable ratios. If the number of available datasets is far greater than the number of unknowns, a linear system can be established to overdetermine the unknowns with the necessary stereochemical restraints (Ren et al., 2013). This analytical strategy is recapped below.

The methodological advance in this work and the companion work (Ren, 2022) is the refinement of each pure intermediate structure that has been deconvoluted from multiple mixtures. Structure factor amplitudes of a single conformation free of heterogeneity are overdetermined. Unlike the conventional crystallography, such a deconvoluted dataset of structure factor amplitudes are not observed directly from any crystal but computed from many experimental datasets. It could be considered as an extrapolated dataset "on steroids" if compared to the traditional extrapolation of small differences, such as, 3Fo-2Fc map, a technique often used to overcome a partial occupancy of an intermediate structure, except that this deconvolution method is an interpolation among many experimental datasets rather than an extrapolation. The standard structural refinement software is taken full advantage of with the built-in stereochemical constraints, e.g. PHENIX (Adams et al., 2010; Liebschner et al., 2019). In case that the computed deconvolution has not achieved a single pure structural species, the structural refinement is expected to make such indication.

*Difference Fourier maps*

A difference Fourier map is synthesized from a Fourier coefficient set of $F_{\text{light}}$-$F_{\text{reference}}$ with the best available phase set, often from the ground state structure. Before Fourier synthesis, $F_{\text{light}}$ and $F_{\text{reference}}$ must be properly scaled to the same level so that the distribution of difference values is centered at zero and not skewed either way. A weighting scheme proven effective assumes that a greater amplitude of a difference Fourier coefficient $F_{\text{light}}$-$F_{\text{reference}}$ is more likely caused by noise than by signal (Ren et al., 2001, 2013; Šrajer et al., 2001; Ursby and Bourgeois, 1997). Both the dark and light datasets can serve as a reference in difference maps. If a light dataset at a certain delay is chosen as a reference, the difference map shows the changes since that delay time but not the changes prior to that delay. However, both the dark and light datasets must be collected in the same experiment. A cross reference from a different experimental setting usually causes large systematic errors in the difference map that would swamp the desired signals. Each difference map is masked 3.5 Å around the entire molecule of bacteriorhodopsin (bR). No lipid density is analyzed.

*Meta-analysis of protein structures*

Structural meta-analysis based on singular value decomposition (SVD) has been conducted in two forms. In one of them, an interatomic distance matrix is calculated from each protein structure in a related collection. SVD of a data matrix consists of these distance matrices enables a large-scale joint structural comparison but requires no structural alignment (Ren, 2013a, 2013b, 2016). In the second form, SVD is performed on a data matrix of electron density maps of related protein structures (Ren, 2019; Ren et al., 2013; Schmidt et al., 2003, 2010). Both difference electron density maps that require a reference dataset from an isomorphous crystal and simulated annealing omit maps that do not require the same unit cell and space group of the crystals are possible choices in a structural meta-analysis (Ren, 2019; Ren et al., 2013). The distance matrices or the electron density maps that SVD is performed on are called core data. Each distance matrix or electron density map is associated with some metadata that describe the experimental conditions under which the core data are obtained, such as temperature, pH, light illumination, time delay, mutation, etc. These metadata do not enter the SVD procedure. However, they play important role in the subsequent interpretation of the SVD result. This computational method of structural analysis takes advantage of a mathematical, yet practical, definition of conformational space with limited dimensionality (Ren, 2013a). Each experimentally determined structure is a snapshot of the protein structure. A large number of such snapshots taken under a variety of experimental conditions, the metadata, would collectively provide a survey of the accessible conformational space of the protein structure and reveal its rection trajectory. Such joint analytical strategy would not be effective in early years when far fewer protein structures were determined to atomic resolution. Recent rapid growth in protein crystallography, such as in structural genomics (Berman et al., 2012; Chandonia and Brenner, 2006) and in serial crystallography (Glynn and Rodriguez, 2019; Schaffer et al., 2021), has supplied the necessarily wide sampling of protein structures for a joint analytical strategy to come of age. The vacancies or gaps in a conformational space between well-populated conformational clusters often correspond to less stable transient states whose conformations are difficult to capture, if not impossible. These conformations are often key to mechanistic understanding and could be explored by a back calculation based on molecular distance geometry (Ren, 2013a, 2016), the chief computational algorithm in nucleic magnetic resonance spectroscopy (NMR), and by a structure refinement based on reconstituted dataset, a major methodological advance in this work (see below). These structures refined to atomic resolution against reconstituted datasets may reveal short-lived intermediate conformation hard to be captured experimentally. Unfortunately, a protein structure refined against a reconstituted dataset currently cannot be recognized by the Protein Data Bank (PDB). Because crystallographic refinement of a macromolecular structure is narrowly defined as a correspondence from one dataset to one structure. A never-observed dataset reconstituted from a collection of experimental datasets does not match the well-established crystallographic template of PDB; let alone a refinement of crystal structure with the NMR algorithm.

A distance matrix contains *M* pairwise interatomic distances of a structure in the form of Cartesian coordinates of all observed atoms. An everyday example of distance matrix is an intercity mileage chart appended to the road atlas. Differences in the molecular orientation, choice of origin, and crystal lattice among all experimentally determined structures have no contribution to the distance matrices. Due to its symmetry, only the lower triangle is necessary. A far more intimate examination of protein structures in PDB is a direct analysis of their electron density maps instead of the atomic coordinates. *M* such (difference) electron densities, often called voxels in computer graphics, are selected by a mask of interest. In the case of difference maps, only the best refined protein structure in the entire collection supplies a phase set for Fourier synthesis of electron density maps. This best structure is often the ground state structure determined by static crystallography. Other refined atomic coordinates from the PDB entries are not considered in the meta-analysis. That is to say, a meta-analysis of difference electron density maps starts from the X-ray diffraction data archived in PDB rather than the atomic coordinates interpreted from the diffraction data, which removes any potential model bias.

*Singular value decomposition of (difference) electron density maps*

An electron density map, particularly a difference map as emphasized here, consists of density values on an array of grid points within a mask of interest. All *M* grid points in a three-dimensional map can be serialized into a one-dimensional sequence of density values according to a specific protocol. It is not important what the protocol is as long as a consistent protocol is used to serialize all maps of the same grid setting and size, and a reverse protocol is available to erect a three-dimensional map from a sequence of *M* densities. Therefore, a set of *N* serialized maps, also known as vectors in linear algebra, can fill the columns of a data matrix **A** with no specific order, so that the width of **A** is *N* columns, and the length is *M* rows. Often, $M \gg N$, thus **A** is an elongated matrix. If a consistent protocol of serialization is used, the corresponding voxel in all *N* maps occupies a single row of matrix **A**. This strict correspondence in a row of matrix

**A** is important. Changes of the density values in a row from one structure to another are due to either signals, systematic errors, or noises. Although the order of columns in matrix **A** is unimportant, needless to say, the metadata associated with each column must remain in good bookkeeping.

SVD of the data matrix **A** results in $\mathbf{A} = \mathbf{UWV}^\mathrm{T}$, also known as matrix factorization. Matrix **U** has the same shape as **A**, that is, *N* columns and *M* rows. The *N* columns contain decomposed basis components $\boldsymbol{U}_k$, known as left singular vectors of *M* items, where $k = 1, 2, \ldots, N$. Therefore, each component $\boldsymbol{U}_k$ can be erected using the reverse protocol to form a three-dimensional map. This decomposed elemental map can be presented in the same way as the original maps, for example, rendered in molecular graphics software such as Coot and PyMol. It is worth noting that these decomposed elemental maps or map components $\boldsymbol{U}_k$ are independent of any metadata. That is to say, these components remain constant when the metadata vary. Since each left singular vector $\boldsymbol{U}_k$ has a unit length due to the orthonormal property of SVD (see below), that is, $|\boldsymbol{U}_k| = 1$, the root mean squares (rms) of the items in a left singular vector is $1/\sqrt{M}$ that measures the quadratic mean of the items.

The second matrix **W** is a square matrix that contains all zeros except for *N* positive values on its major diagonal, known as singular values $w_k$. The magnitude of $w_k$ is considered as a weight or significance of its corresponding component $\boldsymbol{U}_k$. The third matrix **V** is also a square matrix of $N \times N$. Each column of **V** or row of its transpose $\mathbf{V}^\mathrm{T}$, known as a right singular vector $\boldsymbol{V}_k$, contains the relative compositions of $\boldsymbol{U}_k$ in each of the *N* original maps. Therefore, each right singular vector $\boldsymbol{V}_k$ can be considered as a function of the metadata. Right singular vectors also have the same unit length, that is, $|\boldsymbol{V}_k| = 1$. Effectively, SVD separates the constant components independent of the metadata from the compositions that depend on the metadata.

A singular triplet denotes 1) a decomposed component $\boldsymbol{U}_k$, 2) its singular value $w_k$, and 3) the composition function $\boldsymbol{V}_k$. Singular triplets are often sorted in a descending order of their singular values $w_k$. Only a small number of *n* significant singular triplets identified by the greatest singular values $w_1$ through $w_n$ can be used in a linear combination to reconstitute a set of composite maps that closely resemble the original ones in matrix **A**, where $n < N$. For example, the original map in the *i*th column of matrix **A** under a certain experimental condition can be closely represented by the $i$th composite map $w_1v_{1i}\boldsymbol{U}_1 + w_2v_{2i}\boldsymbol{U}_2 + \ldots + w_nv_{ni}\boldsymbol{U}_n$, where ($v_{1i}$, $v_{2i}$, …) is from the $i$th row of matrix **V**. The coefficient set for the linear combination is redefined here as $c_{ki} = w_kv_{ki}/\sqrt{M}$. The rms of the values in a map component, or the average magnitude measured by the quadratic mean, acts as a constant scale factor that resets the modified coefficients $c_{ki}$ back to the original scale of the core data, such as Å for distance matrices and $e^-/Å^3$ for electron density maps if these units are used in the original matrix **A**. Practically, an electron density value usually carries an arbitrary unit without a calibration, which makes this scale factor unnecessary. In the linear combination $c_{1i}\boldsymbol{U}_1 + c_{2i}\boldsymbol{U}_2 + \ldots + c_{ni}\boldsymbol{U}_n$, each component $\boldsymbol{U}_k$ is independent of the metadata while how much of each component is required for the approximation, that is, $c_{ki}$, depends on the metadata.

Excluding the components after $\boldsymbol{U}_n$ in this approximation is based on an assumption that the singular values after $w_n$ are very small relative to those from $w_1$ through $w_n$. As a result, the structural information evenly distributed in all $N$ original maps is effectively concentrated into a far fewer number of $n$ significant components, known as information concentration or dimension reduction. On the other hand, the trailing components in matrix **U** contain inconsistent fluctuations and random noises. Excluding these components effectively rejects noises (Schmidt et al., 2003). The least-squares property of SVD guarantees that the rejected trailing components sums up to the least squares of the discrepancies between the original core data and the approximation using the accepted components.

However, no clear boundary is guaranteed between signals, systematic errors, and noises. Systematic errors could be more significant than the desired signals. Therefore, excluding some components from 1 through $n$ is also possible. If systematic errors are correctly identified, the reconstituted map without these significant components would no longer carry the systematic errors.

*The orthonormal property of SVD*

The solution set of SVD must guarantee that the columns in **U** and **V**, the left and right singular vectors $\boldsymbol{U}_k$ and $\boldsymbol{V}_k$, are orthonormal, that is, $\boldsymbol{U}_h \bullet \boldsymbol{U}_k = \boldsymbol{V}_h \bullet \boldsymbol{V}_k = 0$ (ortho) and $\boldsymbol{U}_k \bullet \boldsymbol{U}_k = \boldsymbol{V}_k \bullet \boldsymbol{V}_k = 1$ (normal), where $h \neq k$ but both are from 1 to $N$. The orthonormal property also holds for the row vectors. As a result, each component $\boldsymbol{U}_k$ is independent of the other components. In other words, a component cannot be represented by a linear combination of any other components. However, two physical or chemical parameters in the metadata, such as temperature and pH, may cause different changes to a structure. These changes are not necessarily orthogonal. They could exhibit some correlation. Therefore, the decomposed components $\boldsymbol{U}_k$ not necessarily represent any physically or chemically meaningful changes (see below).

Due to the orthonormal property of SVD, an $N$-dimensional Euclidean space is established, and the first $n$ dimensions define its most significant subspace. Each coefficient set $\boldsymbol{c}_i = (c_{1i}, c_{2i}, \ldots, c_{ni})$ of the $i$th composite map is located in this $n$-dimensional subspace. All coefficient sets for $i = 1, 2, \ldots, N$ in different linear combinations to approximate the $N$ original maps in a least-squares sense can be represented by $N$ points or vectors $\boldsymbol{c}_1, \boldsymbol{c}_2, \ldots, \boldsymbol{c}_N$ in the Euclidean subspace. This $n$-dimensional subspace is essentially the conformational space as surveyed by the jointly analyzed core data. The conformational space is presented as scatter plots with each captured structure represented as a dot located at a position determined by the coefficient set $\boldsymbol{c}_i$ of the $i$th observed map. When the subspace has greater dimensionality than two, multiple two-dimensional orthographical projections of the subspace are presented. These scatter plots are highly informative to reveal the relationship between the (difference) electron density maps and their metadata.

If two coefficient sets $\boldsymbol{c}_i \approx \boldsymbol{c}_j$, they are located close to each other in the conformational space. Therefore, these two structures $i$ and $j$ share two similar conformations. Two structures located far apart from each other in the conformational space are distinct in their conformations, and distinct in the compositions of the map components. A reaction trajectory emerges in this conformational space if the temporal order of the core data is experimentally determined (Figs. S5, S7, and S9). Otherwise, an order could be assigned to these structures based on an assumed smoothness of conformational changes along a reaction trajectory (Ren, 2013a, 2013b, 2016). Causation and consequence of structural motions could be revealed from the order of the structures in a series, which may further lead to structural mechanism. In addition, an off-trajectory location in the conformational space or a location between two clusters of observed structures represents a structure in a unique conformation that has never been experimentally captured. Such a hypothetical structure can be refined against a reconstituted distance matrix using molecular distance geometry (Ren, 2013a, 2013b, 2016) or a reconstituted electron density map with the method proposed below.

*Rotation in SVD space*

Dimension reduction is indeed effective in meta-analysis of protein structures when many datasets are evaluated at the same time. However, the default solution set of SVD carries complicated physical and chemical meanings that are not immediately obvious. The interpretation of a basis component $\boldsymbol{U}_k$, that is, “what-does-it-mean”, requires a clear demonstration of the relationship between the core data and their metadata. The outcome of SVD does not guarantee any physical meaning in a basis component. Therefore, SVD alone provides no direct answer to “what-does-it-mean”, thus its usefulness is very limited to merely a mathematical construction. However, the factorized set of matrices **U**, **W**, and **V** from SVD is not a unique solution. That is to say, they are not the only solution to factorize matrix **A**. Therefore, it is very important to find one or more alternative solution sets that are physically meaningful to elucidate a structural interpretation. The concept of a rotation after SVD was introduced by Henry & Hofrichter (Henry and Hofrichter, 1992). But they suggested a protocol that fails to preserve the orthonormal and least-squares properties of SVD. The rotation protocol suggested by Ren incorporates the metadata into the analysis and combines with SVD of the core data. This rotation achieves a numerical deconvolution of multiple physical and chemical factors after a pure mathematical decomposition, and therefore, provides a route to answer the question of “what-does-it-mean” (Ren, 2019). This rotation shall not be confused with a rotation in the three-dimensional real space, in which a molecular structure resides.

A rotation in the $n$-dimensional Euclidean subspace is necessary to change the perspective before a clear relationship emerges to elucidate scientific findings. It is shown below that two linear combinations are identical before and after a rotation applied to both the basis components and their coefficients in a two-dimensional subspace of $h$ and $k$. That is,

$$c_h\boldsymbol{U}_h + c_k\boldsymbol{U}_k = f_h\boldsymbol{R}_h + f_k\boldsymbol{R}_k, \tag{1}$$

where $c_h$ and $c_k$ are the coefficients of the basis components $\boldsymbol{U}_h$ and $\boldsymbol{U}_k$ before the rotation; and $f_h$ and $f_k$ are the coefficients of the rotated basis components $\boldsymbol{R}_h$ and $\boldsymbol{R}_k$, respectively. The same Givens rotation of an angle $\theta$ is applied to both the components and their coefficients:

$$\begin{cases} \boldsymbol{R}_h = \boldsymbol{U}_h\cos\theta - \boldsymbol{U}_k\sin\theta; \\ \boldsymbol{R}_k = \boldsymbol{U}_h\sin\theta + \boldsymbol{U}_k\cos\theta. \end{cases} \quad (2)$$

Obviously, the rotated components $\boldsymbol{R}_h$ and $\boldsymbol{R}_k$ remain mutually orthonormal and orthonormal to other components. And

$$\begin{cases} f_h = s_h t_h = c_h\cos\theta - c_k\sin\theta; \\ f_k = s_k t_k = c_h\sin\theta + c_k\cos\theta. \end{cases} \quad (3)$$

Here $s_{h|k} = \sqrt{\sum f_{h|k}^2}$ are the singular values that replace $w_h$ and $w_k$, respectively, after the rotation. They may increase or decrease compared to the original singular values so that the descending order of the singular values no longer holds. $\boldsymbol{T}_{h|k} = (t_{h|k1}, t_{h|k2}, \ldots, t_{h|kN}) = (f_{h|k1}, f_{h|k2}, \ldots, f_{h|kN})/s_{h|k}$ are the right singular vectors that replace $\boldsymbol{V}_h$ and $\boldsymbol{V}_k$, respectively. $\boldsymbol{T}_h$ and $\boldsymbol{T}_k$ remain mutually orthonormal after the rotation and orthonormal to other right singular vectors that are not involved in the rotation.

Eq. 1 holds because the dot product of two vectors does not change after both vectors rotate the same angle. To prove Eq. 1 in more detail, Eqs. 2 and 3 are combined and expanded. All cross terms of sine and cosine are self-canceled:

$$\begin{aligned} f_h\boldsymbol{R}_h + f_k\boldsymbol{R}_k &= (c_h\cos\theta - c_k\sin\theta)(\boldsymbol{U}_h\cos\theta - \boldsymbol{U}_k\sin\theta) + (c_h\sin\theta + c_k\cos\theta)(\boldsymbol{U}_h\sin\theta + \boldsymbol{U}_k\cos\theta) \\ &= c_h\boldsymbol{U}_h\cos^2\theta + c_k\boldsymbol{U}_k\sin^2\theta + c_h\boldsymbol{U}_h\sin^2\theta + c_k\boldsymbol{U}_k\cos^2\theta \pm c_h\boldsymbol{U}_k\sin\theta\cos\theta \pm c_k\boldsymbol{U}_h\sin\theta\cos\theta \\ &= c_h\boldsymbol{U}_h(\cos^2\theta + \sin^2\theta) + c_k\boldsymbol{U}_k(\sin^2\theta + \cos^2\theta) \\ &= c_h\boldsymbol{U}_h + c_k\boldsymbol{U}_k \end{aligned}$$

A rotation in two-dimensional subspace of $h$ and $k$ has no effect in other dimensions, as the orthonormal property of SVD guarantees. Multiple steps of rotations can be carried out in many two-dimensional subspaces consecutively to achieve a multi-dimensional rotation. A new solution set derived from a rotation retains the orthonormal property of SVD. The rotation in the Euclidean subspace established by

SVD does not change the comparison among the core data of protein structures. Rather it converts one solution set $\mathbf{A} = \mathbf{UWV}^{\mathrm{T}}$ to other alternative solutions $\mathbf{A} = \mathbf{RST}^{\mathrm{T}}$ so that an appropriate perspective can be found to elucidate the relationship between the core data and metadata clearly and concisely.

For example, if one physical parameter could be reoriented along a single dimension *k* but not involving other dimensions by a rotation, it would be convincing to show that the left singular vector $\boldsymbol{U}_k$ of this dimension illustrates the structural impact by this physical parameter. Before this rotation, the same physical parameter may appear to cause structural variations along several dimensions, which leads to a difficult interpretation. Would a proper rotation establish a one-on-one correspondence from all physical or chemical parameters to all the dimensions? It depends on whether each parameter induces an orthogonal structural change, that is, whether structural responses to different parameters are independent or correlated among one another. If structural changes are indeed orthogonal, it should be possible to find a proper rotation to cleanly separate them in different dimensions. Otherwise, two different rotations are necessary to isolate two correlated responses, but one at a time.

For another example, if the observed core datasets form two clusters in the conformational space, a rotation would be desirable to separate these clusters along a single dimension *k* but to align these clusters along other dimensions. Therefore, the component $\boldsymbol{U}_k$ is clearly due to the structural transition from one cluster to the other. Without a proper rotation, the difference between these clusters could be complicated with multiple dimensions involved. A deterministic solution depends on whether a clear correlation exists between the core data and metadata. A proper rotation may require a user decision. A wrong choice of rotation may select a viewpoint that hinders a concise conclusion. However, it would not alter the shape of the reaction trajectory, nor create or eliminate an intrinsic structural feature. A wrong choice of rotation cannot eliminate the fact that a large gap exists between two clusters of observed core datasets except that these clusters are not obvious from that viewpoint. A different rotation may reorient the perspective along another direction. But the structural conclusion would be equivalent. See example of before and after a rotation in (Ren, 2016).

This rotation procedure finally connects the core crystallographic datasets to the metadata of experimental conditions and accomplishes the deconvolution of physical or chemical factors that are not always orthogonal to one another after a mathematical decomposition. SVD analysis presented in this paper employs rotations extensively except that no distinction is made in the symbols of components and coefficients before and after a rotation except in this section. This method is widely applicable in large-scale structural comparisons. Furthermore, Ren rotation after SVD is not limited to crystallography and may impact other fields wherever SVD is used. For example, SVD is frequently applied to spectroscopic data, images, and genetic sequence data.

A preliminary analysis of SVD involving all difference maps (Table S1) shows that the short and long delays are in two different subspaces (Fig. S19). That is to say, difference maps in the time regimes before 10 ps and after 10 ns share no common features. This conclusion cannot be reached without a proper rotation of the components. The separation between the short and long delays into different subspaces can only be viewed clearly from this specific perspective shown in Fig. S19. Therefore, these time regimes are analyzed separately.

*Exponential fitting of SVD coefficients*

The time dependencies of the SVD coefficients $c_1(t)$, $c_3(t)$, and $c_6(t)$ for the long delays > 10 ns are modeled with exponential functions (Fig. S9). Each time-dependent coefficient

$$c_k(t) = b_k + p_k \exp\left(-\frac{t}{\tau_1}\right) + q_k \exp\left(-\frac{t}{\tau_3}\right) + r_k \exp\left(-\frac{t}{\tau_6}\right) \quad (4)$$

where $k = 1$, 3, or 6. The time constant $\tau_k$ associated with the major change in each coefficient models each step in the reaction scheme K → L → $M_1$ → $M_2$. Whether the time-dependent coefficients can be successfully fitted with exponentials is also a good test for a proper rotation. Because the fact that the fractional concentrations of intermediate species can be expressed as exponentials indicate that the reaction follows a regular kinetic scheme.

*Structural refinement against reconstituted dataset*

The linear combination $\Delta\rho(t) = f_1(t)\boldsymbol{R}_1 + f_2(t)\boldsymbol{R}_2 + \ldots + f_n(t)\boldsymbol{R}_n$ after a rotation reconstitutes one of the observed difference maps at a specific time point $t$. This time-dependent difference map depicts an ever-evolving mixture of many excited species. A reconstituted difference map $\Delta\rho(E)$ for a time-independent, pure, excited species E = intermediate I′, I, J′, and J deconvoluted from many mixtures would take the same form except that only one or very few coefficients remain nonzero if a proper rotation has been found (Table S2). In order to take advantage of the mature refinement software for macromolecular structures with extensive stereochemical restraints, a set of structure factor amplitudes is needed. Therefore, it is necessary to reconstitute a set of structure factor amplitudes that would produce the target difference map $\Delta\rho(E)$ based on a known structure at the ground state. First, an electron density map of the structure at the ground state is calculated. This calculated map is used as a base map. Second, this base map of the ground state is combined with the positive and negative densities in the target difference map $\Delta\rho(E)$ so that the electron densities at the ground state are skewed toward the intermediate state. Third, structure factors are calculated from the combined map. Finally, the phase set of the calculated structure factors is discarded, and the amplitudes are used to refine a single conformation of the intermediate species E that $\Delta\rho(E)$ represents.

This protocol following the SVD and Ren rotation of components achieves a refinement of a pure structural species without the need of alternative conformations. Several points are noteworthy. First, the minimization protocol in this refinement is performed against a numerically reconstituted amplitude set that has never been directly measured from a crystal. This reconstituted dataset could be considered as an extrapolated dataset “on steroids” if compared to the traditional extrapolation of small differences, such as, the Fourier coefficient set to calculate a 3Fo-2Fc map, a technique often used to overcome a partial occupancy of an intermediate structure. An extrapolation of small differences is not directly observed either but computed by an exaggeration of the observed difference based on an assumption that the intermediate state is partially occupied, such as the doubling of the observed difference in 3Fo-2Fc = Fo + 2(Fo-Fc). In contrast to the conventional technique of extrapolation, the deconvolution method applied here is an interpolation among many experimental datasets rather than an extrapolation. Secondly, the deconvolution is a simultaneous solution of multiple intermediate states mixed together instead of solving a single excited state.

Second, a map calculated from the ground state structure is chosen as the base map instead of an experimental map such as Fo or 2Fo-Fc map. If the second step of the protocol is skipped, that is, no difference map is combined with the ground state map, the refinement would result in an *R* factor of nearly zero, since the refinement is essentially against the calculated structure factors (bR in Table S2). This is to say, the residuals of the refinement are solely due to the difference component instead of the base map. This is desirable since errors in the static structure of the ground state are gauged during its own refinement. On the other hand, if an experimental map is chosen as a base map, the refinement *R* factors would reflect errors in both the base map and the difference map, which leads to a difficulty in an objective evaluation of this refinement protocol.

Third, the combination of the base map and a difference map is intended to represent a pure intermediate species. Therefore, alternative conformations in structural refinement that model a mixture of species would defeat this purpose. However, this combined map could be very noisy and may not represent a single species without a proper rotation. This is particular the case, if the target difference map $\Delta\rho$ is not derived from an SVD analysis and Ren rotation. The SVD analysis identifies many density components that are inconsistent among all observed difference maps and excludes them, which greatly reduces the noise content. Therefore, this refinement protocol may not be very successful without an SVD analysis. Another source of noise originates from the phase set of the structure factors. Prior to the refinement of the intermediate structure, the phase set remains identical to that of the ground state. This is far from the reality when an intermediate structure involves widespread changes, such as those refined in this study. If the rotation after SVD is not properly selected, the target difference map would remain as a mixture minus the ground state. Therefore, the refinement of a single conformation would encounter difficulty or significant residuals, as judged by the *R* factors, the residual map, and the refined structure. A proper solution to this problem is a better SVD solution by Ren rotation rather than alternative conformations. A successful refinement of near perfect *trans* or *cis* double bonds is a good sign to indicate that the reconstituted amplitude set after a rotation reflects a relatively homogeneous structure. If a double bond could not be refined well to near perfect *trans* or *cis* configuration, the dataset of structure factor amplitudes is likely from a mixture of heterogeneous configurations, which occurred frequently in previous studies of bR and photoactive yellow protein (Jung et al., 2013; Lanyi and Schobert, 2007; Nogly et al., 2018). It has been a great difficulty in crystallographic refinement in general that a heterogeneous mixture of conformations cannot be unambiguously refined even with alternative conformations. This difficulty becomes more severe when a mixture involves more than two conformations or when some conformations are very minor.

Lastly, the refinement protocol proposed here could be carried out in the original unit cell and space group of the crystal at the ground state. However, this is not always applicable as the original goal of the meta-analysis is a joint examination of all available structures from a variety of crystal forms. It would be highly desirable to evaluate difference maps of the same or similar proteins from non-isomorphous crystals together by SVD. Alternatively, the refinement protocol could also be performed in the space group of P1 with a virtual unit cell large enough to hold the structure, which is the option in this study (Table S2). This is to say, the entire analysis of SVD-rotation-refinement presented here could be extracted and isolated from the original crystal lattices, which paves the way to future applications to structural data acquired by experimental techniques beyond crystallography, most attractively, to single particle reconstruction in cryo electron microscopy.

#### Supplementary References


Adams, P.D., Afonine, P.V., Bunkóczi, G., Chen, V.B., Davis, I.W., Echols, N., Headd, J.J., Hung, L.-W., Kapral, G.J., Grosse-Kunstleve, R.W., et al. (2010). PHENIX: a comprehensive Python-based system for macromolecular structure solution. Acta Crystallogr. D Biol. Crystallogr. *D66*, 213–221. https://doi.org/10.1107/S0907444909052925.

Balashov, S.P., Govindjee, R., Kono, M., Imasheva, E., Lukashev, E., Ebrey, T.G., Crouch, R.K., Menick, D.R., and Feng, Y. (1993). Effect of the arginine-82 to alanine mutation in bacteriorhodopsin on dark adaptation, proton release, and the photochemical cycle. Biochemistry *32*, 10331–10343. https://doi.org/10.1021/bi00090a008.

Berman, H.M., Kleywegt, G.J., Nakamura, H., and Markley, J.L. (2012). The Protein Data Bank at 40: Reflecting on the past to prepare for the future. Structure *20*, 391–396. https://doi.org/10.1016/j.str.2012.01.010.

Chandonia, J.-M., and Brenner, S.E. (2006). The impact of structural genomics: expectations and outcomes. Science *311*, 347–351. https://doi.org/10.1126/science.1121018.

Dickopf, S., Alexiev, U., Krebs, M.P., Otto, H., Mollaaghababa, R., Khorana, H.G., and Heyn, M.P. (1995). Proton transport by a bacteriorhodopsin mutant, aspartic acid-85-->asparagine, initiated in the unprotonated Schiff base state. Proc. Natl. Acad. Sci. *92*, 11519–11523. https://doi.org/10.1073/pnas.92.25.11519.

Gerwert, K., Hess, B., Soppa, J., and Oesterhelt, D. (1989). Role of aspartate-96 in proton translocation by bacteriorhodopsin. Proc. Natl. Acad. Sci. *86*, 4943–4947. https://doi.org/10.1073/pnas.86.13.4943.

Glynn, C., and Rodriguez, J.A. (2019). Data-driven challenges and opportunities in crystallography. Emerg. Top. Life Sci. ETLS20180177. https://doi.org/10.1042/ETLS20180177.

Govindjee, R., Misra, S., Balashov, S.P., Ebrey, T.G., Crouch, R.K., and Menick, D.R. (1996). Arginine-82 regulates the pKa of the group responsible for the light-driven proton release in bacteriorhodopsin. Biophys. J. *71*, 1011–1023. https://doi.org/10.1016/S0006-3495(96)79302-5.

Grigorieff, N., Ceska, T.A., Downing, K.H., Baldwin, J.M., and Henderson, R. (1996). Electron-crystallographic refinement of the structure of bacteriorhodopsin. J. Mol. Biol. *259*, 393–421. https://doi.org/10.1006/jmbi.1996.0328.

Hackett, N.R., Stern, L.J., Chao, B.H., Kronis, K.A., and Khorana, H.G. (1987). Structure-function studies on bacteriorhodopsin. V. Effects of amino acid substitutions in the putative helix F. J. Biol. Chem. *262*, 9277–9284. https://doi.org/10.1016/S0021-9258(18)48077-5.

Henry, E.R., and Hofrichter, J. (1992). Singular value decomposition: Application to analysis of experimental data. In Numerical Computer Methods, (Academic Press), pp. 129–192.

Jung, Y.O., Lee, J.H., Kim, J., Schmidt, M., Moffat, K., Šrajer, V., and Ihee, H. (2013). Volume-conserving trans–cis isomerization pathways in photoactive yellow protein visualized by picosecond X-ray crystallography. Nat. Chem. *5*, 212–220. https://doi.org/10.1038/nchem.1565.

Lanyi, J.K., and Schobert, B. (2007). Structural changes in the L photointermediate of bacteriorhodopsin. J. Mol. Biol. *365*, 1379–1392. https://doi.org/10.1016/j.jmb.2006.11.016.

Liebschner, D., Afonine, P.V., Baker, M.L., Bunkóczi, G., Chen, V.B., Croll, T.I., Hintze, B., Hung, L.-W., Jain, S., McCoy, A.J., et al. (2019). Macromolecular structure determination using X-rays, neutrons and electrons: recent developments in Phenix. Acta Crystallogr. Sect. Struct. Biol. *75*, 861–877. https://doi.org/10.1107/S2059798319011471.

Logunov, S.L., El-Sayed, M.A., Song, L., and Lanyi, J.K. (1996). Photoisomerization quantum yield and apparent energy content of the K intermediate in the photocycles of bacteriorhodopsin, its mutants D85N, R82Q, and D212N, and deionized blue bacteriorhodopsin. J. Phys. Chem. *100*, 2391–2398. https://doi.org/10.1021/jp9515242.

Marti, T., Otto, H., Mogi, T., Rösselet, S.J., Heyn, M.P., and Khorana, H.G. (1991). Bacteriorhodopsin mutants containing single substitutions of serine or threonine residues are all active in proton translocation. J. Biol. Chem. *266*, 6919–6927. https://doi.org/10.1016/S0021-9258(20)89590-8.

Mogi, T., Stern, L.J., Marti, T., Chao, B.H., and Khorana, H.G. (1988). Aspartic acid substitutions affect proton translocation by bacteriorhodopsin. Proc. Natl. Acad. Sci. *85*, 4148–4152. https://doi.org/10.1073/pnas.85.12.4148.

Nogly, P., Weinert, T., James, D., Carbajo, S., Ozerov, D., Furrer, A., Gashi, D., Borin, V., Skopintsev, P., Jaeger, K., et al. (2018). Retinal isomerization in bacteriorhodopsin captured by a femtosecond x-ray laser. Science *361*, eaat0094. https://doi.org/10.1126/science.aat0094.

Otto, H., Marti, T., Holz, M., Mogi, T., and Stern, L.J. (1990). Substitution of amino acids Asp-85, Asp-212, and Arg-82 in bacteriorhodopsin affects the proton release phase of the pump and the pK of the Schiff base. Proc. Natl. Acad. Sci. *87*, 1018–1022. https://doi.org/10.1073/pnas.87.3.1018.

Ren, Z. (2013a). Reaction trajectory revealed by a joint analysis of Protein Data Bank. PLoS ONE *8*, e77141. https://doi.org/10.1371/journal.pone.0077141.

Ren, Z. (2013b). Reverse engineering the cooperative machinery of human hemoglobin. PLoS ONE *8*, e77363. https://doi.org/10.1371/journal.pone.0077363.

Ren, Z. (2016). Molecular events during translocation and proofreading extracted from 200 static structures of DNA polymerase. Nucleic Acids Res. *6*, 1–13. https://doi.org/10.1093/nar/gkw555.

Ren, Z. (2019). Ultrafast structural changes decomposed from serial crystallographic data. J. Phys. Chem. Lett. *10*, 7148–7163. https://doi.org/10.1021/acs.jpclett.9b02375.

Ren, Z. (2022). Photoinduced isomerization sampling of retinal in bacteriorhodopsin. PNAS Nexus *1*. https://doi.org/10.1093/pnasnexus/pgac103.

Ren, Z., Perman, B., Srajer, V., Teng, T.-Y., Pradervand, C., Bourgeois, D., Schotte, F., Ursby, T., Kort, R., Wulff, M., et al. (2001). A molecular movie at 1.8 Å resolution displays the photocycle of photoactive yellow protein, a eubacterial blue-light receptor, from nanoseconds to seconds. Biochemistry *40*, 13788–13801. https://doi.org/10.1021/bi0107142.

Ren, Z., Chan, P.W.Y., Moffat, K., Pai, E.F., Royer, W.E., Šrajer, V., and Yang, X. (2013). Resolution of structural heterogeneity in dynamic crystallography. Acta Cryst *D69*, 946–959. https://doi.org/10.1107/S0907444913003454.

Schaffer, J.E., Kukshal, V., Miller, J.J., Kitainda, V., and Jez, J.M. (2021). Beyond X-rays: an overview of emerging structural biology methods. Emerg. Top. Life Sci. ETLS20200272. https://doi.org/10.1042/ETLS20200272.

Schmidt, M., Rajagopal, S., Ren, Z., and Moffat, K. (2003). Application of singular value decomposition to the analysis of time-resolved macromolecular X-ray data. Biophys. J. *84*, 2112–2129. https://doi.org/10.1016/S0006-3495(03)75018-8.

Schmidt, M., Graber, T., Henning, R., and Srajer, V. (2010). Five-dimensional crystallography. Acta Crystallogr. A *66*, 198–206. https://doi.org/10.1107/S0108767309054166.

Song, L., El-Sayed, M.A., and Lanyi, J.K. (1993). Protein catalysis of the retinal subpicosecond photoisomerization in the primary process of bacteriorhodopsin photosynthesis. Science *261*, 891–894. https://doi.org/10.1126/science.261.5123.891.

Šrajer, V., Ren, Z., Teng, T.-Y., Schmidt, M., Ursby, T., Bourgeois, D., Pradervand, C., Schildkamp, W., Wulff, M., and Moffat, K. (2001). Protein conformational relaxation and ligand migration in myoglobin: A nanosecond to millisecond molecular movie from time-resolved Laue X-ray diffraction. Biochemistry *40*, 13802–13815. https://doi.org/10.1021/bi010715u.

Stoeckenius, W. (1999). Bacterial rhodopsins: Evolution of a mechanistic model for the ion pumps. Protein Sci. *8*, 447–459. https://doi.org/10.1110/ps.8.2.447.

Stoeckenius, W., Lozier, R.H., and Bogomolni, R.A. (1979). Bacteriorhodopsin and the purple membrane of halobacteria. Biochim. Biophys. Acta BBA - Rev. Bioenerg. *505*, 215–278. https://doi.org/10.1016/0304-4173(79)90006-5.

Tittor, J., Schweiger, U., Oesterhelt, D., and Bamberg, E. (1994). Inversion of proton translocation in bacteriorhodopsin mutants D85N, D85T, and D85,96N. Biophys. J. *67*, 1682–1690. https://doi.org/10.1016/S0006-3495(94)80642-3.

Tsin, A., Betts-Obregon, B., and Grigsby, J. (2018). Visual cycle proteins: Structure, function, and roles in human retinal disease. J. Biol. Chem. *293*, 13016–13021. https://doi.org/10.1074/jbc.AW118.003228.

Ursby, T., and Bourgeois, D. (1997). Improved estimation of structure-factor difference amplitudes from poorly accurate data. Acta Crystallogr. A *53*, 564–575. https://doi.org/10.1107/S0108767397004522.

Zimanyi, L., Varo, G., Chang, M., Ni, B., Needleman, R., and Lanyi, J.K. (1992). Pathways of proton release in the bacteriorhodopsin photocycle. Biochemistry *31*, 8535–8543. https://doi.org/10.1021/bi00151a022.

#### Supplementary Tables

Table S1. Datasets analyzed in this work

| Publication | PDB | Label | Resolution | Main conclusions | New findings in this work |
|---|---|---|---|---|---|
| Nango et al. Science 354, 1552, 2016 | 5b6v | dark5 | 1.77 Å | A newly ordered water on the CP side bridges the SB to Thr89 after isomerization, thus further conducts a proton to Asp85. | These datasets contribute to the overdetermination of the structures K $\rightarrow$ L $\rightarrow$ $M_1$ $\rightarrow$ $M_2$. The early creased retinal is flattened; $C_{20}$ methyl group returns into the original plane of the resting retinal as K transitions to L. The most flattened chromophore is achieved in L. The EC half channel contracts in L while the CP half channel starts to open. Reprotonation of the SB starts in L. Several ordered waters are observed in the enlarged CP half channel in $M_1$. The SB move toward inboard in $M_1$ and $M_2$. |
| | 5b6w | 16ns | 2.0 Å | | |
| | 5h2h | 40ns | 2.0 Å | | |
| | 5h2i | 110ns | 2.0 Å | | |
| | 5h2j | 290ns | 2.0 Å | | |
| | 5b6x | 760ns | 2.0 Å | | |
| | 5h2k | 2μs | 2.0 Å | | |
| | 5h2l | 5.25μs | 2.0 Å | | |
| | 5h2m | 13.8μs | 2.0 Å | | |
| | 5b6y | 36.2μs | 2.0 Å | | |
| | 5h2n | 95.2μs | 2.0 Å | | |
| | 5h2o | 250μs | 2.0 Å | | |
| | 5h2p | 657μs | 2.0 Å | | |
| | 5b6z | 1.725ms | 2.0 Å | | |
| Nogly et al. Science 361, eaat0094, 2018 | 6g7h | dark6 | 1.5 Å | Retinal fully isomerizes at 10 ps. But the SB water dissociates earlier. | The short-delay datasets contribute to the structures of I′ $\rightarrow$ I $\rightarrow$ J′ $\rightarrow$ J. Photoisomerization in J′; retinal binding pocket expansion before 1 ps in I and contraction at 10 ps in J |
| | 6g7i | 49-406fs | 1.9 Å | | |
| | 6g7j | 457-646fs | 1.9 Å | | |
| | 6g7k | 10ps | 1.9 Å | | |
| | 6g7l | 8.33ms | 1.9 Å | | |
| Kovacs et al. Nat. Commun. 10, 3177, 2019 | 6ga1 | dark1 | 1.7 Å | The exceedingly high power density of the pump laser causes two-photon absorption. Vibrational motions were observed. | The sub-ps datasets exhibit extensive vibrations at various frequencies. The vibrational signals are widespread over the entire bR molecule and not associated with any structural elements. Therefore, it is concluded that these global vibrations are intrinsic properties of bR induced by short laser pulses. The vibrational signals are more prominent under higher power density of the laser pulses. However, these vibrations are irrelevant to the light-driven proton pumping function of bR. |
| | 6ga2 | dark2 | 1.8 Å | | |
| | 6rmk | dark3 | 1.8 Å | | |
| | 6ga7 | 240fs | 1.8 Å | | |
| | 6ga8 | 330fs | 1.8 Å | | |
| | 6ga9 | 390fs | 1.8 Å | | |
| | 6gaa | 430fs | 1.8 Å | | |
| | 6gab | 460fs | 1.8 Å | | |
| | 6gac | 490fs | 1.8 Å | | |
| | 6gad | 530fs | 1.8 Å | | |
| | 6gae | 560fs | 1.8 Å | | |
| | 6gaf | 590fs | 1.8 Å | | |
| | 6gag | 630fs | 1.8 Å | | |
| | 6gah | 680fs | 1.8 Å | | |
| | 6gai | 740fs | 1.8 Å | | |
| | 6ga4 | 1ps | 1.8 Å | | |
| | 6ga5 | 3ps | 1.9 Å | | |
| | 6ga6 | 10ps | 1.8 Å | | |
| | 6ga3 | 33ms | 2.1 Å | | |
| Weinert et al. Science 365, 61, 2019 | 6rqp | dark | 1.8 Å | Helices E and F swing outward after 10 ms. | These datasets contribute to the overdetermination of $M_2$, in which an open CP half channel is observed. The SB moves toward inboard. |
| | 6rnj | 0-5ms | 2.6 Å | | |
| | 6rph | 10-15ms | 2.6 Å | | |
| | 6rqo | activated | 2.0 Å | | |

Table S2. Refinement statistics

| Intermediate | | bR | I′ | I | J′ | J | K | L | $M_1$ | $M_2$ |
|---|---|---|---|---|---|---|---|---|---|---|
| Time period | | 0- | < 50 fs | 40-700 fs | 0.5-2 ps | 1-30 ps | 3 ps-2 µs | 1-100 µs | 10-1000 µs | 0.5-5 ms |
| PDB-Dev* entry | | PDBDEV_00000129 | | 138 | 139 | 140 | 144 | 145 | 146 | 147 |
| Short delay components | $c_{10}$ | | | 3,300 | | -4,200 | | | | |
| | $c_{14}$ | | 2,000 | 2,700 | 2,700 | 2,000 | | | | |
| | $c_{17}$ | | 3,000 | | -1,300 | -300 | | | | |
| Long delay components | $c_1$ | | | | | | -700 | | 2,200 | 1,000 |
| | $c_2$ | | | | | | 1,800 | 1,800 | 1,800 | 1,200 |
| | $c_3$ | | | | | | -1,000 | 2,100 | 1,500 | 900 |
| | $c_6$ | | | | | | | -600 | | 1,500 |
| Starting model | | PDB 6g7h | | | | | | | | |
| Resolution range | | 50-2.1 Å | | | | | | | | |
| Space group | | P1 | | | | | | | | |
| Unit cell | | $a$ = 62.32 Å; $b$ = 62.32 Å; $c$ = 111.10 Å; $\alpha$ = 90°; $\beta$ = 90°; and $\gamma$ = 120° | | | | | | | | |
| Unique reflections | | 80,354 in working set + 4,236 in test set = 84,590 total | | | | | | | | |
| Completeness | | 95% in working set + 5% in test set = 100% reconstituted | | | | | | | | |
| $R$ (%) | | 1.8 | 29.4 | 31.0 | 29.1 | 30.0 | 26.0 | 27.7 | 28.4 | 28.5 |
| $R_{free}$ (%) | | 1.9 | 31.1 | 32.4 | 30.4 | 30.7 | 26.3 | 28.8 | 29.6 | 29.7 |
| Refined content | | 230 protein residues + 1 retinal + water molecules | | | | | | | | |
| Number of atoms | | 1,798 | 1,795 | 1,798 | 1,796 | 1,795 | 1,798 | 1,794 | 1,799 | 1,796 |
| Water molecules | | 8 | 5 | 8 | 6 | 5 | 8 | 4 | 9 | 6 |
| RMSD bonds (Å) | | 0.005 | 0.009 | 0.009 | 0.009 | 0.009 | 0.009 | 0.009 | 0.009 | 0.009 |
| RMSD angles (°) | | 0.793 | 1.206 | 1.105 | 1.085 | 1.068 | 1.006 | 1.074 | 1.120 | 1.081 |
| Rama. favored (%) | | 98.7 | 96.5 | 95.6 | 96.1 | 96.5 | 97.8 | 96.5 | 97.4 | 96.1 |
| Rama. outliers (%) | | 0.0 | 0.0 | 0.4 | 0.4 | 0.4 | 0.4 | 0.0 | 0.0 | 0.4 |
| Clash score | | 4 | 9 | 5 | 4 | 6 | 6 | 7 | 10 | 7 |

*pdb-dev.wwpdb.org

#### Supplementary Figures and Legends

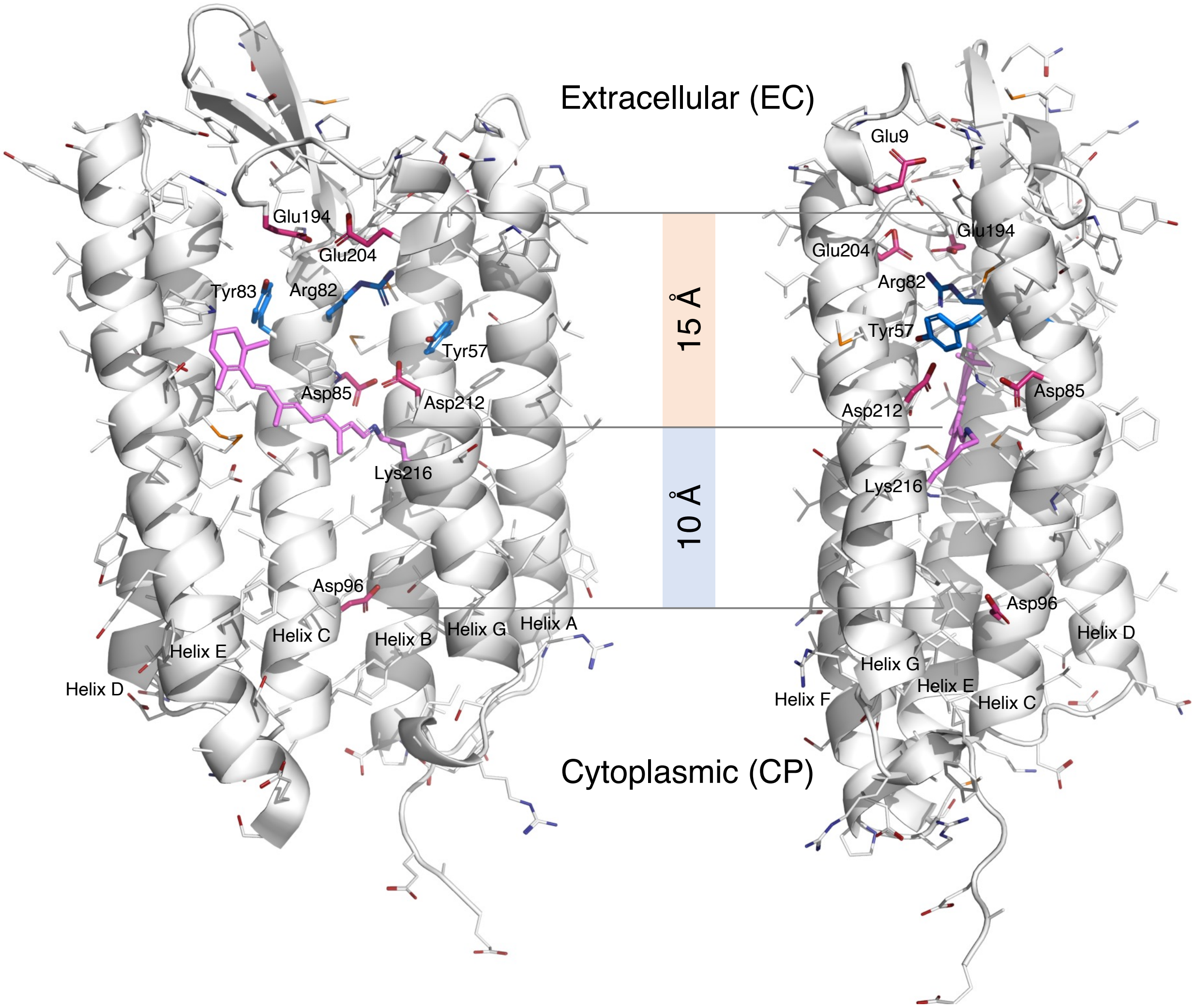


Figure S1. Orthographical views of EC and CP half channels. Parts of the structure are omitted to reveal the interior. The acidic residues in the half channels are highlighted in red. Those in the EC half channel are deprotonated. Asp96 in the hydrophobic CP half channel could be protonated (Gerwert et al., 1989). Arg82 in blue is protonated at neutral pH to form a positive charged guanidinium ion with a p$K_a$ of 12.5. Tyr57 and 83 with a p$K_a$ of 10 in blue could be deprotonated at alkaline pH.

#### Ren: Concentration-driven proton conductance

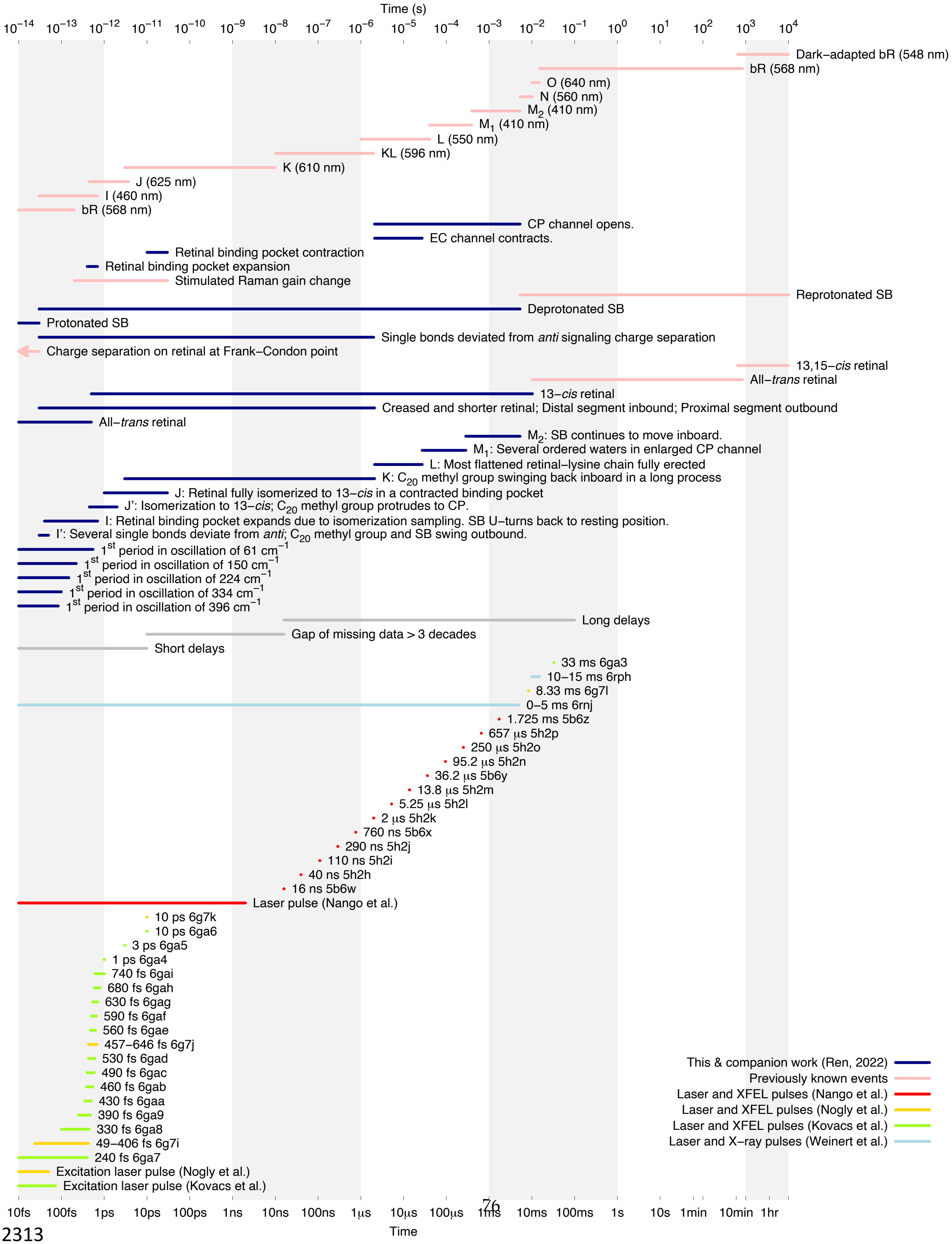




2313

Figure S2. Gantt chart summarizing analyzed datasets and results.

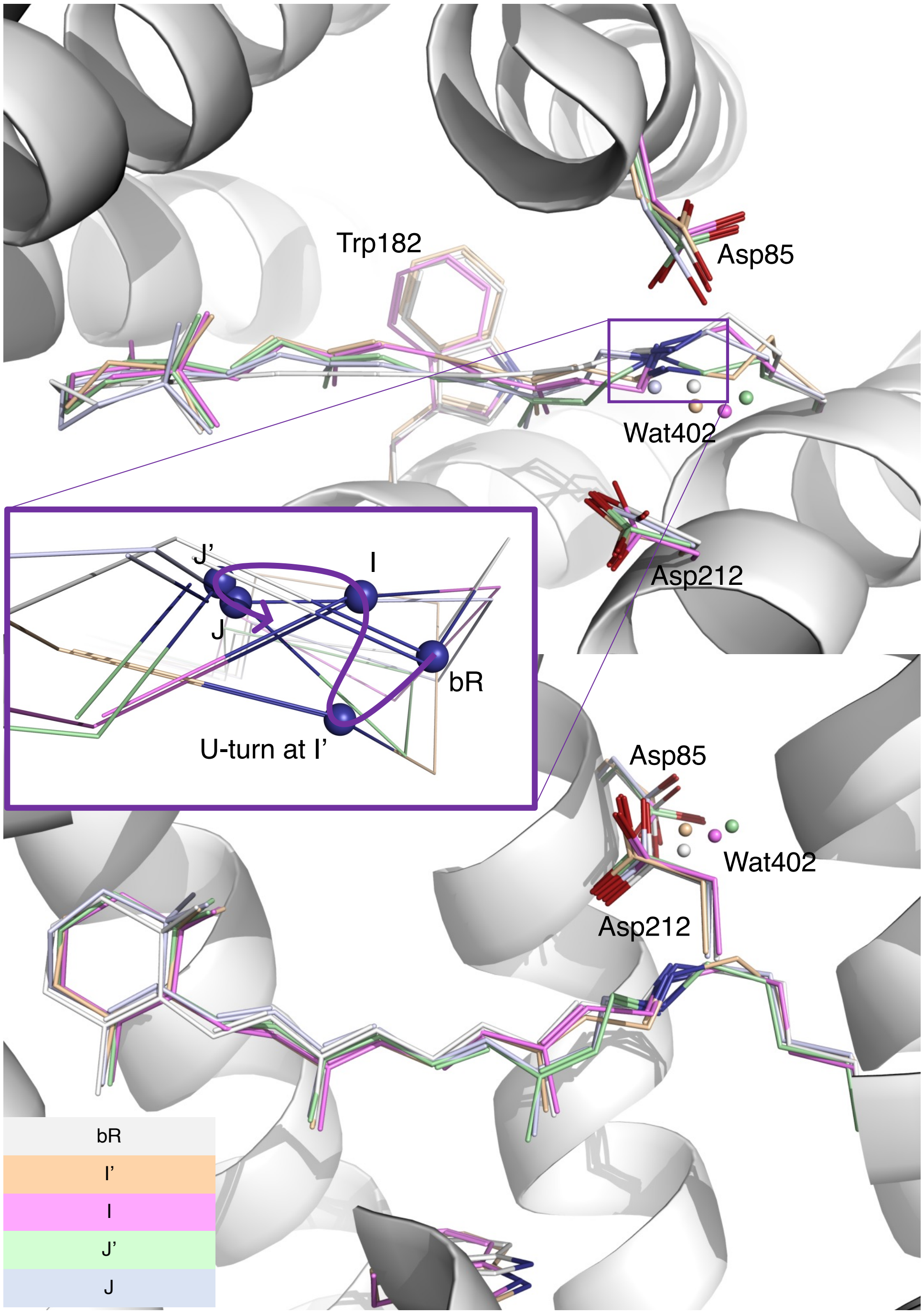


Figure S3. Two orthographical views of early intermediates. The refined conformations of the retinal and several key side chains are rendered in stick models of various colors. These early intermediates (Ren, 2022) are compared with the resting state in white ribbon and stick model. Wat402 is rendered in small spheres. A zoom-in view of the SB is presented in the inset. N$_\zeta$ atom is shown in blue spheres. Its trajectory marked by a spline curve shows a sharp U-turn at I′ state. The SB moves toward the distal direction during I and J due to a creased retinal (Fig. 2b).

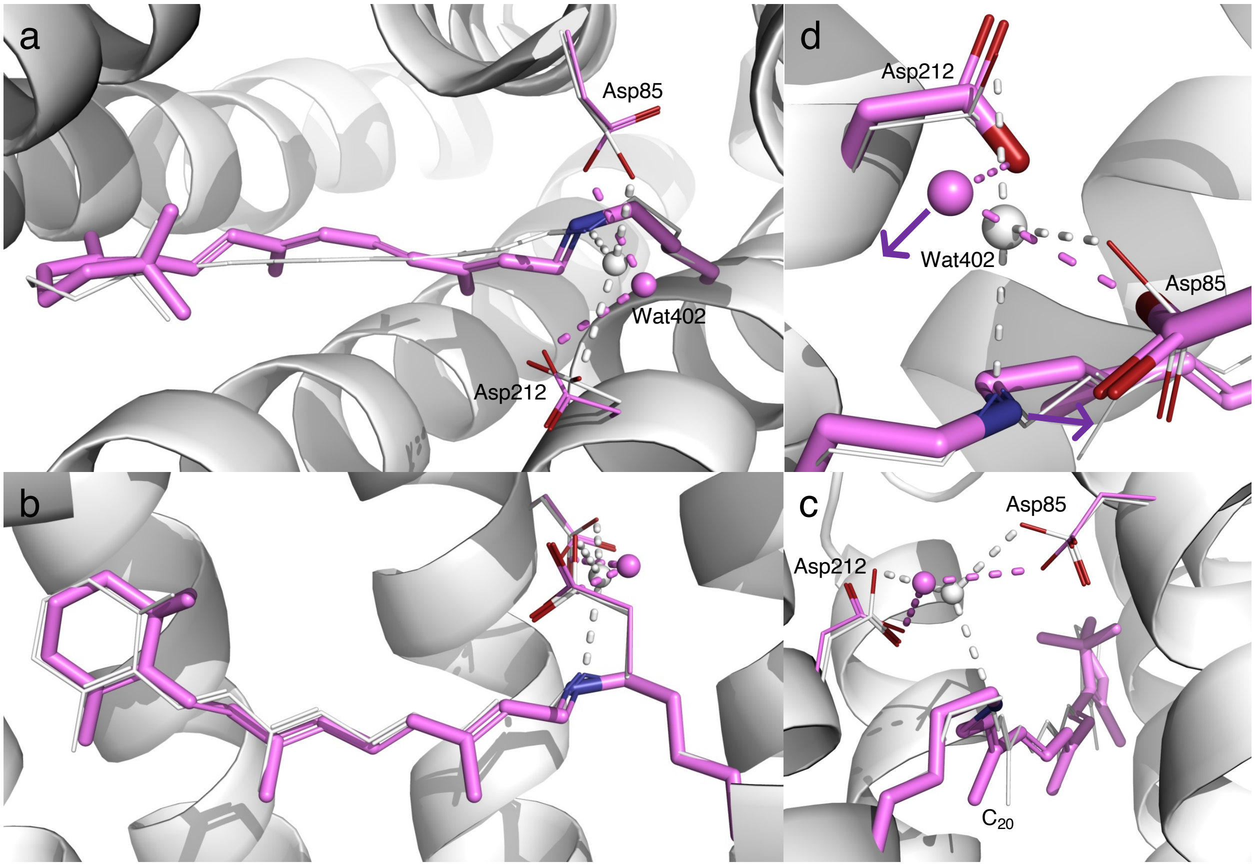


Figure S4. Refined structure of I state. (a, b, and c) Three orthographical views of retinal in I state (Ren, 2022). The refined retinal conformation in purple is compared with the resting state in white. The SB is pointing toward inboard with a perfect *syn* conformation at the single bond $N_\zeta$-$C_\varepsilon$. $C_{20}$ methyl group remains tilted toward outboard. Wat402 with good electron density is displaced toward outboard and remains H-bonded with both Asp85 and 212. (d) Zoom-in view of SB and Wat402. Wat402 is 3.7 Å away from the SB $N_\zeta$. The would-be H-bond directions of the SB and Wat402 are pointing toward the opposite directions as marked by the arrows. This geometry makes not only a H-bond impossible but also any proton transfer difficult.

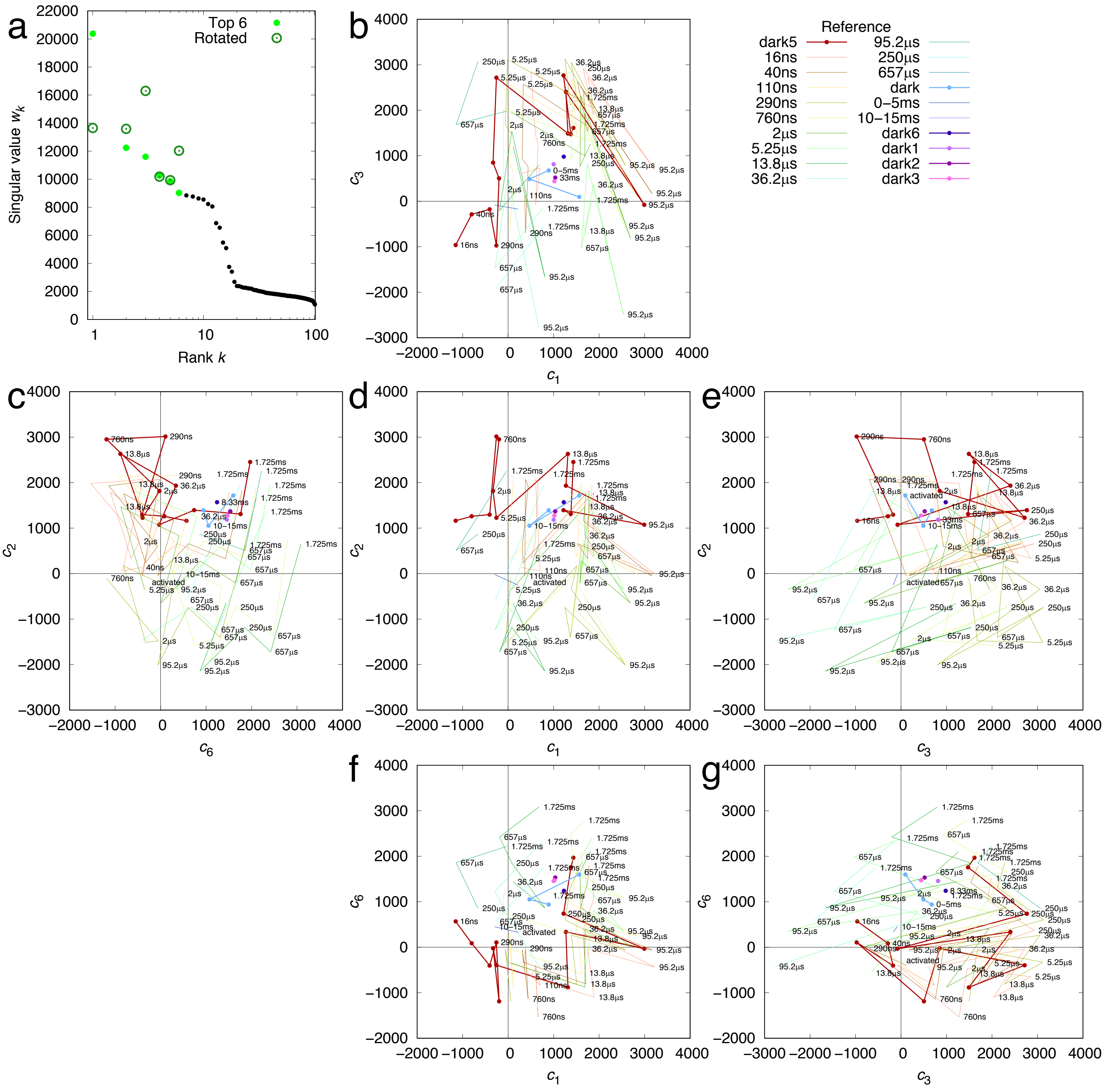


Figure S5. SVD analysis of difference Fourier maps at long delays > 10 ns. SVD analyses of difference Fourier maps result in time-dependent coefficients $c_k(t)$, where $k$ =

1, 2, …, each corresponding to a time-independent components $\boldsymbol{U}_k$. Each raw difference map at a time delay $t$ can be closely represented by a linear combination of these components, $c_1(t)\boldsymbol{U}_1 + c_2(t)\boldsymbol{U}_2 + \ldots$, that is called a reconstituted difference map. Each of these components $\boldsymbol{U}_k$ and the reconstituted difference maps can be rendered in the same way as an observed difference map. The coefficient set $c_k(t)$ is therefore a trace of the photocycle trajectory, when these time-dependent functions are plotted in a multi-dimensional space or plotted together against the common variable $t$. See Methods for detail. (a) Singular values before and after rotation (Ren, 2019, 2022). 19 components stand out. (b-g) Time-dependent coefficients of four components $\boldsymbol{U}_1$, $\boldsymbol{U}_2$, $\boldsymbol{U}_3$, and $\boldsymbol{U}_6$. Each pair of the adjacent orthographical views, horizontal or vertical, can be folded along a straight line between them to erect a three-dimensional space. Each colored trace represents difference maps in a time series calculated with a common reference. Those time series with a dark reference are plotted with thick lines with dots. Other series using a light dataset as a reference are plotted with thin lines without dots.

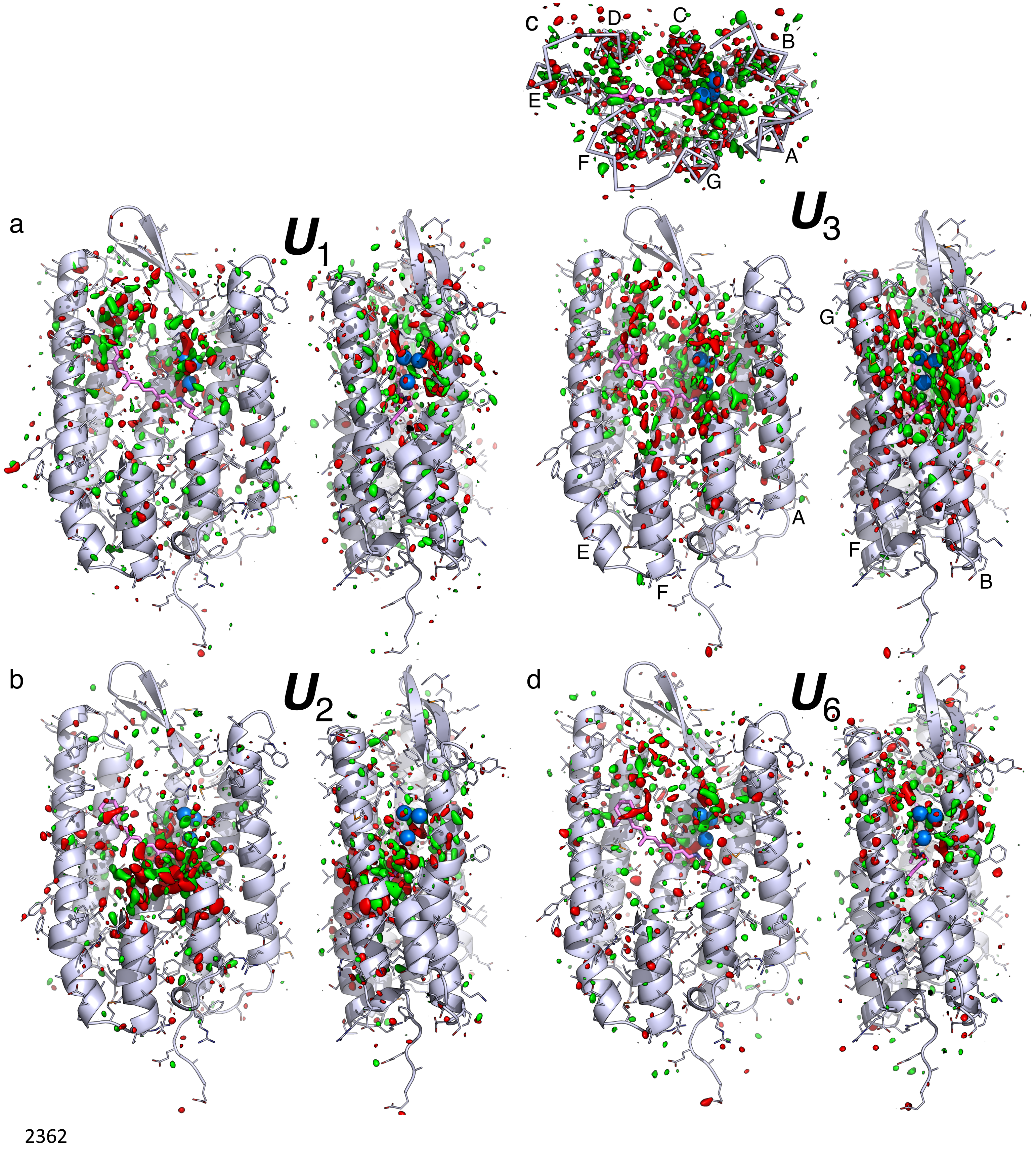
c
D
C
B
E
A
F
G
a
$U_1$
$U_3$
G
A
E
F
F
B
b
$U_2$
d
$U_6$
2362

Figure S6. Component maps $\boldsymbol{U}_1$, $\boldsymbol{U}_2$, $\boldsymbol{U}_3$, and $\boldsymbol{U}_6$ from long delays. The main chain and side chains of the protein are rendered with ribbons and sticks, respectively. The retinal and Lys216 are in purple sticks. Several key waters are in blue spheres. Parts of the structure are omitted to reveal more of the interior. The maps are contoured at $\pm 3\sigma$ in green and red, respectively. (a) Two orthographical views of $\boldsymbol{U}_1$. The signals are distributed over the EC half of the molecule. (b) Two orthographical views of $\boldsymbol{U}_2$. The signals are concentrated around the proximal segment of the retinal. (c) Three orthographical views of $\boldsymbol{U}_3$. Widespread signals are associated with all seven helices. (d) Two orthographical views of $\boldsymbol{U}_6$. The signals are over the EC half of the molecule.

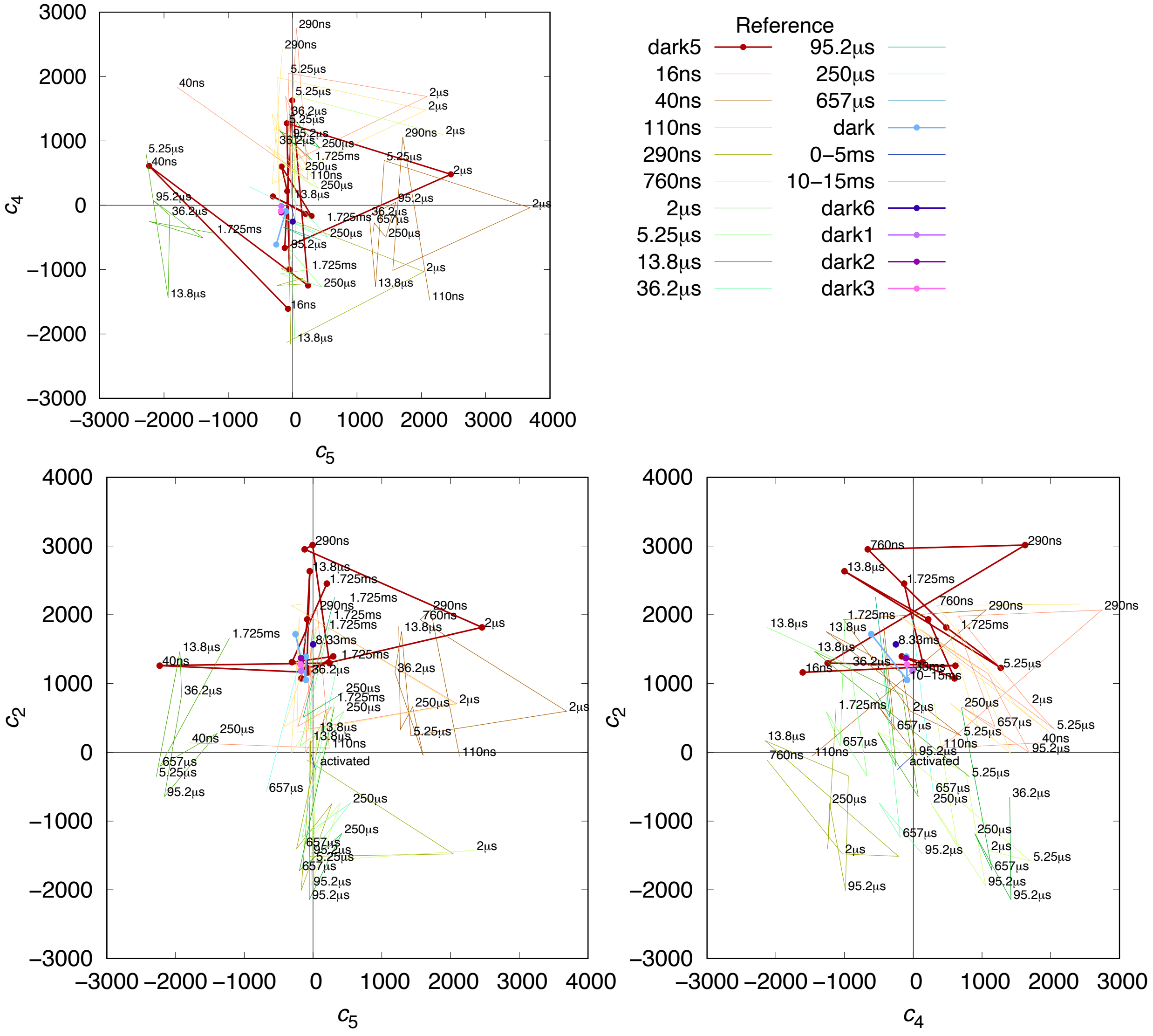


Figure S7. Two components of long delays without clear time dependency. Each pair of the adjacent orthographical views, horizontal or vertical, can be folded along a straight line between them to erect a three-dimensional space. Each colored trace represents difference maps in a time series calculated with a common reference. Those time series with a dark reference are plotted with thick lines with dots. Other series using a light dataset as a reference are plotted with thin lines without dots. $c_4$ fluctuates around zero without a clear time dependency could be caused by inconsistency among difference experiments. $c_5$ is nonzero only for two datasets of 40 ns and 2 μs. The corresponding map components are displayed in Fig. S8. These components are not used in structure refinement. This is an example to demonstrate how SVD identifies and isolates inconsistent systematic errors.

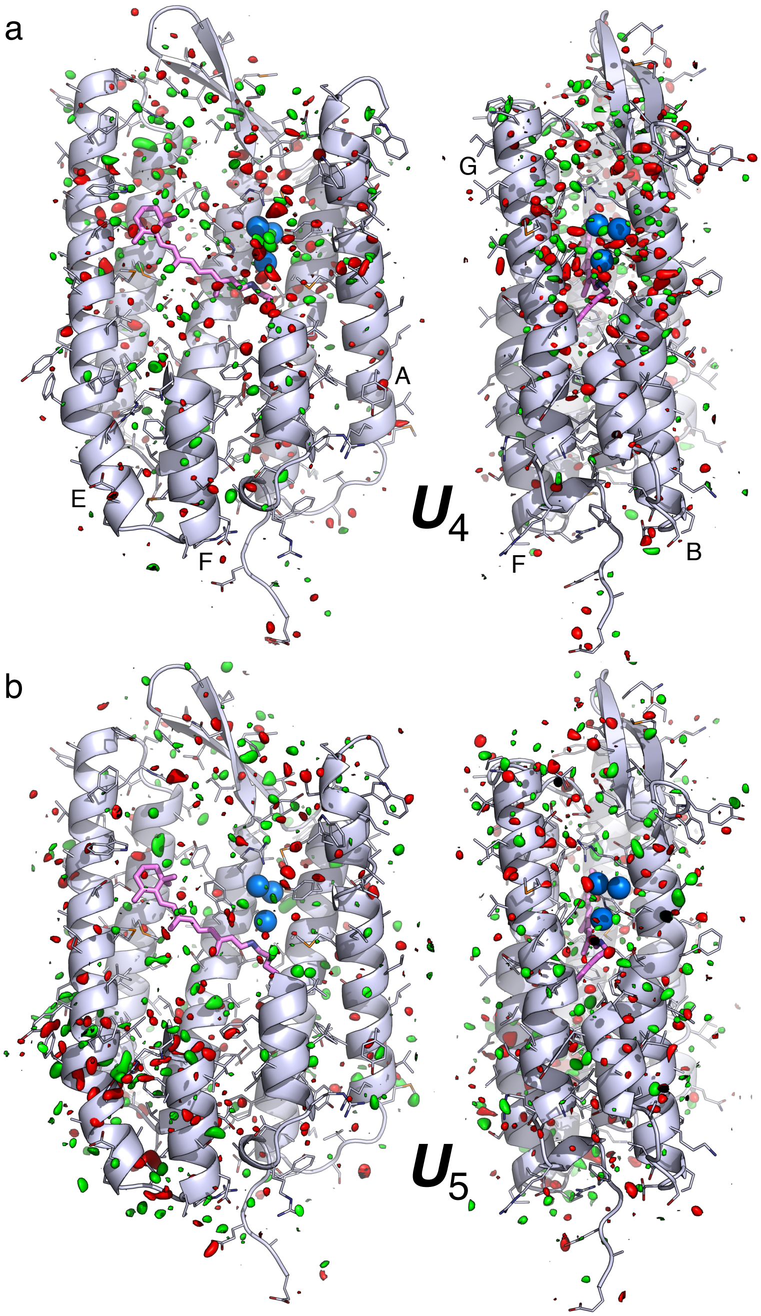


Figure S8. Component maps $\boldsymbol{U}_4$ and $\boldsymbol{U}_5$ from long delays. The main chain and side chains of the protein are rendered with ribbons and sticks, respectively. The retinal and Lys216 are in purple sticks. Several key waters are in blue spheres. Parts of the structure are omitted to reveal more of the interior. The maps are contoured at $\pm 3\sigma$ in green and red, respectively. These components are not associated with a clear time dependency. Therefore, they are isolated but not used in structure refinement. (a) Two orthographical views of $\boldsymbol{U}_4$. Some signals are distributed over the EC half of the molecule. (b) Two orthographical views of $\boldsymbol{U}_5$. Some signals are located around the CP segments of helices E and F.

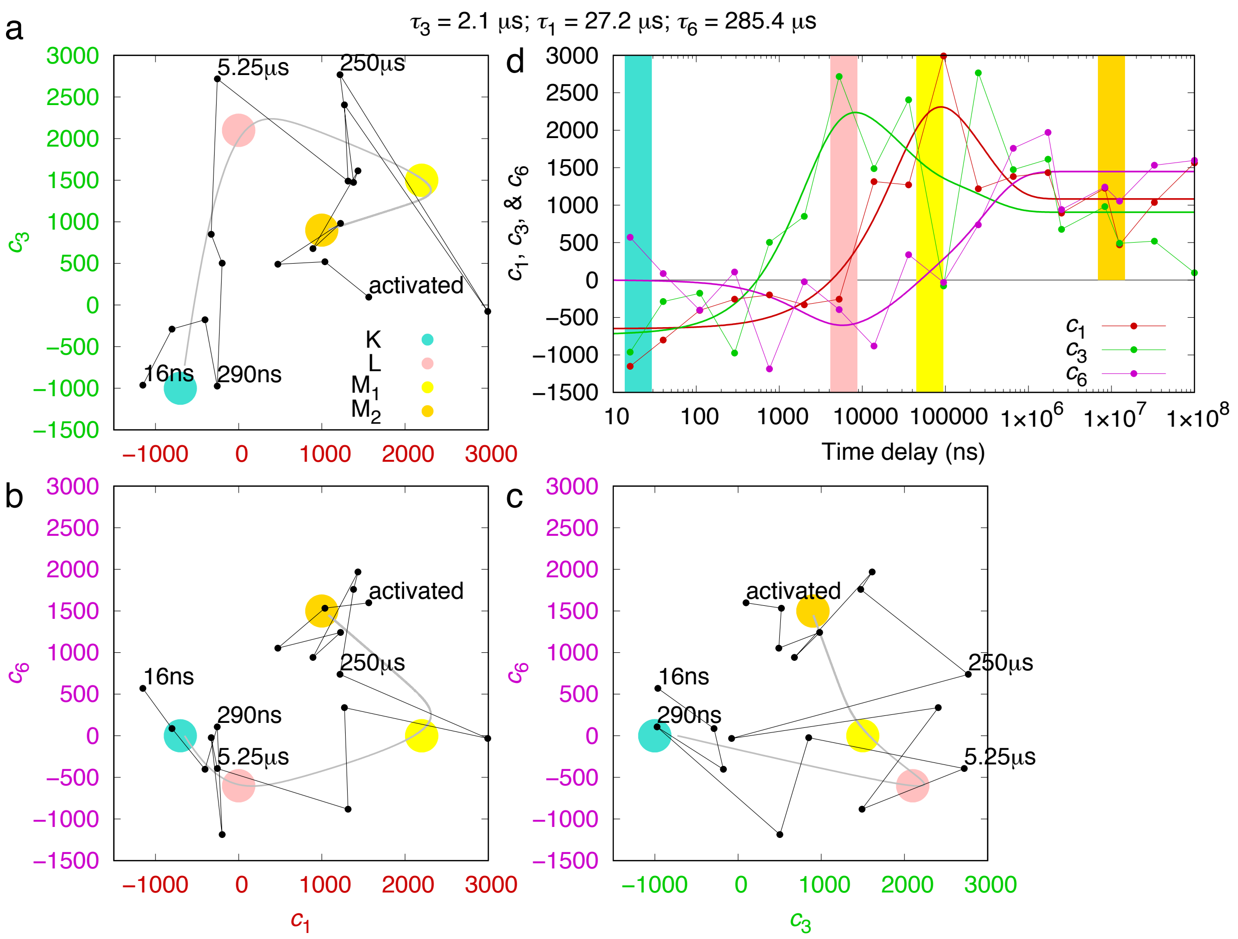


Figure S9. Exponential fitting of SVD coefficients of long delays. (a, b, and c) The time dependencies of the SVD coefficients $c_1(t)$, $c_3(t)$, and $c_6(t)$ of the long delays are modeled with exponential functions (Methods). The time constant $\tau_k$ associated with the major change in each coefficient models a step in the reaction scheme K → L → $M_1$ → $M_2$. $\tau_3$ = 2.1±1 μs or a rate of 470 $ms^{-1}$ is the rate of increase in $c_3(t)$. $\tau_1$ = 27±10 μs or a rate of 37 $ms^{-1}$ describes the jump in $c_1(t)$. $\tau_6$ = 290±100 μs or a rate of 4 $ms^{-1}$ seems to separate $M_1$ and $M_2$. The fitted exponential functions are plotted as the gray curves through the black dots derived from SVD in three orthographical views of the three-dimensional subspace of $c_1$, $c_3$, and $c_6$. The large colored dots mark the relatively pure states of K, L, $M_1$, and $M_2$. (d) SVD coefficients and their exponential fittings as functions of time. The relatively pure states are marked as color bars at the approximate time. The result of exponential fitting shows the time-dependency clearer. The second component map $\boldsymbol{U}_2$ seems to be a constant component without a time-dependent trend (Fig. S5cde). It describes significant changes near the SB and the retinal anchor (Figs. 4ab and S6b). The third component map $\boldsymbol{U}_3$ increases sharply and consistently around 1 μs (Figs. S5beg). This component contains widespread signals associated with all seven helices (Fig. S6c), therefore describes the global changes during K $\rightarrow$ L transition. The first component $\boldsymbol{U}_1$ increases at 10 μs (Figs. S5bdf), which could reflect the changes during L $\rightarrow$ M transition. $\boldsymbol{U}_1$ clearly contains signals over the EC half of the molecule (Fig. S6a), which indicates events related to the proton release. Similarly, the sixth component $\boldsymbol{U}_6$ also shows additional signals on the EC half to a lesser extent (Fig. S6d). This component seems to increase at a rather late time around hundreds of μs (Figs. S5cfg), which splits the M state into $M_1$ and $M_2$.

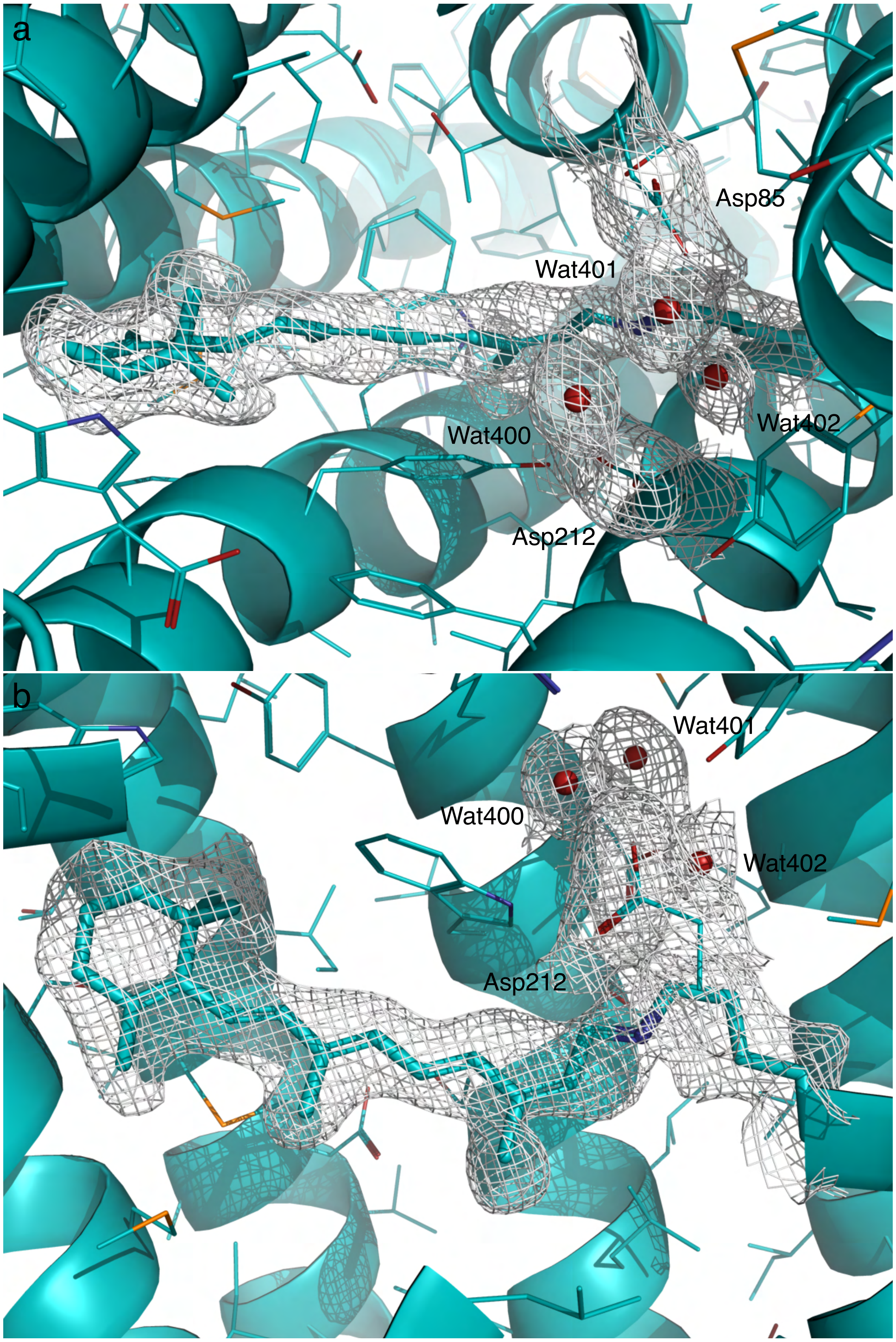


Figure S10. Two orthographical views of the 2Fo-Fc map of K contoured at 6$\sigma$. Here Fo is the reconstituted structure factor amplitudes rather than observed amplitudes (Table S2). Fc is the structure factor amplitudes calculated from the refined structure (Methods). The same applies to the other 2Fo-Fc maps.

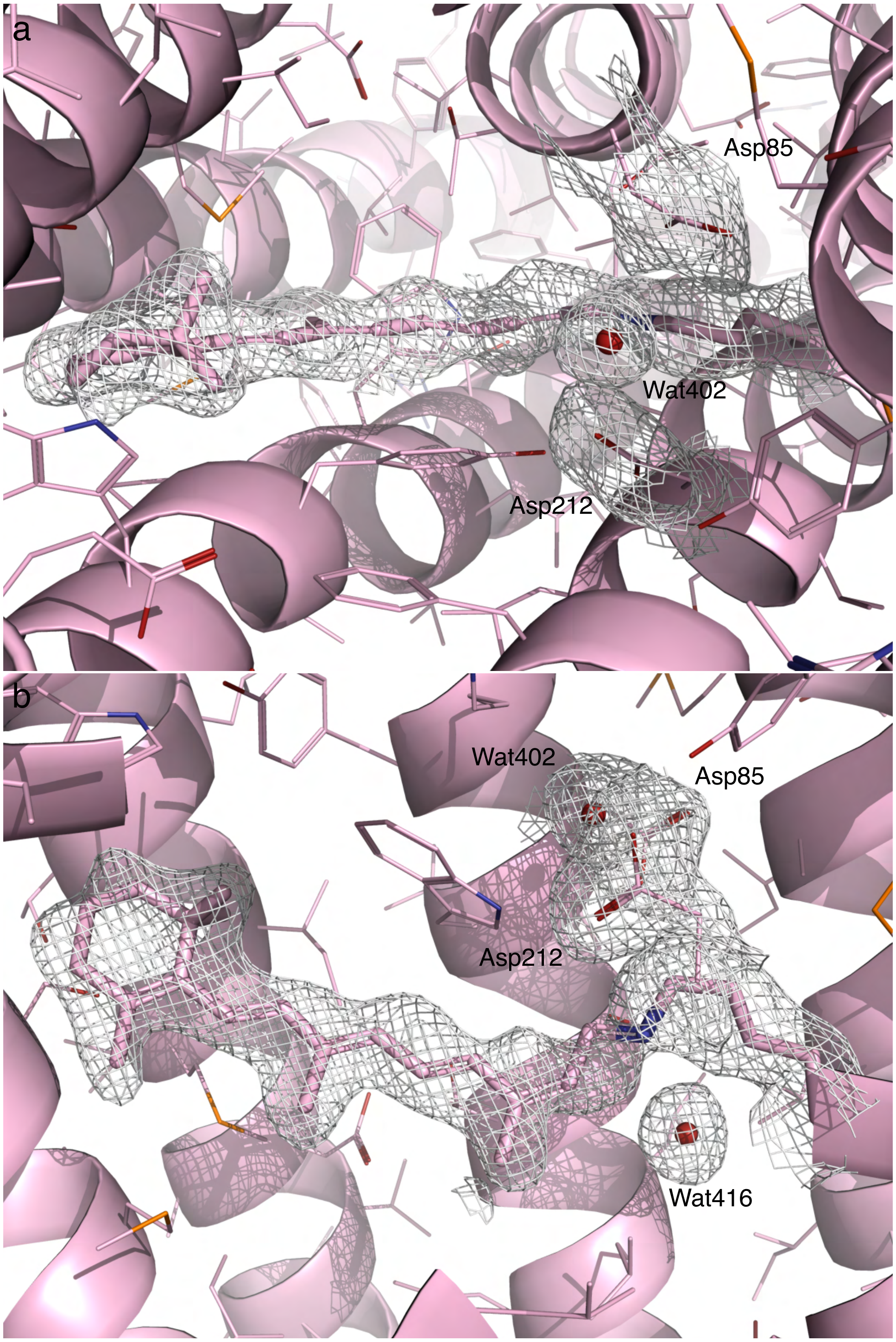


Figure S11. Two orthographical views of the 2Fo-Fc map of L contoured at 3.5$\sigma$. Here Fo is the reconstituted structure factor amplitudes rather than observed amplitudes (Table S2). Fc is the structure factor amplitudes calculated from the refined structure (Methods).

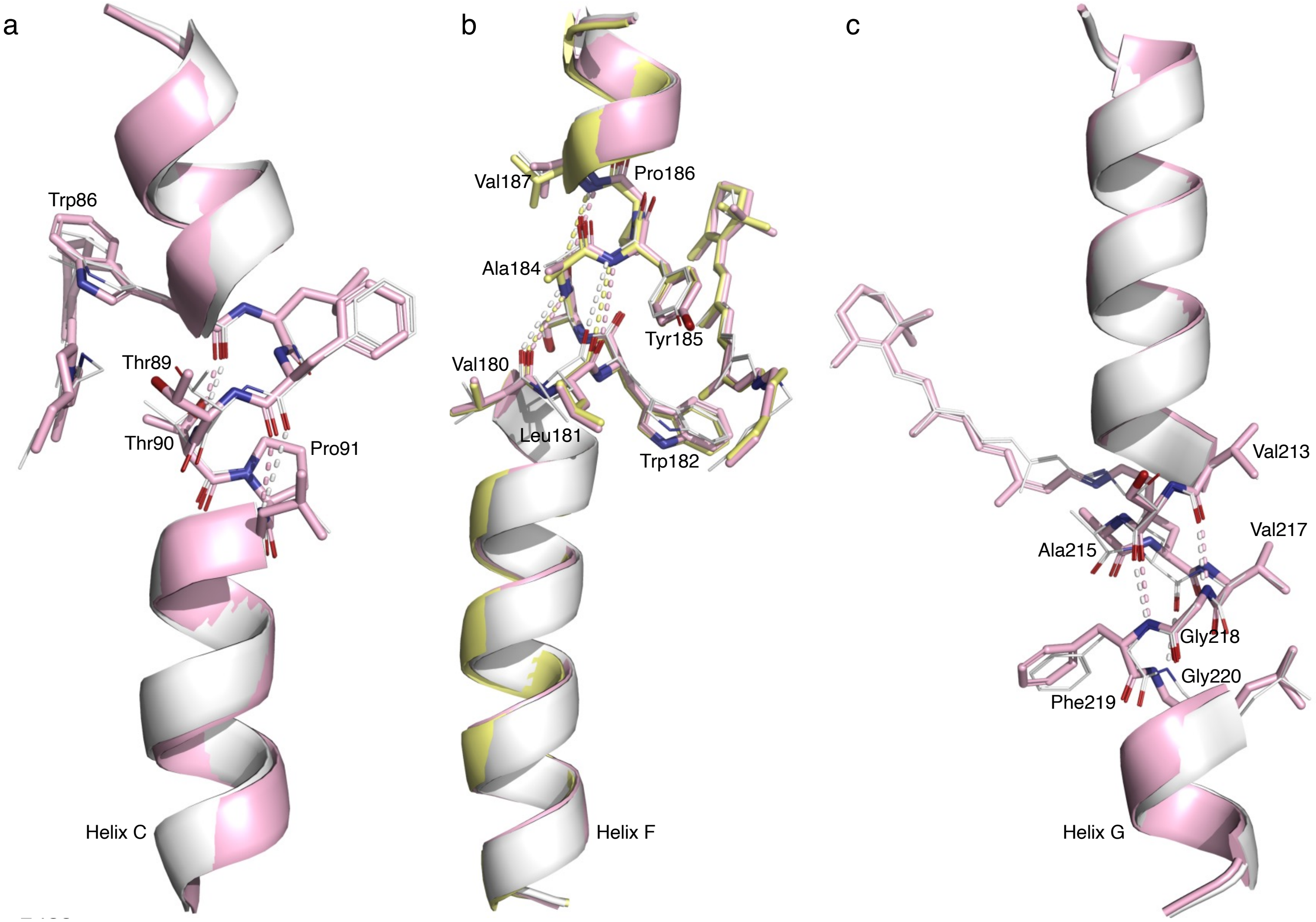


Figure S12. Irregularities in helices C, F, and G. bR in the resting state, L, and M are rendered in white, pink, and yellow, respectively. (a) Kinked helix C. All main chain H-bonds are regular except that Pro91 in helix C causes a kink. This kink was noticed as soon as the bR structure was determined to a reasonable resolution (Grigorieff et al., 1996). In the resting retinal, the corner of $C_\varepsilon$ makes a contact with Thr89 and pushes the helix inboard. In bR, J, K, and L, $C_\varepsilon$ moves more and more outboard (Fig. 4f). So does Thr89. The opposite motions occur in L, $M_1$, $M_2$, and back to bR (Fig. S16c). Thr90 also contributes to the seal. Pro91 is located exactly at right place to facilitate this spring-loaded bend. (b) Stretched helix F. Pro186 removes one H-bond in helix F. Several would-be H-bonds in the main chain around Pro186 are stretched to 3.6-4.2 Å in the resting state, such as Val180O-Ala184N, Leu181O-Tyr185N, and Ser183O-Val187N.

Pro186 and these stretched H-bonds marked by dashed lines effectively breaks helix F into two segments. In J, K, and L, Trp182 is pushed by $C_{20}$ methyl group more and more toward the CP for more than five decades. The breakage in helix F stretches even more. P186L mutant is likely to feature an intact α helix F with uninterrupted main chain H-bonds. Its proton pumping activity is severely affected (Hackett et al., 1987). A more rigid helix F would increase the difficult to displace Trp182 toward the CP direction and hinder the opening of the CP half channel for proton uptake. This stretched segment springs back in $M_1$ in yellow. (c) π helical segment in helix G. The retinal anchor is in a π helical segment with two main chain H-bonds Val213O-Gly218N and Ser214O-Phe219N. Unlike the regular α helix, a π helix features H-bonds from the carbonyl of residue *n* to the amide N of residue *n*+5. During the transition from the α helix to the π helical segment, Val217N does not have a H-bond partner. As the π helical segment transitions back to the regular α helical conformation, Ala215O does not join the H-bond pattern for helical formation. Loss of main chain H-bonds on both sides of the retinal anchor allows the anchor to move 1.2 Å in L. In addition, two nearby Gly218 and 220 increase the flexibility of the anchor.

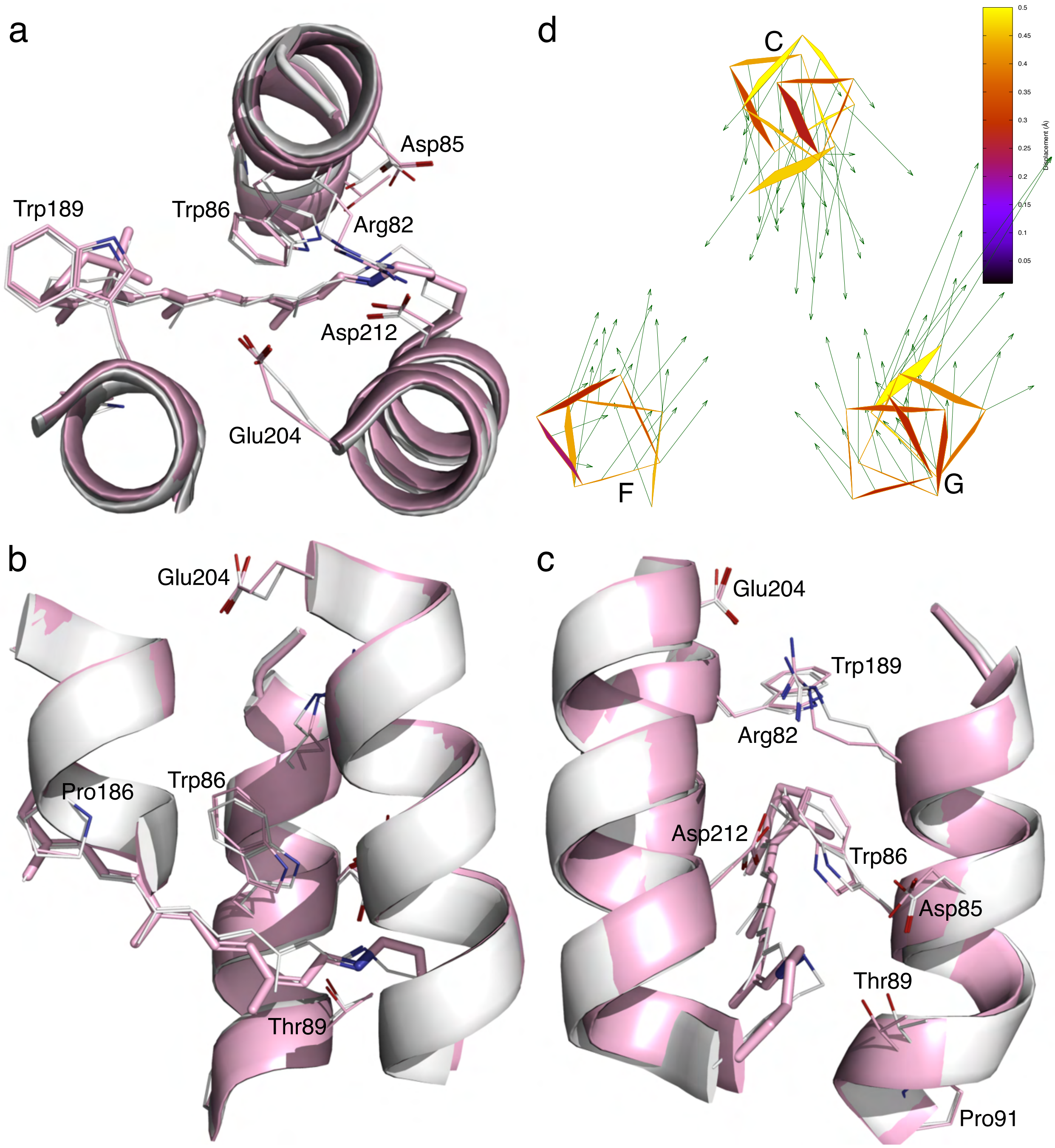


Figure S13. Tighter EC half channel in L. (a-c) Three orthographical views of the EC half channel. The resting state and L state are rendered in white and pink, respectively. (d) Atomic displacements in the main chain around the EC half channel from the resting state to L are marked with arrows 10× as long as the actual values.

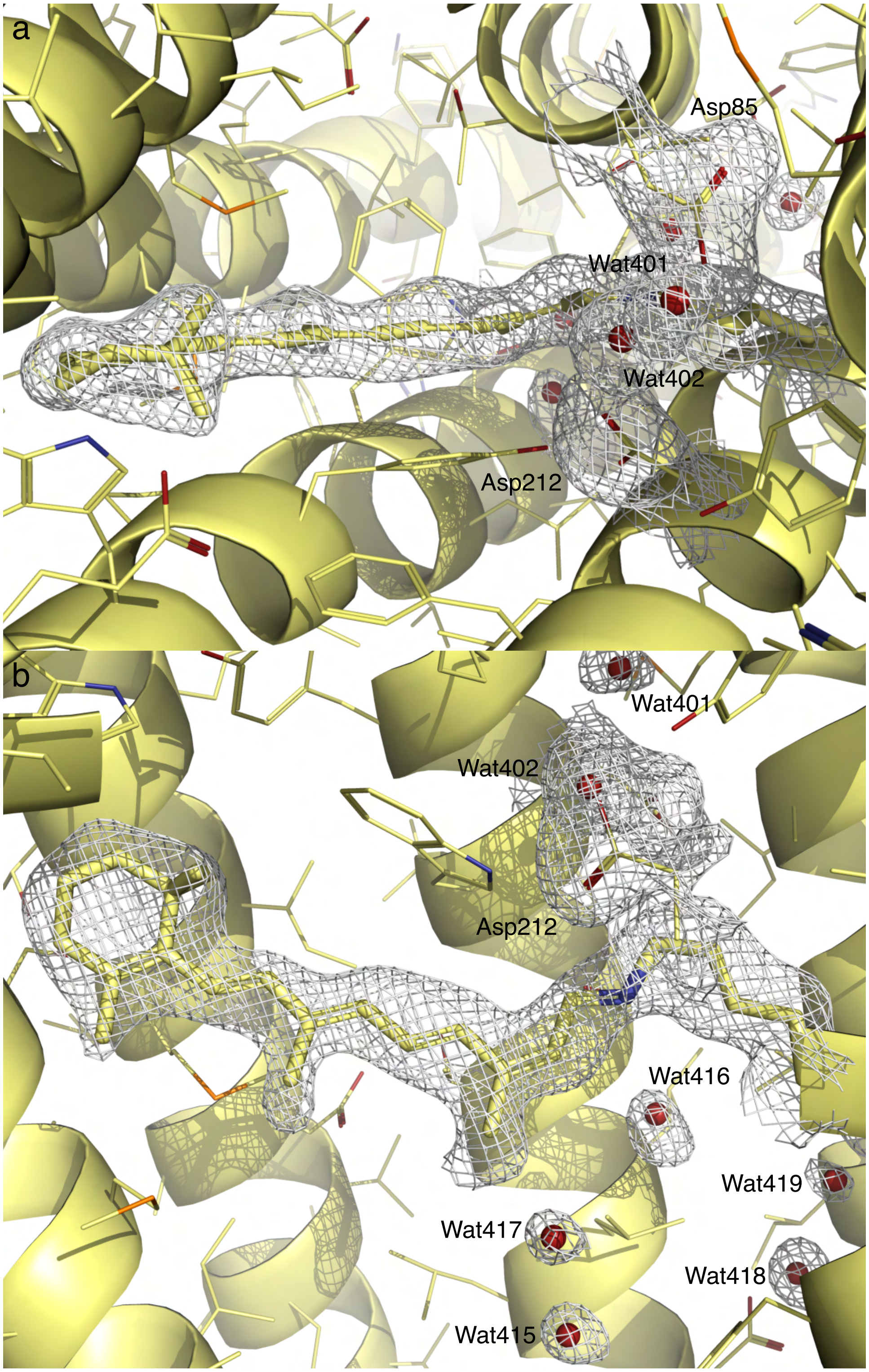


Figure S14. Two orthographical views of the 2Fo-Fc map of $M_1$ contoured at $3\sigma$. Here Fo is the reconstituted structure factor amplitudes rather than observed amplitudes

(Table S2). Fc is the structure factor amplitudes calculated from the refined structure (Methods).

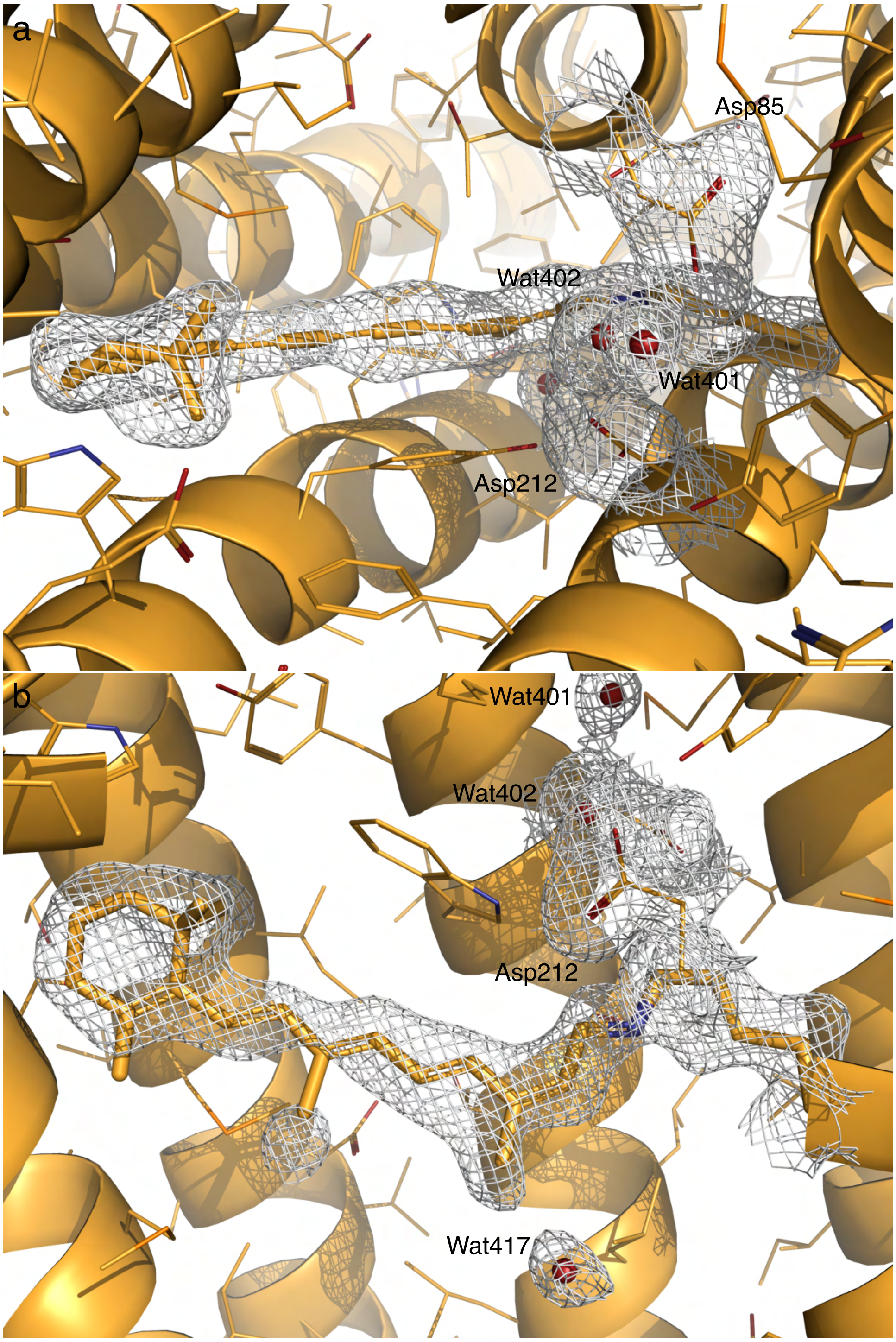


Figure S15. Two orthographical views of the 2Fo-Fc map of $M_2$ contoured at $5\sigma$. Here Fo is the reconstituted structure factor amplitudes rather than observed amplitudes (Table S2). Fc is the structure factor amplitudes calculated from the refined structure (Methods).

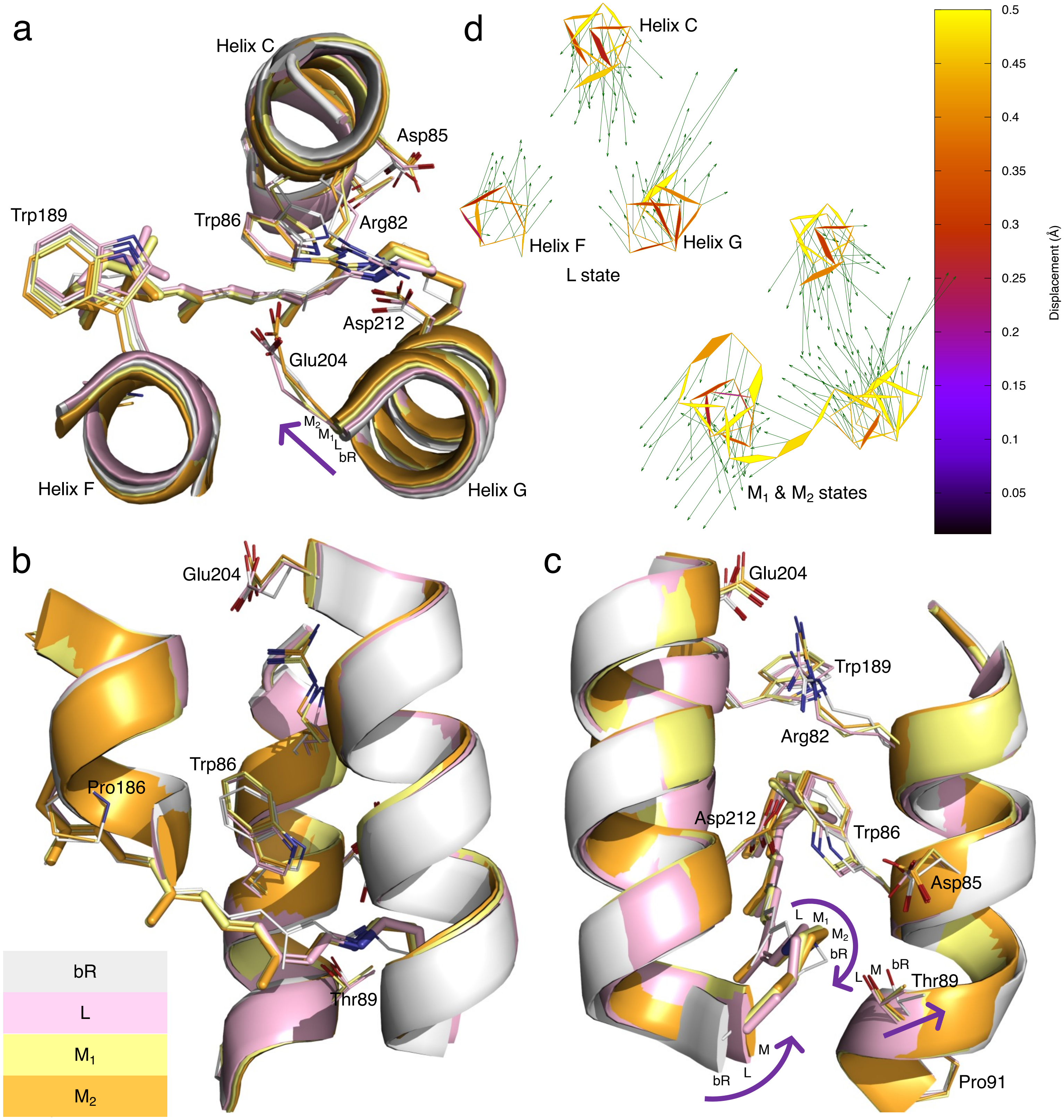
a
Helix C
Asp85
Trp189
Trp86
Arg82
Asp212
Glu204
M2
M1
L
bR
Helix F
Helix G
d
Helix C
Helix F
Helix G
L state
M1 & M2 states
0.5
0.45
0.4
0.35
0.3
0.25
0.2
0.15
0.1
0.05
Displacement (Å)
b
Glu204
Trp86
Pro186
Thr89
bR
L
M1
M2
c
Glu204
Trp189
Arg82
Asp212
Trp86
Asp85
L M1
M2
bR
L M bR
Thr89
M
bR L
Pro91

Figure S16. Intermediates L, $M_1$, and $M_2$. (a, b, and c) Three orthographical views of EC half channel. The refined structures of L, $M_1$, and $M_2$ in pink, yellow, and orange, respectively, are compared to the resting state in white. Several arrows mark the motions from the ground state bR to these intermediates or from these intermediates back to bR. The SB is moving more and more inboard since L and finally establishes the corner at $C_\varepsilon$ protruding inboard in the ground state by isomerization back to all-*trans*. Meanwhile, Thr89 and helix C are pushed away. (d) Atomic displacements in the main chain around the EC half channel from the resting state to L and M are marked with arrows 10× as long as the actual displacements. The EC segments of helices C and G move closer to each other as Arg82 swings out. The same portion is displayed as in (a-c).

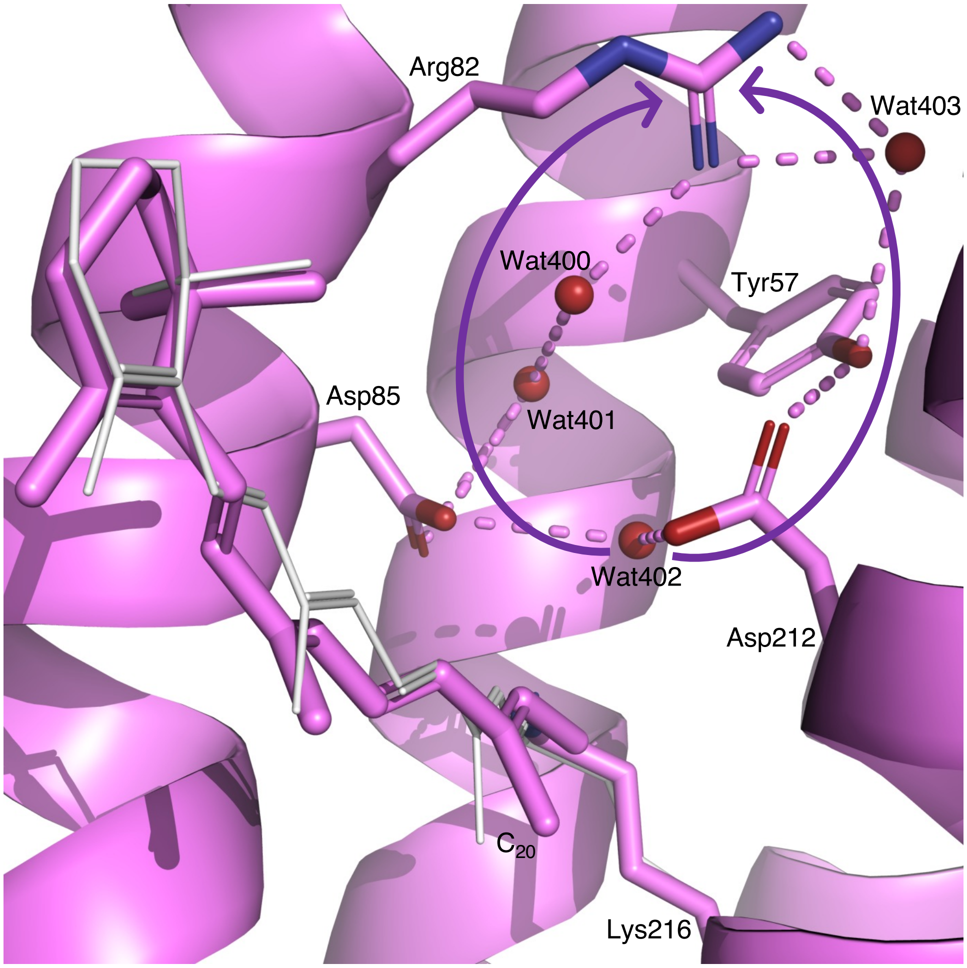


Figure S17. H-bond network at end of EC half channel in I state. Two curved arrows mark two possible proton conductance pathways. At more acidic EC pH, the pathway through Asp85 is in use and that through Asp212 is blocked by Tyr57. At more alkaline EC pH, Tyr57 could deprotonate thus the pathway through Asp212 may help to increase the proton flow. Both pathways conduct protons only under a concentration gradient of protons.

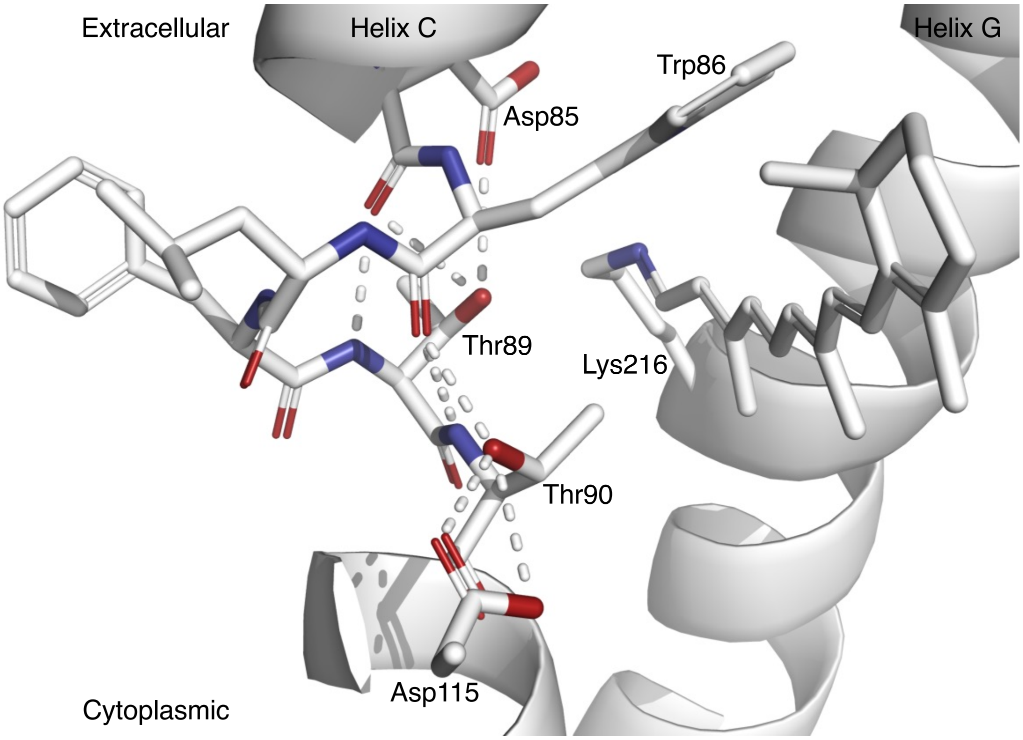


Figure S18. H-bond network involving Thr89 and Thr90. Both Thr residues are important to the function of the proton pump bR. Each $O_\gamma$ hydroxyl group is H-bonded with the main chain carbonyl of residue *n* – 4, where *n* is the residue number of the Thr. This is a common way to fix the rotamer position of the Thr side chain by forming a heptagon ring structure -$O_\gamma$-H:$O_{n-4}$:H-N-$C_\alpha$-$C_\beta$-, where all atoms belong to the Thr residue *n* except $O_{n-4}$ from the main chain carbonyl group of residue *n* – 4. Thr90 also forms H-bonds with Aps115. Its side chain conformation is important to the suppression of the byproducts during the photoinduced isomerization sampling (Ren, 2022). Thr89 is also H-bonded to Asp85. Therefore, the side chain of Thr89 is fixed toward the SB at a distance of 3.4 Å in the ground state. During the photocycle, Thr89 is constantly pressing on the SB in motion due to the irregular, bent helix C that has the tendency to be straightened (Fig. S12a).

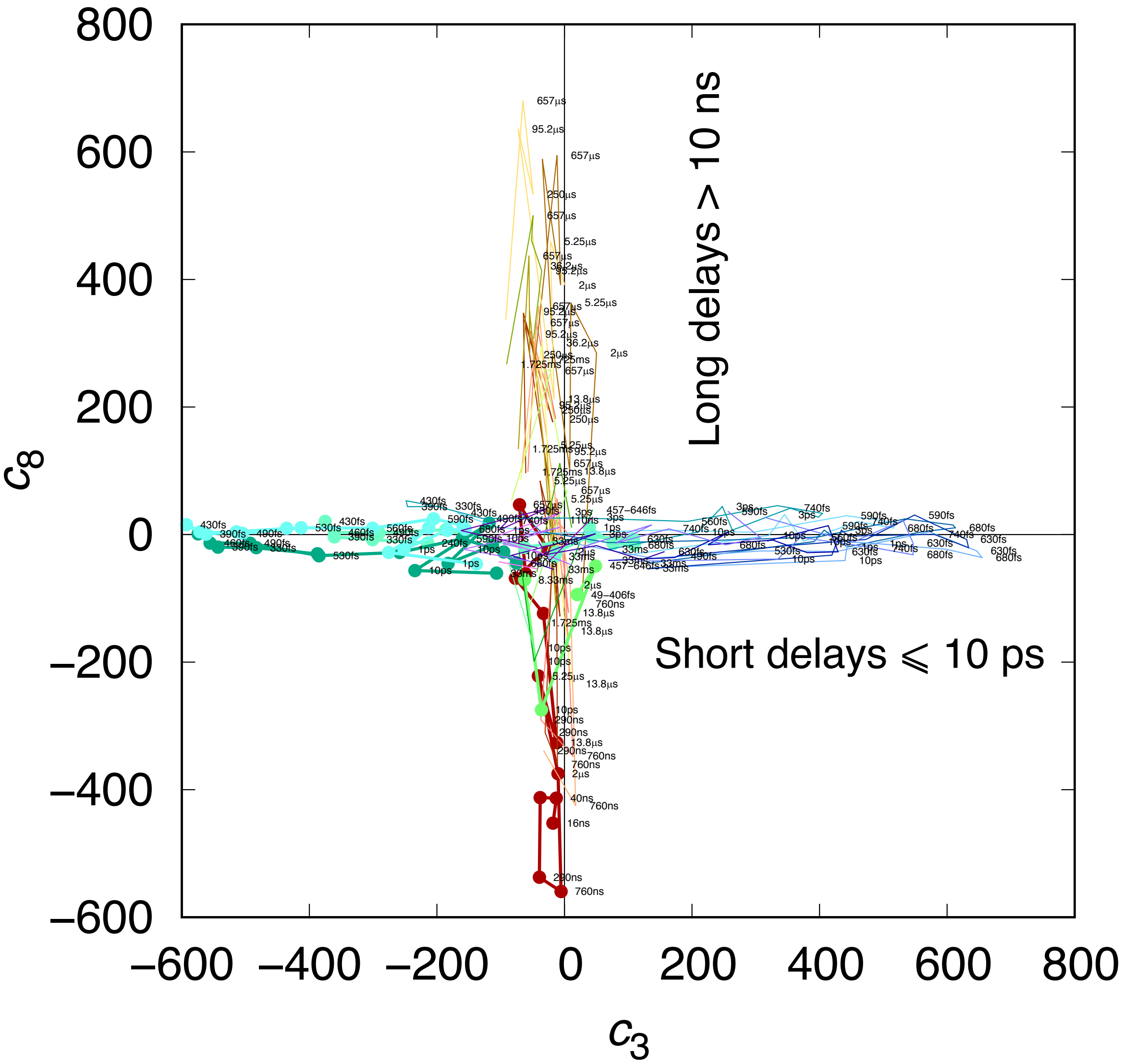


Figure S19. Short and long delays in two orthogonal subspaces. A preliminary SVD analysis shows that the difference Fourier maps at short and long delays do not share common features among the major components. This is an example to show that the third and eighth components are in two orthogonal subspaces. Each colored trace represents difference maps in a time series calculated with a common reference. Those time series with a dark reference are plotted in thick lines with dots. Other series are in thin lines without dots. Short delays are in cool colors and long delays are in warm colors.

*From:*
*Subject: PNAS Nexus MS# PNASNEXUS-2022-00209 Decision Notification*
*Date: April 5, 2022 at 1:48:55 PM CDT*
*To:*
*Reply-To:*

*April 5, 2022*

*Title: "Directional proton conductance in bacteriorhodopsin is driven by concentration gradient, not affinity gradient"*
*Tracking #: PNASNEXUS-2022-00209*
*Authors: Ren*

*Dear Professor Ren,*

*After an initial evaluation of its general suitability for PNAS Nexus, your manuscript was assigned to an editor with expertise in the subject area of your work, who then examined the manuscript and obtained reviews from 2 authorities in the field. These reviews, which appear below, were then considered by the editor, who determined the comments were of sufficient significance to recommend rejection.*

*Because we receive many more manuscripts for consideration than we can accept for publication, critical comments from expert reviewers that cannot be easily and rapidly addressed in a revision are sufficient cause for rejection; our goal to publish important research rapidly prevents us from undertaking multiple rounds of revision and re-review.*

*Although we expect that this outcome is disappointing, we hope that the rationale for our decision process is clear and that you will find the comments of the reviewers helpful when revising your work for submission elsewhere. We thank you for choosing PNAS Nexus as a potential venue for publishing your work and we hope that you'll continue to consider PNAS Nexus for future submissions.*

*Yours,*
*Karen E. Nelson*
*PNAS Nexus Editor-in-Chief*

********************
*Three expert scientists in the field agreed to review this work, but only two submitted their evaluations. Nevertheless, both reviewers found serious fault with the model presented by the*

*authors. Both reviewers indicated that the authors' conclusions were not sufficiently justified and that the overall quality was insufficient for publication in PNAS Nexus. Although one of the reviewers stated that the authors elegantly succeeded in analyzing more than 40 structures, the reviewer added that the suggested model was quite different from the standard model and that further experimental evidence or computational support would be necessary to reconcile the proposed model. The second reviewer states that the authors make extensive claims that contradict what is solidly known. Both reviewers provide extensive comments and concerns. Consequently, it is my opinion that the current submission would be inappropriate for publication.*

*Reviewer #1 Comments for the Author:*

*Bacteriorhodopsin (BR) is the best characterized light-driven proton pump that can convert photon energy to the proton motive force to energize the cells with compact seven-transmembrane architecture and an all-trans-retinal chromophore. The mechanism of H+ pumping by BR was studied in detail by a pile of biochemical, spectroscopic, theoretical, and structural biological methods for five decades leading a sophisticated model in which H+ is sequentially transferred between retinal Schiff base (RSB) and acidic residues (Asp/Glu) along with the appearance of spectrally distinguishable multiple photo intermediates called K, L, M, and so on. Especially, the time-resolved serial X-ray crystallography using X-ray free-electron laser or synchrotron radiation enabled real-time observation of 3-D structural change of the entire body of the protein at room temperature and provided unprecedented detailed structural information. The structures observed at specific time points after the light illumination, however, are mixtures of several intermediate species so that it is difficult to correspond the observed conformational changes to the respective intermediates.*

*In this work, Dr. Ren elegantly succeeded in extracting the pure structures of each photo intermediate by applying SVD analysis for more than 40 structures previously reported in different serial X-ray crystallography studies. This would make a new basis for the methodology of the analysis of serial X-ray crystallographic data and be useful for understanding the dynamic events of a wide range of proteins. On the other hand, the suggested model involving ultrafast deprotonation of RSB in femtosecond time region and concentration driven H+ release is quite different from the standard model (e.g. shown in Ernst et al. Chem. Rev. (2014) in the reference list in the manuscript) in which deprotonation of RSB occurs in ~50 μs on the formation of the M intermediate from the L, and dynamic pKa change of RSB and acidic residues is the key element to drive the vectorial H+ transfer.*

*One of the biggest concerns about this work is the precise position of H+ was not observed in the X-ray crystallographic data so that the formation of hydronium ion was not directly observed in the results. This was suggested by only the theory based on Newton's law. Although the author demonstrated that H+ is left out from the RSB due to its ultrafast movement by the same physics as tablecloth trick, the H+ does not weakly interact with RSB like a dining set on a tablecloth but is covalently bound with RSB. Therefore, it can quickly move with RSB as a single object. In order to make the author's model sure, further experimental evidence or quantum chemical calculation is needed.*

*Although the author explained the reason for the discrepancy of his model and the previous interpretation of RR study in the 2nd paragraph on page 10 (line 314-330), it is not sufficiently convincing unless further experimental or computational support is shown. Especially, although the position of C=N vibration in the RR spectra correlates with the absorption wavelength of each photo intermediate as the author mentioned, the downshift with deuterium can be explicitly observed so that the assignment of the protonation state of RSB by RR spectroscopy is extremely solid and consistent with FTIR spectroscopy which also observed N-H vibration of RSB not only in the dark state but also in the K and other intermediates. Further experimental or computational results are needed to reconcile the author's model with these vibrational spectroscopy studies.*

*The author demonstrated that a H+ is pooled in the extracellular part of RSB during the photocycle initiated by the first photon, and further supplements of H+ during subsequent photocycle(s) are needed for the efficient H+ release to the extracellular solvent. However, if it is the case, because the absorption of the retinal chromophore is highly sensitive to the electric charge around RSB (e.g. Melaccio et al., J. Chem. Theory. Comput. (2016)), a new state exhibiting an absorption significantly shifted, probably toward the red side, from the initial state must be observed at the end of the first photocycle by the transient absorption spectroscopy. But, to my knowledge, no supportive result was reported so far. Also, the H+ release to the solvent was observed in a single-photon photocycle using pH indicator (pyranine) (Varo et al. Biochemistry (1993)). This also cannot support the presence of the super-acidic inner EC channel.*

*Page 17, Line 549-550*
*"The observed kinetics of FTIR continua could explain excess protons harbored by the water cluster in the inner EC channel"*
*The FTIR spectroscopy observed continuum band for the "initial" state (assigned to the protonated water cluster between Glu194 and Glu 204) not for any photo intermediate. Hence, it cannot support the hypothesis of the author that hydronium ion is generated in the EC cavity during the first photocycle.*

*These concerns need to be clarified before considering the model of this work publishable. Otherwise, I strongly recommend to the story be significantly changed to focus only on the structural changes other than the position of H+, which can be firmly observed by X-ray structural analysis, without mentioning H+ transfer different from the standard model. Since the author's SVD analysis is highly effective to extract the pure structural change occurring in each photo intermediate and would be useful also for the serial X-ray crystallography in the future, it would be better to avoid indicating the fact which cannot be explicitly supported by the results.*

*-Minor points*
*Page 11, Line 340*
*What is the meaning of "reliable"? Why does it need to be emphasized so?*

*Page 12, Line 389*
*The J-to-K conversion occurs at 3-5 ps (e.g. Nuss et al. Chem. Phys. Lett. (1985)). Therefore, The J' and J in this article should be assigned to the J and K, respectively.*

*Page 14, Line 455-456*
*"the transition times between intermediates K, L, M1, and M2 known from spectroscopy"*
*Please add the relevant references.*

*Page 15, Line 498*
*"83" would be better to be "Tyr83".*

*Page 18, Line 592-593*
*"Even if these changes in proton affinity are achievable, there will be 10% failed attempts of proton transfers at a prolonged equilibrium"*
*How was the degree of failure (10%) was estimated?*

*Figure 1*
*The N and O intermediates have 13-cis and all-trans retinal, respectively.*

*Reviewer #2 Comments for the Author:*

*The manuscript revises crystallographic data from recent serial femtosecond crystallography experiments on bacteriorhodopsin. On this basis the results are re-interpreted and a different conclusion is reached. The conclusions are questioning the current understanding of the photocycle in bacteriorhodopsin. This work is stimulating and interesting. However, I see one major inconsistency. If the bacteriorhodopsin photocycle would indeed start with the deprotonation of the Schiff base in retinal, then the spectral properties would drastically change. A deprotonated Schiff base absorbs at 380 nm, instead of 568 nm from protonated Schiff base in bacteriorhodopsin. But ultrafast spectroscopic studies do not show such a spectral shift. Therefore there is no evidence for what is proposed by the author. In addition, a change in the protonation would also lead to a change which can be detected by vibrational spectroscopy, which is also not described in the literature as far as I know. Therefore it is important that the author will describe how the new mechanism can be verified using time resolved spectroscopy or how it fits with a large body of published spectroscopic results.*

*Further points:*
*1) The author has mentioned several times that previous studies were mistaken. But it was not clear to me what the mistake was.*
*2) The inertia of the Schiff base proton is mentioned several times. But the mass of the proton is much lighter than of a nitrogen atom. So I would expect that the Schiff base nitrogen has more inertia.*
*3) Did the author compare the energy of the hydrogen bond with that of the photon that excites bR? The energy of the photon should be much larger than the hydrogen bond. It answers the question about the driving force of the proton pump, but it is not discussed in the manuscript.*

*Reviewer #3 Comments for the Author:*

*Ren provides a manuscript that feeds into published time-resolved X-ray crystallographic data on the bacteriorhodopsin photocycle to determine more accurate structural models for the intermediate states. Then, he uses the obtained structural model to provide a very personal view on how bacteriorhodopsin proton pumps.*
*To give a background, the general "believe" is that photoisomerization of the all-trans retinal with a protonated SB remains in a protonated 13-cis conformation until the L intermediate. In the L to M transition (tens of microseconds) the SB deprotonated in favor of Asp85, and soon a proton is released from the proton release complex (PRC) in the EC domain. The SB reprotonates from Asp96, a residue in the CP domain. Later, Asp96 recovers it proton from the CP medium, completing the proton pumping. The initial state is recovered in few tens of milliseconds as the protonated 13 cis-retinal reisomerizes to all-trans and Asp85 transfers its proton to the PRC.*
*Ren introduces several ground-breaking ideas: 1) very fast (femtoseconds) deprotonation of the retinal SB, instead of taking place in tens of microseconds. 2) The metastable proton acceptor of the SB is not Asp85 but a water molecule. 3) The photocycle does not pump protons in its first round, but forms a super-acid initial state. When this super-acidic initial state absorbs light, then the photocycle pumps protons.*
*While I am the first one that think some speculative science help to bring new ideas to the discussion and helps the progress, the problem is that Ren makes innumerable claims that*

*contradict what it is solidly known. As I will made clear below by commenting some of the claims in the paper that I think are wrong, the present MS dismisses, omits, or ignores many solid experimental solid results that contradicts the core of his proposed proton-pumping mode. These critical comments are mixed with other comments and questions I had while reading the MS. All of them are provided below.*

*-Line 53. "This integral membrane protein of 28 kDa singlehandedly achieves the exact same goal of photosynthesis and cellular respiration combined."*
*Photosynthesis uses light to create a proton electrochemical gradient, but also to transfer electrons from an initial donor (H2O or other inorganic molecule) to a final acceptor (NADP+) against their redox potential, and cellular respiration uses electron transfer from an organic donor (NADH or other) to an inorganic acceptor (O2 or other) in favor of their redox potential to build a proton electrochemical gradient. These two processes combined do not achieve the same goal as light-driven proton pump by BR. Please, rewrite.*

*-Line 56: "A trimeric form of bR on the native purple membrane shares the retinal chromophore and the same protein fold of seven transmembrane helices A-G (Fig. S1) with large families of microbial and animal rhodopsins (Ernst et al., 2014; Kandori, 59 2015)"*
*This statement might suggest that animal rhodopins and other microbial rhodopsins also adopt a trimeric form, which is not the general case. Please, rewrite.*

*-Line 68 "It has been shown that the species I or a collection of species prior to the photoisomerization arises before 30 fs and remains in 13-trans instead of a near 90° configuration about C13=C14 (Zhong et al., 1996)."*
*Why a collection of species? In the cited work the authors determined a single rising component at 18[illegible]10 fs.*

*-Line 80. "However, the event of SB deprotonation has been mistakenly inferred from the observed protonation of the carboxylate of Asp85."*
*SB deprotontation has been mainly studied by the rise of the retinal absorption at 400 nm in UV-vis spectroscopy, while Asp85 protonation has been studied by the rise of the C=O of its carboxylic group at 1760 cm-1. By far, the most habitual technique used has been visible spectroscopy and, thus, SB deprotonation has been rarely deduced by Asp85 protonation, but the other way around: people doing visible spectroscopy often assume Asp85 protonation when they observe SB deprotonation.*

*-Line 82. "It has been widely quoted in the literature that the proton transfer from the SB to the proton accepter occurs as the M formation."*
*Independent studies by both Gewert and coworkers (10.1016/S0006-3495(92)81722-8; 10.1016/S0006-3495(93)81264-5) and Lorenz-Fonfria and co-workers (10.1021/jp201739w )*

*have shown that both the rise of the absorbance at 400 nm and at 1760 cm-1 follow roughly the same kinetics. Mutagenesis studies also point to Asp85 as the proton acceptor. So, there are good reasons to think that a proton is transferred from the SB to Asp85 in the M intermediate.*

*-Line 169: "Therefore, the SB proton is associated with both N and O simultaneously in Nz:H+:OH2 with both interactions much weaker than an ordinary single bond (Fig. 2c)."*
*Add "atoms" after N and O to avoid confusion with the N and O intermediates.*

*-Line 174: "The difficulty encountered by this hypothesis is the negative pKa values of a hydronium ion and a protonated threonine that the proton has to go through."*
*I agree that it intuitively it does not seem plausible, but science should not be based on just intuitions. Bondar and co-workers have several papers showing that such proton transfer from a CP SB to Asp85 has an energy barrier low enough to be possible (check her review: 10.1007/s00232-010-9329-3 ).*

*-Line 181: "This interpretation of these highly twisted double bonds is a direct consequence of the inability to unscramble a mixed observation of multiple conformations."*
*Without further explanation this sounds as an opinion dressed as a fact. Being an opinion, it should better read as: "This interpretation of these highly twisted double bonds could be a direct consequence of the inability to unscramble a mixed observation of multiple conformations."*

*.Line 186: "Q1) How does the protonated SB with a pKa of 13.3 transfer its proton to the acidic acceptors with a pKa of 2.2 and when?"*
*I think that the "when" has been known for long time: in the L to M transition. The author can check numerous reviews on BR that have been published by different authors.*

*-Line 188: "Q2) Which molecular event, the isomerization, or another event, switches the accessibility of the SB from EC to CP and how does the SB avoid reprotonation on the wrong side?"*
*Several works from Varo & Lanyi (for instance 10.1021/bi00234a024 ), and other studies, point to the irreversible M1 to M2 transition as the switching event. Results from Balashov (reviewed in 10.1016/s0005-2728(00)00131-6 ) show that the pKa of Asp85 is coupled to the protonation state of the proton release complex. Lorenz-Fonfria & Kandori (10.1021/ja900334c ) showed that the proton release complex deprotonates in the M1-to-M2 transition. All these observations indicate that deprotonation of the PRC increases the pKa of Asp85 and, in combination with structural chanages, forces the SB to recover its proton from a CP group.*

*Line 190: "Q3) How are protons conducted > 15 Å through the rollercoasting pKa values and released into the EC medium from the SB of a high pKa through aspartic acids of low pKa,*

*arginine (strictly speaking, its guanidinium ion) and tyrosines of high pKa, and glutamic acids of low pKa again?*
*The released proton does not originate from the SB, but from the proton release complex (PRC). This is well-established, as the author can confirm reading any review. What it is less clear is how the PRC recovers its proton from Asp85 in the O to bR transition, but simulation studies from the group of Elstner and co-workers have addressed this question (10.1021/ja809767v ).*

*-Line 194: "No convincing evidence can directly demonstrate a photoinduced reversal of the drastically different proton holding abilities along the EC half channel to facilitate a spontaneous proton conductance that would require an ascending order of pKa's."*
*I do not understand what the author means here.*

*-Line 197: "The fundamental flaw in the previous line of thinking is multiple steps of equilibrium driven by proton affinities as the values of pKa describe."*
*Not clear why it is a flawed assumption. Proton transfers are rather fast events, so for transitions between intermediates in the microsecond (K-to L, L-to-M, M-to-N, N-to-O and O-to-bR) it makes perfect sense to assume an equilibration of the protonation state governed by the pKa.*

*-Lines 211-218: "Here the ultrafast molecular events revealed from the serial crystallographic datasets demonstrate that the photochemical reaction of bR transfers a proton held by the SB to form a strong acid that barely has any ability to hold a proton. However, this proton transfer is not achieved by a chemical equilibrium therefore not governed by pKa's. Instead, before the isomerization occurs, this proton transfer takes place due to the SB displacement within ~30 fs in a single molecular event. The proton's inertia prevents it from accelerating as fast as the SB under the same physics of the tablecloth 218 trick (Q1)."*
*These are bold claims about what happens to the SB proton by just looking at electron densities from X-ray crystallography, virtually blind to hydrogen atoms.*

*-Line 240. "Eight intermediate structures along the photocycle, now isolated from one another, are refined against reconstituted structure 242 factor amplitudes (Methods; Table S2)."*
*For the long delays data, 6 significant SVD components are identified. However, Table S2 shows that intermediates from K to M2 are reconstructed with only four components: 1, 2, 3 and 6. What happens with components 4 and 5? In addition, looking at Fig. S4a it is not clear why only 6 components are selected. Components 6 and 7 (and the following) have almost the same contribution in describing the data set.*
*It is also surprising that for the early delay, which should include the K intermediate and thus 5 intermediates, only 3 significant components are found (rank of 3). This would rather suggest that there are only three structural intermediates in this time window: I, J and K. In other words, which is the evidence or reason to include I'and J' as intermediates? They do not seem to be*

*needed to describe the data when considering that SVD indicates a rank of 3. If the five intermediates co-exist in this time window, should not the rank found by SVD be equal to 5? There are more issues about the application of SVD to the early delay data. SVD assumes that the data can be decomposed in three matrices, as explained by the author. For the present application this assumption translates into assuming that the experimental data can be described as the linear combination of electron densities of a discrete number of intermediates. While this is true at longer time scales, where the energy barriers force the system to populate only a discrete number of intermediate states, in the very fast regime the structural changes are ballistic and show coherence effects: the system does not necessarily experience a transition between intermediates but show a continuous change, and the notion of discrete intermediate might lose sense and, thus, the applicability of SVD. This regime can be identified by the presence of coherent oscillations in the positions of atoms. I guess these oscillations are present in the data, right?*

*-Lines 276-297. The authors provide some arguments that "proof" that the SB deprotonates few fs after light absorption, but these arguments are very speculative. The N-H vibrational frequency of the SB in the ground state of BR has been determined by the group of Kandori in D2O (corresponding to the N-D vibration, 10.1021/bi025585j ). Considering a typical isotopic effect, it is possible to deduce the N-H frequency for the SB. This frequency is related to the force of the bond holding the atoms N and H together. My rough calculation from the N-H frequency and the reduced mass gives me a force of 600 N/m for the bond, meaning that if the N atom moves 1 pm but the H atom does not move, the H atom will feel a force of 600 pN, enough to get the acceleration it seems to need to keep moving with the N atom.*
*Is it still possible that the SB could deprotonate by a different mechanism or reason after photoexcitation, but the group of Oluvicci have made dynamics simulations of the retinal isomerization in BR and other microbial rhodopsins at the QM level and they have not reported a SB deprotonation following light excitation to my best knowledge. Also, if the SB was to deprotonate after photoexcitation, the electron density of the retinal would rapidly adjust to the new deprotonated state and the I intermediate would be largely blue shifted, as the M intermediate is.*

*-Lines 299-312: The author quotes references to FTIR studies describing broad bands assigned to protonated water clusters in support this idea that the SB proton is early accepted by a water molecule. However, the author does not mention that the reported continua bands in BR are in all cases negative, i.e., protonated water clusters present in the initial BR state transiently deprotonate during the photocycle. Also, continua band in BR have been only reported to occur in the L to M transition and N-to-O transition, i.e., in the microseconds-milliseconds.*

*-Lines 314-330. The author suggests that the believe that the SB remains protonated until the L to M transition is only (mostly?) based on Resonance Raman results. RR has used the*

*sensitivity of the C=N frequency to deuteration as a proxy for both the H-bonding and protonation state of the SB. The problem is that both a H-bonding free and a deprotonated SB are H/D insensitive, meaning that it cannot distinguish a protonated SB losing its H-bond and a deprotonated SB. But the author seems to ignore that the N-H frequency of the SB has been study by FTIR spectroscopy by the group of Kandori. This vibration can discriminate between changes in H-bonding (the band shifts) and deprotonation (the band disappears). In the K intermediate the authors observed that the N-H frequency upshifts, meaning that in the K intermediate the SB is protonated and is less H-bonded than in the initial state (10.1021/bi025585j ).*
*In addition, the intensity of retinal C-C vibrations in FTIR experiments it is known to be very sensitive to the protonation state of the SB. By this means FTIR experiments can follow changes in the protonation state of the SB, which seems to deprotonate in the L-to-M transition, and not earlier (10.1021/jp201739w ). Lastly, as already mentioned, the retinal absorption in the visible is notably sensitive to the protonation state of the SB, with results totally consistent that the M intermediate is the only intermediate state (at least the only one significantly populated) with a deprotonated SB.*

*-Line 341: "The deprotonation event [from the SB] cannot be directly inferred from the vibrational signatures due to the protonation of the carboxylic proton acceptors in the Asp residues."*
*Why not? As mentioned above there are several spectroscopic signatures (N-H vibration, C-C intensities, lamda max of the retinal in the UV-vis) that inform about the protonation state of the retinal SB.*

*-Line 343: "the "proton transfer" from the SB to the proton acceptor, as commonly stated in the literature, is not a single event. Instead, it spans ten decades in time with multiple asynchronous molecular events."*
*I disagree. The proton transfer from the SB to Asp85 takes place in the L to M transition. It is true that this transition does not follow a single exponential and, thus, is more complex and expands certain time. But the deprotonation of the SB does not start until the microseconds accordingly to what it is known.*

*-Line 359. "This ultrafast U-turn is how the SB gets rid of the proton by a mechanism of the same physics that governs the tablecloth trick (Q1; Fig. 361 S2).*
*First, as I mentioned, protons are not "visible" in X-ray crystallography (unless the resolution is exceptional, which is not the case). Second, as also mentioned, there is no reason to think that the hydrogen bound to the SB N atom will not be able to follow it as it moves during retinal photoisomerization: the N-H bond can provide enough force if moved from its equilibrium distance by few pm.*

*-Line 367: "The U-turn of the SB marks the peak power, or the maximum rate of enthalpy change dH/dt, of the entire photochemical reaction (see Concluding 369 Remarks)."*
*This statement is far from evident. I do not see why.*

*Line 371: "If a deprotonation is unsuccessful at the peak power of the photocycle, there will be no further opportunity to translocate a proton from a fair base to a decent acid uphill against 11 pH units under equilibrium."*
*This statement is not supported by any evidence, besides of the assumption that the photocycle relays on this early deprotonation to pump protons, which I do not think to be the case.*

*-Line 383: "Incidentally, all previous hypotheses on the mechanism of SB deprotonation imply a formation of hydronium at one point or another (Lanyi and Schobert, 2007; Nango et al., 385 2016). It would require a donor group with a pKa more negative than -1.7 to reliably form a hydronium by the means of equilibrium."*
*Basically, all proton transfer reactions in solution are mediated by the transient protonation of water. Simulations at the QM level in BR also show that proton transfer reaction between groups in BR are possible and mediated by the transient protonation of water molecules, as shown by different authors (see works by Bondar and co-workers, by Gerwert and co-workers, or by Elstner and co-workers). The key is that this "hydronoium ions" hold the proton for a very small fraction of time, while the author seems to suggests that between the deprotonation of the SB in few fs until the potonation of Asp85 in tenths of microseconds, W401 stays protonated.*

*Line 450: "Four of them contain outstanding signals of structural changes (Fig. S5). Two others also carry structural signals but without a clear time dependency (Fig. S6)."*
*The time dependency is mentioned but not shown in any case. I would be nice to see the time dependence of all the 6 components, the 4 used to reconstruct the pure intermediates and the 2 with structural detail but discarded.*

*-Line 501 to 506: "Yet, the question remains: How are protons conducted through the proton barrier of the guanidinium group of Arg82 that is already protonated and the phenol hydroxyl groups of the Tyr residues that cannot be protonated anymore (Brown et al., 1993)? No evidence shows that such rollercoasting pKa values would change in any meaningful way by the light-induced conformations. Therefore, the previous proposal on an affinity-driven proton conductance is fundamentally flawed."*
*The protons are not conducted through this proton barrier, because the analyzed data reaches only until the M2 intermediate. As the author probably knows, the proton released to the EC medium does not come from the SB or Asp85, but from a different group known as the proton release complex (PRC). For the reprotonation of the PRC from Asp85, which take place much later in the O-to-BR transition, the proton does need to go through Arg82 and so on. But at this point some structural changes have take place that make it possible. The author can check*

*publications from the group of Eltsner addressing this late proton transfer (for instance 10.1021/ja809767v ).*

*-Line 508: "Proton conductance after SB deprotonation through the EC half channel not necessarily operates on the basis of one event of proton release per photocycle."*
*I am quite surprised with this statement. It is true that the proton-pump stoichiometry was for a time in the past an open debate, but work on the late eighties (mainly from Dencher but also by other) settled down the current view of one proton pumped per deprotonated SB (strictly, one proton released to the medium for each BR reaching an M intermediate).*

*-Line 510: "It was previously suggested that a proton pumped in one photocycle could be released in the next or subsequent photocycles (Braiman et al., 1991)."*
*Initial FTIR results from Asp96 carboxylic C=O bands lead to Braiman and co-workers to consider the possibility of a partial or total deprotonation of Asp96 in the L intermediate. This was latter shown to be an artefact caused by overlapping contributions from Asp96 and Asp115 in the carboxylic region (Gerwert and co-workers, and Maeda and co-workers).*

*-Line 511: "However, a flawed common assumption persists that the pumping and releasing of each proton are somehow scheduled in different periods of a photocycle."*
*This is not an assumption, but an empirical fact. The use of pH sensitive shows that a proton is released first in the EC domain and a proton is taken later in the CP domain, with the uptake completing the proton pumping (see for instance a review by Heberle: 10.1016/s0005-2728(00)00064-5 ).*

*-Line 516: "The first a few photocycles build up a proton concentration at the end of the EC half channel without any proton release."*
*How the author knows about this from the analysis of the time-resolved X-ray data? I can say that I have been single shot experiments on BR where proton release could be detected by pH sensitive dyes. Moreover, the signal from the pH sensitive dye remains the same with every photoexcitation: every photocycle releases the same number of protons. This is the reason the data is averaged, so published data is the average from multiple measurements. But, as I say, proton release takes also place in single short experiments.*

*-Line 521: "That is to say, two states sharing the same or very similar all-trans retinal are differentiated – the resting state in dark (Fig. 5b) and the beginning state of the photocycle in light with a super charged inner EC channel (Fig. 5c). The structure of the all-trans light state is not observed directly. Its existence can only be implied from the spectroscopic data."*
*The spectroscopic data refutes the above claims. If you excite BR with a laser pulse in tens of microseconds the protein returns to the same initial state. How do I know this hypothetical super charged initial state does not exist? If the final state after a single laser pulse was a super charged*

*state as the one shown in Fig.5 it would be easily detected in an FTIR difference spectrum. However, after a single laser excitation the light-induced FTIR difference spectrum goes back to zero, which indicates that the initial and final state are the same.*

*-Line 527: "Although the first a few photocycles are productive, they are not capable of releasing any proton before an established proton concentration gradient"*
*Wrong. Proton release in single short experiments can be measured, as already mentioned. The first photocycle is already able to release a proton.*

*-Line 530: "Proton release from the wildtype bR was observed before proton uptake as if it is coupled with, if not faster than, the protonation of the proton acceptor Asp85 (Braiman et al., 1988, 1991; Gerwert et al., 1990; Zimanyi et al., 1992), which strongly suggests that the newly pumped proton is not the one released to the EC medium in the same photocycle."*
*I am not sure if I understood that the author wants to say here. It is well known that in BR a proton is released first to the EC side and later a proton is uptaken from the CP side, completing by his way proton pump. The fact that proton release precedes proton uptake is not a problem, and it is perfectly explained by the order of the proton transfers inside BR.*
*It also seems that the author states here that before Asp85 protonates, a proton is already released by BR. A recent comparison of protonation of Asp85 and the release of protons show that the former takes place slightly before the second, not after (10.1073/pnas.1707993114 ).*

*-Line 546: "It appears that Asp85 and 212 are continuously protonated by proton pumping thus neutralized in the inner EC channel."*
*Protonation of Asp212 was only experimentally observed in the photocycle of wt BR at very low pH values and in some mutants (10.1021/bi990873+ and 10.1006/bbrc.2001.4730 ), in both cases in the the last steps of the photocycle (O to BR transition). Asp212 is, therefore, not continuously protonated.*

*-Line 548: "Several hydronium ions make the inner EC channel positively charged. The observed kinetics of FTIR continua could explain excess protons harbored by the water cluster in the inner EC channel (Garczarek et al., 2004, 2005; Lorenz-Fonfria et al., 2017; Wang and El-Sayed, 2001).*
*As I already noted, these continua are all negative and not positive in light-induced FTIR difference spectroscopy, meaning that they cannot explain the building of hydronium ions during the photocycle. They indicate that a protonated water clusters are transient deprotonated and final reformated during the photocycle..*

*-Line 573: "Neither a hydraulic powered noria or an animal powered sakia in the old time, nor an electric powered fluid pump in the modern days, works under such an operating principle that requires energy for a reconstruction of the landscape."*

*Please, explain what do you mean by "requires energy for a reconstruction of the landscape".*

*-Line 576: "Quite the contrary, the landscape is a given constant and energy is absorbed by the to-be-pumped substance to gain its potential energy, such as water is lifted by a noria or pressurized by an electric pump. The energized substance will then flow freely along the largely constant landscape governed by thermodynamics. In the meanwhile, it is critical to prevent a backflow along a far steeper gradient than the forward-driving gradient."*
*I am lost. I do not know that the author is taking about. Please, refocus it.*

*-Line 585: "Alteration of affinities to the pumped substance cannot be an effective strategy because it is against the chemical nature along the translocation pathway and therefore energetically difficult to achieve."*
*Confusing sentence. Please, rewrite it.*

*-Lines 588 to 612: The rest of the Concluding Remarks are confusing and hard to follow divagations about the view of the author about how BR works.*

Dear Editor-in-Chief Nelson,

I am writing to appeal the decision to reject my manuscript PNASNEXUS-2022-00209. Contrary to the summary, the reviewers did not find “serious fault” in my study, and they didn’t explicitly recommend rejection either. Reviewer 1 finds my analysis a success and recommends my method as “a new basis” for future analysis of serial crystallographic data. Reviewer 2 finds this work “stimulating and interesting”. Reviewer 3 finds the intermediate structures resulted by my work “more accurate”. It was not their conclusion that my finding “was quite different from the standard” and “contradict(s) what is solidly known”. These are the precise messages that I convey to the readers. These cannot be used as the reasons to reject my manuscript; these are the very basis to publish this manuscript.

The widely disseminated perception of the light-driven uphill transport of protons by bacteriorhodopsin is far from what I have observed in this study. I traced back the history where discrepancy could have occurred. An early error took place in 1960s when the molecular mechanism was imagined without much of experimental evidence (Jardetzky, 1966). This is not surprising as no one could have guessed it right for the first time when direct sightings of such biochemical reaction were extremely scarce. The fundamental error occurred repeatedly ever since, that is, everybody was trying to prove the early hypothesis, and no one questioned it even when more and more data contradicted the theory. This is because peer reviews only praise those studies that conclude consistently with the previous publication, as who peers are. High-profile journals like PNAS only publish findings that confirm what is already in the literature. The word “only” in the previous statements is perhaps a little too strong. However, this tendency is overwhelming. Academic publishing promotes mutual confirmation and verification, but overwhelmingly hinders falsification. The root course arises from the necessity of mutual support for scientific funding. However, a direct consequence is that everything published becomes the perpetual truth and even a shaky opinion can be reinforced over time into solid evidence. Rejecting this manuscript would be another round of the same mistake.

I will have to lay out every detail in my point-by-point response later. In this appeal letter, I can only focus on several major issues in the big picture. The premise of the debate is the new structures I report in this manuscript that depict unscrambled intermediate states in the photocycle of bacteriorhodopsin. The reviewers agree that these new models are “more accurate” because they are “pure structures” extracted from “mixtures of several intermediate species”. Reviewer 1 also praises the elegance and effectiveness of the methodology and predicts that this would be “a new basis” for future analysis of serial X-ray crystallographic data. The doubt arises when I present a very different mechanism of proton pumping based on these more accurate, pure, intermediate structures. “How the author knows about this from the analysis of the time-resolved X-ray data?” asked Reviewer 3. This question can be applied throughout the manuscript. Time-resolved X-ray data reveal visuals in motion. Each frame is associated with a precise time stamp. It is true that a standalone proton does not scatter X-rays hence not visible in X-ray data. But its association with the surrounding structural elements are visible.

A proton is not visible in vibrational spectroscopy either. What they really measure is the stretching frequency of a chemical bond that is influenced by its association with a proton. Protonation or not is even more questionable in absorption spectroscopy, especially in a rapidly changing protein environment. Spectroscopy does not produce visuals of molecular image that we can see. Spectroscopy critically relies on “assignment”. If a spectral feature is assigned to a structural element correctly, spectroscopy could be very sensitive. The bulk of the spectroscopic results on bacteriorhodopsin were derived in 1970s, 80s, and 90s before a detailed structure of this protein was revealed in 1999 (Luecke et al., 1999). The greater problem is that the ground state structure and a very limited number of cryo-trapped intermediate structures did not trigger a reevaluation of the earlier spectroscopic data for the very reason I point out in the first place. It is simply not the academic culture to change an early story. Structural biologists cannot change the story in spectroscopy either. Interpretation of spectroscopic data under an erroneous structural basis could be very misleading. The obvious example in this case is that the Schiff base has departed from the extracellular half channel, the chamber that houses the resting Schiff base in the ground state, ever since half picosecond, a very early time point. It is critically important to the function of this proton pump that the Schiff base can no longer access anything in its home space since that early time point until nearly the end of the photocycle. Unfortunately, spectroscopic data were largely interpreted in its home space based on a few inaccurate, early indications of the resting structure. It is interesting to learn that Reviewer 3 blames the lack of later intermediate structures after M state for not understanding how protons can be conducted through Arg82. However, when nine accurate structures of bR, I′, I, J′, J, K, L, $M_1$, and $M_2$ that cover the majority of the photocycle, Reviewer 3 pays no interest in what molecular mechanism they could reveal as long as the old story is not changed.

The old story that the reviewers claim as "solidly known" is only a firm belief in the community that originated from an ill-conceived imagination and corroborated blindly by early interpretations of spectroscopic data without a solid structural basis. The root of this belief sounds very convincing. Just like water flowing from high ground to low ground, protons are conducted from chemical groups with low proton affinity to those with high affinity. The essence of the belief is that the proton pump bacteriorhodopsin uses light energy to reverse the gradient of proton affinity so that protons are driven in the direction that they would not flow in dark. This belief deceived everybody into seeking evidence to prove it. According to this laughable pumping principle, energy should be used to dig a canal deeper and deeper from the river front into the village on the hill. Upon the completion of such canal, the villagers would immediately realize that they cannot reach the water deep down in the canal despite the fact that the water does reach the town center. No working water pump, ancient or modern, follows such allosteric model of Jardetzky. In case of successful water pumps, energy has to be used to elevate the water by old designs or to pressurize the water by modern fluid pumps powered by electricity. Bacteriorhodopsin uses light energy to increase the potential energy of the proton, in stark contrast to what was believed that light energy is used to alter proton affinities.

The reviewers seem to admit that there are indeed remaining unknowns about this protein. However, this work could be publishable as long as "without mentioning H+ (proton) transfer different from the standard model". By responding to my question when and how the Schiff base transfers its proton uphill across 11 pH units, Reviewer 3 thinks that "the 'when' has been known for long time". I am free to explain "how" as I wish. But do not change the story of "when" because they have published that. Too bad, when and how are integrated in the same mechanism – the proton gains its potential energy over 11 pH units from the absorbed photon, that is, overcoming one-hundred-billion-fold of proton affinity. This extremely steep uphill gain is only possible at the peak power of this photophysical process within tens of femtosecond. The reviewers insisted that the proton would follow the motion of the Schiff base instead of departing from it until a very late stage in M state. Reviewer 1 made a factual error when stating that the proton "is covalently bound with RSB (retinal Schiff base)". One must understand that the proton, that is, a hydrogen ion, does not carry an electron, therefore by definition, the bond is not covalent. The proton is only attracted to the lone pair of electrons of the Schiff base nitrogen, a much weaker association compared to a covalent bond. Reviewer 2 argues why the Schiff base nitrogen is not stationary if the proton is, as "the Schiff base nitrogen has more inertia." The Schiff base nitrogen is covalently bound with two carbons with one single bond and one double bond. These covalent bonds are far stronger than its association with the proton. As a result, the nitrogen is an integral part of the chromophore, but the proton is only an attachment. Reviewer 3 presented a back-of-the-envelope calculation that results in 600 pN force exerted on the proton if it moves every picometer, that is, 0.01 Å. The intermediate structures refined in this study show 0.6 Å relative motion of the proton within the first 30 fs that would generate a force of 36 nN. A force at the magnitude of hundreds pN or nN is more than sufficient to stretch a protein into a linear polypeptide chain (Takahashi et al., 2018). A non-covalent association of the proton cannot sustain a force at this magnitude. Therefore, the reviewers grossly overestimated the strength of the proton association with the Schiff base.

Most importantly, they chose to ignore my explanation judged by zero mention to the text and figures on this most crucial point: The proton does not follow the departing Schiff base not because it associates with the Schiff base weakly. The determining factor is that the proton associates with both the Schiff base nitrogen and the oxygen of the water simultaneously like this [⩾N:$H^+$:O<], where $H^+$ is the proton. Both associations are comparable in strength and weaker than a regular covalent bond because the proton is charged and does not contribute any electron in either association. One of these two associations must break within the first 30 fs. The proton has to stay with the stationary partner instead of accelerating with the departing partner due to its inertia. The reviewers must not ignore this fact and explain why the proton could break its association with the water and accelerate with the Schiff base if they insist that deprotonation is impossible at this early stage.

It may take years or even decades for this community to reinterpret existing data, to gather additional evidence, and to conduct theoretical simulations to finally clarify the mechanism of proton pumping by bacteriorhodopsin, one way or the other. I request a reevaluation of a revised manuscript of 209. I hope the editors would agree that my numerical analysis of the XFEL data and my crystallographic refinement of these intermediate structures are a complete piece of work that ought to stand alone and be publishable on Nexus. These eight intermediates in the photocycle of bacteriorhodopsin are pure structures at room temperature. Such accurate depiction of the intermediates has never been determined before. They are the indispensable foundation eagerly awaited by the community. Future advance in structural studies, spectroscopy, or computation require such structural basis. For example, quantum mechanical calculation is clearly needed and will be a separate piece of work carried out by experts. Intermediate species described in 208 (Ren, 2022) and 209, such as the transient expansion and contraction of the retinal binding pocket, are the guiding structures for these computational studies.

From a different perspective, Nexus as a newly launched journal also needs to publish this manuscript. I wish that the editors can see this as a turning point in both bacteriorhodopsin research and applications of time-resolved serial crystallography. This work, if published, no doubt will trigger a broad-based debate, thus will be highly cited. The interest will not be limited in bacteriorhodopsin and related channels and transporters. The success in isolating pure conformational species from inevitable mixtures, as agreed by the reviewers, will attract even wider attention. Because experimental observations of concurrent molecular events are common.

Sincerely yours,

Zhong Ren

References

Jardetzky, O. (1966). Simple allosteric model for membrane pumps. Nature *211*, 969–970. https://doi.org/10.1038/211969a0.

Luecke, H., Schobert, B., Richter, H.-T., Cartailler, J.-P., and Lanyi, J.K. (1999). Structure of bacteriorhodopsin at 1.55 Å resolution. J. Mol. Biol. *291*, 899–911. https://doi.org/10.1006/jmbi.1999.3027.

Ren, Z. (2022). Photoinduced isomerization sampling of retinal in bacteriorhodopsin. PNAS Nexus *1*. https://doi.org/10.1093/pnasnexus/pgac103.

Takahashi, H., Rico, F., Chipot, C., and Scheuring, S. (2018). $\alpha$-helix unwinding as force buffer in spectrins. ACS Nano *12*, 2719–2727. https://doi.org/10.1021/acsnano.7b08973.

### Directional Proton Conductance in Bacteriorhodopsin Is Driven by Concentration Gradient, Not Affinity Gradient

Zhong Ren

Department of Chemistry, University of Illinois at Chicago, Chicago, IL 60607, USA
Renz Research, Inc., Westmont, IL 60559, USA


ORCID 0000-0001-7098-3127

**Response to peer review**

I would like to express my appreciation to the editor and all reviewers. Their comments are constructive and representative, from which I learn a great deal on how to revise my manuscript so that the newly refined structures and the proposed mechanism can be presented more effectively to the diverse community of bacteriorhodopsin studies. I believe that the revised manuscript has been much improved by incorporating every suggestion and by answering every question. The mainstream view on the subject of bacteriorhodopsin (bR) had the historical chance to develop into an overwhelmingly strong status during the past 50 years. Here I plead with the reviewers to listen to my lone voice open-mindedly. Sometimes a newcomer could bring a fresh perspective from outside the box. I would like to attract the reviewers' attention to the coherence in my answers to many questions posed in this and the companion manuscripts (Ren, 2022). These answers seemingly addressed to some unrelated questions actually support one another. I would also like to note that the discussion on several key mutants provide crucial cross validation to the proposed mechanism. However, due to the length limit, I have to move these discussion into Supplementary. Before I dive into all the points raised by the reviewers, I would like to sketch an overview to focus on several major issues in the big picture. The comments are then copied verbatim in *italics* followed by my response. The topics are arranged by relevance to one another.

***Deconvoluted intermediate structures***

The premise of the debate is the new structures I report in this and the companion manuscripts that depict unscrambled intermediate states in the photocycle of bacteriorhodopsin (Ren, 2022). The reviewers agree that these new models are "more accurate" because they are "pure structures" extracted from "mixtures of several intermediate species". Reviewer 1 also praises the elegance and effectiveness of the methodology and predicts that this would be "a new basis" for future analysis of serial X-ray crystallographic data. Reviewer 2 finds this analysis "stimulating and interesting". Are the newly refined intermediate structures merely an incremental improvement from the published ones? Or could they reveal transformative understanding of the pumping mechanism? What is the difference anyway?

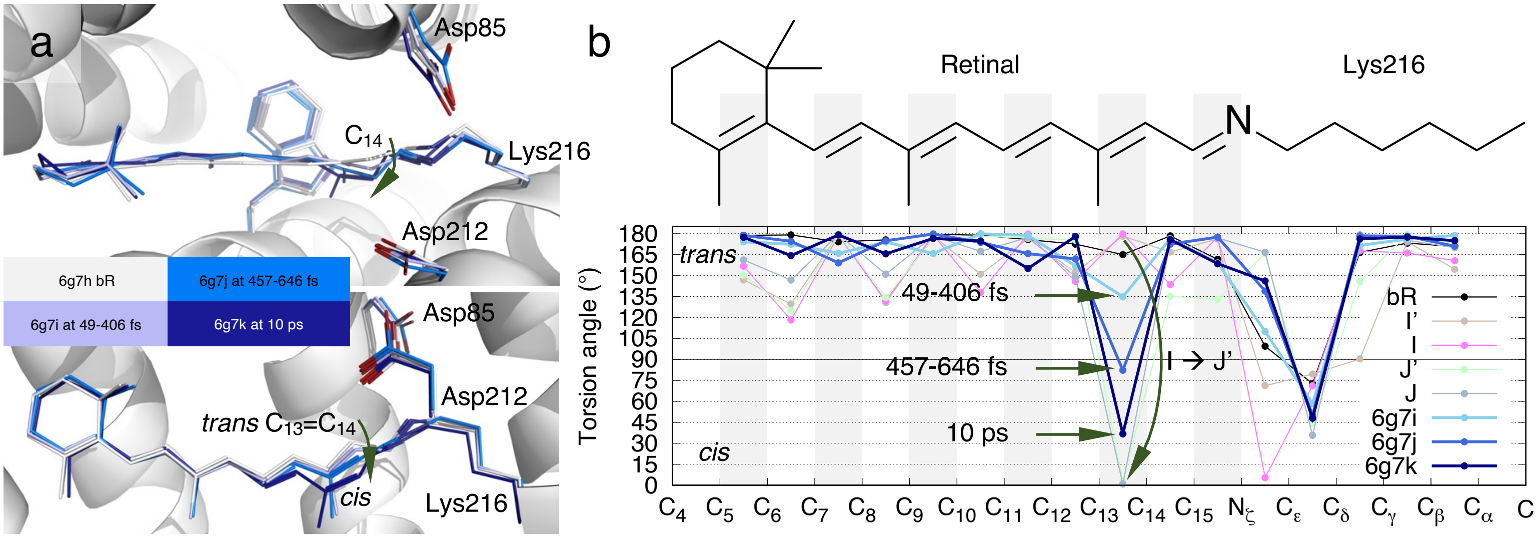


Figure A. Photoisomerization at $C_{13}$=$C_{14}$ of retinal in bR presented by Nogly et al. (a) Published structures of Nogly et al. The chromophore conformations in darker and darker blues are overlayed with the resting state in white. Photoisomerization is interpreted as a gradual change in conformation from tens to hundreds of fs to ps. Even at 10 ps, $C_{13}$=$C_{14}$ is not yet in perfect *cis* (Nogly et al., 2018). Kovacs et al. presented a similar rotation. (b) Torsion angles calculated from the published structures of Nogly et al. The chemical structure of the chromophore on top is aligned to the horizontal axis. Double bonds are shaded in gray. The torsion angles indicate no significant rotation of single bonds captured by the structures of Nogly et al. However, $C_{13}$=$C_{14}$ is transitioning gradually from *trans* to *cis*. In contrast, the refined structures of I and J' in my analysis show a complete photoisomerization from perfect *trans* to perfect *cis* directly.

The previous interpretations of the mixed signals of multiple intermediate species presented a rotation of the $C_{13}$=$C_{14}$ double bond in the counterclockwise direction if viewed from the Schiff base (SB) end of the retinal (Fig. Aa) (Kovacs et al., 2019; Nogly et al., 2018). This rotation starts at the time point of 49-406 fs (6g7i) and proceeds continuously through hundreds of fs to ps. At 10 ps time point (6g7k), this rotation still does not complete and results in the remaining twist of 37° in the torsion angle of

$C_{13}=C_{14}$ (Fig. Ab). An experimentally determined double bond with a torsion angle of 82°, for example 6g7j, indicates that this twisted double bond conformation is more populated than any other conformation at hundreds of fs. Such interpretation is in direct contradiction of the current view in molecular dynamics. In theory, a double bond conformation spends most time in either energy well of *trans* or *cis*. The transition time between these energy wells is relatively short. A twisted double bond conformation cannot be significantly populated compared to two discrete populations in *trans* and *cis*, that is, cannot be observed, unless an additional energy well emerges near the torsion angle of 90°, which no evidence supports. The gradual rotation of $C_{13}=C_{14}$ double bond presented in the refined structures of Nogly et al. and Kovacs et al. was a misinterpretation of the mixed signals of multiple intermediate species as a single conformer. The gradual rotation does not occur in isomerization, rather it merely reflects the gradual population shift from discrete states of *trans* to *cis*. Reviewer 2 asked "what the mistake was" in previous studies. Here is one of the mistakes. The same mistake occurred repeatedly in structural refinement and interpretation without any regard to the basic thermodynamic principle. For another example, a cryo-trapped structure after light illumination was refined to two severely twisted double bonds $C_{13}=C_{14}$ and $C_{15}=N_{\zeta}$ far from *trans* and *cis* (2ntw), which formed the structural basis for the erroneous conclusion that the H-bond network involving the SB in the dark resting state persists until the proton transfer in M state (Lanyi and Schobert, 2007).

In stark contrast, my analysis presents that perfect *trans* to perfect *cis* isomerization occurs in a single move during the transition of I → J′, the first report of a perfect 13-*cis* retinal in bR at hundreds of fs (Fig. Ab). This is not to say that the transitioning double bond never rotates through a twisted conformation passing 90° dihedral angle. This interpretation obeys statistical mechanics and acknowledges that a twisted conformation near 90° is never populated significantly, therefore cannot be observed. Are these two different interpretations of the same data equivalent and inconsequential to mechanistic understanding? The discrete states in *trans* and *cis* presented here result in two sharp U-turns of the SB $N_{\zeta}$ atom not observed in the previous interpretations, the first one at I′ around 30 fs and the second one during I → J′ isomerization at ~500 fs (Fig. 2a and S3). The first U-turn of $N_{\zeta}$ at ~30 fs marks the peak power of the entire photocycle that was not observed before. Its implication is enormously important to the mechanistic understanding of the photophysical deprotonation from the SB (see below).

***Time-resolved crystallography***

"How the author knows about this from the analysis of the time-resolved X-ray data?" asked Reviewer 3. This question can be applied throughout the manuscript. Time-resolved X-ray data reveal visuals in motion. Each frame is associated with a precise time stamp. A conformational change observed at a precise time point indicates the velocity of the conformation change, which further hints the minimal acceleration and force required to achieve the observed conformational change. It is true that a standalone proton does not scatter X-rays hence invisible in X-ray data. But its association with the surrounding structural elements are visible. It is foolish to dismiss a proton location in a crystallographic structure on the grounds of the lack of X-ray scattering by protons. Can we dismiss a hydrogen atom near the midpoint of a good H-bonding distance? Can we dismiss three hydrogen atoms on a methyl group since no electron density is observed?

Unfortunately, the great promise of cryo-trapping crystallography, time-resolved crystallography at room temperature, especially, the ultrafast studies using X-ray free electron lasers, was not delivered sooner. The rapid molecular motion blurs the visual picture because multiple conformations are captured simultaneously – exact the same difficulty encountered in spectroscopy. This work deconvolutes pure species from mixtures. "How [does] the author know[s] about this from the analysis of the time-resolved X-ray data?" Now I can see. The readers can see. Even if I cannot see every hydrogen or proton, prior knowledge in chemistry is the foundation for either a deduction or speculation, whichever the readers can judge.

***Spectroscopy***

A proton is not visible in vibrational spectroscopy either. What they really measure is the stretching frequency of a chemical bond that might be influenced by its association with a proton. Protonation or not is even more questionable in absorption spectroscopy, especially in a rapidly changing protein environment. The absorption property of bR during its entire photocycle has been thoroughly and repeatedly measured. Unfortunately, the implication of the absorption property was mistakenly inferred ever since the very beginning, even if the original authors had drawn the right conclusions. In 1973, Oesterhelt & Hess first demonstrated that bR photoconverts between two absorption maxima 568 and 412 nm (Oesterhelt and Hess, 1973), the ground state and the later named M state, respectively. The next year, Lewis et al. corrected a previous mistake (see below) and concluded from their resonance Raman (RR) experiments that the SB is protonated in the ground state and deprotonated in the 412-nm-absorbing state (Lewis et al., 1974). These conclusions stood the test of time. However, the strong blue shift of 568→412 nm and the SB deprotonation were mistaken as two correlated events, despite the clear statements of Lewis et al.: "the retinylidene lysine linkage in bacteriorhodopsin is protonated … the proton is removed before bacteriorhodopsin reaches the 412 nm intermediate". There is absolutely no hard evidence to correlate an absorption maximum at 412 nm to the molecular event of SB deprotonation – continued answers to the question of "what the mistake was". A loose correlation was derived largely from studies on model compounds. But no strict correlation was ever established (Heyde et al., 1971). An environmental change occurring to the same isomer of retinal could cause greater blue shift than a change in protonation state. For example, the absorption of an unprotonated, neutral SB in an all-*trans* retinal *in vacuo*, that is, without interaction with solvent, peaks at 487 nm, while the absorption of a protonated SB in an all-*trans* retinal peaks at 444 nm in methanol and in near UV in ethanol, way further blue shifted from the neutral SB (Andersen et al., 2005; Heyde et al., 1971). What factors dominate the absorption property of retinal? These factors in an approximately descending order of influence are coplanarity, charge distribution, environment, protonation, isomers in *trans* or *cis*, etc. Surprisingly, protonation state and isomers are rather minor factors compared to the geometry and environment of the chromophore. Therefore, absorption maxima other than 412 nm in no way indicate a protonated SB. Here I suggest transitional color shift instead of color shift with respect to the ground state, an arbitrary reference in the photocycle. It becomes more apparent that the periodic color change repeats twice in the photocycle (Fig. B). Each period corresponds to one of the two protonation states. Both start with a dramatic swing from a strong transitional blue shift to a strong transitional red shift. This is due to proton transfer and violent conformational changes associated with isomerization. Both periods gradually settle to a strong blue shift. However, the first period during the unprotonated SB ends with a deep-violet-absorbing M state, while the second period during the protonated SB finishes in a yellow-absorbing ground state. Far more likely, the stepwise blue shifts during $J^{625} \rightarrow K^{610} \rightarrow KL^{596} \rightarrow L^{550} \rightarrow M^{412}$ indicate that the strong influences from the stereochemical distortion and charge redistribution over the chromophore subside gradually and give way to the influence of the neutral, unprotonated SB throughout this process over 12 decades. These four consecutive transitional blue shifts indicate the early deprotonation at the beginning of the photocycle. The blue-absorbing I state before isomerization marks the initial deprotonation from the SB. However, immediately after photoisomerization, J′ and J species are highly unusual and strained in their conformation. The normal blue shift induced by the deprotonated SB is only a minor factor hence masked by other major factors until the chromophore is relaxed enough. This relaxation process is well depicted in the refined structures of J′, J, K, L, $M_1$, and $M_2$.

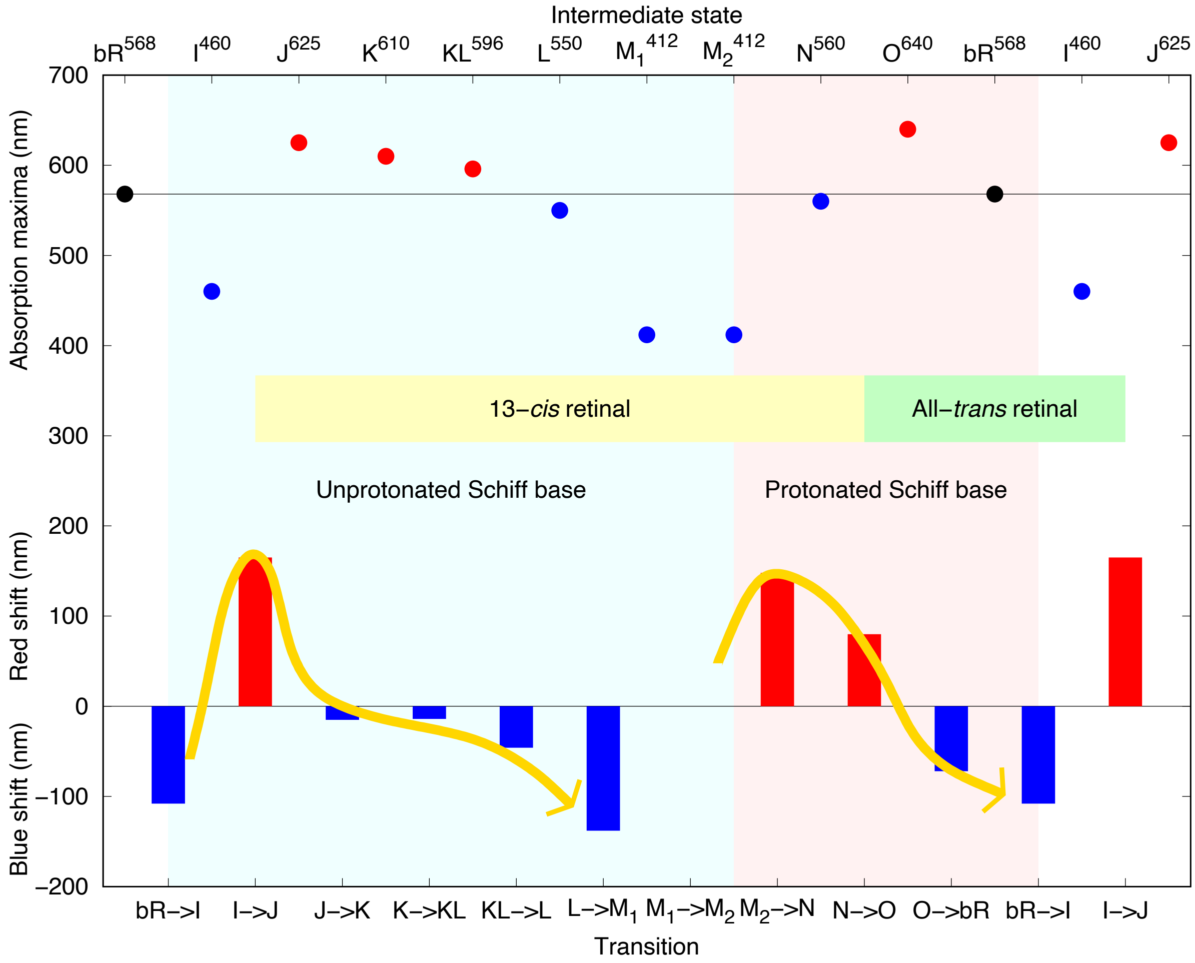


Figure B. Color shifts in photocycle of bacteriorhodopsin. Absorption maxima are plotted in the top part. With respect to the ground state, red or blue shift of each intermediate state is represented by a colored dot. Intermediate states with all-*trans* and 13-*cis* retinal are marked with horizontal bars. Red or blue shift of a transition between two consecutive states is plotted in the bottom part with red or blue bars. Protonation states are shaded with pink and light blue background.

The bulk of the spectroscopic results on bR were derived in 1970s, 80s, and 90s before a detailed structure of this protein in the ground state was revealed in 1999 (Luecke et al., 1999). Interpretation of spectroscopic data under an erroneous structural basis could be very misleading. The obvious example in this case is that the SB has departed from the extracellular half channel, the chamber that houses the resting SB in the ground state, ever since half picosecond, a very early time point. It is critically important to the function of this proton pump that the SB can no longer access anything in its home space since that early time point until nearly the end of the photocycle. That is to say, there is no water connected to the SB ever since isomerization at hundreds fs. Even if there is, that is a water other than Wat402. Arg82, Asp85, Asp212, and everything used to be around the SB are no longer relevant. Unfortunately, spectroscopic data were largely interpreted in its home space, or an environment similar to the ground state, based on a few inaccurate, early indications of the resting structure, let alone any consideration of the creased S-shape of the retinal gradually flattening in the expanded and contracted retinal binding pocket (Ren, 2022). See specific examples below.

Spectroscopy does not produce visuals of molecular image that we can see. Spectroscopy critically relies on “assignment”. If a spectral feature is assigned to a structural element correctly, spectroscopy could be very sensitive. As Reviewer 2 argued, “if the bacteriorhodopsin photocycle would indeed start with the deprotonation of the Schiff base in retinal”, the deprotonated SB would absorb “at 380 nm, instead of 568 nm” since the beginning of the photocycle. The intermediates during the photocycle J, K, L, M, etc. were differentiated by their absorption maxima. Their corresponding structures, including conformation, protonation, and charge distribution, were never directly observed but assigned or interpreted. Without a correct structural basis, the stepwise blue shifts since photoisomerization until reprotonation would never been interpreted as the indication for the very early deprotonation.

There is in fact no disagreement on an unprotonated, neutral SB in M, nor any dispute on the pronation of Aps85 in M. The spectroscopic evidence on a protonated SB persisting until M is actually weak. One of the earliest RR studies already noticed that the signature of deprotonation from the SB does not follow the kinetic rise of M state, but appears before 8 μs when there is no sign of M, where 8 μs is very much limited by the technology back then (Marcus and Lewis, 1977). The literature on the subject of deprotonation from the SB is chaotic and largely evades direct evidence that supports a lasting protonated SB prior to M formation. For example, the frequency difference of the SB double bond stretching in water and heavy water has been considered as an indication of H-bond strength to the SB. Very small difference $\Delta\nu_{H\text{-}D}$ prior to M formation compared to the ground state indicates the broken H-bond between Wat402 and the SB (Kandori et al., 2002). If the H-bonding has been lost before M formation, how does the SB deprotonate when M forms? Does it support an early deprotonation before the disappearance of the H-bond?

***Mechanistic understanding of proton pump***

The unscrambled pure intermediates reported here allow me to observe the structural basis of the bR photocycle from tens of fs to $M_2$ that shows an evolution of structural events far from the widely disseminated perception of the light-driven uphill transport of protons. I traced back the history where discrepancy could have originated. An early error took place in 1960s when the molecular mechanism was imagined without much of experimental evidence (Jardetzky, 1966) – more answers to the question of "what the mistake was". This is not surprising as no one could have guessed it right for the first time when direct sightings of such biochemical reaction were extremely scarce. The fundamental error occurred repeatedly ever since, that is, everybody was trying to prove the early hypothesis, and no one questioned it even when more and more data contradicted the theory. What the reviewers claim as "solidly known" is only a firm belief in the community that originated from this ill-conceived imagination and corroborated blindly by early interpretations of spectroscopic data without a solid structural basis. The strong faith originates from a doctrine similar to water flowing from high ground to low ground, that is, protons are conducted from chemical groups with low proton affinity to those with high affinity. The essence of the belief is that the proton pump bR uses light energy to alter proton affinities so that protons are driven in the desired direction. This belief is explicitly stated by Stoeckenius et al.: "The Schiff base should further drastically change its p*K* during that movement and inject its proton into the external part of the channel… It should then return to its original position and relatively high p*K* to accept a proton from the cytoplasmic side of the membrane via the cytoplasmic segment of the channel. The energy necessary for the proton transfer against its electro-chemical gradient must be derived from the photon energy absorbed and stored in transient conformational changes of the protein." (Stoeckenius et al., 1979). This belief deceived everybody into seeking evidence to prove it. According to this laughable pumping principle, energy should be used to dig a canal deeper and deeper from the river front into the village on the hill. Upon the completion of such canal, the villagers would immediately realize that they cannot reach the water deep down in the canal despite the fact that the water does reach the town center. No working water pump, ancient or modern, follows such allosteric model of Jardetzky. In case of successful water pumps, energy has to be used to elevate the water by old designs or to pressurize the water by modern fluid pumps powered by electricity. It is the common sense that bR uses light energy to increase the potential energy of the proton (Fig. 1), in stark contrast to what was believed that light energy is used to alter proton affinities.

The reviewers seem to admit that there are indeed remaining unknowns about this protein. However, my work could be publishable as long as "without mentioning H+ (proton) transfer different from the standard model". "Focus only on the structural changes", recommended Reviewer 1. By responding to my question when and how the SB transfers its proton uphill across 11 pH units, Reviewer 3 thinks that "the 'when' has been known for long time". Although "how" is not clear, the story of "when" has been published many times by many people. Therefore, there is no room to change; don't even question. Too bad, when and how must be integrated in a single theory of pumping mechanism regardless of what that theory is. How could a proton gain its potential energy over 11 pH units from the absorbed photon, that is, overcoming one-hundred-billion-fold of proton affinity, escape from the protonated SB with a p$K_a$ of 13.3, and form a carboxylic acid of a p$K_a$ of 2.2? How does this seemingly impossible molecular event occur? The mainstream insists that this occurs at tens of μs because the photoinduced conformational changes reduce the p$K_a$ of the protonated SB but increase that of the carboxylic acid. However, no one could explain how to change p$K_a$ and demonstrate how much change is achievable. Let's assume each partner of this proton transfer does achieve a change of six p$K_a$ units, that is, a million-fold of proton affinity. Now the protonated SB somehow achieves a p$K_a$ of 7.3 and the carboxylic acid has a p$K_a$ of 8.2, approximately one unit higher than that of the SB. Therefore, the proton transfers spontaneously. At a prolonged equilibrium, most of the carboxylates are neutralized. But a significant fraction of the SB remains protonated. That fraction could range from a quarter to a third depending on pH. More decisive proton transfer would require even greater p$K_a$ changes. No one is close to demonstrate that a million-fold change in proton affinity occurs simultaneously to each partner of the proton transfer. Because this is against the chemical nature of a protonated SB and a carboxylic acid, therefore remotely unachievable, regardless of what conformational changes have been induced by the photon absorption. Nature does not work like that.

Why do we have to keep our faith in the allosteric model of Jardetzky that does not explain the working principle of any pump, let alone the molecular pump bR? The extremely steep uphill gain of > 11 pH units is only possible at the peak power of this photophysical process within tens of femtosecond, not a spontaneous equilibrium. The peak power of the photocycle achieves an uphill gain of > 15 pH units to form a strong acid of hydronium. The rest is a spontaneous downhill flow (Fig. 1). The formation of the carboxylic acid at tens of μs agrees very well with many spectroscopic results. However, the departure of the proton from the SB is where the doubt arises. It could be instinctive to accept that the proton would follow the motion of the Schiff base instead of departing from it. Reviewer 1 made a factual error when stating that the proton "is covalently bound with RSB (retinal Schiff base)". One must understand that the proton, that is, a hydrogen ion, does not carry an electron, therefore by definition, the bond is not covalent. The proton is only attracted to the lone pair of electrons of the SB nitrogen, a much weaker association compared to a covalent bond. Reviewer 2 argues why the SB nitrogen is not stationary if the proton is, as "the Schiff base nitrogen has more inertia." The SB nitrogen is covalently bound with two carbons with one single bond and one double bond. These covalent bonds are far stronger than its association with the proton. As a result, the SB nitrogen is an integral part of the chromophore, but the proton is only an attachment. Reviewer 3 presented a back-of-the-envelope calculation that results in 600 pN force exerted on the proton if it moves every picometer, that is, 0.01 Å. The intermediate structures refined in this study show 0.6 Å relative motion of the proton within the first 30 fs that would generate a force of 36 nN. A force at the magnitude of hundreds pN or nN is more than sufficient to stretch a protein into a linear polypeptide chain and could be strong enough to rupture covalent bonds (Grandbois, 1999; Takahashi et al., 2018). A non-covalent association of the proton cannot sustain a force at this magnitude. Therefore, the reviewers grossly overestimated the strength of the proton association with the SB.

However, the proton association with the SB weaker than a covalent bond is not the reason why it departs from the SB. The reviewers chose to ignore my explanation judged by zero mention to the text and figures on this most crucial point: The proton associates with both the SB nitrogen and the oxygen of the water simultaneously like this $\geqslant N_{\zeta}:H^{+}:O<H_2$, where $H^+$ is the proton. Both associations are weaker than a regular covalent bond because the proton is charged and does not contribute any electron in either association. The proton association with the water is weaker than that with the SB if the negative p$K_a$ of a hydronium is considered. However, the positive charge of the proton is more distributed than we realize although we still call that proton proton. A proton in an aqueous system associates with waters in significantly covalent nature (Swanson and Simons, 2009). Therefore, two proton associations in $\geqslant N_{\zeta}:H^{+}:O<H_2$ could be comparable in strength due to the delocalized positive charge like this $\geqslant N_{\zeta}:H\text{-}O^{+}<H_2$. One of these two associations must break within the first 30 fs. The proton has to stay with the stationary partner instead of accelerating with the departing partner due to its inertia. The conformation of the retinal, the anchor Lys216, Wat402, and its interactions with Asp85 and 212 are all well resolved in the electron density maps immediately before and immediately after photoisomerization (Ren, 2022). The location of the proton is well deducible. This analysis based on basic knowledge in chemistry, mechanics, and thermodynamics cannot be easily labeled as “speculative” on the grounds of invisible proton in X-ray data. I do not pretend an explicit observation of the electron density well isolated for this proton with partially delocalized charge. Instead, I explicitly state in the manuscript that this conjecture “is not based on an observation of the scattered X-rays by the proton” (Line 375-376) so that the readers can make their own judgement.

It may take years, hopefully not decades, for this community to reinterpret existing data, to gather additional evidence, and to conduct theoretical simulations to finally clarify the mechanism of proton pumping by bR, one way or the other. I agree completely that quantum mechanical calculation is necessary and will be a separate piece of work carried out by experts. Intermediate species revealed in this work, such as the transient expansion and contraction of the retinal binding pocket, are the guiding structures for these computational studies. I hope the editors and reviewers would agree that my numerical analysis of the XFEL data and my crystallographic refinement of these intermediate structures are a complete piece of work that ought to stand alone and be publishable in Nexus. These eight intermediates in the photocycle of bR are pure structures at room temperature. Such accurate depiction of the intermediates has never been determined before. They are the indispensable foundation eagerly awaited by the community. Future advance in structural studies, spectroscopy, or computation require such structural basis. I wish that the editors and reviewers can see this as a historical turning point in both bR research and applications of time-resolved serial crystallography. The interest will not be limited in bR and related channels and transporters. The success in isolating pure conformational species from inevitable mixtures will attract even wider attention. Because experimental observations of concurrent molecular events are common.

***Point-by-point response***

*Reviewer 1: In this work, Dr. Ren elegantly succeeded in extracting the pure structures of each photo intermediate by applying SVD analysis for more than 40 structures previously reported in different serial X-ray crystallography studies. This would make a new basis for the methodology of the analysis of serial X-ray crystallographic data and be useful for understanding the dynamic events of a wide range of proteins.*

*Reviewer 2: The manuscript revises crystallographic data from recent serial femtosecond crystallography experiments on bacteriorhodopsin. On this basis the results are re-interpreted and a different conclusion is reached. The conclusions are questioning the current understanding of the photocycle in bacteriorhodopsin. This work is stimulating and interesting.*

I appreciate the encouraging words of the reviewers.

*Reviewer 1: Bacteriorhodopsin (BR) is the best characterized light-driven proton pump that can convert photon energy to the proton motive force to energize the cells with compact seven-transmembrane architecture and an all-trans-retinal chromophore. The mechanism of H+ pumping by BR was studied in detail by a pile of biochemical, spectroscopic, theoretical, and structural biological methods for five decades leading a sophisticated model in which H+ is sequentially transferred between retinal Schiff base (RSB) and acidic residues (Asp/Glu) along with the appearance of spectrally distinguishable multiple photo intermediates called K, L, M, and so on. Especially, the time-resolved serial X-ray crystallography using X-ray free-electron laser or synchrotron radiation enabled real-time observation of 3-D structural change of the entire body of the protein at room temperature and provided unprecedented detailed structural information. The structures observed at specific time points after the light illumination, however, are mixtures of several intermediate species so that it is difficult to correspond the observed conformational changes to the respective intermediates.*

Reviewer 1 provides a brief but nice recap of the bacteriorhodopsin research and points out the key difficulty that the entire field faces in both spectroscopic and structural studies – mixtures. Because of mixture of intermediates, misinterpretation of data persists throughout the history. For example, the resting SB was first interpreted as unprotonated and ready to take a proton (Mendelsohn, 1973). This misinterpretation of data had to be corrected regardless of the possible cause – either a misassignment of a vibration mode or a later intermediate mixed into the sample. As we know now, the resting SB is already protonated (Lewis et al., 1974; Mendelsohn, 1976). Mendelsohn should have doubted his original conclusion as soon as he drew it. Just like water, very small fraction of the molecules would lose a proton under a neutral pH. A resting SB would hold on to a proton because both water and protonated SB have similar p$K_a$ of ~14. That is to say, their Raman experiments only confirmed something of extremely high probability. When we conclude something extremely improbable in thermodynamics, shall we doubt ourselves first? However, the lesson in history was not learned.

*Reviewer 3: To give a background, the general "believe" is that photoisomerization of the all-trans retinal with a protonated SB remains in a protonated 13-cis conformation until the L intermediate. In the L to M transition (tens of microseconds) the SB deprotonated in favor of Asp85, and soon a proton is released from the proton release complex (PRC) in the EC domain. The SB reprotonates from Asp96, a residue in the CP domain. Later, Asp96 recovers it proton from the CP medium, completing the proton pumping. The initial state is recovered in few tens of milliseconds as the protonated 13 cis-retinal reisomerizes to all-trans and Asp85 transfers its proton to the PRC.*

This is a nice recap of the mainstream view that had its historical opportunity to develop into the current status. This mainstream view is indeed what I challenge in this study.

*Reviewer 3: Ren provides a manuscript that feeds into published time-resolved X-ray crystallographic data on the bacteriorhodopsin photocycle to determine more accurate structural models for the intermediate states. Then, he uses the obtained structural model to provide a very personal view on how bacteriorhodopsin proton pumps.*

Indeed, this manuscript describes my personal view on the pumping mechanism of bR. This coherent mechanism will soon replace the primitive, yet stubborn, view in the mainstream.

*Reviewer 2: Further points: 1) The author has mentioned several times that previous studies were mistaken. But it was not clear to me what the mistake was.*

Usually, people are angry at me because I point out mistakes too clearly. If this is the request, I will try to point out explicitly where the mistakes are. But this is going to annoy a lot of people.

*Reviewer 3: While I am the first one that think some speculative science help to bring new ideas to the discussion and helps the progress, the problem is that Ren makes innumerable claims that contradict what it is solidly known. As I will made clear below by commenting some of the claims in the paper that I think are wrong, the present MS dismisses, omits, or ignores many solid experimental solid results that contradicts the core of his proposed proton-pumping mode. These critical comments are mixed with other comments and questions I had while reading the MS. All of them are provided below.*

I welcome all criticism and comments. As I respond to each point raised, many "solid experimental results" turn out to be misassignment, misinterpretation, or misunderstanding. A common cause of mistakes is the mixture of intermediates or concurrent molecular events. In crystallography, intermediate structures were often refined against diffraction data from mixtures of multiple species induced by light. In spectroscopy, observed spectral features were often interpreted as a single cause, most likely the one that would be consistent with previous publications. Transient structural distortion was either unknown or was not taken into account, which caused the greatest misunderstanding for the absorption property of bR during its photocycle. Once a correct interpretation is realized, "innumerable" experimental observations suddenly fall into place in agreement with everything. I have expended my interpretation of the previous experiments considerably. This is the coherence that I point out at the beginning.

*Reviewer 1: These concerns need to be clarified before considering the model of this work publishable. Otherwise, I strongly recommend to the story be significantly changed to focus only on the structural changes other than the position of H+, which can be firmly observed by X-ray structural analysis, without mentioning H+ transfer different from the standard model. Since the author's SVD analysis is highly effective to extract the pure structural change occurring in each photo intermediate and would be useful also for the serial X-ray crystallography in the future, it would be better to avoid indicating the fact which cannot be explicitly supported by the results.*

I appreciate the recommendation. Yes, I agree completely that we conclude as far as our data allow and not extrapolate beyond what the data could support. However, this is not because of the preexistence of a "standard model" out there. We must evaluate what the "standard model" was based on. I traced back the root of the "standard model". The allosteric model of Jardetzky and the common line of thinking on proton affinity led by Stoeckenius blatantly disregard the principles in thermodynamics. Just like Mendelsohn's original conclusion that the SB is to be protonated in the ground state, the "standard model" is thermodynamically impossible.

*Reviewer 3: Line 186: "Q1) How does the protonated SB with a pKa of 13.3 transfer its proton to the acidic acceptors with a pKa of 2.2 and when?" I think that the "when" has been known for long time: in the L to M transition. The author can check numerous reviews on BR that have been published by different authors.*

I pose the question not because I am unaware of the literature. I pose the question exactly because people in this field have been fooled by the literature for too long. So, I traced back the history. I found that the first mistake was innocent. No one could have guessed it right in the first place without much of experimental observation. The real mistake is that people believe in the literature instead of question the literature.

*Reviewer 1: On the other hand, the suggested model involving ultrafast deprotonation of RSB in femtosecond time region and concentration driven H+ release is quite different from the standard model (e.g. shown in Ernst et al. Chem. Rev. (2014) in the reference list in the manuscript) in which deprotonation of RSB occurs in ~50 μs on the formation of the M intermediate from the L, and dynamic pKa change of RSB and acidic residues is the key element to drive the vectorial H+ transfer.*

Clearly, the first major contradiction between my analysis and the consensus view is the time of SB deprotonation, ~30 fs at the beginning of the photocycle vs. ~50 μs at M formation. I have expanded my arguments considerably. The second major contradiction is more fundamental as the title states, concentration driven, not affinity driven. However, these two aspects are inseparable. None of these contradictions are due to my ignorance of the literature or carelessness. These are the precise messages I would like to convey to the readers. This field of bacteriorhodopsin studies has made a series of mistakes throughout the history. Some of them could be technical errors. But the fundamental ones are due to misunderstanding or disregard of the principles in thermodynamics. This is why this work together with the already published companion article (Ren, 2022) and upcoming ones represent a turning point in our understanding of the operating mechanism of this molecular pumping device. I hope this string of papers would trigger a reevaluation of the old thinking.

*Reviewer 3: -Line 343: "the "proton transfer" from the SB to the proton acceptor, as commonly stated in the literature, is not a single event. Instead, it spans ten decades in time with multiple asynchronous molecular events." I disagree. The proton transfer from the SB to Asp85 takes place in the L to M transition. It is true that this transition does not follow a single exponential and, thus, is more complex and expands certain time. But the deprotonation of the SB does not start until the microseconds accordingly to what it is known.*
*Reviewer 3: Line 371: "If a deprotonation is unsuccessful at the peak power of the photocycle, there will be no further opportunity to translocate a proton from a fair base to a decent acid uphill against 11 pH units under equilibrium." This statement is not supported by any evidence, besides of the assumption that the photocycle relays on this early deprotonation to pump protons, which I do not think to be the case.*

One has to offer at least a speculation on how such a proton transfer takes place during L → M transition at tens of μs. This proton transfer uphill by 11 orders of magnitude does not occur spontaneously. But it does happen in bR. 50 years of research has to attempt some answers. This field can no longer bypass this problem. This work offers a coherent answer. Unfortunately, a number of previous assumptions must be overturned. It is good that what I overturned are those that disobey thermodynamic rules.

*Reviewer 3: -Line 194: "No convincing evidence can directly demonstrate a photoinduced reversal of the drastically different proton holding abilities along the EC half channel to facilitate a spontaneous proton conductance that would require an ascending order of pKa's." I do not understand what the author means here.*

It means that we cannot disregard the principles in thermodynamics. A spontaneous proton conductance, as an ensemble, could only go one way – from a chemical group of lower proton affinity to that of a higher proton affinity. Without an ascending order of p$K_a$, one must explain how the proton conductance takes place by force instead of spontaneously. Blatant disregard for the thermodynamic principles occurs everywhere in the literature. For example, a proton conductance was suggested as SB (13.3) → water (-1.7). → Thr89 (-3) → Asp85 (2.2), where the p$K_a$ values of the corresponding acids are in parentheses (Nango et al., 2016). Without an explanation on how the p$K_a$ values could change into an ascending order or how some other unknown force magically functions, the claim of such proton conductance was successfully published in Science magazine. How could Iwata, Neutze, and co publish something like that? Because they were not the first one; everybody says so. Often a quantum mechanics or molecular dynamics simulation is used as a justification that such an event is possible. No doubt such a rare event at a single molecular level could be "possible". However, one must understand that for every proton successfully conducted in the wishful direction, a billion protons will flow in the opposite direction if the affinity gradient is against your wish. Declaration of the possibility does not help the readers to understand the mechanism of the directional proton conductance.

*Reviewer 3: -Line 383: "Incidentally, all previous hypotheses on the mechanism of SB deprotonation imply a formation of hydronium at one point or another (Lanyi and Schobert,*

*2007; Nango et al., 385 2016). It would require a donor group with a pKa more negative than -1.7 to reliably form a hydronium by the means of equilibrium." Basically, all proton transfer reactions in solution are mediated by the transient protonation of water. Simulations at the QM level in BR also show that proton transfer reaction between groups in BR are possible and mediated by the transient protonation of water molecules, as shown by different authors (see works by Bondar and co-workers, by Gerwert and co-workers, or by Elstner and co-workers). The key is that this "hydronoium ions" hold the proton for a very small fraction of time, while the author seems to suggests that between the deprotonation of the SB in few fs until the potonation of Asp85 in tenths of microseconds, W401 stays protonated.*

One must understand what it means by quantum mechanically possible. It means that a certain event could take place, even if it takes place once. It doesn't describe the rate or frequency of such event. If a hydronium is required in the reaction, one must show which group with a $pK_a < -1.7$ would be the source of the proton. Otherwise, the rate of hydronium production, although quantum mechanically possible, is far slower than the rate of the hydronium giving away its proton. If the hydronium is only an intermediate, one must show the proton acceptor has a $pK_a$ greater than that of the source of the proton. None of these were shown, not even a slightest indication. However, it was shown that the $pK_a$ of the carboxylic acid of Asp85 decreases by a half unit after the retinal isomerizes to 13-*cis*, in the opposite direction of the wishful thinking (Balashov et al., 1996).

*Reviewer 3: -Line 573: "Neither a hydraulic powered noria or an animal powered sakia in the old time, nor an electric powered fluid pump in the modern days, works under such an operating principle that requires energy for a reconstruction of the landscape." Please, explain what do you mean by "requires energy for a reconstruction of the landscape".*

To lead water from one place to another, people have to dig a canal with a downward slope. This construction requires energy. To move protons from one chemical group to another, proton affinities along the pathway must be changed into an ascending order. These changes require energy. Proton affinities of various chemical groups in a protein form a landscape on which protons are conducted. A directional proton conductance requires an ascending order of proton affinities. If this is not a given, energy is required to make one. This is the basic premise of the mainstream view on how the directional proton conductance in bR is achieved – affinity driven.

*Reviewer 3: -Line 576: "Quite the contrary, the landscape is a given constant and energy is absorbed by the to-be-pumped substance to gain its potential energy, such as water is lifted by a noria or pressurized by an electric pump. The energized substance will then flow freely along the largely constant landscape governed by thermodynamics. In the meanwhile, it is critical to prevent a backflow along a far steeper gradient than the forward-driving gradient." I am lost. I do not know that the author is taking about. Please, refocus it.*
*Reviewer 3: -Line 585: "Alteration of affinities to the pumped substance cannot be an effective strategy because it is against the chemical nature along the translocation pathway and therefore energetically difficult to achieve." Confusing sentence. Please, rewrite it.*

However, no successful pump of any kind adopts the strategy imagined by the mainstream thinking. Because it is energetically too costly to reconstruct the proton affinity landscape against the chemical nature. Proton affinities are considered as a constant, given landscape even if some small changes have been detected. A sufficient yet small dose of energy from the absorbed photon has to be pinpointed to one proton to be pumped just like other pumps for water. Energy is not only focused spatially but also temporally to achieve a high peak power at the beginning of the photoisomerization. Once the to-be-pumped substance, proton or water, is energized. It would flow downhill freely. Therefore, it is critically important to prevent backflow – an issue that the mainstream didn't even care.

*Reviewer 3: -Line 197: "The fundamental flaw in the previous line of thinking is multiple steps of equilibrium driven by proton affinities as the values of pKa describe." Not clear why it is a flawed assumption. Proton transfers are rather fast events, so for transitions between intermediates in the microsecond (K-to L, L-to-M, M-to-N, N-to-O and O-to-bR) it makes perfect sense to assume an equilibration of the protonation state governed by the pKa.*

If p$K_a$ is indeed the driving force of the proton conductance in bR, why not demonstrate an ascending order of p$K_a$ values along the proton pathway? Because nobody can. Because the proton conductance is not governed by p$K_a$ values, at least in two critical steps (Steps 1 and 3 in Fig. 1). First, the deprotonation from the SB is a photophysical process that does not depend on any p$K_a$ values. Second, the overflow through Arg82 in the proton barrier is driven by concentration gradient, not by p$K_a$ values. Needless to say, other downhill steps are indeed governed by p$K_a$ values.

*Reviewer 3: -Line 174: "The difficulty encountered by this hypothesis is the negative pKa values of a hydronium ion and a protonated threonine that the proton has to go through." I agree that it intuitively it does not seem plausible, but science should not be based on just intuitions. Bondar and co-workers have several papers showing that such proton transfer from a CP SB to Asp85 has an energy barrier low enough to be possible (check her review: 10.1007/s00232-010-9329-3).*

The energy barrier could be low. Even without any barrier, this proton transfer is impossible. Because it is an uphill transfer, uphill by 11 orders of magnitude. Someone needs to show this somehow becomes a downhill. I agree completely that science is not intuition. I give three examples in this study. First, the only deep-violet-absorbing M state does not mean it is the only deprotonated state. Second, the proton is more of an integral part of the water rather than an integral part of the protonated SB. Third, the proton barrier surrounding the inner EC channel is not to prevent protons from flowing out. It is to block protons flowing in. Intuition does not work in these cases. However, if you change a perspective, all of them make sense.

*Reviewer 3: Ren introduces several ground-breaking ideas: 1) very fast (femtoseconds) deprotonation of the retinal SB, instead of taking place in tens of microseconds. 2) The metastable proton acceptor of the SB is not Asp85 but a water molecule. 3) The photocycle does not pump protons in its first round, but forms a super-acid initial state. When this super-acidic initial state absorbs light, then the photocycle pumps protons.*

This is an incomplete summary of the findings in this study. I need to clarify 2) and 3). 2) It is OK to call the water molecule a proton acceptor. What is really accepting the protons is a water cluster of at least four waters 400-403 and two carboxylates of Asp85 and 212 (Fig. 5c). Therefore, the full capacity of the proton acceptors is six protons. What is unclear is to what extent the capacity is used during active pumping. Notice that the traditional answer to this question is one. 3) I define a pumping event as the translocation of a proton from the cytoplasmic half channel into the extracellular half channel. Therefore, the first photocycle does pump a proton. However, the first proton occupying one of the six proton capacitors, most likely the carboxylate of Aps85 due to its largest p$K_a$ among the six positions, is not capable to pass the proton barrier. Therefore, no proton release can occur until the proton capacitors are charged up by the subsequent photocycles.

*Rewiewer 3: -Line 546: "It appears that Asp85 and 212 are continuously protonated by proton pumping thus neutralized in the inner EC channel." Protonation of Asp212 was only experimentally observed in the photocycle of wt BR at very low pH values and in some mutants (10.1021/bi990873+ and 10.1006/bbrc.2001.4730), in both cases in the the last steps of the photocycle (O to BR transition). Asp212 is, therefore, not continuously protonated.*

Again, it is not known to what extent the proton capacity of the inner EC channel is occupied during active pumping. It is possible that Asp85 is neutralized first and then transfers its proton to Asp212. This sentence is reworded (Line 859-862).

*Reviewer 1: One of the biggest concerns about this work is the precise position of H+ was not observed in the X-ray crystallographic data so that the formation of hydronium ion was not directly observed in the results. This was suggested by only the theory based on Newton's law. Although the author demonstrated that H+ is left out from the RSB due to its ultrafast movement by the same physics as tablecloth trick, the H+ does not weakly interact with RSB like a dining set on a tablecloth but is covalently bound with RSB. Therefore, it can quickly move with RSB as a single object. In order to make the author's model sure, further experimental evidence or quantum chemical calculation is needed.*

It is a factual error to consider that the proton is covalently attached to the SB. The proton does not contribute an electron to the bonding since it doesn't have one. The proton is only attracted to the lone pair of electrons available on the SB nitrogen. On the other hand, the proton does not contribute an electron to its hydrogen bond with the water either. However, the three O-H bonds in the water including the hydrogen bond are equivalent. It is indistinguishable which of the three is a hydrogen bond. In addition, the positive charge of the proton very likely has been delocalized (Swanson and Simons, 2009). Therefore, between two non-covalent associations with the proton, the O-H association is more likely to take a covalent nature. Therefore, both interactions with the proton are comparable in strength. It is instinctive to think the proton is a part of the SB. Quite the contrary, it is more likely a part of the water. Why would the SB rip a proton away from the water when it departs since no electron density was observed that the water remains intact or not? We cannot easily dismiss the existence of a proton or hydrogen along a good hydrogen bonding distance on the basis of no electron density is observed. We have to acknowledge that a proton or hydrogen is located near the midpoint of the observed hydrogen bond. Well, how do I know the SB departs without the proton? The equivalent question should also be asked: How do you know the SB brings the proton with it? Which interaction survives the abrupt tear is determined by the inertia of the proton. This is not a speculation, rather a conclusive deduction based on chemistry, mechanics, and thermodynamics (Fig. 1 inset). I agree that a quantum chemical calculation is needed. My analysis is the foundation of such calculation. The rest will require an independent expert to carry out.

*Reviewer 2: 2) The inertia of the Schiff base proton is mentioned several times. But the mass of the proton is much lighter than of a nitrogen atom. So I would expect that the Schiff base nitrogen has more inertia.*

This is an important argument. The SB nitrogen has much greater inertia. Why would it accelerate if not the proton? I have explained this in the overview above. But fundamentally, covalent bonds are much stronger than coordination bonds and hydrogen bonds. The SB nitrogen has two covalent bonds to carbons, one single, one double. It is an integral part the chromophore. On the other hand, the proton has already become an integral part of the water due to the delocalized charge. There is no reason, that is, no force, why the proton could accelerate with the departing SB.

*Reviewer 3: Considering a typical isotopic effect, it is possible to deduce the N-H frequency for the SB. This frequency is related to the force of the bond holding the atoms N and H together. My rough calculation from the N-H frequency and the reduced mass gives me a force of 600 N/m for the bond, meaning that if the N atom moves 1 pm but the H atom does not move, the H atom will feel a force of 600 pN, enough to get the acceleration it seems to need to keep moving with the N atom. Is it still possible that the SB could deprotonate by a different mechanism or reason after photoexcitation, but the group of Oluvicci have made dynamics simulations of the retinal*

*isomerization in BR and other microbial rhodopsins at the QM level and they have not reported a SB deprotonation following light excitation to my best knowledge.*
*Reviewer 3: Second, as also mentioned, there is no reason to think that the hydrogen bound to the SB N atom will not be able to follow it as it moves during retinal photoisomerization: the N-H bond can provide enough force if moved from its equilibrium distance by few pm.*

I already respond to this in the overview and above. But briefly, a weak association to the proton is not the reason for deprotonation. The real reason is that the proton inertia determines which of the two weak associations will have to break. Why is the non-covalent association $N_\zeta$:H strong enough to tear apart the near covalent bond H-O?

*Reviewer 3: -Lines 211-218: "Here the ultrafast molecular events revealed from the serial crystallographic datasets demonstrate that the photochemical reaction of bR transfers a proton held by the SB to form a strong acid that barely has any ability to hold a proton. However, this proton transfer is not achieved by a chemical equilibrium therefore not governed by pKa's. Instead, before the isomerization occurs, this proton transfer takes place due to the SB displacement within ~30 fs in a single molecular event. The proton's inertia prevents it from accelerating as fast as the SB under the same physics of the tablecloth 218 trick (Q1)." These are bold claims about what happens to the SB proton by just looking at electron densities from X-ray crystallography, virtually blind to hydrogen atoms.*
*Reviewer 3: -Line 359. "This ultrafast U-turn is how the SB gets rid of the proton by a mechanism of the same physics that governs the tablecloth trick (Q1; Fig. S2). First, as I mentioned, protons are not "visible" in X-ray crystallography (unless the resolution is exceptional, which is not the case).*

"Looking at electron densities from X-ray crystallography" is the first step. The second step is SVD and deconvolution of pure intermediate species. The third step is refinement of the pure conformations. The fourth step is calculation and comparison of conformational parameters. It would be a rookie mistake to consider crystallographically determined conformation discardable based on little to no X-ray scattering from a hydrogen or proton (see above).

*Reviewer 1: -Minor points Page 11, Line 340 What is the meaning of "reliable"? Why does it need to be emphasized so?*

As I hypothesized above, if the molecular event of deprotonation from the SB is a chemical equilibrium, the proton affinity of the SB must decrease and that of the carboxylic acid must increase a million-fold simultaneously. Even if these are achievable, there will be a significant fraction of SB remain protonated at the equilibrium, that is, failed proton transfer. A chemical reaction is not reliable. It depends on the forward and backward rates. Ultimately, it depends on the potential energy landscape. The photophysical process of the SB deprotonation due to the proton inertia is a far more reliable molecular event. It doesn't depend on proton affinity of anything. Nevertheless, I also discuss the possibility of reprotonation on the wrong side, a very rapid chemical equilibrium, that would undo the successful deprotonation. This is why the photoisomerization in bR is so fast at ~500 fs, a barrierless relaxation from Frank-Condon point (Gozem et al., 2017).

*Reviewer 1: Page 18, Line 592-593 "Even if these changes in proton affinity are achievable, there will be 10% failed attempts of proton transfers at a prolonged equilibrium" How was the degree of failure (10%) was estimated?*

If two chemical groups share the same proton affinity, both are equally protonated at the equilibrium of proton exchange. If one has ten times the proton affinity of the other, that is, one p$K_a$ is one unit higher than the other. The equilibrium will lean towards one side so that the group with higher proton affinity would be protonated more. But the other group with less proton affinity would remain a smaller fraction still protonated. That is to say, a chemical equilibrium based on proton affinity is not a reliable way to deprotonate. 10% is only an approximation. It can be shown precisely that this is pH dependent. This is reworded (Line 908-910).

*Reviewer 3: -Line 80. "However, the event of SB deprotonation has been mistakenly inferred from the observed protonation of the carboxylate of Asp85." SB deprotontation has been mainly studied by the rise of the retinal absorption at 400 nm in UV- vis spectroscopy, while Asp85 protonation has been studied by the rise of the C=O of its carboxylic group at 1760 cm-1. By far, the most habitual technique used has been visible spectroscopy and, thus, SB deprotonation has been rarely deduced by Asp85 protonation, but the other way around: people doing visible spectroscopy often assume Asp85 protonation when they observe SB deprotonation.*

This is a very constructive point. I have expanded my argument around this point substantially (Line 487-540). It was concluded that M is the only deprotonated state because it is the only state that absorbs in deep violet. This was probably the greatest misinterpretation. The strongest evidence was the absorption properties of model compounds. However, the correlation between an unprotonated SB and near UV to deep violet absorption was never strictly established. Such conclusion that dominated the mainstream thinking for decades was naïve and unsophisticated. As I demonstrate in this study with eight pure intermediate structures, the absorption property of bR is largely governed by the transient evolution of the chromophore going through a complicated relaxation from a very strained conformation. This is particularly true in the early photocycle and repeats again after reprotonation and reisomerization. It is the consecutive and stepwise blue shifts that indicate the unprotonated SB during the relaxation (Fig. B). However, it was not necessary to know the transient conformations of the chromophore throughout the photocycle to cast a doubt on the naïve conclusion. How could the mainstream assume that every intermediate structure has relaxed enough to show its protonation effect in the absorption property? The M absorption in deep violet was simply misinterpreted big time.

*Reviewer 2: However, I see one major inconsistency. If the bacteriorhodopsin photocycle would indeed start with the deprotonation of the Schiff base in retinal, then the spectral properties would drastically change. A deprotonated Schiff base absorbs at 380 nm, instead of 568 nm from protonated Schiff base in bacteriorhodopsin. But ultrafast spectroscopic studies do not show such a spectral shift. Therefore there is no evidence for what is proposed by the author.*

See above. Where a SB absorbs was never solidly established, protonated or unprotonated. There were always exceptions. Environment is a significant factor. Distortion of the retinal is another. It was assumed that every intermediate state is a fully relaxed state. Therefore, the absorption maxima should follow what are known from model compounds. This assumption was far from the reality. A cross validation was that the correlation between absorption maxima and protonation states were artificially assigned based on coincidences that two observations were obtained during the same period of the photocycle, here absorption maxima and vibration modes. The same misassignment could occur to bond stretching frequencies in a strained conformation. Far stronger factors to influence absorption and vibration properties were ignored.

*Reviewer 3: -Line 367: "The U-turn of the SB marks the peak power, or the maximum rate of enthalpy change dH/dt, of the entire photochemical reaction (see Concluding 369 Remarks)." This statement is far from evident. I do not see why.*

Why the absorption spectrum of bR changes during the photocycle? We could tribute that observed spectral change to changes in chromophore and protein conformation, charge distribution, protonation state, etc. All these changes are associated with energy conversion or flow. Therefore, the temporal rate of the spectral change reflects the rate of changes in conformation, protonation, charge distribution, etc., and therefore, a power function of time. The change in absorption maxima clearly shows the peak power of the photocycle occurs at the beginning – a 108-nm blue shift followed by a 165-nm red shift (Fig. B). These two opposite swings take place within the first 500 fs. Spectrograms offer a much more complete view that shows the peak power is reached within the first tens of fs (Kahan et al., 2007; Kobayashi et al., 2001).

*Reviewer 2: 3) Did the author compare the energy of the hydrogen bond with that of the photon that excites bR? The energy of the photon should be much larger than the hydrogen bond. It answers the question about the driving force of the proton pump, but it is not discussed in the manuscript.*

No doubt the absorbed photon energy is more than sufficient to drive the pump. But that doesn't explain anything. The main point in this study is not the total energy but the peak power throughout the photocycle. It is the peak power that drives the pump. Both energy and peak power are discussed throughout the manuscript (Line 442-448, 527-533, 657-659, and 683-691).

*Reviewer 1: Although the author explained the reason for the discrepancy of his model and the previous interpretation of RR study in the 2nd paragraph on page 10 (line 314-330), it is not sufficiently convincing unless further experimental or computational support is shown. Especially, although the position of C=N vibration in the RR spectra correlates with the absorption wavelength of each photo intermediate as the author mentioned, the downshift with*

*deuterium can be explicitly observed so that the assignment of the protonation state of RSB by RR spectroscopy is extremely solid and consistent with FTIR spectroscopy which also observed N-H vibration of RSB not only in the dark state but also in the K and other intermediates. Further experimental or computational results are needed to reconcile the author's model with these vibrational spectroscopy studies.*
*Reviewer 2: In addition, a change in the protonation would also lead to a change which can be detected by vibrational spectroscopy, which is also not described in the literature as far as I know. Therefore it is important that the author will describe how the new mechanism can be verified using time resolved spectroscopy or how it fits with a large body of published spectroscopic results.*

There is solidly evidence to prove an unprotonated SB in M state, which is not being disputed here. Where is the evidence other than the absorption property discussed above to show that the molecular event of deprotonation from the SB synchronizes with the M formation? An unprotonated SB in M state is often mixed with SB deprotonation at M formation. The former does not imply the latter. Quite the opposite, the down shift of C=N stretching frequency upon deuteration was found very small in K state, indicating that the SB has lost its hydrogen bond before K (Kandori et al., 2002). How does proton transfer occur without the hydrogen bond? It was noticed very early on that the SB deprotonates before M formation (Lewis et al., 1974; Marcus and Lewis, 1977). However, in order to reconcile with the fact that M is the only state featuring deep-violet absorption and its predecessor L absorbs in green, they had to stretch the conclusion. See below for N-H frequency.

*Reviewer 3: -Lines 276-297. The authors provide some arguments that "proof" that the SB deprotonates few fs after light absorption, but these arguments are very speculative. The N-H vibrational frequency of the SB in the ground state of BR has been determined by the group of Kandori in D2O (corresponding to the N-D vibration, 10.1021/bi025585j).*
*Reviewer 3: -Lines 314-330. The author suggests that the believe that the SB remains protonated until the L to M transition is only (mostly?) based on Resonance Raman results. RR has used the sensitivity of the C=N frequency to deuteration as a proxy for both the H-bonding and protonation state of the SB. The problem is that both a H-bonding free and a deprotonated SB are H/D insensitive, meaning that it cannot distinguish a protonated SB losing its H-bond and a deprotonated SB. But the author seems to ignore that the N-H frequency of the SB has*

*been study by FTIR spectroscopy by the group of Kandori. This vibration can discriminate between changes in H-bonding (the band shifts) and deprotonation (the band disappears). In the K intermediate the authors observed that the N-H frequency upshifts, meaning that in the K intermediate the SB is protonated and is less H-bonded than in the initial state (10.1021/bi025585j).*

I agree with the reviewer on the ambiguity of RR results. More of the point, the RR results did show that the SB is unprotonated in M. But they have difficulty to pin down that the SB retains its proton before M. The "observed" vibration frequencies at 2468 and 2495 $cm^{-1}$ of the N-D bond along the membrane plane in K were a misassignment (Kandori et al., 2002). This result is self-defeating because the orientation of the bond already casts a doubt on the frequencies of the bond. There is no such bond N-D or N-H in K. Even if there is, that bond would be more perpendicular to the membrane than parallel. Furthermore, the same work found that the SB is no longer hydrogen bonded with a water in K. If that hydrogen bond is already broken in K, how does the proton transfer later? Quite the opposite, the proton has already departed before K. Therefore, there is no N-D or N-H bond in K. The small C=N frequency down shift in K when deuterated indicates not only the water has been lost but also the proton or deuteron.

*Reviewer 3: In addition, the intensity of retinal C-C vibrations in FTIR experiments it is known to be very sensitive to the protonation state of the SB. By this means FTIR experiments can follow changes in the protonation state of the SB, which seems to deprotonate in the L-to-M transition, and not earlier (10.1021/jp201739w).*

The work of Lorenz and co basically shows that some subjectively "selected IR bands" would follow the known kinetics of bR photocycle (Lórenz-Fonfría et al., 2011). Sure. I am not commenting whether such subjective selection has any significance. There are far more statistically sound methods to deal with such problem. But why does this prove deprotonation from the SB at M formation? The only resort falls back to the absorption in deep violet in M.

*Reviewer 3: Lastly, as already mentioned, the retinal absorption in the visible is notably sensitive to the protonation state of the SB, with results totally consistent that the M intermediate is the only intermediate state (at least the only one significantly populated) with a deprotonated SB.*

Yes, the last resort is the deep-violet-absorbing M state already discussed above.

*Reviewer 3: -Line 341: "The deprotonation event [from the SB] cannot be directly inferred from the vibrational signatures due to the protonation of the carboxylic proton acceptors in the Asp residues." Why not? As mentioned above there are several spectroscopic signatures (N-H vibration, C-C intensities, lamda max of the retinal in the UV-vis) that inform about the protonation state of the retinal SB.*

Ultimately, there is only one – the deep-violet-absorbing M state. Everything else only synchronizes with M formation instead of indicates the deprotonation event. And the deep violet absorption at 412 nm was misinterpreted (see above).

*Reviewer 3: Also, if the SB was to deprotonate after photoexcitation, the electron density of the retinal would rapidly adjust to the new deprotonated state and the I intermediate would be largely blue shifted, as the M intermediate is.*

I intermediate is the second largest blue-shifted state. It is very strained so that the full effect of deprotonation has not shown yet (Fig. B). But the 108-nm blue shift is a good indication for deprotonation. I wonder why this was not considered before. I species was not known until a very late time, too late to change the story in the mainstream. Also, it was unfortunate that I species absorbs in blue unlike M that absorbs in deep violet.

*Reviewer 1: The author demonstrated that a H+ is pooled in the extracellular part of RSB during the photocycle initiated by the first photon, and further supplements of H+ during subsequent photocycle(s) are needed for the efficient H+ release to the extracellular solvent. However, if it is the case, because the absorption of the retinal chromophore is highly sensitive to the electric charge around RSB (e.g. Melaccio et al., J. Chem. Theory. Comput. (2016)), a new state exhibiting an absorption significantly shifted, probably toward the red side, from the initial state must be observed at the end of the first photocycle by the transient absorption spectroscopy. But, to my knowledge, no supportive result was reported so far.*

That is probably why the ground state is yellow absorbing at 568 nm and O is the most red-absorbing state at 640 nm. A protonated SB in a model compound absorbs as far as green light < 520 nm. The ground state absorbs yellow light probably because the super-acidic inner EC channel is already established after light adaptation. The protonated SB is back into the super-acidic pool with an additional proton in O state. Therefore, the most red-absorbing O state corresponds to the most acidic inner EC channel throughout the photocycle. By the way, I never found an explanation in the literature on the most red-absorbing O state.

*Reviewer 1: Page 17, Line 549-550 "The observed kinetics of FTIR continua could explain excess protons harbored by the water cluster in the inner EC channel." The FTIR spectroscopy observed continuum band for the "initial" state (assigned to the protonated water cluster between Glu194 and Glu 204) not for any photo intermediate. Hence, it cannot support the hypothesis of the author that hydronium ion is generated in the EC cavity during the first photocycle.*

The so-called initial state could be understood as an established proton pool after light adaptation. This is the all-*trans* super charged ground state in light (Fig. 5c). The negative going FTIR continua indicate the discharge from this proton pool. It is unknown how many excess protons are sufficient to drive the proton conductance. It could be as few as three including the original protons on the SB and the guanidinium ion, that is, one extra. It could be as many as eight or six extras. Nevertheless, this explains the complexity of the kinetics of the continuum signals. Originally, the signal was thought coming from within the active site near the SB (Wang and El-Sayed, 2001). Later, the signal was misassigned to the proton release complex because the mutant E204Q affected the observed signal. That does not mean that the assignment is a fact. The assignment is without structural basis. There is no proton barrier around the proton release complex to establish a water cluster with more than one excess proton. Quite the contrary, such proton barrier surrounding the inner EC channel is well-known.

*Reviewer 3: -Lines 299-312: The author quotes references to FTIR studies describing broad bands assigned to protonated water clusters in support this idea that the SB proton is early accepted by a water molecule. However, the author does not mention that the reported continua bands in BR are in all cases negative, i.e., protonated water clusters present in the initial BR state transiently deprotonate during the photocycle. Also, continua band in BR have been only reported to occur in the L to M transition and N-to-O transition, i.e., in the microseconds-milliseconds.*

This is an excellent point. The negative going FTIR continua in μs-ms indicate discharge from the proton pool maintained by consecutive pumping. The timing agrees perfectly.

*Reviewer 3: -Line 548: "Several hydronium ions make the inner EC channel positively charged. The observed kinetics of FTIR continua could explain excess protons harbored by the water cluster in the inner EC channel (Garczarek et al., 2004, 2005; Lorenz-Fonfria et al., 2017; Wang and El-Sayed, 2001)." As I already noted, these continua are all negative and not positive in light-induced FTIR difference spectroscopy, meaning that they cannot explain the building of hydronium ions during the photocycle. They indicate that a protonated water clusters are transient deprotonated and final reformated during the photocycle.*

Yes, the continua are not due to the building up of charges in the water cluster. They depict the transient discharge of the proton pool. The water cluster is constantly super charged as soon as light adapted. Only change that could be detected is the periodic discharge.

*Reviewer 1: Also, the H+ release to the solvent was observed in a single-photon photocycle using pH indicator (pyranine) (Varo et al. Biochemistry (1993)). This also cannot support the presence of the super-acidic inner EC channel.*

A single photon photocycle could certainly release a proton. A first photon photocycle after a long dark period is not capable of proton release. Because the first proton left in the inner EC channel cannot pass the proton barrier (Fig. 1). The inner EC channel must be saturated with many protons before proton release. An easier way to understand this is that protons are squeezed out by continuous and forceful pumping. This is why the photophysical nature of the SB deprotonation is important. This is something a chemical equilibrium based on p$K_a$'s cannot achieve.

*Reviewer 3: -Line 516: "The first a few photocycles build up a proton concentration at the end of the EC half channel without any proton release." How the author knows about this from the analysis of the time-resolved X-ray data? I can say that I have been single shot experiments on BR where proton release could be detected by pH sensitive dyes. Moreover, the signal from the pH sensitive dye remains the same with every photoexcitation: every photocycle releases the same number of protons. This is the reason the data is averaged, so published data is the average from multiple measurements. But, as I say, proton release takes also place in single short experiments.*

I love this question: "How [does] the author know[s] about this from the analysis of the time-resolved X-ray data?" This is the power of seeing. The super-acidic inner EC channel could only be established by the SB deprotonation mechanism proposed here. The mainstream thinking based on proton affinity could never realize a proton concentration high enough to conduct a proton through the proton barrier. Single photon photocycle is different from first photon photocycle. First photon photocycle is a single photon photocycle after dark adaptation. One bR molecule absorbs a single photon and enters a photocycle. If this event occurs after a long dark period, this is the first photocycle. The first photocycle is a single photon photocycle. But a single photon photocycle is not necessarily the first photocycle. It could be the second, the third, … The entire manuscript is discussing single photon photocycle. I have commented on multiphoton photocycles. They do not carry the physiological function (Ren, 2022).

*Reviewer 3: -Line 527: "Although the first a few photocycles are productive, they are not capable of releasing any proton before an established proton concentration gradient" Wrong. Proton release in single short experiments can be measured, as already mentioned. The first photocycle is already able to release a proton.*

If the single shot experiment starts from a light-adapted state, every photocycle would release proton. The buildup of the proton concentration during the startup is a part of the light adaptation. Evidence on this point has been substantially expanded (Line 781-825).

*Reviewer 3: Line 190: "Q3) How are protons conducted > 15 Å through the rollercoasting pKa values and released into the EC medium from the SB of a high pKa through aspartic acids of low*

*pKa, arginine (strictly speaking, its guanidinium ion) and tyrosines of high pKa, and glutamic acids of low pKa again?" The released proton does not originate from the SB, but from the proton release complex (PRC). This is well-established, as the author can confirm reading any review. What it is less clear is how the PRC recovers its proton from Asp85 in the O to bR transition, but simulation studies from the group of Elstner and co-workers have addressed this question (10.1021/ja809767v).*
*Reviewer 3: -Line 501 to 506: "Yet, the question remains: How are protons conducted through the proton barrier of the guanidinium group of Arg82 that is already protonated and the phenol hydroxyl groups of the Tyr residues that cannot be protonated anymore (Brown et al., 1993)? No evidence shows that such rollercoasting pKa values would change in any meaningful way by the light-induced conformations. Therefore, the previous proposal on an affinity-driven proton conductance is fundamentally flawed." The protons are not conducted through this proton barrier, because the analyzed data reaches only until the M2 intermediate. As the author probably knows, the proton released to the EC medium does not come from the SB or Asp85, but from a different group known as the proton release complex (PRC).*

The released proton ultimately originates from the SB. Because the SB brings the proton from the CP side to the EC side, that is, the pump. The proton would have to pass the proton barrier. It would require multiple protons accumulated in the inner EC channel to pass the proton barrier.

*Reviewer 3: For the reprotonation of the PRC from Asp85, which take place much later in the O-to-BR transition, the proton does need to go through Arg82 and so on. But at this point some structural changes have take place that make it possible. The author can check publications from the group of Eltsner addressing this late proton transfer (for instance 10.1021/ja809767v).*

This is another wishful thinking without any experimental observation just like those imaginations in the early days. The last two intermediate states N and O are not going to have dramatic conformational changes. Because I already see the trend approaching the ground state (Figs. 4 and S16). The rest are going to return to the ground state as well. There will be no change to the proton barrier. Its function is to maintain as a good barrier. The protons will have to pass this barrier. The way they could pass is first to accumulate in the inner EC channel.

*Reviewer 3: -Line 508: "Proton conductance after SB deprotonation through the EC half channel not necessarily operates on the basis of one event of proton release per photocycle." I am quite surprised with this statement. It is true that the proton-pump stoichiometry was for a time in the past an open debate, but work on the late eighties (mainly from Dencher but also by other) settled down the current view of one proton pumped per deprotonated SB (strictly, one proton released to the medium for each BR reaching an M intermediate).*
*Reviewer 3: -Line 510: "It was previously suggested that a proton pumped in one photocycle could be released in the next or subsequent photocycles (Braiman et al., 1991)." Initial FTIR results from Asp96 carboxylic C=O bands lead to Braiman and co-workers to consider the possibility of a partial or total deprotonation of Asp96 in the L intermediate. This was latter shown to be an artefact caused by overlapping contributions from Asp96 and Asp115 in the carboxylic region (Gerwert and co-workers, and Maeda and co-workers).*
*Reviewer 3: -Line 511: "However, a flawed common assumption persists that the pumping and releasing of each proton are somehow scheduled in different periods of a photocycle." This is not an assumption, but an empirical fact. The use of pH sensitive shows that a proton is released first in the EC domain and a proton is taken later in the CP domain, with the uptake completing the proton pumping (see for instance a review by Heberle: 10.1016/s0005-2728(00)00064-5).*

One proton release per photocycle is only true when the necessary concentration gradient is established in the light-adapted state. During the startup period of the first a few photocycles, no proton can be released before the inner EC channel becomes super-acidic. To avoid confusion, this is reworded (Line 781-825).

*Reviewer 3: -Line 521: "That is to say, two states sharing the same or very similar all-trans retinal are differentiated – the resting state in dark (Fig. 5b) and the beginning state of the photocycle in light with a super charged inner EC channel (Fig. 5c). The structure of the all-trans light state is not observed directly. Its existence can only be implied from the spectroscopic data." The spectroscopic data refutes the above claims. If you excite BR with a laser pulse in tens of microseconds the protein returns to the same initial state. How do I know this hypothetical super charged initial state does not exist? If the final state after a single laser pulse was a super charged state as the one shown in Fig.5 it would be easily detected in an FTIR difference spectrum. However, after a single laser excitation the light-induced FTIR difference spectrum goes back to zero, which indicates that the initial and final state are the same.*

This is further clarified (Line 818-825). Vibrational spectroscopy has always measured the ground state as the super charged state after light adaptation. After a photocycle, everything goes back to the super-charged ground state at the beginning. The discharged resting state in dark is harder to observe.

*Reviewer 3: -Line 530: "Proton release from the wildtype bR was observed before proton uptake as if it is coupled with, if not faster than, the protonation of the proton acceptor Asp85 (Braiman et al., 1988, 1991; Gerwert et al., 1990; Zimanyi et al., 1992), which strongly suggests that the newly pumped proton is not the one released to the EC medium in the same photocycle." I am not sure if I understood that the author wants to say here. It is well known that in BR a proton is released first to the EC side and later a proton is uptaken from the CP side, completing by his way proton pump. The fact that proton release precedes proton uptake is not a problem, and it is perfectly explained by the order of the proton transfers inside BR. It also seems that the author states here that before Asp85 protonates, a proton is already released by BR. A recent comparison of protonation of Asp85 and the release of protons show that the former takes place slightly before the second, not after (10.1073/pnas.1707993114).*

There is no disagreement here. The proton released is not the proton transferred into the EC half channel by the SB and accepted by Asp85 in the same photocycle. Rather it is the proton transferred by the SB and accepted by Aps85 several photocycles ago. As Reviewer 3 said that it "is not a problem". It supports my theory perfectly.

*Reviewer 3: -Line 82. "It has been widely quoted in the literature that the proton transfer from the SB to the proton accepter occurs as the M formation." Independent studies by both Gewert and coworkers (10.1016/S0006-3495(92)81722-8; 10.1016/S0006-3495(93)81264-5) and Lorenz-Fonfria and co-workers (10.1021/jp201739w) have shown that both the rise of the absorbance at 400 nm and at 1760 cm-1 follow roughly the same kinetics. Mutagenesis studies also point to Asp85 as the proton acceptor. So, there are good reasons to think that a proton is transferred from the SB to Asp85 in the M intermediate.*

There is no dispute on the synchronization of the rise of M indicated by 412-nm absorption and the formation of -COOH indicated by 1760 $cm^{-1}$ vibration frequency. That observation simply says the carboxylate is neutralized in M state. It doesn't say where the proton is transferred from.

*Reviewer 3: -Line 188: "Q2) Which molecular event, the isomerization, or another event, switches the accessibility of the SB from EC to CP and how does the SB avoid reprotonation on the wrong side?" Several works from Varo & Lanyi (for instance 10.1021/bi00234a024), and other studies, point to the irreversible M1 to M2 transition as the switching event. Results from Balashov (reviewed in 10.1016/s0005-2728(00)00131-6) show that the pKa of Asp85 is coupled to the protonation state of the proton release complex. Lorenz-Fonfria & Kandori (10.1021/ja900334c) showed that the proton release complex deprotonates in the M1-to-M2 transition. All these observations indicate that deprotonation of the PRC increases the pKa of Asp85 and, in combination with structural chanages, forces the SB to recover its proton from a CP group.*

No one disputes these findings: the irreversible $M_1 \rightarrow M_2$ transition, deprotonation from PRC during $M_1 \rightarrow M_2$ transition, and the p$K_a$ of Asp85 coupled to the protonation state of PRC. However, none of these answers the question of the accessibility switch of the SB. None of these is even relevant to the question. The controversy originates from what structural basis is referenced. These spectroscopic studies mentioned by Reviewer 3 interpreted their data based on a bR structure similar to its ground state, which is far from the reality. During $M_1 \rightarrow M_2$ transition, the SB is to be protonated in 13-*cis* configuration pointing to the CP side (Figs. 4, S14, and S15). It has no access to anything in the EC half channel. Asp85 and PRC on the EC side have nothing to do with where the SB could capture its next proton. At this point, the SB has no choice but to gain a proton from the CP side. If this is successful, this photocycle is productive. Otherwise, this is a futile photocycle. The question on the accessibility of the SB is how it could avoid a reprotonation while it is still in the EC half channel before isomerization.

*Reviewer 1: Page 12, Line 389 The J-to-K conversion occurs at 3-5 ps (e.g. Nuss et al. Chem. Phys. Lett. (1985)). Therefore, The J' and J in this article should be assigned to the J and K, respectively.*

It is a good point. As I clarified further, the assignment of the intermediate states to my refined structures are solely based on the transition time. There are no absorption data associated with the X-ray data. Therefore, misassignment is possible. I have warned the readers explicitly (Line 303-307). However, the long lifetime of K is very certain.

The contraction of the retinal binding pocket captured in J cannot last that long because it is simply a recoil from the previous expansion. Therefore, I stand by my assignment so far.

*Reviewer 3: -Line 240. "Eight intermediate structures along the photocycle, now isolated from one another, are refined against reconstituted structure factor amplitudes (Methods; Table S2)." For the long delays data, 6 significant SVD components are identified. However, Table S2 shows that intermediates from K to M2 are reconstructed with only four components: 1, 2, 3 and 6. What happens with components 4 and 5? In addition, looking at Fig. S4a it is not clear why only 6 components are selected. Components 6 and 7 (and the following) have almost the same contribution in describing the data set.*
*Reviewer 3: Line 450: "Four of them contain outstanding signals of structural changes (Fig. S5). Two others also carry structural signals but without a clear time dependency (Fig. S6)." The time dependency is mentioned but not shown in any case. I would be nice to see the time dependence of all the 6 components, the 4 used to reconstruct the pure intermediates and the 2 with structural detail but discarded.*

An additional figure shows the lack of time dependency of components 4 and 5 (Fig. S7). Component 4 fluctuates a lot although it seems to carry signals over the EC half of the molecule (Fig. S8a). Component 5 is only required by two datasets at 40 ns and 2 μs although it may carry signals on the CP segments of helices E and F (Fig. S8b). These signals could reflect some systematic differences from one experiment to another. Due to the lack of consistency, these components are not used in structural refinement. This is a good example to demonstrate how SVD identify and isolate inconsistent systematic errors.

*Reviewer 3: It is also surprising that for the early delay, which should include the K intermediate and thus 5 intermediates, only 3 significant components are found (rank of 3). This would rather suggest that there are only three structural intermediates in this time window: I, J and K. In other words, which is the evidence or reason to include I' and J' as intermediates? They do not seem to be needed to describe the data when considering that SVD indicates a rank of 3. If the five intermediates co-exist in this time window, should not the rank found by SVD be equal to 5?*

It is a good point that the K intermediate could be mixed in the early delays. The difficulty is the very few early delays available. All short delays from Novacs et al. are useless except complicated oscillations (Ren, 2022). Luckily, the last short delay is 10 ps, less than a decade after K supposed to appear. Therefore, K should not be involved significantly. If five intermediates coexist, should the outstanding rank be 5? Yes, if these five intermediates are truly independent of one another, that is, the dot product of any pair of the electron density maps among these five is zero. This is not guaranteed. In fact, the consecutive intermediates could be very correlated. For example, I and J are found in an expanded and contracted retinal binding pocket, respectively, while the pocket in I′ and J′ are normal. Therefore, I′ and J′ are necessary. It only requires one component to describe the retinal binding pocket for all four states since expansion and contraction are depicted as positive and negative component of $\boldsymbol{U}_{10}$. A normal pocket is depicted as $0\boldsymbol{U}_{10}$ (Ren, 2022).

*Reviewer 3: There are more issues about the application of SVD to the early delay data. SVD assumes that the data can be decomposed in three matrices, as explained by the author. For the present application this assumption translates into assuming that the experimental data can be described as the linear combination of electron densities of a discrete number of intermediates. While this is true at longer time scales, where the energy barriers force the system to populate only a discrete number of intermediate states, in the very fast regime the structural changes are ballistic and show coherence effects: the system does not necessarily experience a transition between intermediates but show a continuous change, and the notion of discrete intermediate might lose sense and, thus, the applicability of SVD.*

In a very fast time regime, if we focus on the double bond isomerization, the conformational change is very discrete *trans* and *cis*. This is the key difference that this work stands out from the previous interpretations of intermediate conformations (Fig. A). However, if we consider the expansion and contraction of the retinal binding pocket, the changes are likely to follow a ballistic trajectory without any discrete states. We don't really have the luxury to study such a ballistic trajectory in great detail. We only have a handful of time points. For the sake of argument, if we indeed have many time points along a ballistic trajectory, would SVD still be effective? Reviewer 3 is right that SVD becomes less effective compared to its application in discrete states. However, it depends on how extensive the ballistic trajectory is. If the trajectory is as short as the retinal binding pocket expansion, SVD is still very effective. The greatest difficulty is an extensive trajectory with continuously populated conformations. In this case, the dimensionality reduction becomes less effective.

*Reviewer 3: This regime can be identified by the presence of coherent oscillations in the positions of atoms. I guess these oscillations are present in the data, right?*

Exactly. I have demonstrated in two separate cases that oscillatory signals could be extensive (Ren, 2019, 2022). However, I have never found these signals correlated with any function and secondary structure of proteins. These oscillatory signals take a lot of components to describe. Therefore, I conclude that these oscillations are due to multiphoton absorption and the decays from physiologically irrelevant potential energy surfaces (Miller et al., 2020; Ren, 2022).

*Reviewer 3: -Lines 588 to 612: The rest of the Concluding Remarks are confusing and hard to follow divagations about the view of the author about how BR works.*

What we can learn from this molecular proton pump is far beyond bR. It is necessary to discuss at a different level.

*Reviewer 3: -Line 53. "This integral membrane protein of 28 kDa singlehandedly achieves the exact same goal of photosynthesis and cellular respiration combined." Photosynthesis uses light to create a proton electrochemical gradient, but also to transfer electrons from an initial donor (H2O or other inorganic molecule) to a final acceptor (NADP+) against their redox potential, and cellular respiration uses electron transfer from an organic donor (NADH or other) to an inorganic acceptor (O2 or other) in favor of their redox potential to build a proton electrochemical gradient. These two processes combined do not achieve the same goal as light-driven proton pump by BR. Please, rewrite.*

This long sentence of Reviewer 3 "Photosynthesis uses light to … to build a proton electrochemical gradient" already proves that my statement is correct. Everything in the mid-sentence is means, not goal. Higher organisms had to develop more sophisticated means, but the goal remains the same, to acquire energy from the sun.

*Reviewer 3: -Line 181: "This interpretation of these highly twisted double bonds is a direct consequence of the inability to unscramble a mixed observation of multiple conformations." Without further explanation this sounds as an opinion dressed as a fact. Being an opinion, it should better read as: "This interpretation of these highly twisted double bonds could be a direct consequence of the inability to unscramble a mixed observation of multiple conformations."*

Further explanation is expanded (Line 168-192).

*Reviewer 1: Page 14, Line 455-456 "the transition times between intermediates K, L, M1, and M2 known from spectroscopy" Please add the relevant references.*

Done.

*Reviewer 1: Figure 1 The N and O intermediates have 13-cis and all-trans retinal, respectively.*

Fixed.

*Reviewer 3: -Line 68 "It has been shown that the species I or a collection of species prior to the photoisomerization arises before 30 fs and remains in 13-trans instead of a near 90° configuration about C13=C14 (Zhong et al., 1996)." Why a collection of species? In the cited work the authors determined a single rising component at 18±10 fs.*

This is clarified (Line 71-77).

*Reviewer 3: -Line 56: "A trimeric form of bR on the native purple membrane shares the retinal chromophore and the same protein fold of seven transmembrane helices A-G (Fig. S1) with large families of microbial and animal rhodopsins (Ernst et al., 2014; Kandori, 2015)". This statement might suggest that animal rhodopins and other microbial rhodopsins also adopt a trimeric form, which is not the general case. Please, rewrite.*

Done (Line 56-59).

*Reviewer 1: Page 15, Line 498 "83" would be better to be "Tyr83".*

Done.

*Reviewer 3: -Line 169: "Therefore, the SB proton is associated with both N and O simultaneously in Nz:H+:OH2 with both interactions much weaker than an ordinary single bond (Fig. 2c)." Add "atoms" after N and O to avoid confusion with the N and O intermediates.*

Done.

#### References

Andersen, L.H., Nielsen, I.B., Kristensen, M.B., El Ghazaly, M.O.A., Haacke, S., Nielsen, M.B., and Petersen, M.Å. (2005). Absorption of Schiff-base retinal chromophores in vacuo. J. Am. Chem. Soc. *127*, 12347–12350. https://doi.org/10.1021/ja051638j.

Balashov, S.P., Imasheva, E.S., Govindjee, R., Sheves, M., and Ebrey, T.G. (1996). Evidence that aspartate-85 has a higher pKa in all-trans than in 13-cis bacteriorhodopsin. Biophys. J. *71*, 1973–1984. https://doi.org/10.1016/S0006-3495(96)79395-5.

Gozem, S., Luk, H.L., Schapiro, I., and Olivucci, M. (2017). Theory and simulation of the ultrafast double-bond isomerization of biological chromophores. Chem. Rev. *117*, 13502–13565. https://doi.org/10.1021/acs.chemrev.7b00177.

Grandbois, M. (1999). How Strong Is a Covalent Bond? Science *283*, 1727–1730. https://doi.org/10.1126/science.283.5408.1727.

Heyde, M.E., Gill, D., Kilponen, R.G., and Rimai, L. (1971). Raman spectra of Schiff bases of retinal (models of visual photoreceptors). J. Am. Chem. Soc. *93*, 6776–6780. https://doi.org/10.1021/ja00754a012.

Jardetzky, O. (1966). Simple allosteric model for membrane pumps. Nature *211*, 969–970. https://doi.org/10.1038/211969a0.

Kahan, A., Nahmias, O., Friedman, N., Sheves, M., and Ruhman, S. (2007). Following photoinduced dynamics in bacteriorhodopsin with 7-fs impulsive vibrational spectroscopy. J. Am. Chem. Soc. *129*, 537–546. https://doi.org/10.1021/ja064910d.

Kandori, H., Belenky, M., and Herzfeld, J. (2002). Vibrational frequency and dipolar orientation of the protonated Schiff base in bacteriorhodopsin before and after photoisomerization. Biochemistry *41*, 6026–6031. https://doi.org/10.1021/bi025585j.

Kobayashi, T., Saito, T., and Ohtani, H. (2001). Real-time spectroscopy of transition states in bacteriorhodopsin during retinal isomerization. Nature *414*, 531–534. https://doi.org/10.1038/35107042.

Kovacs, G.N., Colletier, J.-P., Grünbein, M.L., Yang, Y., Stensitzki, T., Batyuk, A., Carbajo, S., Doak, R.B., Ehrenberg, D., Foucar, L., et al. (2019). Three-dimensional view of ultrafast dynamics in photoexcited bacteriorhodopsin. Nat. Commun. *10*, 3177. https://doi.org/10.1038/s41467-019-10758-0.

Lanyi, J.K., and Schobert, B. (2007). Structural changes in the L photointermediate of bacteriorhodopsin. J. Mol. Biol. *365*, 1379–1392. https://doi.org/10.1016/j.jmb.2006.11.016.

Lewis, A., Spoonhower, J., Bogomolni, R.A., Lozier, R.H., and Stoeckenius, W. (1974). Tunable laser resonance Raman spectroscopy of bacteriorhodopsin. Proc. Natl. Acad. Sci. *71*, 4462–4466. https://doi.org/10.1073/pnas.71.11.4462.

Lórenz-Fonfría, V.A., Kandori, H., and Padrós, E. (2011). Probing specific molecular processes and intermediates by time-resolved Fourier transform infrared spectroscopy: Application to the bacteriorhodopsin photocycle. J. Phys. Chem. B *115*, 7972–7985. https://doi.org/10.1021/jp201739w.

Luecke, H., Schobert, B., Richter, H.-T., Cartailler, J.-P., and Lanyi, J.K. (1999). Structure of bacteriorhodopsin at 1.55 Å resolution. J. Mol. Biol. *291*, 899–911. https://doi.org/10.1006/jmbi.1999.3027.

Marcus, M.A., and Lewis, A. (1977). Kinetic resonance Raman spectroscopy: Dynamics of deprotonation of the Schiff base of bacteriorhodopsin. Science *195*, 1328–1330. https://doi.org/10.1126/science.841330.

Mendelsohn, R. (1973). Resonance Raman spectroscopy of the photoreceptor-like pigment of Halobacterium halobium. Nature *243*, 22–24. https://doi.org/10.1038/243022a0.

Mendelsohn, R. (1976). Thermal denaturation and photochemistry of bacteriorhodopsin from Halobacterium cutirubrum as monitored by resonance raman spectroscopy. Biochim. Biophys. Acta BBA - Protein Struct. *427*, 295–301. https://doi.org/10.1016/0005-2795(76)90305-6.

Miller, R.J.D., Paré-Labrosse, O., Sarracini, A., and Besaw, J.E. (2020). Three-dimensional view of ultrafast dynamics in photoexcited bacteriorhodopsin in the multiphoton regime and biological relevance. Nat. Commun. *11*, 1240. https://doi.org/10.1038/s41467-020-14971-0.

Nango, E., Royant, A., Kubo, M., Nakane, T., Wickstrand, C., Kimura, T., Tanaka, T., Tono, K., Song, C., Tanaka, R., et al. (2016). A three-dimensional movie of structural changes in bacteriorhodopsin. Science *354*, 1552–1557. https://doi.org/10.1126/science.aah3497.

Nogly, P., Weinert, T., James, D., Carbajo, S., Ozerov, D., Furrer, A., Gashi, D., Borin, V., Skopintsev, P., Jaeger, K., et al. (2018). Retinal isomerization in bacteriorhodopsin captured by a femtosecond x-ray laser. Science *361*, eaat0094. https://doi.org/10.1126/science.aat0094.

Oesterhelt, D., and Hess, B. (1973). Reversible photolysis of the purple complex in the purple membrane of Halobacterium halobium. Eur. J. Biochem. *37*, 316–326. https://doi.org/10.1111/j.1432-1033.1973.tb02990.x.

Ren, Z. (2019). Ultrafast structural changes decomposed from serial crystallographic data. J. Phys. Chem. Lett. *10*, 7148–7163. https://doi.org/10.1021/acs.jpclett.9b02375.

Ren, Z. (2022). Photoinduced isomerization sampling of retinal in bacteriorhodopsin. PNAS Nexus https://doi.org/10.1093/pnasnexus/pgac103.

Stoeckenius, W., Lozier, R.H., and Bogomolni, R.A. (1979). Bacteriorhodopsin and the purple membrane of halobacteria. Biochim. Biophys. Acta BBA - Rev. Bioenerg. *505*, 215–278. https://doi.org/10.1016/0304-4173(79)90006-5.

Swanson, J.M.J., and Simons, J. (2009). Role of charge transfer in the structure and dynamics of the hydrated proton. J. Phys. Chem. B *113*, 5149–5161. https://doi.org/10.1021/jp810652v.

Takahashi, H., Rico, F., Chipot, C., and Scheuring, S. (2018). $\alpha$-helix unwinding as force buffer in spectrins. ACS Nano *12*, 2719–2727. https://doi.org/10.1021/acsnano.7b08973.

Wang, J., and El-Sayed, M.A. (2001). Time-resolved Fourier transform infrared spectroscopy of the polarizable proton continua and the proton pump mechanism of bacteriorhodopsin. Biophys. J. *80*, 961–971. https://doi.org/10.1016/S0006-3495(01)76075-4.

*From:*
*Subject: PNAS Nexus MS# PNASNEXUS-2022-00209R-A Decision Notification*
*Date: September 14, 2022 at 10:45:33 AM CDT*
*To:*
*Reply-To:*

*September 14, 2022*

*Title: "Directional proton conductance in bacteriorhodopsin is driven by concentration gradient, not affinity gradient"*
*Tracking #: PNASNEXUS-2022-00209R-A*
*Authors: Ren*

*Dear Professor Ren,*

*After an initial evaluation of its general suitability for PNAS Nexus, your manuscript was assigned to an editor with expertise in the subject area of your work, who then examined the manuscript and obtained reviews from 2 authorities in the field. These reviews, which appear below, were then considered by the editor, who determined the comments were of sufficient significance to recommend rejection.*

*Because we receive many more manuscripts for consideration than we can accept for publication, critical comments from expert reviewers that cannot be easily and rapidly addressed in a revision are sufficient cause for rejection; our goal to publish important research rapidly prevents us from undertaking multiple rounds of revision and re-review.*

*Although we expect that this outcome is disappointing, we hope that the rationale for our decision process is clear and that you will find the comments of the reviewers helpful when revising your work for submission elsewhere. We thank you for choosing PNAS Nexus as a potential venue for publishing your work and we hope that you'll continue to consider PNAS Nexus for future submissions.*

*Yours,*
*Yannis C. Yortsos*
*PNAS Nexus Interim Editor-in-Chief*

*********************
*Associate Editor Comments:*

*This manuscript was originally submitted and reviewed by 3 scientists in the field, with an initial decision of rejection on the basis of the reviewers' comments. The author appealed the decision, and upon review, the appeal was accepted. The author then submitted a revised version of the work in response to the comments and concerns of the three reviewers, one of whom declined the invitation to review the revised work. Although several other scientists were approached to review the revised manuscript in place of the reviewer who declined, none of them agreed to review the work. The final decision was therefore based on the comments of the 2 original reviewers.*

*Both of the two reviewers commented that the author failed to sufficiently substantiate their claims of a new model of a protein-pumping mechanism for bacteriorhodopsin. The reviewers both submitted extensive comments for the author's consideration.*

*Consequently, it is recommended to reject the revised submission.*

*Reviewer #1 Comments for the Author:*

*Page 5 in Response to peer review*
*despite the clear statements of Lewis et al.: "the retinylidene lysine linkage in bacteriorhodopsin is protonated … the proton is removed before bacteriorhodopsin reaches the 412 nm intermediate".*
*Page 27 in Response to peer review*
*Where is the evidence other than the absorption property discussed above to show that the molecular event of deprotonation from the SB synchronizes with the M formation?*

*The paper by Lewis et al. states that it deprotonates before M, but does not say at what stage of deprotonation. Therefore, this alone is not a strong basis for the present paper. Smith et al. (PNAS 1984) showed a photocycle of L→M deprotonation (Fig. 6 in their paper), which is consistent with Lewis et al. Furthermore, the peak of the C=N stretching vibration of the L intermediate is clearly lower shifted in D2O, indicating the retinal Schiff base of the L intermediate is protonated. In other words, deprotonation of the Schiff base occurs in the L→ M process.*

*Page 8 in Response to peer review*
*If the H-bonding has been lost before M formation, how does the SB deprotonate when M forms?*

*It was suggested to occur through Thr89 which can interact with Schiff base and Asp85 (Figs. 4 and 5 in Nango et al. Science 2016). A similar mechanism was suggested by a computational study (Bondar et al. Photochem. Photobiol. Sci. 2006). So, the disruption of H bond between SB and water does not interfere with the proton transfer.*

*Page 10 in Response to peer review*
*No one is close to demonstrate that a million-fold change in proton affinity occurs simultaneously to each partner of the proton transfer.*

*The elevation of pKa of Asp85 to 13.2 in the M intermediate was explained by theoretical calculation (Saito et al. J. Phys. Chem. B 2022). Not "no one has demonstrated this issue", and it can be explained in the range of the standard model and it is consistent with the conclusion obtained by the FTIR spectroscopy.*

*Page 10 in Response to peer review*
*Reviewer 1 made a factual error when stating that the proton "is covalently bound with RSB (retinal Schiff base)". One must understand that the proton, that is, a hydrogen ion, does not carry an electron, therefore by definition, the bond is not covalent. The proton is only attracted to*

*the lone pair of electrons of the SB nitrogen, a much weaker association compared to a covalent bond.*

*I cannot agree with this opinion. H+ does not carry an electron. But, it shares (not only be attracted) the electrons from a lone pair of the nitrogen atom of the Schiff base upon the protonation leading to the formation of a covalent bond between H and N like the formation of NH4+ from NH3 + H+. If the H+ is so weakly bound, the frequency of N-H vibration must appear in the non-typical region or not observable, but it was observed in the normal N-H or N-D (when the protein in D2O) region indicating it is a typical covalent bond.*

*Page 11 in Response to peer review*
*This analysis based on basic knowledge in chemistry, mechanics, and thermodynamics cannot be easily labeled as "speculative" on the grounds of invisible proton in X-ray data.*

*This is correct, but we need more solid results. If the author believes that the proton is necessarily passed to water by basic chemistry and physics and wants to demonstrate as one of the main findings of this work, QM/MM calculations using the 3D structures which the author obtained should be performed. Otherwise, the possibilities of ultrafast proton dissociation and proton pooling models should be mentioned only briefly in the discussion section, and future work with theoretical studies is needed to substantiate the idea.*

*Page 13 in Response to peer review*
*Reviewer 1 provides a brief but nice recap of the bacteriorhodopsin research and points out the key difficulty that the entire field faces in both spectroscopic and structural studies – mixtures.*

*I mentioned that difficulties due to mixtures exist in time-resolved x-ray crystallography, but not for spectroscopy. In the visible (Chizhov et al. Biophys. J. 1996) and infrared (Lórenz-Fonfría and Kandori J. Am. Chem. Soc. 02009) regions, SVD-based component separation methods are well established, allowing spectra of specific intermediates to be obtained and firmly argued*

*Page 16 in Response to peer review*
*This field of bacteriorhodopsin studies has made a series of mistakes throughout the history. Some of them could be technical errors. But the fundamental ones are due to misunderstanding or disregard of the principles in thermodynamics.*

*I cannot agree that rhodopsin researchers have made a series of mistakes "throughout" the history. While there were some misinterpretations and errors in specific early studies, numerous spectroscopic studies have led to conclusions that are consistent with each other and with subsequent structural studies, and are supported by numerous theoretical computational studies.*
*Also, although the author states that the standard model with large pKa shifts is thermodynamically difficult, it should be kept in mind that many points of his argument is based on the pKa of water in the solution phase, Schiff bases in the dark state, and typical amino acids and that there are theoretical studies that show these values vary drastically in proteins (e.g. Saito et al. J. Phys. Chem. B 2022).*

*Page 16 in Response to peer review*
*How do you know the SB brings the proton with it?*

*As I already mentioned that resonance Raman spectroscopy clearly showed that the Schiff base is protonated up to the L, leading the shift of C=N str. mode upon changing the solvent from H2O to D2O.*

*Page 16 in Response to peer review*
*Therefore, the most red-absorbing O state corresponds to the most acidic inner EC channel throughout the photocycle.*

*The O intermediate is formed by a single photon photocycle meaning only one proton is stored on the extracellular side. If pooling of multiple protons is needed for the pump function as the author demonstrates, a more red-shifted state must appear upon next proton transfer induced by subsequent photon absorption. Such a state, however, was not observed in any study.*

*Page 16 in Response to peer review*
*The so-called initial state could be understood as an established proton pool after light adaptation. This is the all-trans super charged ground state in light (Fig. 5c).*

*If the author considers so-called initial state represents the state in which excess proton is pooled on the extracellular side, in the structural model obtained by the author, the excess proton must be present somewhere from the beginning, and after deprotonation of the Schiff base, there will be an additional excess proton (i.e., two excess protons for each intermediate), but this is not mentioned in the text.*

*Page 36 in Response to peer review*
*However, the long lifetime of K is very certain.*

*There is no disagreement about the lifetime of K. But, its emerging time is still described as "Emerged from J around 10 ps, K lasts... (Line 652)". This is significantly longer than the result of spectroscopy (3-5 ps, e.g. Nuss et al. Chem. Phys. Lett. (1985)). The lifetime of J and the emerging time of K are described as too long throughout the paper.*

*Reviewer #2 Comments for the Author:*

*The present manuscript is well written and the text can be followed easily, except for the motion of the retinal. I have a hard time to follow it. It would be much better if the author would provide pdb files such that a movie of the new mechanism can be visualized.*

*The manuscript, the SI and the review replies are extensive. It is difficult to address all the points. It is also counterproductive that the author is mixing different reviewer questions and not reproducing the questions.*

*Overall, I conclude that the evidence for deprotonation of the Schiff base is not provided in this work. The use of old experiments with inferior setups and samples is outdated. The paper from Anderson on retinal absorption in gas phase from 2005 is also wrong, as reported by the same author in 2010 (see discussion below).*

*I'm starting the response to the reviews:*

*page 1 of the response letter: I agree with the author, his analysis is refreshing and innovative. The same is true for the mechanism. However, it doesn't mean that the older data can be ignored. I noticed that the author is selecting some papers which are rather old to support his findings. For example, the paper from Nuss from 1985. Nowadays, there is much cleaner and better time-resolved data, like [https://doi.org/10.1039/C4CP01826E](https://doi.org/10.1039/C4CP01826E)*
*Why not use this instead?*

*p. 2-3: I agree with the author about the twisted structure. It should be very short lived in the excited state and hard to capture. However, I strongly disagree about the perfect cis and trans. Due to the protein environment a structure with 0 and 180 degrees is nearly impossible to achieve. A twist around 10-30 degrees is possible for a double bond.*

*p. 3: One recurring point in the discussion is the U-turn motion. It is not clear to me. Can the author make a movie or provide pdb files? I don't understand the mechanism of such an atomic motion. Usually, the isomerization is described by torsional angles and not by atom displacements. I recommend to choose appropriate torsional angles and describe their evolution.*

*p. 4: I disagree with the author on the discussion of the proton movement. The example of the hydrogen atoms of the methyl group is not chosen well, because a methyl group is not getting deprotonated. In this manuscript the author claims that the Schiff base is getting deprotonated before the isomerization. I think such a process would be called excited state proton transfer. However, protons cannot be seen. So the question is on which data, which distances exactly are pointing towards a proton transfer from the Schiff base to the water? This is not clearly explained.*

*p.4: Spectroscopy: the author is right that vibrational spectroscopy cannot see protons. But this method is used for visualization of atomic structure. It can however, detect the certain bonds that have characteristic frequency. And this method is very sensitive. Here is a recent review: doi:10.1021/acs.jpcb.6b09222*

*If a proton would move from the Schiff base, it would have been reported by many time resolved IR experiments. This is my main point. I don't see how the author can explain the lack of signal that would indicate a deprotonation.*

*The same is also true for UV/Vis absorption spectroscopy. It doesn't see proton directly but Schiff base deprotonation would induce a large spectral shift to the UV region of the spectrum. This similar to cyanine dyes, where a charge is delocalized over the molecule. If the Schiff base proton will move, there is no charge and therefore no charge delocalization. It will cause a large spectral shift.*

*The author should also note that the discussion of spectral shifts is usually done in nm, which is not a good choice because it is not changing the energy linearly. In electron volts the spectral shift will be much larger upon deprotonation and is therefore a clear indication of Schiff base deprotonation. This is also a strong point which cannot be explained by the newly proposed mechanism in this work.*

*In addition, there is not explanation or evidence for this statement of the author in the manuscript:*

*"This event of SB deprotonation has been mistakenly inferred from the strong blue shift of M state, the lack of deuteration effect in C15=N. stretching frequency in M, and the observed protonation of the carboxylate of Asp85 during L -> M transition." Why is this a mistake? What is the evidence that this is a mistake?*

*The work from Oesterhelt and Hess from 1973 is not a good evidence for little spectral change upon Schiff base deprotonation. The spectroscopic setup (hardware and software) was developed and has improved significantly in the following decades.*

*The work of Anderson from 2005 is not a good reference for neutral retinal Schiff base in gas phase. The same author has reported a mistake in the experimental setup in 2010, in this work:* [http://dx.doi.org/10.1002/anie.200905061](http://dx.doi.org/10.1002/anie.200905061)

*In addition, there is a positive charge on the neighboring nitrogen atom (Scheme 1b in 2005 paper). Hence, the retinal cannot be called neutral (=deprotonated). The charge is just distributed differently.*

*p.6: In ultrafast spectroscopy, the analysis is done using similar tools as used by the author. Hence, it is not clear what exactly is "masked" and why it cannot be see in absorption spectroscopy.*

*p.7-8: The discussion about the assignment of spectroscopic data is also true for the analysis of the crystallographic data by the author. Nowadays, it is common to integrate crystallography, spectroscopy and computer simulation to get a unified picture.*

*p.8-9: The discussion of the energetics and the proton translocation indicates some misunderstanding. I think the consideration from Stockenius is not wrong. The photon energy is converted into chemical energy by photoisomerization, which is a non-radiative process. It means the photon energy is not emitted by rather stored in the isomerized retinal. It happens on the ultrafast time scale, hence there is not enough time to dissipate the energy to the environment. Regarding the translocation of the proton: Is the author suggesting that the same proton is moving in one direction? This is indeed rather unlikely. It would rather resemble a domino effect, where one change propagates through a network.*

*p. 9-10: The model from Jardetzky is not central in the mechanism of bacteriorhodopsin. It is not clear why the author is relying on this old study.*

*While the reviewers have agreed on remaining open questions, it doesn't mean that a suggestion which is not inline with a wide range of repeatable and confirmed spectroscopic studies by various groups provides the answers. As I wrote before, a deprotonated preceeding the isomerization would have different spectroscopic signatures.*

*p. 17: The author writes in the response (but also in the manuscript):*
*One has to offer at least a speculation on how such a proton transfer takes place during L -> M transition at tens of μs. This proton transfer uphill by 11 orders of magnitude does not occur spontaneously. But it does happen in bR. 50 years of research has to attempt some answers.*

*This is quite straightforward to explain. Bacteriorhodopsin is out of equilibrium because of the converted photon energy. Hence, the proton transfer is a response to atomistic rearrangements which are initially triggered by photoisomerization and a vibrationally excited retinal. It doesn't require an ultrafast deprotonation of retinal Schiff base.*

*p. 18: The author describes the function of a channel, while bacteriorhodopsin is a pump. The mechanism of a pump and channel are very different, because the latter requires an opening of a pore.*

*p.22: I disagree with the reply to the question about the hydrogen inertia. The hydrogen forms a covalent bond with the Schiff base nitrogen, it is not a hydrogen bond as suggested by the author!!! It is true that a charge transfer takes place in retinal, but it doesn't mean that a covalent bond is breaking. Even if it would break, the inertia from a proton is very small. So it should be rather the attraction from the lone pair of the oxygen of the water than inertia. But in this case the electrostatic interaction between the deprotonated counterion and the proton should be much higher.*

*p.25: Author writes "Where a SB absorbs was never solidly established" - This reply about the spectroscopic signature of deprotonated Schiff base is incorrect. The signature in the UV region of the spectrum is very well established! The selected references by the author are rather old from experiments that are obsolete or by studies that have been corrected like Anderson 2005 paper (see above).*
